# Phosphorylation of S396 and S400 Promotes Tau Self-Assembly and Favors the Chronic Traumatic Encephalopathy (CTE) Protofilament Fold

**DOI:** 10.64898/2026.05.19.726321

**Authors:** Wyatt C. Powell, Nicholas Yan, Eric Tse, Arthur A. Melo, Nathaniel Sin, Daniel R. Southworth, Jason E. Gestwicki

**Affiliations:** Institute for Neurodegenerative Diseases, University of California, San Francisco, San Francisco, California 94158, United States; Department of Biochemistry & Biophysics, University of California, San Francisco, San Francisco, California 94158, United States; Department of Pharmaceutical Chemistry, University of California, San Francisco, San Francisco, California 94158, United States

## Abstract

Tau hyperphosphorylation is linked to tauopathy aggregates, but the effects of individual sites on tau assembly remain unclear. Here, we used protein semisynthesis to generate defined Tau(297–407) proteoforms containing specific combinations of phosphorylation within the PHF1 epitope. Spontaneous aggregation revealed that phosphorylation of S396, and S400 to a lesser extent, promoted nucleation, while phosphorylation of T403 and S404 suppressed assembly. A similar reactivity trend of pS396 > pS400 > WT > pT403 > pS404 was observed for seeded assembly, and all proteoforms, even anti-aggregation ones, were incorporated. The fibrils have similar thermodynamic stability, suggesting that phosphorylation selectively tunes reactivity and not thermodynamics. Cryo-EM revealed that pS400 produces a chronic traumatic encephalopathy (CTE) protofilament conformation. Strikingly, one of the structures observed in the pS400 sample appeared to capture a secondary nucleation step. Together, these studies reveal the importance of positional effects of phosphorylation on tau self-assembly.

## Introduction

Tauopathies, including Alzheimer’s disease (AD) and chronic traumatic encephalopathy (CTE), are neurodegenerative diseases characterized by the accumulation of tau pathology. Tau proteins within these deposits are extensively modified by post-translational modifications (PTMs), with hyperphosphorylation as a notable hallmark. The close link between tau hyper-phosphorylation and disease progression has led to the hypothesis that these modifications might play active roles, such as driving tau nucleation and/or promoting fibril formation.^1–6^ More recently, a potential role for phosphorylation in shaping fibril structure has also been suggested; for example, specific modification signatures have been identified in material isolated from patients with different tauopathies and these patterns are proposed to play an important role in the maturation of the distinct fibril structures observed in these diseases.^4,6–11^ However, it has been difficult to rigorously test these hypotheses, in part, because tau is subject to a high number of PTMs, with modification stoichiometry up to 12 phosphorylation sites per molecule across 55 residues, yielding a staggering number of theoretical proteoforms.^12^ Moreover, traditional methods to add PTMs to tau, such as incubation with disease-associated kinases, yields heterogeneous mixtures, making it hard to deconvolute the contributions of individual modifications. Alternatively, the use of phosphorylation-mimicking mutations (*i.e.,* Asp) gives site-selective information, but fails to capture the true chemical nature of native phosphorylation.^13^

A long-standing literature points to tau phosphorylation as a driver of its nucleation. Full-length tau (such as the 2N4R variant) does not readily self-assemble at supersaturation, whereas phosphorylation of this protein is sufficient to promote its assembly.^14–30^ Unlike the full length protein, the protease-resistant ordered core of paired helical filaments (PHFs), composed of tau(297-391), readily assembles into AD-like fibrils at high concentrations.^15,31–38^ This observation is important because the protease-sensitive, disordered regions, termed the “fuzzy coat”, are highly cationic and they contain many of the disease-associated phosphorylation sites. Thus, it has been hypothesized that phosphorylation overcomes charge repulsion to initiate assembly via a polyampholyte-like mechanism. In other words, the disordered anionic charges in the fuzzy coat are thought to behave as a polyelectrolyte brush,^39^ and act in a way that is reminiscent of how polyanions induce tau assembly.^40–48^ Indeed, phosphorylation and polyanions have competing effects on the propensity for assembly.^24,49–51^ These results suggest that tau assemblies are driven by electrostatic interactions with disordered multivalent anions (*e.g.* fuzzy coat phosphorylation or polyanions), with subsequent contributions from the hydrophobic interactions of the ordered core. However, this simplistic model does not seem to fully explain the role of phosphorylation. For example, histochemical studies also suggest that neurofibrillary tangles (NFTs) are not always hyper-phosphorylated,^52^ suggesting that the site of phosphorylation, and not just its levels, might be important. Moreover, tau phosphorylation occurs in healthy adults and infants, and not just tauopathy patients, further suggesting that there might be some positional effects that remain mysterious.^53^

Here, we asked whether native phosphorylation of the PHF-1 epitope has positional effects on assembly kinetics, aggregate stability, or fibril structure. We used chemical protein synthesis to prepare a library of phosphorylated Tau(297-407) proteoforms containing all combinations of zero to three modifications from the disease-associated PHF1 motif (**Fig 1**). Using inducer-free assembly conditions, we revealed strong position-dependent effects on assembly kinetics: pS396 and pS400 promote nucleation, whereas pT403 and pS404 are strongly inhibitory. This general reactivity trend of pS396 > pS400 > wild type > pT403 > pS404 is also retained in seeded assembly conditions, further demonstrating the pro- and anti-aggregation effects in that model. We found that all of the fibrils had similar thermodynamic stability, as revealed by equilibrium solubility and chemical denaturation assays. Thus, position-selective effects on tau assembly kinetics occurred independently of any change in thermodynamic stability. Finally, we used cryo-EM to study the structure of the pS400 and pS396/pT403 fibrils, revealing that the pS400 protofilaments adopt a conformation identical to the CTE fold and similar to those in AD. We also identified a hetero-dimeric structure in the pS400 sample that appears to capture a proposed secondary nucleation step in tau self-assembly. Together, these findings demonstrate that specific phosphorylation sites have remarkably variable impacts on folding propensity, providing insight for the development of precision medicine and diagnostics targeting tau’s post-translational modifications.

**Figure 1.**
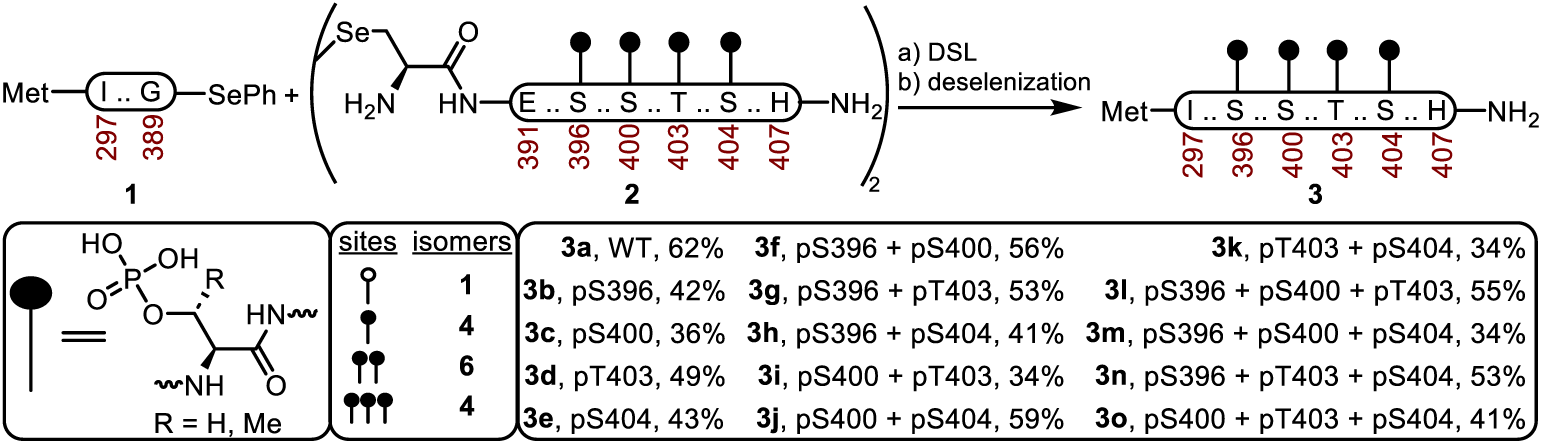
Synthetic strategy for the creation of a library of phosphorylated tau proteins. Conditions for one pot diselenide-selenoester-ligation (DSL)-deselenization. a) Ligation: 6 M guandinium hydrochloride (Gnd·HCl), 0.1 M sodium phosphate pH 6.8, 1 h, 23 ⁰C then diphenyl diselenide extraction with hexanes. b) Deselenization: adjust to 100 mM tris(2-carboxyethyl)phosphine hydrochloride (TCEP·HCl), 50 mM dithiothreitol (DTT) pH 6.8 and mix at 23 ⁰C for 24 h. The isolated yields of each proteoform are reported after high-performance liquid chromatography purification. See **Supporting Information S1-3** for details and analytical characterization.

## Results

### Semi-synthesis of a chemically defined library of pTau(297-407) proteins

It has been shown that Tau(297-407) with four mutations: S396D S400D T403D S404D, hereby referred to as Tau(297-407)-4D, assembles into AD-like structures at supersaturation, with the fuzzy coat phosphomimetic mutations as facilitators its assembly.^25,37^ Thus, we considered this system to be a useful one in which to address the key question of positional effects and to differentiate any impact of true phosphorylation from pseudo-phosphorylation. To test these ideas, we prepared chemically defined, discrete Tau(297-407) proteoforms using solid-phase peptide synthesis of phosphorylated PHF1 epitope residues, combined with intein-mediated, protein semi-synthesis. Briefly, the synthesis involves ligation of phenyl-selenoester **1** and a synthetic phosphorylated peptide diselenide **2**, followed by radical deselenization of selenocysteine into native alanine 390 (**Figure 1**).^54^ The production of Tau(297-389)-SePh **1** was accomplished upon a bacterial expression as a fusion protein with a C-terminal MxeGyrA intein, which was spliced to an acyl hydrazide, and then converted to the phenyl selenoester **1** through an acyl pyrazole intermediate (**Supplemental S1**).^55–60^ The diselenide peptides **2** containing the phosphorylated residues were synthesized as C-terminal amides using SPPS on Rink-amide resin (**Supplemental S2**). These fragments represent each PHF phosphorylation site separately, as well as combinations of two or three phosphorylation sites. Our goal was to identify the effects of individual phosphorylations, as well as revealing any additive or synergistic effects. The segments were then merged using the diselenide-selenoester ligation (DSL)-deselenization, a one-pot protocol, which, upon completion of the ligation, is treated with tris(2-carboxyethyl)phosphine (TCEP) to induce selective deselenization of A390Sec in the presence of an unprotected cysteine residue (C322), furnishing the phosphorylated Tau(297-407) **3** variants in 34-62% isolated yield.^61–63^ The semi-synthesis method successfully yielded 15-30 mg of the Tau(297-407) library, and high-resolution mass spectra demonstrate excellent purity (>95%) (**Supplemental S3**). We also expressed and purified the pseudo-phosphorylated protein, Tau(297-407)-4D, to allow direct comparisons. Two phosphorylation sites promote fibril formation, while the others limit aggregation.

We first measured the ability of the phosphorylated Tau(297-407) proteins to form fibrils under quiescent, inducer-free conditions. These experiments were designed to reveal positional effects on the primary nucleation barrier. Accordingly, the assembly rate of the various proteoforms was compared using the time at half saturation (t_1/2_) from thioflavin T kinetic traces (**Supplemental S4**). Strikingly, we found that monophosphorylated pS396 (t_1/2_ ∼2 days) and, to a lesser extent, pS400 (t_1/2_ ∼3.5 days), promoted assembly as compared to the WT, non-phosphorylated protein (t_1/2_ ∼5 d), while the pT403 and pS404 proteoforms did not assemble under these conditions (t_1/2_ > 7 days; **Figure 2**). This result shows that pS396 is a strong nucleator, while pS400 is a mild promoter and both pT403 and pS404 inhibit cofactor-free assembly. When we tested each of the combinations of phosphorylation sites to probe whether these relative activities would be additive, we found that the combination of pS396 with inhibitory phosphorylations (di-phosphorylated pS396 + pT403/pS404 (t_1/2_ ∼2.2 days) behaves similar to WT. Likewise, the combination of pS396 + pS400 (t_1/2_ ∼1.5 days) is more aggregation prone than either alone. Thus, the effects from individual phosphorylations seem to combine in an approximately linear way (*e.g,* pro- and anti-aggregation marks cancel each other, while two pro-aggregation marks are roughly additive). Importantly, the di- and tri-phosphorylated proteoforms lacking pS396 failed to assemble, thereby indicating that pS396 is uniquely able to overcome the suppressive effects of pT403 and pS404.

**Figure 2.**
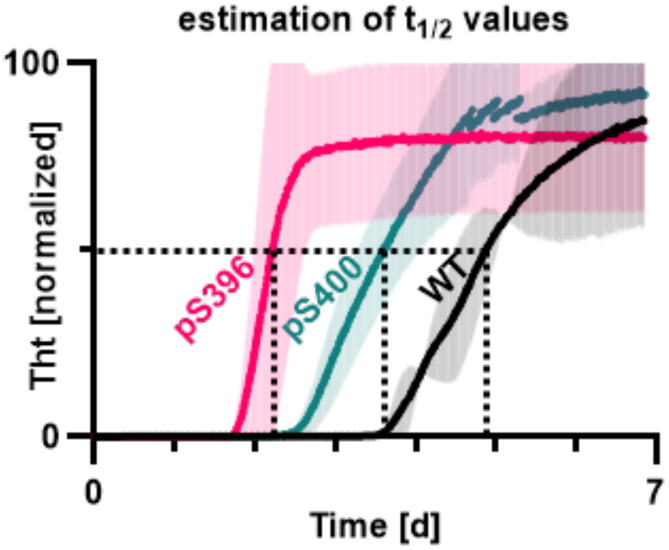
Phosphorylation of pS396 or pS400 promotes aggregation, while pT403 or pS404 prevent aggregation. Semi-synthetic proteins were assembled under quiescent conditions without co-factors and fibril formation estimated from thiofavin T (ThT) fluorescence (see Methods). The cross sections of the dashed lines represent the t_1/2_ values, Reactions were performed in biological triplicates, with 6 technical replicates per reaction. Representative experiments are shown. The Tau(297-407) pT403 and pS404 proteins failed to form fibrils under these conditions (t_1/2_ > 7 days).

The assembly of tau fibrils can be catalyzed by the addition of fibril fragments (termed “seeds”). These seeds are thought to template recruitment of tau monomers into the growing fibril. Hyperphosphorylation has been reported to play a role in maintaining the autocatalytic, prion-like properties of tau fibrils; for example, treatment with phosphatases reduces the seeding activity of PHFs.^16,64–66^ Therefore, we next used our Tau(297-407) library to examine how phosphorylation impacts the seeded assembly barrier. In these experiments, we created seeds from either WT Tau(297-407) or Tau(297-407) containing the two, pro-aggregation marks (pS396 and pS400) and then used those seeds to accelerate fibril formation for the phosphorylated Tau(297-407) library. The results indicate that all proteoforms undergo seeded assembly (**Figure 3b, c**), demonstrating that anti-aggregation monomers are not able to fully suppress seeded assembly. Another important observation is that the WT seeds had generally lower activity than pS396 + pS400 seeds, further supporting an important role of the pS396 and pS400 sites (**Figure 3b,c, Supplemental S5**). Overall, the relative seeding kinetics generally followed the same trend observed in the primary nucleation reactions: pS396 > pS400 > WT > pT403 > pS404 (**Figure 3b,c**). For example, as monomers, pS396 and Tau(297-407)-4D had the greatest propensity to assemble, while WT, pS400, and pT403 had similar seeding kinetics, and pS404 clearly has inhibitory effects. Diphosphorylated and triphosphorylated proteoforms containing pS396 had assembly kinetics similar to pS396 alone, suggesting that pS396 has a dominant effect on the seeded assembly barrier. Moreover, pT403 + pS404 clearly shows the additive nature of the modifications, as both sites seemed to contribute to relative suppression. Thus, the positional effects of PHF1 phosphorylations on seeded barriers tended to follow those observed in primary nucleation.

**Figure 3.**
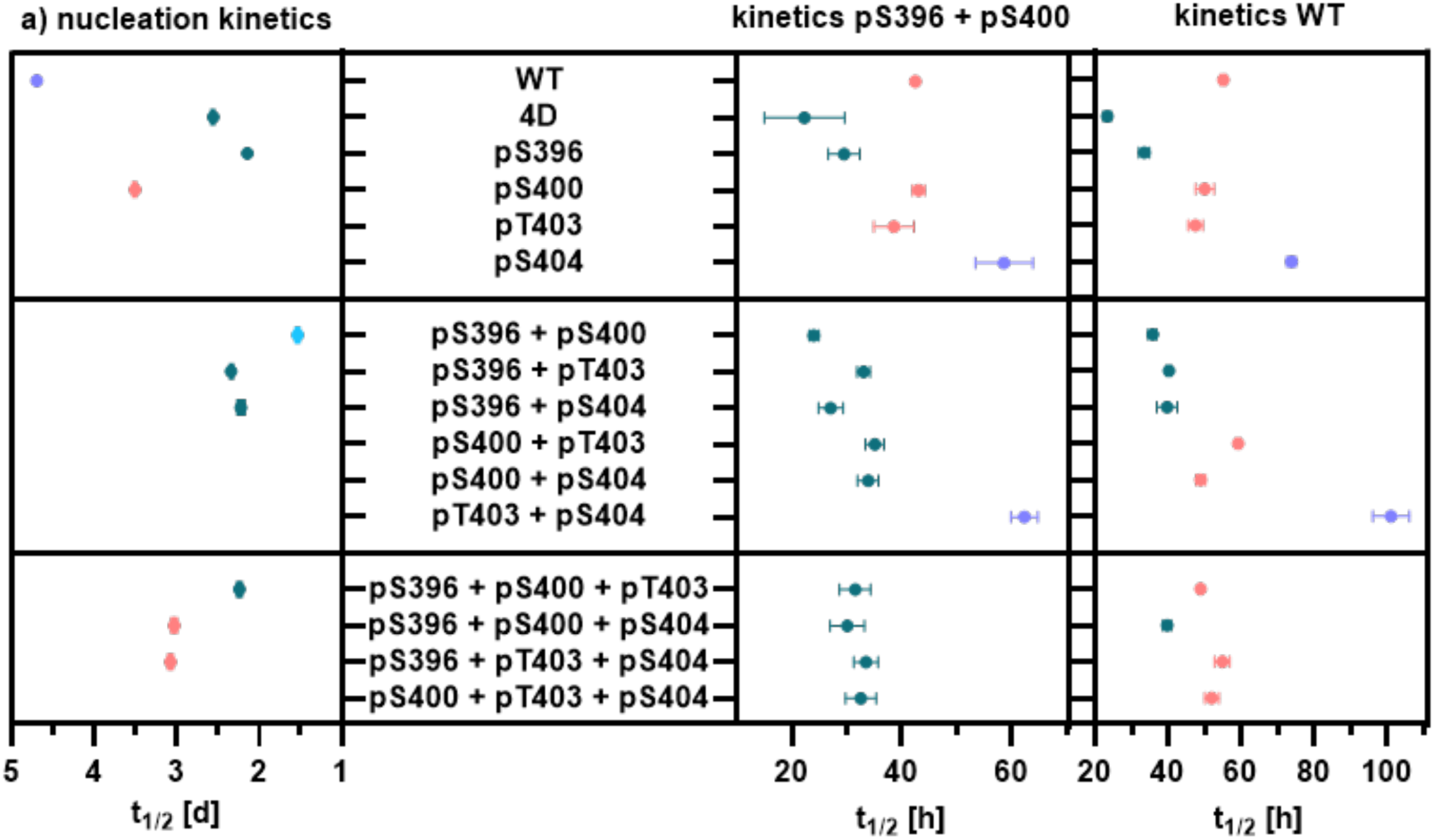
Kinetics of primary nucleation and seeded assembly reveals strong positional effects and approximately additive impacts on aggregation kinetics. a) Primary nucleation kinetics. Conditions: 200 µM Tau(297-407), 200 mM potassium citrate tribasic, 10 mM DTT, 3 µM thioflavin T, 10 mM potassium phosphate pH 7.4, 37 ⁰C, 7 days, no mixing (see supplemental S4 for details). b) Seeding kinetics with pS396 + pS400 seeds or c) wild type (WT) seeds. The buffer conditions are the same as in a), but with 50 µM Tau(297-407) proteoforms and 5 µM Tau(297-407) seeds. The reactions were performed in triplicate, with 6 technical replicates per reaction. The average t_1/2_ values were obtained from the normalized assembly kinetics (see supplemental S5), and the error bars represent the standard deviation. Values are color coded to indicate relative position compared to WT. 4D = Tau(207-407)-4D (see Methods).

Many studies suggest that hyperphosphorylation stabilizes tau filaments.^15,67–72^ To test for potential, positional effects of PHF1 phosphorylation on fibril stability, we collected the fibrils from the previous experiments and measured their properties using multiple assays. Firstly, we employed sedimentation assays to estimate equilibrium solubility, which is directly related to stability.^73–76^ In these experiments, the free monomer remaining after direct nucleation reactions (C_sat_) was measured to be in a relatively broad range (∼25 to 100 µM), and there were not significant trends between the proteoforms (**Figure 4a**). This variability is likely due to the fact that tau fibrils have non-equilibrium and gel-like characteristics under these conditions,^77^ such that minor differences in collection time or other factors might increase variability. Therefore, we move to measuring the solubility upon seeded assembly, with the goal of obtaining less variability. Indeed, the measured C_sat_ values were now in a narrower range (2 to 7 µM). These results showed that phosphorylation had no significant effect on the seeded equilibrium solubility, yielding similar C_sat_ values across all proteoforms, regardless of whether they were seeded with pS396/pS400 or WT (**Figure 4b-c, Supplemental S6**). While this result suggested that phosphorylation only impacts kinetics, but not fibril stability, we wanted to use an independent measurement. Accordingly, we measured the relative stability of the fibrils using chemical denaturation, which reflects thermodynamic stability. These studies showed that the phosphorylated proteoforms all produce fibrils that have similar stability (**Supplemental S7**). Therefore, we conclude that phosphorylation of the PHF-1 epitope does not significantly alter the thermodynamic stability of tau fibrils. Thus, phosphorylation in this region seems to preferentially impact tau fibril kinetics.

**Figure 4.**
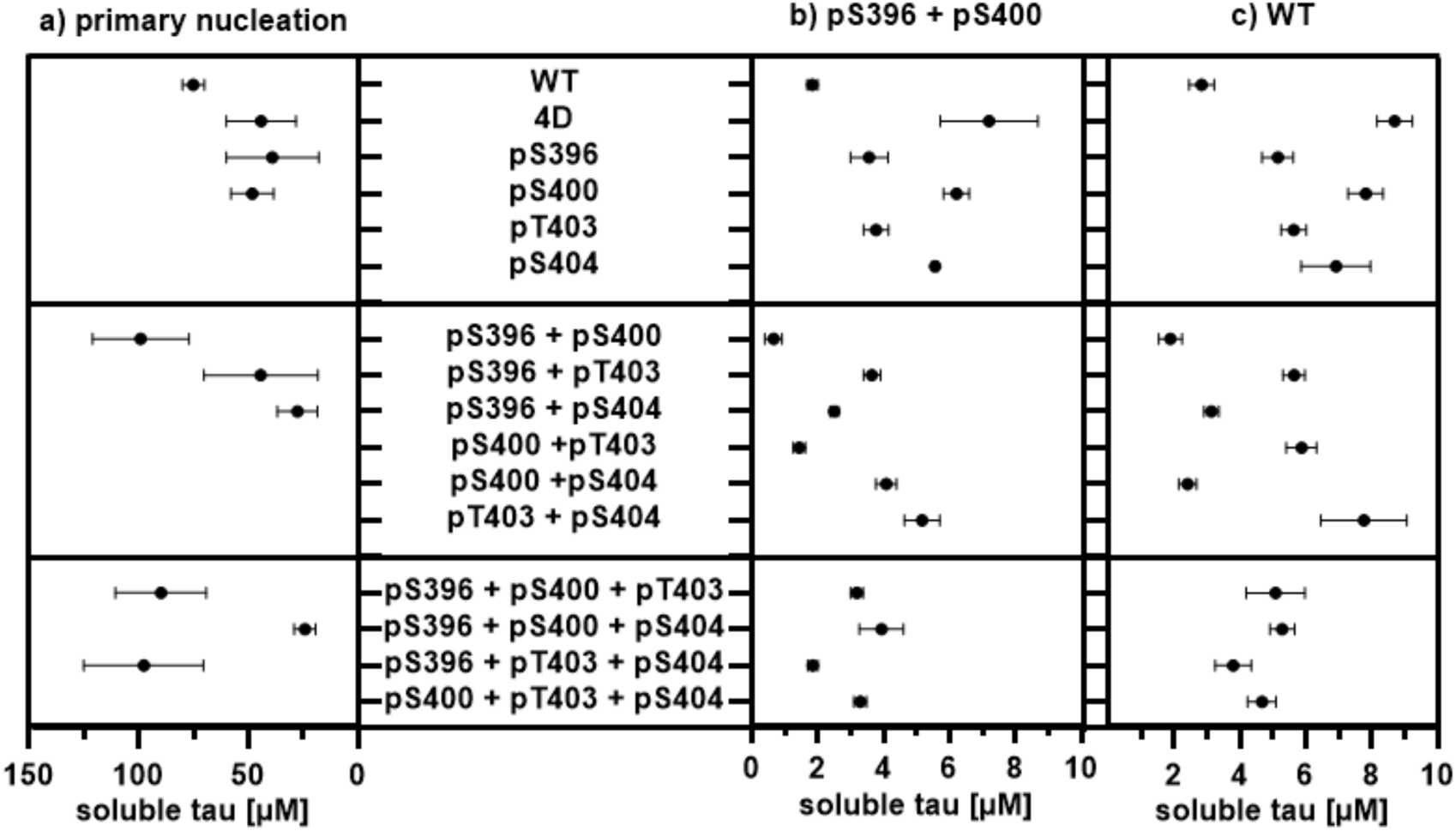
Equilibrium solubility indicates that the phosphorylated proteoforms exhibit similar thermodynamic stability. a) Equilibrium solubility upon completion of primary nucleation (see Figure 3a for conditions). b) Equilibrium solubility upon completion of seeded assembly with pS396 + pS400 seeds or c) WT seeds (see Figure 3b, c for conditions). Determination of C_sat_: ultracentrifugation at 100 k x g for 60 min, and then the monomer concentration in the supernatant was determined using high-performance-liquid-chromatography at 214 nm (see Supplemental S*6* for detailed experimental protocols).

### Cryo-EM indicates that phosphorylation accelerates thermodynamically driven rearrangements

A large body of literature suggests that phosphorylation contributes to tau fibril structure; for example, enzymatic dephosphorylation alters fibril morphology.^6, 55–60^ However, it has been challenging to identify position-specific effects due to heterogeneity. To assess the structural impact of PHF-1 phosphorylation, we solved the cryo-EM structures of Tau(297-407) fibrils. In preliminary studies, the pS396 + pT403 and pS400 samples exhibited helical features that would be promising for structural determination, so we moved forward with these two examples, while pS396 + pS404 filaments were predominantly straight and pS396 + pS400 + pT403 exhibited worm-like morphology that was difficult to resolve (**Supplemental S8**). To obtain structures closer to thermodynamic equilibrium,^77^ the samples were aged for an additional 4 weeks at 23 °C (after the 7 days of initial assembly) before preparing the cryo-EM grids. From 2D classification, approximately 21% of the pS396 + pT403 filaments contained a helical twist (**Figure 5**). In this sample, the cross-section is composed of two identical protofilaments arranged in an approximate C2_1_ screw symmetry. In contrast, the pS400 sample contained two resolvable polymorphs (termed pS400-1 and pS400-2) with similar abundances (33% and 28%, respectively). The pS400-1 example was composed of two identical protofilaments with an asymmetrical interface, and pS400-2 contained two non-identical protofilaments. Moreover, pS400-2 is distinct because it has a noticeably asymmetrical interface. This level of asymmetry is uncommon, but it has been observed in brain derived alpha synuclein (MSA) and recombinant tau filaments.^78,79^ We next used atomic modeling to study these structures in more detail. For the pS396 + pT403 fibrils, two J-shaped protofilaments were observed, with structured regions comprising residues S305 through R379 (**Figure 6**). These protofilaments contain a “head” region connected through a “neck” to an elongated “tail” and the two protofilaments were arranged with contacts along the tails (**Figure 6**). To understand whether these protofilaments resemble disease-associated structures, we aligned the tails with protofilaments from AD and CTE.^80,81^ This comparison revealed that the tail regions are superimposable, whereas the head regions exhibit different levels of relative curvature or bending (**Figure 7a**). Specifically, the head region of the pS396 + pT403 protofilament adopts an intermediate level of curvature: more closed/curled than CTE, but less curled than AD. A closer examination reveals that this conformation is caused by the solvent exposure of S356 and L357 side chains in the neck region, as S356 and L357 are buried within the core of the pS396 + pT403 protofilament, but one of these residues is solvent-exposed in AD, and the other residue is exposed in CTE. Another interesting aspect is that residues S324-H330 in the pS396-pT403 fold adopt the same conformation as the AD and CTE folds, whereas in the pseudo-phosphorylated, Tau(297-407)-4D filaments, these residues show less similarity to the disease conformations (**Figure 7b**). Thus, while pS396 + pT403 does not form the native CTE or AD conformations under these conditions, it yields protofilaments that more closely resemble disease-specific conformations than Tau(297-407)-4D.

**Figure 5.**
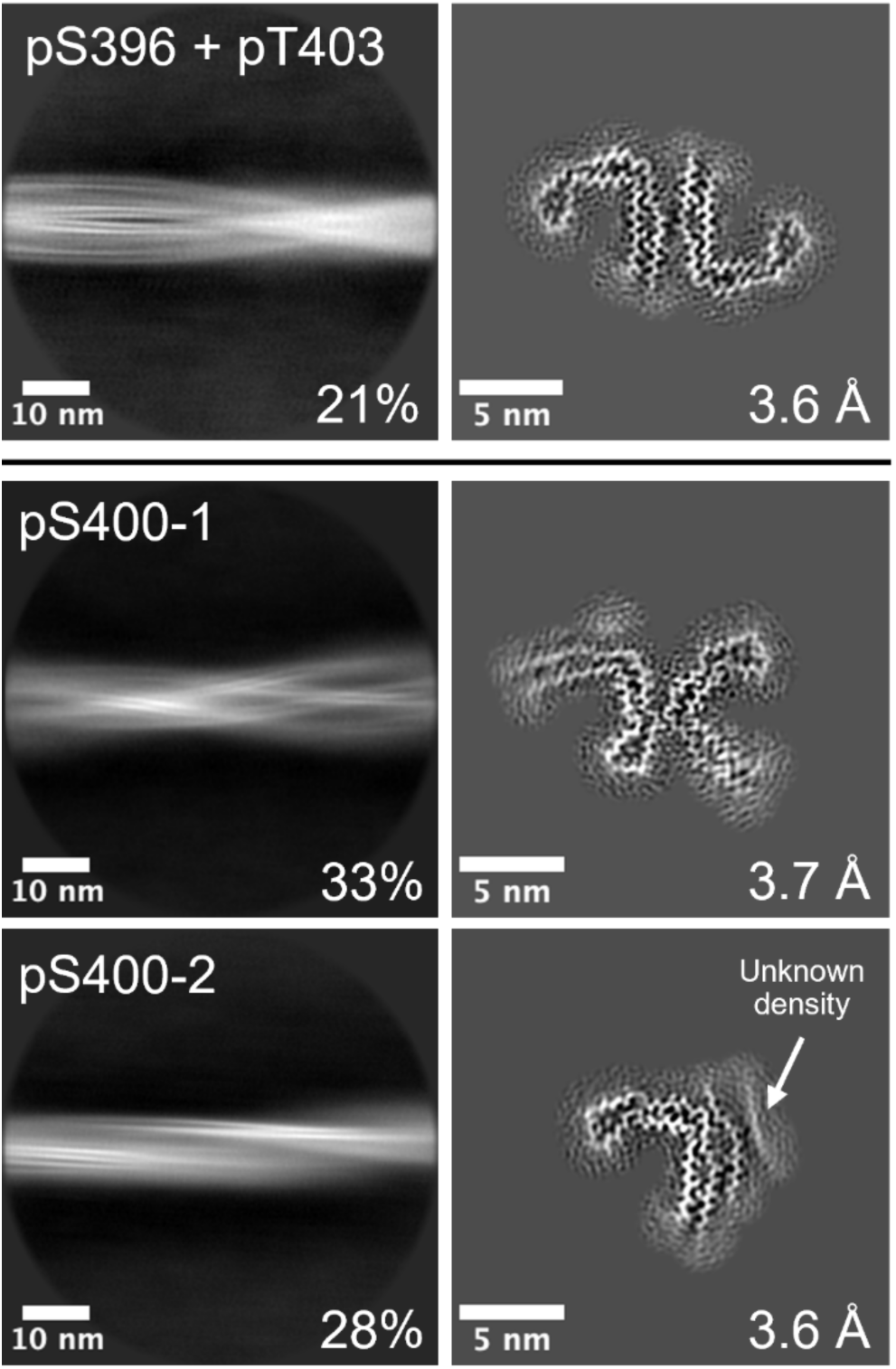
Cryo-EM of select tau fibrils. Cryo-EM grids were prepared after incubation for 1 week at 37 °C, followed by an additional 4 weeks at 23 °C (35 days total). The 2D classifications reveal one twisted, well-defined polymorph of pS396 + pT403, and two polymorphs for pS400. The polymorph totals were determined by 2D classification, and the remaining filaments in each dataset were non-twisting or difficult to resolve. An unknown density is observed by the shorter protofilament in pS400-2.

**Figure 6.**
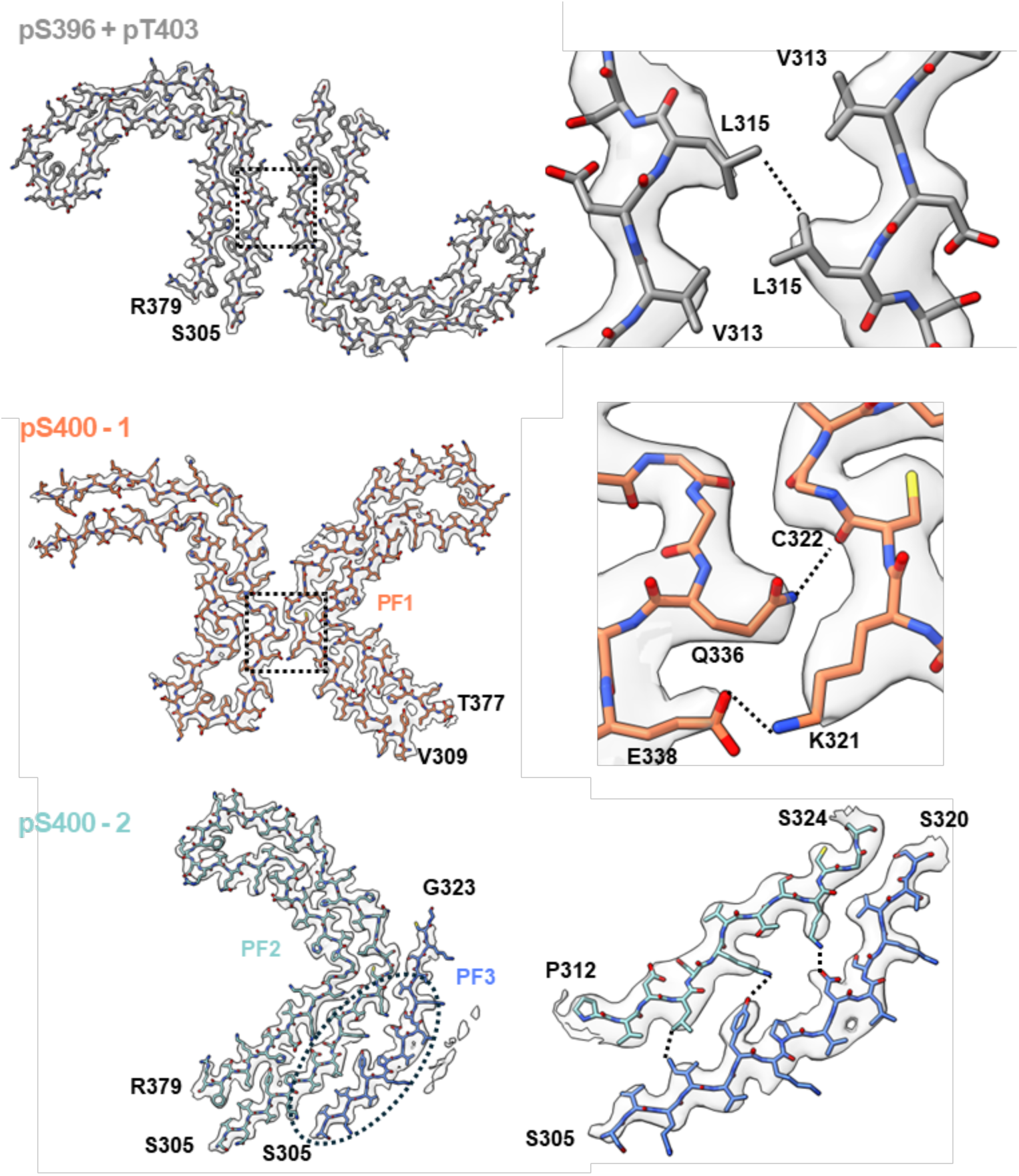
Modelling and structural analysis of the pTau filaments. Atomic models of the filament that have been refined to the density, and close up views of the protofilament interfaces. The pS396 + pT403 polymorph contains two identical protofilaments, and it has an approximate C2_1_ screw symmetry. The pS400-1 polymorph consists of two identical protofilaments. The pS400-2 polymorph contains two non-identical protofilaments, and the protofilaments are arranged with an asymmetrical interface, in a parallel fashion.

**Figure 7.**
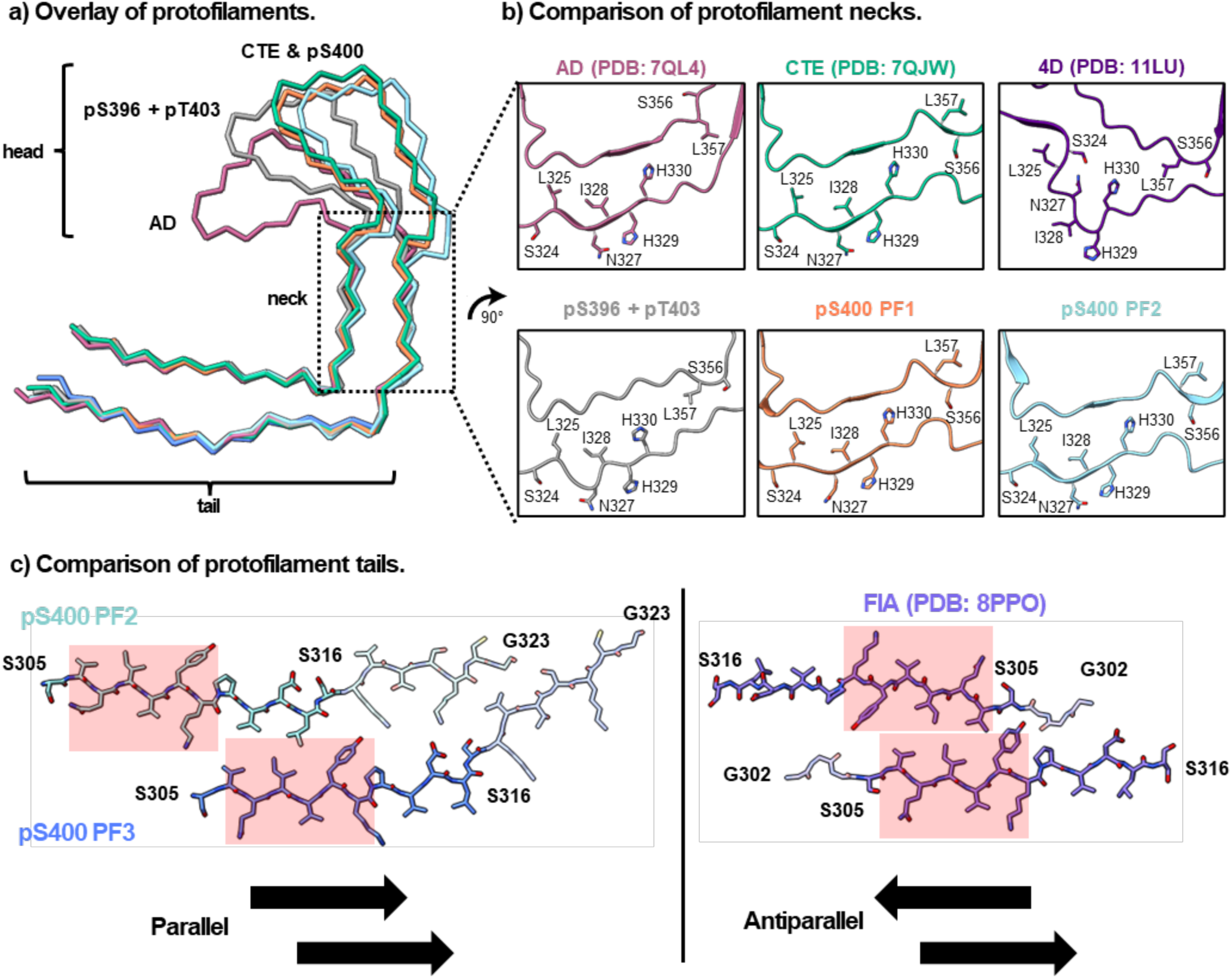
Structural analysis of the pTau filaments. a) The protofilaments are aligned with the AD and CTE folds. The protofilament head and tail regions are identical, while the conformational differences are localized to the neck region (S324-H330, S356-L357). b) The protofilament neck regions contain the conformational differences. The only difference between AD and CTE is the solvent exposure of L357 and S356. A tau(297-407)-4D fold is also shown for comparison, indicating conformational differences in S324-H330 and suggesting that phosphorylated tau fibrils are a more disease-specific conformation. c) The pS400-2 protofilament tail region interface is compared with the first intermediate amyloid (FIA). The solid color residues (S305-S316) are seen in both structures. The ^306^VQIVYK^311^ motifs are highlighted in red boxes. Additionally, the pS400-2 protofilaments adopt a parallel conformation, while FIA has an antiparallel conformation.

Next, we turned our attention to the pS400-1 fibril conformation. Atomic modelling of the major pS400-1 polymorph revealed that it contains two protofilaments (termed PF1), both consisting of residues V309 to T377, although one protofilament tail is less well-resolved than the other tail (**Figure 6**). These protofilaments have an asymmetrical protofilament interface, akin to the straight filaments from AD brains.^80^ This polymorph is nearly identical to a “late intermediate amyloid” (LIA-3, PDB 8Q9F) that was observed during time-resolved cryo-EM of tau(297-391) filaments under CTE-fold formation conditions, with an overall Cα RMSD of 1.2 Å.^79^ Strikingly, these protofilaments are superimposable with the CTE fold (**Figure 7a**). In the neck region, the position of sidechains in S356 and L357 are aligned with the CTE fold, and good alignment can also be observed in residues 324 through 330 (**Figure 7b**). Thus, under these conditions, phosphorylation at pS400 is sufficient to produce a disease-relevant tau fibril fold.

The second pS400-2 polymorph also contains a CTE protofilament (termed PF2) with a similar arrangement to that of PF1 in pS400-1, but paired with a second non-equivalent protofilament (termed PF3). The PF2 filament is again nearly identical to those observed in CTE (**Figure 7b**). In contrast, PF3 consists of two discontinuous layers that are likely connected by an unresolved chain. Atomic modelling revealed that only the first layer (closest to the PF2 filament) is resolved at high resolution, comprising residues S305-G323 (**Figure 7c**), which is superimposable on the tail region of the AD and CTE protofilaments. The protofilament interface between PF2 and PF3 is arranged in a parallel conformation, as opposed to the more common antiparallel steric zippers, including the first intermediate amyloid (termed FIA), which was previously reported from self-assembly of Tau(297-391).^79,82^ Notably, both PF2 and PF3 contain the VQIVYK motif, which has long been believed to be important for assembly.^83^ The VQIVYK motifs are

displaced in pS400-2 such that only the PF3 VQIVYK is in the resolved contact region, whereas both VQIVYK motifs in FIA form the core of the interface with each other. In pS400-2, the interface therefore involves non-identical residues, which is unusual in reported tau fibril structures. We speculate that pS400-2 might represent a long-hypothesized, secondary nucleation step (see Discussion).

## Discussion

Aberrant tau hyperphosphorylation is widely associated with the pathology of tauopathies.^1–6^ Despite decades of research, numerous questions remain regarding how (or if) individual phosphorylation events contribute to tau aggregation, stability, and filament polymorphism. This uncertainty partially arises from the proteoform heterogeneity of tau, which is hard to control through traditional methods. Consequently, it has been difficult to decipher which modifications are mechanistically important and which are merely correlated with pathology or even potentially protective of disease. Moreover, there are likely some modifications that contribute to bulk physical properties of tau (*e.g.* overall charge), whereas others play molecularly defined, discrete roles.^84^ Here, we take advantage of advances in intein splicing technology and solid phase peptide synthesis to create chemically defined libraires of phosphorylated tau proteoforms. When we investigated the positional effects of phosphorylation at the PHF-1 epitope (S396, S400, T403, S404) using this semi-synthetic library of Tau(297-407) proteoforms, we were able to isolate how individual modifications (and combinations) regulate the primary assembly barrier, the seeded reaction barrier, the thermodynamic stability of resulting fibrils, and how a subset of the phosphorylation sites contribute to the structure of tau filaments.

### Phosphorylation Controls Reactivity, Not Fibril Stability

A central finding of this study is that PHF-1 phosphorylation exerts position-dependent effects on assembly kinetics, without measurable impact on thermodynamic stability of the fibrils. Monophosphorylation at S396, and to a lesser extent S400, markedly lowers the primary nucleation barrier, whereas phosphorylation at T403 and S404 suppresses assembly (see **Figure 2 and 3**). When combined into di- and tri-phosphorylated proteins, these effects are approximately additive, with pS396 acting as a dominant promoter, which overcomes the inhibitory influence of pT403 and pS404. Likewise, pS400, which has a more modest pro-aggregation effect, can be suppressed by pT403 and pS404. Similar findings were observed in both primary nucleation and seeded reactions, suggesting that PHF-1 phosphorylation plays important roles in both. Despite this dramatic impact on aggregation kinetics, equilibrium solubility measurements and chemical denaturation assays revealed no significant differences in the stability of the resulting fibrils. Thus, we conclude that PHF1 phosphorylation strongly modulates the rate at which tau assembles, but it does not change the free energy of the aggregated state once formed. These data argue against models in which PHF-1 phosphorylation stabilizes fibrils thermodynamically,^15,67–72^ and instead support a model in which site-specific phosphorylation primarily acts on the kinetic barriers governing assembly. In cells, it is possible that PHF-1 phosphorylation contributes to binding of polyanions or other partners, which does impact fibril stability, but these modifications alone do not directly contribute.

### Similarities and Differences Between Fuzzy Coat Phosphorylation and Polyanionic Inducers

We do not yet understand the mechanism by which phosphorylation at pS396 or pS400 promotes tau aggregation or why pT403 and pS404 counteract this process. Because these residues are not present in the structured core that is visible by cryo-EM, we can only speculate that intra- or inter-molecular contacts along the assembly pathway might be involved. For example, one possibility is that pS396 could make polar contacts with lysine sidechains in the precursor to the CTE fold, while the nearby pT403 might promote an alternative contact that is less favorable or even off-pathway. The pS396 modification seems able to overcome the anti-aggregation marks at pT403 and pS404, suggesting that the pS396 contact is likely preferred. While the molecular details are not yet clear, it is important to note that our findings are inconsistent with a purely property-based model for contributions of PHF1 modifications, in which phosphorylation at any site simply acts to generally suppress cationic charge repulsion. Rather, there are strong positional effects that cannot be readily explained by bulk electrostatic properties.

In this way, there are parallels between the pro-aggregation effects of PHF1 phosphorylation and polyanionic cofactors. Both seem to act as negatively charged species that transiently associate with cationic regions of tau.^40,85,86^ Also, we observe that the pattern of anionic modifications has an important effect on raising (pT403 and pS404) or lowering (pS396 and pS400) the nucleation barrier, which is similar to how the sulfation/phosphorylation/carboxylation patterns on polyanionic inducers modulate their relative effect on tau self-assembly kinetics.^44,87^ However, there are important differences as well. For example, unlike polyanions, PHF-1 phosphorylation does not thermodynamically stabilize fibrils or serve as a stoichiometrically associated element of the filament structure.^41,47,48,88–93^ Thus, phosphorylation and polyanionic cofactors are similar in some respects, but they diverge in their contributions to fibril stability.

### Preformed Fibrils Catalyze Incorporation of Inhibitory Modifications into the Filaments

The inhibitory effects of pT403 and pS404 on primary nucleation raise questions about their abundance in AD filaments. Why would these inhibitory phosphorylation sites be incorporated into post-mortem fibrils? We found that pT403- and pS404-containing proteoforms resist direct nucleation but that they remain substrates for elongation in seeding reactions (see **Figure 3**). This observation might help define roles of opposing phosphorylation effects in the classic, two-stage model of tau aggregation.^4,6–10^ In the first stage, we envision that a small subset of kinetically favorable tau proteoforms —most notably those containing pS396— could initiate fibril formation. In the second stage, templated growth enables the incorporation of other tau proteoforms, even strongly anti-aggregation ones. It is possible that cells might even attempt to add pT403 and pS404 modifications in an attempt to prevent tau’s aggregation, but that strongly pro-aggregation sites, such as pS396, can overwhelm this protective mechanism. Under this potential framework, pS396 and pS400 represent causative nucleator modification sites, whereas pT403 and pS404 represent correlative PTMs that accumulate during fibril growth, but actively inhibit pathology. The balance and timing of these phosphorylation events would, therefore, ultimately shape tau aggregation processes.

### Phosphorylation Accelerates Folding Towards Disease-Relevant Conformations

There has been intense interest in developing models that reproducibly create disease-relevant tau folds over the last several decades. Here, we find that pS400 is sufficient in Tau(297-407) to produce fibrils with the CTE protofilament under quiescent, inducer-free conditions (**Figure 7a**). Filaments assembled from authentically phosphorylated tau adopt protofilament conformations that more closely resemble AD and CTE folds than those formed by phosphomimetic mutants (see **Figure 7b**). Consistent with the literature, this finding may suggest that the location of phosphorylation and neighboring residues contribute to local conformational changes.^94,95^ Importantly, phosphorylation at pS400 does not appear to encode a unique fold or alter the folding trajectory; instead, we propose that it only accelerates the thermodynamically driven rearrangements of the aggregates toward low-energy, disease-relevant conformational states.

### Asymmetrical pS400 Polymorphs Might Represent Secondary Nucleation

We were struck by how the shorter pS400-2 protofilament, PF3, resembles a structure termed the first amyloid intermediate (FIA) that was observed in prior studies (see **Figure 7c**).^79^ In that work, FIA appeared early in the aggregation process and then became less common as the sample progressed towards CTE or AD folds. This observation led to the hypothesis that the FIA might represent a primary nucleation step, which helps direct (or template) the protofilaments. However, in our case, the pS400 sample was aged for 5 weeks to promote thermodynamic maturation, and so PF3 likely represents a relatively late intermediate formed via side-association-driven secondary nucleation rather than an initial nucleation product. Although this structure may also represent a non-productive intermediate, it likely still represents secondary nucleation. Therefore, this is an important finding because secondary nucleation has long been hypothesized to contribute to fibril growth and structure, yet it has not been directly visualized at atomic resolution.^96–100^

The interface between the PF2 and PF3 filaments might, therefore, shed initial light on how secondary nucleation occurs and how it might be targeted by therapeutics. For example, we note that the VQIVYK motif in PF3, which has long been known to be critical for tau self-assembly,^83^ is featured at the PF2-PF3 interface. However, we also find that PF3 has density for residues 317-323 (see **Figure 7c**), which is more than observed in FIA. It is not clear if these “extra” residues differentiate early self-assembly events from later, secondary nucleation events. Regardless, we speculate that this structure could be useful in considering antibodies and other therapeutics that target secondary nucleation.

### Implications for the Identification of Causative Biomarkers for Precision Medicine

In addition to the structure of the PF2-PF3 interface, these findings have other implications for the development of diagnostics and therapeutics for tauopathy. While previous studies have speculated on the role of pS396 in pathology,^101–103^ we directly demonstrated that pS396 (and, secondarily, pS400) promotes aggregation (**Figure 3**). This work provides mechanistic support for designating these sites as causative biomarkers, which might be therapeutic targets for immunotherapy^104–107^, targeted dephosphoryation^108–112^, and as diagnostic indicators.^113–115^ In contrast, pT403 and pS404—despite their prevalence in pathological tau—exert protective effects on nucleation and should be considered correlative biomarkers. Therefore, strategies that decrease the levels of pT403 and pS404 or that use these marks to monitor therapeutic benefits might be confounded by their contribution to tau aggregation. More broadly, therapeutic strategies that indiscriminately target all phosphorylated tau proteoforms may risk removal of neutralizing modifications that are not pathogenic drivers.

In summary, this study establishes that phosphorylation at the PHF-1 epitope regulates tau aggregation in a highly site-specific manner. Phosphorylation of S396 and S400 lowers the kinetic barriers to cofactor-free assembly, while pT403 and pS404 inhibit nucleation, but all of these are also incorporated during seeded growth in the presence of catalytic quantities of pre-formed fibrils. PHF-1 phosphorylation does not stabilize fibrils thermodynamically but accelerates structural maturation toward disease-relevant conformations. Structural studies show that pS400 is sufficient to form CTE-like folds, while also revealing a potential example of secondary nucleation. Future studies are needed to extend these findings to larger tau constructs and additional PTMs (*e.g.,* phosphorylation, ubiquitination, acetylation, glycosylation). Thus, truncated tau constructs may not fully capture the complexity of human pathology. Yet, these results highlight the importance of positional effects in tau phosphorylation and emphasize the need for precision targeting of pathogenic modifications in tauopathies.

## Methods

### Expression and purification of Tau(297-389)-MxeGyrA-His6

*Escherichia coli* BL21 Rosetta 2 (DE3) cells were transformed with a PET28a (+) vector containing Tau(297-389)-Mxe GyrA-His6 intein and plated on a kanamycin-resistant agar plate. Expressions were conducted on a 6 x 1 L scale. A culture was prepared from a single colony in 20 mL of Luria broth supplemented with 50 µg mL^−1^ kanamycin in a 50 mL Falcon tube. This culture was grown at 37 °C until turbidity was observed, then transferred to 300 mL of medium and incubated overnight at 37 °C. Each 1 L culture was inoculated with 50 mL of overnight culture and incubated at 37 °C until optical density (OD_600_) = 0.6 to 0.9. Expression was induced with 0.8 mM isopropyl-β-D-1-thiogalactopyranoside for 4 h at 37 °C. Bacteria were pelleted by centrifugation (4,000 × g, 30 min, 4 °C) and then resuspended in 15 mL of lysis buffer (50 mM (4-(2-hydroxyethyl)-1-piperazineethanesulfonic acid) (HEPES), 50 mM sodium chloride, pH 7.2, 0.1 mM phenylmethylsulfonyl fluoride, oComplete ethylenediaminetetraacetic acid-free protease inhibitor) per liter of culture. Resuspended cells were lysed with a probe sonicator (35% amplitude, 30 s on, 30 s off, 15 min) in an ice bath at 4 °C, and then the cell debris was removed by centrifugation (20,000 × g, 30 min, 4 °C). The supernatant was gravity-purified over a nickel nitrilotriacetic acid resin. The lysate was loaded onto an equilibrated resin bed (∼10 mL of resin, 50 mM HEPES, 200 mM sodium chloride, pH 7.2). After loading, purification was performed using 30 mL portions of elution buffer (0 mM, 200 mM, 400 mM, 600 mM, 800 mM, 1000 mM imidazole in 50 mM HEPES, 200 mM sodium chloride, pH 7.2). For more experimental details and purity information, see **Supplementary Information S1.**

### Intein splicing and hydrazinolysis to produce recombinant Tau(297-389)-NHNH2

Protein samples were collected in 50 mL conical Falcon tubes, precipitated with 0.3 g/mL ammonium sulfate, and collected by centrifugation (4000 x g, 60 min, 4 ⁰C). The protein was then dissolved in splicing buffer (30 mL per conical tube, 2% v/v hydrazine, 100 mM dithiothreitol, 50 mM HEPES, 200 mM sodium chloride, pH 7.2) and incubated at 23 °C for 16 h. To remove any remaining impurities, the reaction was treated with 0.3 mL of formic acid, and insoluble material removed by centrifugation (4000 x g, 60 min, 4 °C). The supernatant was treated with 0.3 g/mL ammonium sulfate, and Tau(297-389)-NHNH_2_ and spliced intein was collected by centrifugation (4 k x g, 60 min, 4 ⁰C). Tau was separated from the intein by gently mixing the pellet in 3 mL of 6 M guandinium hydrochloride, a spatula tip of tris(2-carboxyethyl)phosphine hydrochloride, and 100 µL of formic acid for 1 h until the pellet was completely resuspended, and then it was diluted to 30 mL with buffer (50 mM HEPES, 200 mM sodium chloride pH 7.2). The insoluble intein was removed by centrifugation (4000 x g, 60 min, 4 ⁰C), and Tau(297-389)-NHNH_2_ was precipitated from the supernatant with 0.3 g/mL ammonium sulfate and collected by centrifugation (4000 x g, 60 min, 4 ⁰C). The pellet was dissolved in 3 mL of 6 M guandinium hydrochloride, a spatula tip of tris(2-carboxyethyl)phosphine hydrochloride and 100 μL of formic acid, filtered with a syringe filter (0.22 µm, nylon), and it was purified over a semi-preparative C18 column (10 × 250 mm, 5 µM, 100 Å, 15–50% acetonirile/H_2_O/0.1% trifluoroacetic acid over 60 min, 3 mL/min). The tau band was collected into 2 mL Eppendorf tubes, the fractions were analyzed by ultra-high-performance-liquid-chromatography-mass-spectromatry, and the pure protein was collected and lyophilized to afford Tau(297-389)-NHNH_2_ (108 mg, 18 mg/L). For details on the preparation and characterization of the recombinant Tau(297-389)-NHNH_2_, see **Supplemental Information S1**.

### Selenoesterification to produce recombinant Tau(297-389)-SePh

A 20 mL vial with a stir-bar containing Tau(297-389)-NHNH_2_ (∼100 mg) and diphenyl diselenide (47 mg) was dissolved in argon-purged buffer (3.0 mL, 6 M guandinium hydrochloride, 0.2 M tris(2-carboxyethyl)phosphine hydrochloride, 0.2 M HEPES, pH 2.0) and treated with acetyl acetone (5.4 μL). After the heterogeneous reaction was mixed vigorously at 23 ⁰C for 1.5 h, the diphenyl diselenide was extracted with hexanes (5 x 1 mL), and it was purified over a C18 column (10 x 250 mm, 10 μm, 100 Å, 3.0 mL/min) 5%-65% acetonitrile/H_2_O/0.1% trifluoroacetic acid over 60 min. The fractions were immediately frozen to prevent aqueous hydrolysis, and the pure fractions were lyophilized to afford **1** (72% yield). For details on the preparation and characterization of the recombinant Tau(297-389) selenoester **1**, see **Supplemental Information S1**.

### Synthesis of phosphorylated tau fragments

All peptides containing the individual PHF1 phosphorylations and combinations were synthesised by solid-phase peptide synthesis on Rink-amide resin, and the selenocysteine was installed with a paramethoxybenzyl protective group. The diselenides were prepared upon oxidative cleavage of the paramethoxybenzyl-protected selenides with trifluoroacetic acid/dimethylsulfoxide. For information on the synthesis of these tau fragments, including overall yield and analytical characterization, see **Supplementary Information S2.**

### Diselenide-selenoester-ligation-deselenization to produce semi-synthetic tau proteins

Using air-free techniques to prevent oxidation, a 20 mL vial with a stir-bar containing the selenoester **1** (4 μmol) and the diselenide **2** (3-4 μmol) was treated with argon-purged buffer (1.0 mL, 6 M guandinium hydrochloride, 0.1 M sodium phosphate, pH 6.8), and the vial was carefully sealed under argon. After the reaction was mixed at 23 ⁰C for 2 h, the diphenyl diselenide was carefully extracted with argon-purged hexanes (5 x 1 mL) while maintaining the reaction blanketed under a stream of argon. The reaction was treated with argon-purged deselenization buffer (2.0 mL, 200 mM tris(2-carboxyethyl)phosphine hydrochloride, 100 mM dithiothreitol, 6 M guandinium hydrochloride, 100 mM HEPES, pH 6.9) and it was mixed at 23 ⁰C. After 24 h, the delelenization was filtered through a 0.22 µm syringe filter, and it was purified by sample displacement mode chromatography (Microsorb C18, 4.6 x 250 mm, 5 μm, 100 Å, 50 °C, 1 mL/min, 5%-65% acetonitrile/H_2_O/0.1% trifluoroacetic acid over 60 min), and the fractions that were deemed pure by intact protein mass spectrometry were lyophilized to afford phosphoryated Tau(297-407) **3** (10-30 mg, 34-62% yield). For details on the preparation and characterization of the semi-synthetic phosphoryated Tau(297-403) proteins, see **Supplemental Information S3**.

### Fibril assembly assay

Thioflavin T assays were performed in black, non-treated 96-well microplates (Corning, 3915), sealed with adhesive microplate film (VWR, 7659). Each well was directly filled with 110 μL of reaction mixture (200 µM Tau, 10 mM potassium phosphate buffer, 200 mM potassium citrate tribasic, 10 mM dithiothreitol, 2 µM thioflavin T, pH 7.5) and the plate was incubated at 37 ⁰C in the microplate reader (Molecular Devices M5) without shaking for 7 days. The fluorescence was measured every 10 minutes (Ex = 440 nm, EM = 480 nm, PMT gain = medium, 3 flashes per read). Reactions were performed in biological triplicates, with 6 technical replicates per reaction. Occasionally, wells were visibly evaporated and these were removed from the analysis. The assembly kinetics were estimated by fitting the thioflavin T fluorescence data to a Boltzmann sigmoidal function to obtain t_1/2_ values. For additional information and raw curves, see **Supplementary Information S4**.

### Seeded assembly reactions

Briefly, tau protein (50 µM) was incubated with pre-formed Tau(297-407)-4D fibril seeds (5 µM; see above) in 10 mM potassium phosphate buffer, 200 mM potassium citrate tribasic, 10 mM dithiothreitol, 3 µM thioflavin T) at 37°C for 4 days. The assembly reactions were performed in black, non-binding, low-volume 384-well microplates (Greiner 784900), sealed with adhesive microplate film (VWR 7659). Thioflavin T fluorescence was measured every 10 minutes (Ex = 440 nm, EM = 480 nm, PMT gain = medium, 3 flashes per read), without shaking, in a Molecular Devices M5 plate reader. The reactions were performed in biological triplicates. For additional information and raw curves, see **Supplementary Information S5.**

### Solubility assay

Upon completion of the fibril assembly assays (above), the reactions (22 µL) were transferred to 1.6 mL Eppendorf tubes (Beckmann Coulter, Ref 357488) and centrifuged at 100 k x g for 60 min at 4 °C. The supernatant (20 µL) was collected and transferred to a 0.3 mL autosampler vial. The concentration of tau remaining in the supernatant was determined by high-performance-liquid-chromatography using a single 10 µL injection per sample. The monomer concentrations are reported as the average, and the error bars represent the standard deviation. For additional information and calibration curves, see **Supplementary Information S6.**

### Chemical denaturation assays

Denaturation assays (2 µM Tau(297-407) seeds, 10 mM potassium phosphate buffer pH 7.4, 10 mM dithiothreitol, 10 µM thioflavin T, 23 °C, 16 h) were performed in black, non-binding, low-volume 384-well microplates (Greiner 784900), sealed with adhesive microplate film (VWR 7659). A 1.6 mL Eppendorf tube was filled with 2x fibril master mix, and three wells were filled with 5 μL of 2x fibril and 2x denaturant, for each denaturant concentration. The plate was incubated at 23 °C for 16 h, then analyzed the fluorescence using a SpectraMax M5 microplate reader. For more information and raw curves, see **Supplementary Information S7.**

### Cryo-electron microscopy sample preparation (cryo-EM)

Various dilutions (i.e. 1:5, 1:10, 1:20) of fibrillized phosphorylated Tau(297-407) in fibrillization buffer (10 mM potassium phosphate pH 7.5, 200 mM potassium citrate tribasic, 10 mM dithiothreitol, 3 μM thioflavin T) were prepared after 1 week of quiescent incubation at 37 °C and followed by an additional 4 weeks of quiescent incubation at 23 °C. At the time points, 3 µL of diluted sample was applied to the front side of holey carbon grids (Quantifoil R1.2/1.3 on a gold 300 mesh support) that were glow-discharged for 120 s. Then, 1.5 µL of fibrillization buffer was applied to the back side. The grids were back-blotted (i.e., the blotting paper contacted the back side only) for 8 s at 4 °C and 100% humidity using a Leica GP2 (Leica Microsystems, Waltham, MA), followed by plunge freezing in liquid ethane.

All grids were screened on a Glacios (Thermo Fisher Scientific, Waltham, MA) operated at 200 kV. Two constructs (pS396+pT403; pS400) resulted in sufficient quantities of twisted filaments with well-defined crossovers suitable for structural determination, while all other samples gave non-twisting ribbons or filaments with irregular widths and crossovers. For the pS396 + pT403 sample, a total of 10,331 movies were collected at a nominal magnification of 165,000× (physical pixel size: 0.728 Å/pixel) on a Titan Krios G4 (Thermo Fisher Scientific, Waltham, MA) operated at 300 kV and equipped with a Falcon 4i direct electron detector and Selectris X energy filter (Thermo Fisher Scientific, Waltham, MA) set to a slit width of 20 eV. A defocus range of −0.8 to −1 μm was used with a total exposure time of 4.5-5 s fractionated into 90 TIFF frames. The total dose for each movie was 45 electrons/Å^2^. Movies were motion corrected using MotionCor2 ^116^ in Scipion.^117^ Motion-corrected and dose-weighted micrographs were manually curated in Scipion to remove micrographs lacking filaments, with too high filament density, at low resolution, or with significant ice contamination, resulting in 832 remaining micrographs. For the pS400 sample, a total of 9,823 movies were collected using the same settings as the pS396+pT403 sample. After curation, 1,805 micrographs were used for further processing. For additional information on the cryo-EM sample preparation, see **Supplementary Information S8.**

### Cryo-EM data processing

All image processing was done in RELION 5 and 5.1. ^118–121^ Dose-weighted summed micrographs were imported into RELION 5. The contrast transfer function was estimated using CTFFIND-4.1.^122^ For the pS396+pT403 sample, filaments were picked manually, and segments were extracted from the CTF-corrected micrographs with a box size of 900 pixels downscaled to 300 pixels, resulting in 122,440 segments. Reference-free 2D classification was used to remove contaminants and segments contributing to straight filaments, resulting in 26,038 remaining segments. From these 2D classes, we generated a 3D ab initio model using RELION’s relion_helix_inimodel2d feature. The segments were then re-extracted with a box size of 288 pixels without downscaling. One round of 3D classification with image alignment was performed on the segments using the ab initio model, 3 classes (i.e. k=3), a regularization parameter (T) of 15, and allowing helical rise and twist parameters to vary, starting from an initial rise of 4.78 Å and twist of -1.3°. After 35 iterations, the highest-resolution class (12,980 segments) was subjected to one round of 3D auto-refinement imposing C2_1_ symmetry with the helical parameters and using Blush regularization with the AmyBlush (amy-v1.0) network in RELION 5.1. The refined map underwent standard mask creation and post-processing, to give the final pS396+pT403 map with a global resolution of 3.6 Å, a helical rise of 2.39 Å, and a twist of 179.47°.

For the pS400 sample, filaments were picked manually and segments were extracted with a box size of 900 pixels downscaled to 300 pixels, resulting in 372,025 segments. Multiple rounds of reference-free 2D classification were then performed to identify 2 different major polymorphs: polymorph 1 (122,580 segments, 33% of the data) and polymorph 2 (103,301 segments, 28% of the data). From the highest resolution 2D classes, we generated separate 3D ab initio models as with the pS396+pT403 sample and re-extracted segments with a box size of 288 pixels.

Segments in polymorph 1 underwent two rounds of 3D classification. The first round used the ab initio model as a reference, 25 iterations, k=5, T=10, initial twist -1.2°, initial rise 4.75 Å, allowing helical parameters to vary. The best class (38,134 segments) was used for a second round of 3D classification (40 iterations, k=3, T=20, Blush regularization with the AmyBlush network, initial twist -1.4°, initial rise 4.76 Å, allowing helical parameters to vary). The best class (15,578 segments) underwent 3D auto-refinement using Blush regularization (AmyBlush) then standard mask creation and post-processing, to give the final pS400-1 map with a global resolution of 3.7 Å, a helical rise of 4.76 Å, and a twist of -1.40°.

Segments in polymorph 2 underwent three rounds of 3D classification. The first round used the ab initio model as a reference, 38 iterations, k=5, T=10, initial twist -1.1°, initial rise 4.75 Å, allowing helical parameters to vary. The best class (52,936 segments) was used for a second round of 3D classification (25 iterations, k=4, T=10, initial twist -1.0°, initial rise 4.78 Å, fixed helical parameters). The best class from the second round (32,780 segments) was used for a third round of 3D classification (40 iterations, k=4, T=10, Blush regularization with AmyBlush network, initial twist -1.0°, initial rise 4.78 Å, allowing helical parameters to vary). The best class from the third round (24,468 segments) underwent 3D auto-refinement using Blush regularization (AmyBlush) then standard mask creation and post-processing, to give the final pS400-2 map with a global resolution of 3.6 Å, a helical rise of 4.77 Å, and a twist of -1.36°. For additional information on the cryo-EM data processing, see **Supplementary Information S8.**

### Model building and analysis

Initial model building was performed by fitting existing in vitro tau filament models into the refined maps (pS396+pT403: PDB 7QJW, pS400-1: PDB 8Q9F, pS400-2: 8QJJ). Subsequent model fitting and refinement was performed using ISOLDE.^123^ For pS400-2, ModelAngelo was used to determine the identity of the 19-residue peptide flanking the main protofilament. ^124^ After fitting into the density, each protofilament dimer was translated to give a stack of three rungs (6 strands total per model). After Ramachandran parameters, rotamers, and clashes were satisfied in the middle rung, the outer two rungs were deleted, and the middle rung was re-translated into a new eight-rung stack (16 strands per model). Afterwards, the ISOLDE command “isolde write phenixRsrInput # <MODEL> <MAP resolution> # <MAP>” was used to export the model and generate a rigid-body refinement settings file for real-space refinement in PHENIX,^125^ giving the final refined models. (Note that the validation of 3-rung models often gave spurious map-model FSC curves that started below FSC 0.5). Data were deposited into the Protein Data Bank (PDB) under accession codes 13GX (pS396+pT403), 13GY (pS400-1), 13GZ (pS400-2) and into the Electron Microscopy Data Bank (EMDB) under accession codes EMD-77062 (pS396+pT403), EMD-77063 (pS400-1), and EMD-77064 (pS400-2).

Amyloid packing difference (APD) scores were calculated and associated figures were generated as previously described. ^126^ Other structural figures were made using UCSF ChimeraX.^127^ For additional information on the cryo-EM model building and analysis, see **Supplementary Information S8.**

### Statistical information

All biophysical experiments are performed in biological triplicates and are reported as the average of 3-6 technical replicates. The error of the measurements and error bars are reported as standard deviation or standard error of the mean.

## Data Availability

For details on the experimental methods, raw data and analytical characterization, see the supplemental information.

## Contributions

W.C.P., N.Y., D.R.S., and J.E.G. designed the work. W.C.P., N.Y., E.T., A.A.M., and N.S. completed data acquisition, analysis, and interpretation. W.C.P., N.Y., D.R.S., and J.E.G. drafted the work. D.R.S. and J.E.G. completed the funding aquesition.

## Acknowledgements

We thank Dancia Galonic Fujimori and Darius Mcardle (UCSF) for assistance with intact protein mass spectrometry. We are grateful for the technical support from Ny Sin, Sergei Bolushevsky, Alex Chong, Jay Conrad, Nick Paras and Stanley Prusiner (UCSF). This work was supported by the Tau Consortium (to J.E.G. and D.R.S.).

## Supplementary Information

The cryo-EM data was deposited into the Protein Data Bank (PDB) under accession codes **PDB 13GX** (pS396 + pT403), **PDB 13GY** (pS400-1), **PDB 13GZ** (pS400-2) and into the Electron Microscopy Data Bank (EMDB) under accession codes **EMD-77062** (pS396 + pT403), **EMD-77063** (pS400-1), and **EMD-77064** (pS400-2).

For details on the experimental methods, raw data and analytical characterization, see the supplemental information:

Supplementary Information S1. Preparation of the recombinant tau proteins.

Supplementary Information S2. Preparation of synthetic tau fragments.

Supplementary Information S3. Preparation of semi-synthetic phosphorylated Tau(297-407) proteins.

Supplementary Information S4. Primary nucleation assays.

Supplementary Information S5. Seeded assembly reactions.

Supplementary Information S6. Solubility equilibrium

Supplementary Information S7. Chemical denaturation assays.

Supplementary Information S8. Details of the cryo-EM studies.

