## Supplemental Information for "Phosphorylation of S396 and S400 Promotes Tau Self-Assembly and Favors the Chronic Traumatic Encephalopathy (CTE) Protofilament Fold"

### ABBREVIATIONS.

AA, amino acid; AcAc, acetylacetone; AD, Alzheimer's Disease; Ar, argon; Boc, tert-butyloxycarbonyl; CTE, chronic traumatic encephalopathy; cryo-EM, cryo-electron microscopy; $C_{sat}$, equilibrium solubility; DCM, dichloromethane; DIPEA, diisopropylethylamine; DMF, dimethylformamide; DMSO, dimethylsulfoxide; DSL, diselenide-selenoester-ligation; DPDS, diphenyldiselenide; DTT, dithiothreitol; EDTA, ethylenediaminetetraacetic acid; EMDB, electron-microscopy data bank; FA, formic acid; Fmoc, fluorenylmethyloxycarbonyl; Gnd·HCl, guanidinium hydrochloride; HATU, hexafluorophosphate azabenzotriazole tetramethyl uranium; HEPES, (4-(2-hydroxyethyl)-1-piperazineethanesulfonic acid); HOBt, hydroxybenzotriazole; HPLC, high-performance liquid chromatography; HRMS, high-resolution mass spectrometry; IPTG, Isopropyl-β-D-1-thiogalactopyranoside; KPB, potassium phosphate buffer; MeCN, acetonitrile; Mob, 4-methoxybenzyl; NFT, neurofibrillary tangle; NTA, nitrilotriacetic acid; oxyma, ethyl cyanohydroxyiminoacetate; Pbf, 2,2,4,6,7-pentamethyldihydrobenzofuran-5-sulfonyl; PDB, protein-data bank; PF, protofilament; PHF, paired helical filament; PMB, paramethoxybenzyl; PMSF, phenylmethylsulfonyl fluoride; PTMs, post-translational modifications; Sec, selenocysteine; SPPS, solid-phase-peptide synthesis; TCEP·HCl, Tris(2-carboxyethyl)phosphine hydrochloride; TIPSH, triispropylsilane; TFA, trifluoroacetic acid; ThT, thioflavin T; Trt, trityl; UPLC, ultra-high performance liquid chromatography.

### GENERAL INFORMATION.

All chemicals were purchased as reagent grade and used as received without further purification. All buffers were filtered (0.22 μm) before use and stored at 4 ºC for no longer than 2 months. All biochemical experiments were conducted in milliQ $H_2O$. All additives (*e.g.,* citrates, DTT, salts) were dissolved in $H_2O$ and used immediately. Gel electrophoresis was performed in Tris-glycine running buffer using a Bio-Rad electrophoresis system. Ultracentrifugation was performed on Beckman Coulter Optima systems. Protein quantification (A280) was performed on a Thermo Scientific NanoDrop 200c spectrophotometer. Thioflavin T measurements were conducted in multimode SpectraMax M5 plate readers (Molecular Devices). Analytical and preparative HPLC was performed on an Agilent 1100 Series HPLC System (American Laboratory Trading, G1311A Quaternary Pump, G1313A Autosampler, G1315B Diode Array Detector, G1316A Column Compartment, and G1379A Degasser with Solvent Tray). Ultra-high-performance liquid chromatography (UPLC) and high-resolution mass spectrometry (HRMS) were performed using a Waters XEVO G2-S QTof Mass Spectrometer with an Acquity UPLC system. Electron microscopy grids were glow-discharged on a PELCO easiGlow system. Cryo-electron microscopy (cryo-EM) was conducted on a Thermo Scientific Titan Krios G4 with a Thermo Scientific Falcon 4i camera. Cryo-EM grids were prepared on a Lecia EM GP2 plunge freezer.

### S1. PREPARATION OF THE RECOMBINANT TAU FRAGMENT.

#### S1a. Expression of Tau(297-389)-$NHNH_2$.

**Tau(297-389)-$NHNH_2$ (S2).**

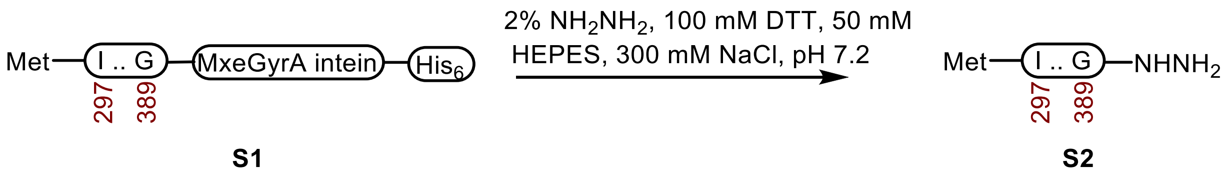

**Preparation of PET28a (+) tau plasmids.** A gene containing the tau construct was installed into a PET28a (+) vector using the Gibson assembly. The plasmid was then grown in XL1-Blue Competent Cells (Agilent) and isolated using the QIAprep Spin Miniprep Kit (250, Qiagen) according to the manufacturer's protocol. The purified plasmid was sequenced (plasmidsaurus) to validate that the gene was successfully inserted and that no mutations occurred.

**Tau(297-389).**
MIKHVPGGGSVQIVYKPVDLSKVTSKCGSLGNIHHKPGGGQVEVKSEKLDFKDRVQSKIGSLDNITHVPGGGN
KKIETHKLTFRENAKAKTDHG

**MxeGyrA inten.**

CITGDALVALPEGESVRIADIVPGARPNSDNAIDLKVLDRHGNPVLADRLFHSGEHPVYTVRTVEGLRVTGTAN
HPLLCLVDVAGVPTLLWKLIDEIKPGDYAVIQRSAFSVDCAGFARGKPEFAPTTYTVGVPGLVRFLEAHHRDPD
AQAIADELTDGRFYYAKVASVTDAGVQPVYSLRVDTADHAFITNGFVSHATGLTGIHHHHHHH

**Expression of Tau(297-389)-MxeGyrA-His6.** *Escherichia coli* BL21 Rosetta 2 (DE3) cells were transformed with a PET28a (+) vector containing Tau(297-389)-Mxe GyrA-His6 intein and plated on a kanamycin-resistant agar plate. Expressions were conducted on a 6 x 1 L scale. A morning culture was prepared from a single colony in 20 mL of Luria broth (LB) supplemented with 50 µg $ml^{-1}$ kanamycin in a 50 mL Falcon tube. It was grown at 37 °C until turbidity was observed, then transferred to 300 mL of medium and incubated overnight at 37 °C. Each 1 L culture was inoculated with 50 mL of overnight culture and incubated at 37 ºC until optical density ($OD_{600}$) = 0.6-0.9. Expression was induced with 0.8 mM IPTG for 4 h at 37 °C. Bacteria were pelleted by centrifugation (4,000 × g, 30 min, 4 °C) and then resuspended in 15 mL of lysis buffer (50 mM HEPES, 50 mM NaCl, pH 7.2, 0.1 mM PMSF, oComplete EDTA-free protease inhibitor) per liter of culture. Resuspended cells were lysed with a probe sonicator (35% amplitude, 30 s on, 30 s off, 15 min) in an ice bath at 4ºC, and then the impurities were removed by centrifugation (20,000 × g, 30 min, 4 °C).

**Purification.** The supernatant was gravity-purified over a Ni-NTA column. The lysate was loaded onto an equilibrated resin bed (~10 mL of resin, 50 mM HEPES, 200 mM NaCl, pH 7.2). After loading, purification was performed using 30 mL portions of elution buffer (0 mM, 200 mM, 400 mM, 600 mM, 800 mM, 1000 mM imidazole in 50 mM HEPES, 200 mM NaCl, pH 7.2). The fractions were analyzed by SDS-PAGE.

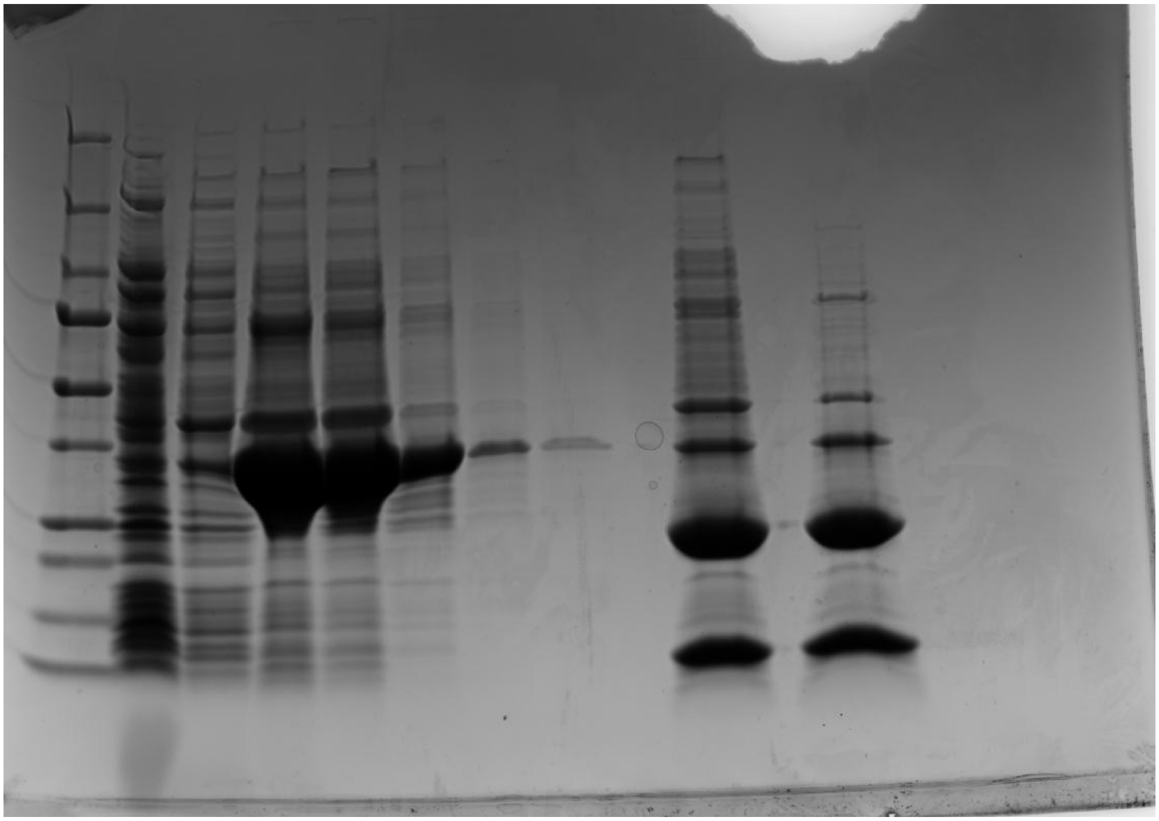

SDS PAGE (4–20% Mini-PROTEAN® TGX™ Precast Protein Gels, Bio-Rad 4561096, tris-glycine running buffer, 180 V for 45 min at 23 ºC) of IMAC affinity purification and intein splicing. Lanes: 1. Precision Plus Protein Dual Color Standards (Bio-Rad 1610374); 2. flow through; 3. 0 mM imidazole; 4. 200 mM imidazole; 5. 400 mM imidazole; 6. 600 mM imidazole; 7. 800 mM imidazole; 8. 1000 mM imidazole; 10. Representative SDS-PAGE of intein splicing products after 16 h of incubation; 12. Representative SDS-PAGE of intein splicing products after formic acid treatment and centrifugation, which removes most of the remaining bacterial proteins.

**Intein splicing.** The fractions containing the protein (lanes 4-6) were collected in 50 mL conical Falcon tubes, precipitated with 0.3 g/mL ammonium sulfate, and collected by centrifugation (4 k x g, 60 min, 4 ºC). The protein was dissolved in splicing buffer (30 mL per conical tube, 2% v/v hydrazine, 100 mM DTT, 50 mM HEPES, 200 mM NaCl, pH 7.2) and incubated at 23 °C for 16 h (see lane 10 of the SDS-PAGE). To remove impure bacterial proteins, the reaction was treated with 0.3 mL of formic acid, and insoluble impurities were removed by centrifugation (4 k x g, 60 min, 4 °C) (see lane 12 of the SDS-PAGE). The supernatant was treated with 0.3 g/mL ammonium sulfate, and Tau(297-389)-$NHNH_2$ and spliced intein was collected by centrifugation (4 k x g, 60 min, 4 ºC). Tau was separated from the intein by gently mixing the pellet in 3 ml of 6 M Gnd·HCl, a spatula tip of TCEP·HCl, and 100 µL of formic acid for 1 h until the pellet was completely resuspended, and then it was diluted to 30 mL with buffer (50 mM HEPES, 200 mM NaCl pH 7.2). The insoluble intein was removed by centrifugation (4 k x g, 60 min, 4 ºC), and Tau(297-389)-$NHNH_2$ was precipitated from the supernatant with 0.3 g/mL ammonium sulfate and collected by centrifugation (4 k x g, 60 min, 4 ºC). The pellet was dissolved in 3 ml of 6 M Gnd·HCl, a spatula tip of TCEP·HCl, and 100 µL of formic acid, filtered with a syringe filter (0.22 µm, nylon), and it was purified over a semi-preparative C18 column (10 × 250 mm, 5 µM, 100 Å, 15–50% MeCN/$H_2O$/0.1% TFA over 60 min, 3 mL/min). The tau band was collected into 2 mL Eppendorf tubes, the fractions were analyzed by UPLC-HRMS, and the pure protein was collected and lyophilized to afford Tau(297-389)-$NHNH_2$ **S2** (108 mg, 18 mg/L).

- The hydrazide yield ranges from 14.3 to 29.0 mg/L.

- Incomplete hydrolysis of the first methionine start codon occurs. A 3-4 h expression at 37 ºC has lower levels of methionine cleavage than an overnight expression at 16 ºC.
- HPLC purification is required to remove the incomplete partial methionine cleavage product, as well as the proteolysis products.

**UPLC-HRMS:** Measured on an ACQUITY UPLC Protein BEH C4 column (2.1 x 50 mm; 1.7 µm; 300 Å; 55 °C; 0.6 mL/min) with a gradient of 5% MeCN/$H_2O$/0.1% FA until 0.1 min, 5-95% MeCN/$H_2O$/0.1% FA until 1.0 min, 95% MeCN/$H_2O$/0.1% FA until 1.1 min.

**Calculated Mass:** $C_{442}H_{729}N_{135}O_{132}S_2$ 10110.627, 10110.0 found.

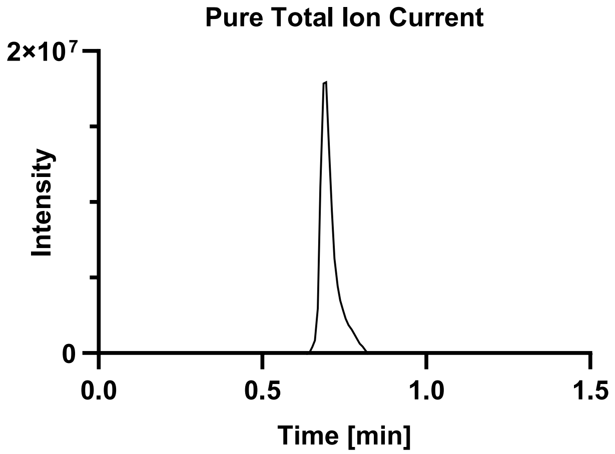

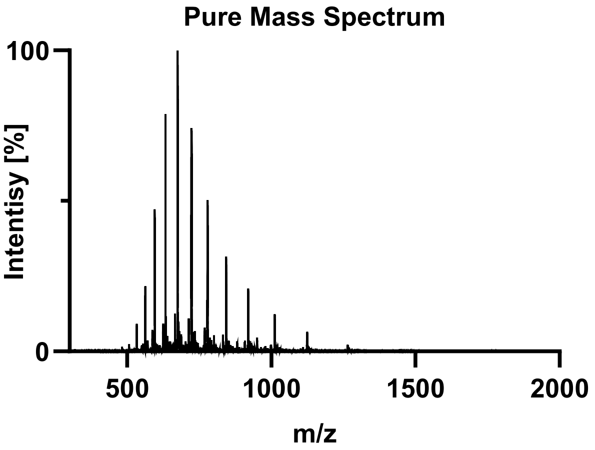

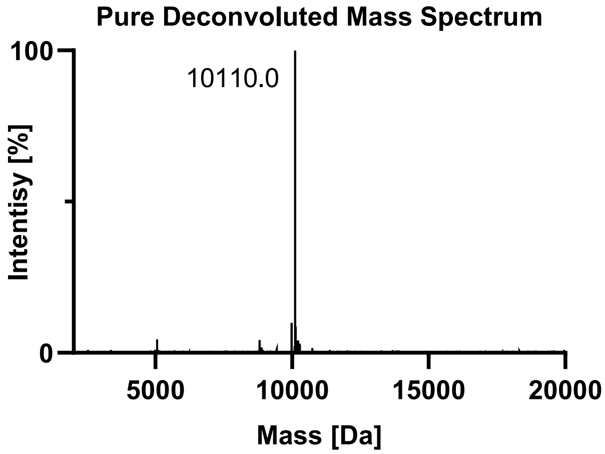

#### S1b. Synthesis of Met-Tau(297-389)-SePh.

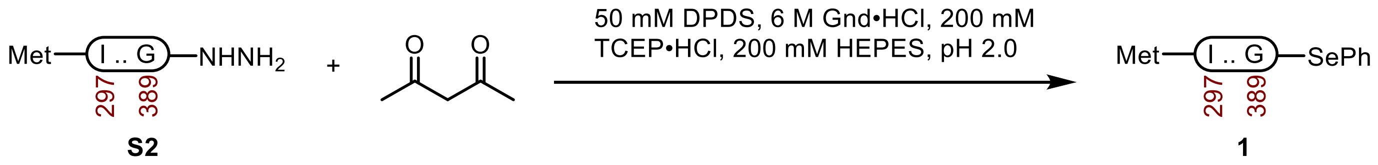

**Tau(297-389)-SePh (1).** A 20 mL vial with a stir-bar containing **S2** (108 mg, 10.5 µmol) and DPDS (47 mg, 151 µmol) was dissolved in Ar-purged buffer (3.0 mL, 6 M Gnd·HCl, 0.2 M TCEP·HCl, 0.2 M HEPES, pH 2.0) and treated with AcAc (5.4 µL, 52.5 µmol). After the heterogeneous reaction was mixed vigorously at 23 ºC for 1.5 h, the DPDS was extracted with hexanes (5 x 1 mL), and it was purified over a C18 column (10 x 250 mm, 10 µm, 100 Å, 3.0 mL/min) 5%-65% MeCN/$H_2O$/0.1% TFA over 60 min. The fractions were lyophilized to afford **1** (77.7 mg, 72% yield).

- The HPLC fractions were immediately frozen at -80 ºC after elution from the column to prevent hydrolysis.
- For setups with 5 mL HPLC sample loops, the reaction is poorly scalable past ~100 mg because the maximal concentration is ~4 mM. Higher concentrations of the hydrazide yield the acyl pyrazole intermediate, and extended reaction times produce hydrolyzed product instead of selenoester.
- The acyl pyrazole intermediate has similar stability as the selenoester towards HPLC purification, and it can be recovered and subjected to the same reaction conditions.

**UPLC-HRMS:** Measured on an ACQUITY UPLC Protein BEH C4 column (2.1 x 50 mm; 1.7 µm; 300 Å; 55 °C; 0.6 mL/min) with a gradient of 5% MeCN/$H_2O$/0.1% FA until 0.1 min, 5-95% MeCN/$H_2O$/0.1% FA until 1.0 min, 95% MeCN/$H_2O$/0.1% FA until 1.1 min.

**Calculated Mass:** $C_{442}H_{729}N_{135}O_{132}S_2$ 10235.666, 10235.0 found.

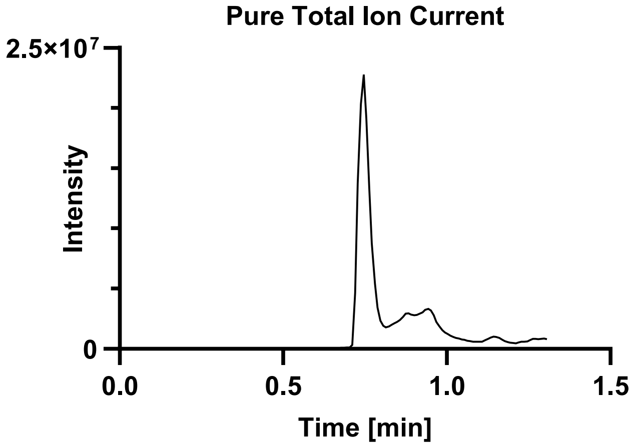

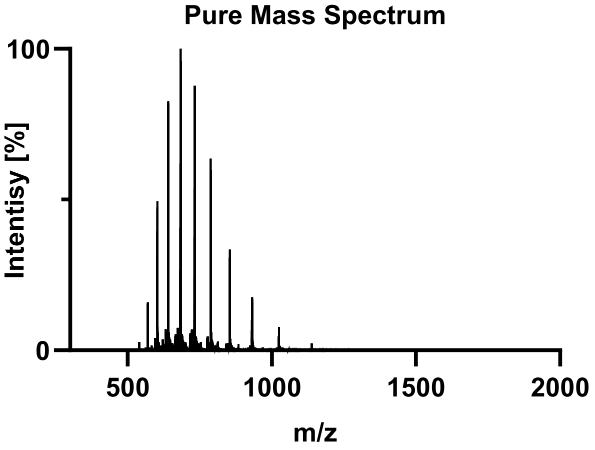

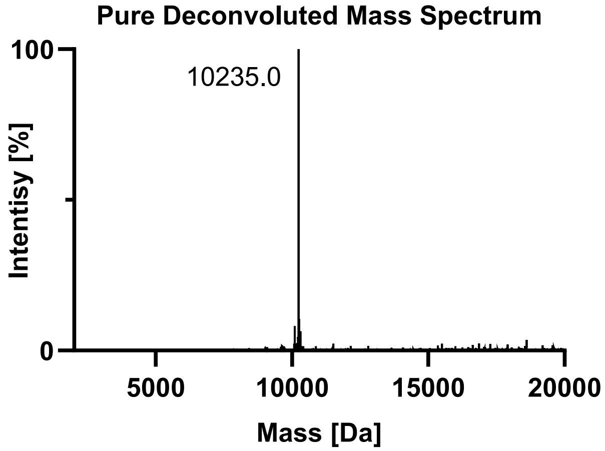

### S2. SYNTHESIS OF PHOSPHORYLATED TAU FRAGMENTs.

Tau(390-407) WT (**S3a**).

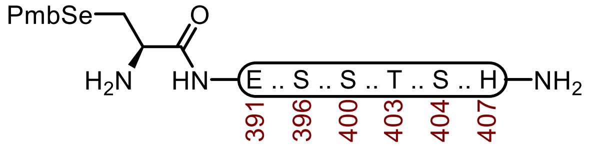

**S3a**

**Sequence:** H-C(SePmb)EIVYKSPVVSGDTSPRH-$NH_2$

**Amino Acids:** Fmoc-L-SeC(Mob)-OH, Fmoc-L-Glu(O*t*Bu)-OH, Fmoc-Ile-OH, Fmoc-Val-OH, Fmoc-Tyr(*t*Bu)-OH, Fmoc-Lys(Boc)-OH, Fmoc-Ser(*t*Bu)-OH, Fmoc-Pro-OH, Fmoc-Val-OH, Fmoc-Gly-OH, Fmoc-Thr(*t*Bu)-OH, Fmoc-Arg(Pbf)-OH, Fmoc-His(Trt)-OH.

**Resin:** Fmoc-RinkAmide Protide resin from CEM (100-200 mesh, 0.6 mmol/g, 0.833 g, 0.50 mmol) was used.

**Coupling:** Amino acids were double coupled using a preactivated solution of Fmoc-AA-OH (0.25 M, 4 equiv.), HATU (0.5 M, 4 equiv.), and DIPEA (5 equiv.) at 23 °C for 1 h. The resin was washed with DMF (2 x 35 mL)

**Selenocysteine Coupling:** Unique amino acids were single coupled using 0.25 M amino acid in DMF (2 equiv.), 0.5 M HATU (2 equiv.), oxyma (2 equiv.) and DIPEA (7 equiv.) at 23 °C for 16 h.

**Caping:** After double coupling, the resin was capped with $Ac_2O$ (5 equiv.) and DIPEA (10 equiv.) in DMF (20 mL) for 15 min. The resin was washed with DMF (3 x 20 mL).

**Deprotection:** 20% piperidine + 0.1 M HOBt in DMF (30 mL) at 23 ºC (2 x 15 min). The resin was washed with DMF (5 x 20 mL).

**Global Deprotection:** The dry resin in a 250 mL polypropylene vial was treated with TFA:$H_2O$:thioanisole:TIPSH (85:5:5:5, 15 mL) and mixed on an overhead stirrer. The mixture was stirred at 23 ºC for 3 h before filtering, and the resin was washed with an additional portion of TFA (3 x 4 mL). The pooled filtrate was cooled to 0°C and treated with $Et_2O$ to precipitate the peptide. The precipitate was centrifuged, and the resulting pellet was washed twice more with $Et_2O$ before it was dissolved in 40% MeCN and lyophilized.

**Crude UPLC-MS:** Measured on a HALO C18 column (3.0 x 30 mm; 2.7 µm; 90 Å; 40 °C; 1.5 mL/min) with a gradient of 5% MeCN/$H_2O$/0.1% TFA until 0.3 min, 5-95% MeCN/$H_2O$/0.1% TFA until 3.0 min, 95% MeCN/$H_2O$/0.1% TFA until 4.0 min.

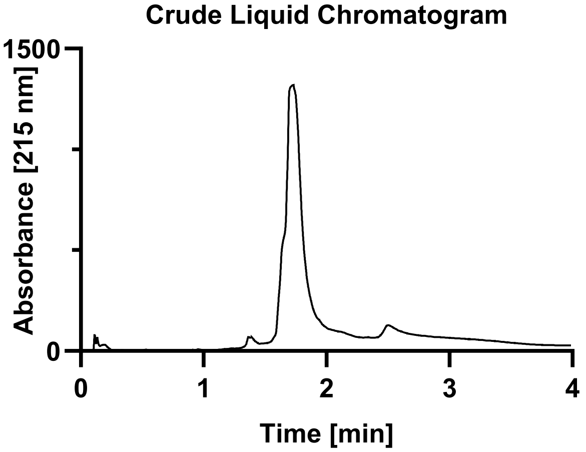

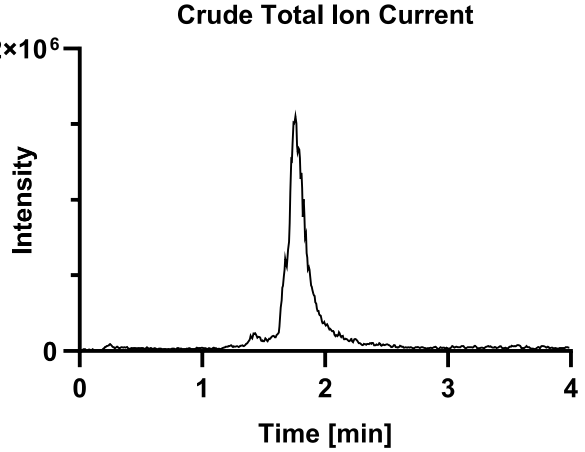

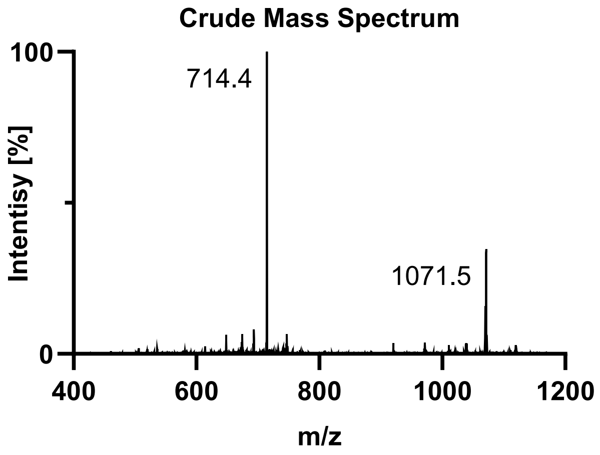

**Purification**: The material was purified over a semi-prep C4 column (10 x 250 mm, 10 µm, 300 Å, 5.0 mL/min) 18-45% MeCN/$H_2O$/0.1% TFA over 30 min. The fractions were lyophilized to afford **S3a** (93.0 mg, 9% yield).

**UPLC-MS:** Measured on a HALO C18 column (3.0 x 30 mm; 2.7 µm; 90 Å; 40 °C; 1.5 mL/min) with a gradient of 5% MeCN/$H_2O$/0.1% TFA until 0.3 min, 5-95% MeCN/$H_2O$/0.1% TFA until 3.0 min, 95% MeCN/$H_2O$/0.1% TFA until 4.0 min.

**Chemical Formula:** $C_{93}H_{145}N_{25}O_{28}Se$
**Molecular Weight:** 2140.3010 g/mol
**LRMS for [M+2H+]:** $C_{93}H_{147}N_{25}O_{28}Se^{2+}$ 1071.5, found 1070.9.
**LRMS for [M+3H+]:** $C_{93}H_{148}N_{25}O_{28}Se^{3+}$ 714.7, found 714.5.

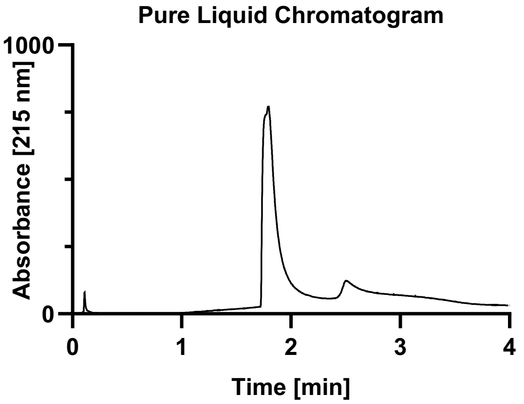

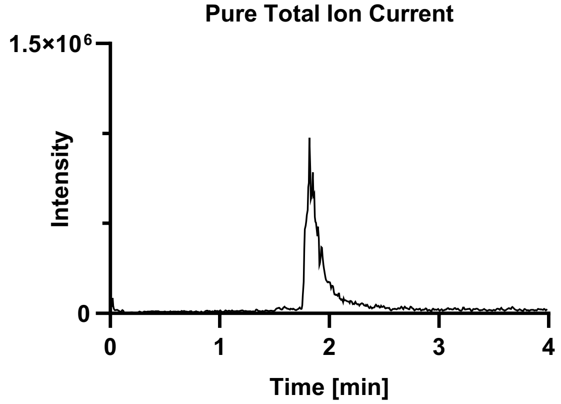

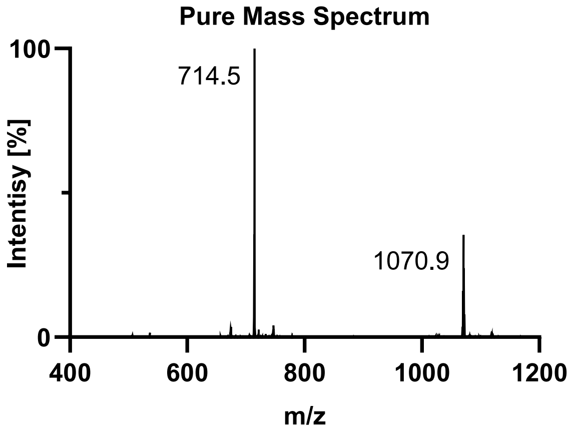

Tau(390-407) pS396 (**S3b**).

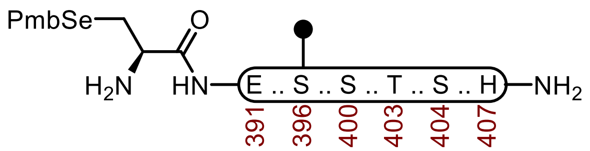

**S3b**

**Sequence:** H-C(SePmb)EIVYKS(OPO$_3$HBn)PVVSGDTSPRH-NH$_2$

**Amino Acids:** Fmoc-L-SeC(Mob)-OH, Fmoc-L-Glu(O*t*Bu)-OH, Fmoc-Ile-OH, Fmoc-Val-OH, Fmoc-Tyr(*t*Bu)-OH, Fmoc-Lys(Boc)-OH, Fmoc-Ser(*t*Bu)-OH, Fmoc-Pro-OH, Fmoc-Val-OH, Fmoc-Ser(OPO$_3$HBn)-OH, Fmoc-Gly-OH, Fmoc-Thr(*t*Bu)-OH, Fmoc-Arg(Pbf)-OH, Fmoc-His(Trt)-OH.

**Resin:** Fmoc-RinkAmide Protide resin from CEM (100-200 mesh, 0.6 mmol/g, 0.833 g, 0.50 mmol) was used.

**Coupling:** Amino acids were double coupled using a preactivated solution of Fmoc-AA-OH (0.25 M, 4 equiv.), HATU (0.5 M, 4 equiv.), and DIPEA (5 equiv.) at 23 °C for 1 h. The resin was washed with DMF (2 x 35 mL)

**Phosphoserine, phosphothreonine, and Selenocysteine Coupling:** Unique amino acids were single coupled using 0.25 M amino acid in DMF (2 equiv.), 0.5 M HATU (2 equiv.), oxyma (2 equiv.) and DIPEA (7 equiv.) at 23 °C for 16 h.

**Caping:** After double coupling, the resin was capped with Ac$_2$O (5 equiv.) and DIPEA (10 equiv.) in DMF (20 mL) for 15 min. The resin was washed with DMF (3 x 20 mL).

**Deprotection:** 20% piperidine + 0.1 M HOBt in DMF (30 mL) at 23 ºC (2 x 15 min). The resin was washed with DMF (5 x 20 mL).

**Global Deprotection:** The dry resin in a 250 mL polypropylene vial was treated with TFA:H$_2$O:thioanisole:TIPSH (85:5:5:5, 15 mL) and mixed on an overhead stirrer. The mixture was stirred at 23 ºC for 3 h before filtering, and the resin was washed with an additional portion of TFA (3 x 4 mL). The pooled filtrate was cooled to 0°C and treated with Et$_2$O to precipitate the peptide. The precipitate was centrifuged, and the resulting pellet was washed twice more with Et$_2$O before it was dissolved in 40% MeCN and lyophilized.

**Crude UPLC-MS:** Measured on a HALO C18 column (3.0 x 30 mm; 2.7 µm; 90 Å; 40 °C; 1.5 mL/min) with a gradient of 5% MeCN/H$_2$O/0.1% TFA until 0.3 min, 5-95% MeCN/H$_2$O/0.1% TFA until 3.0 min, 95% MeCN/H$_2$O/0.1% TFA until 4.0 min.

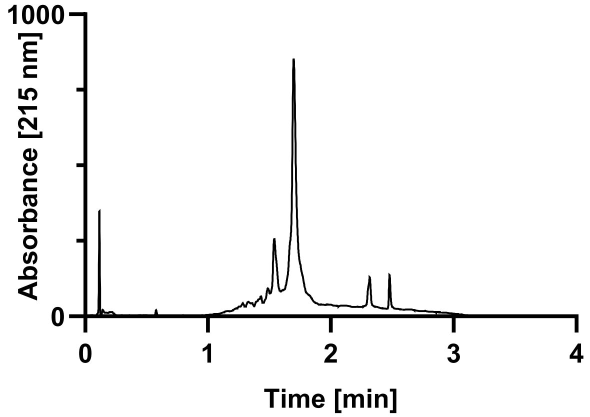

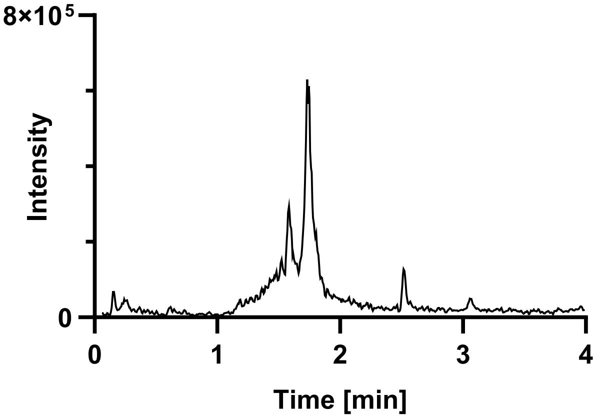

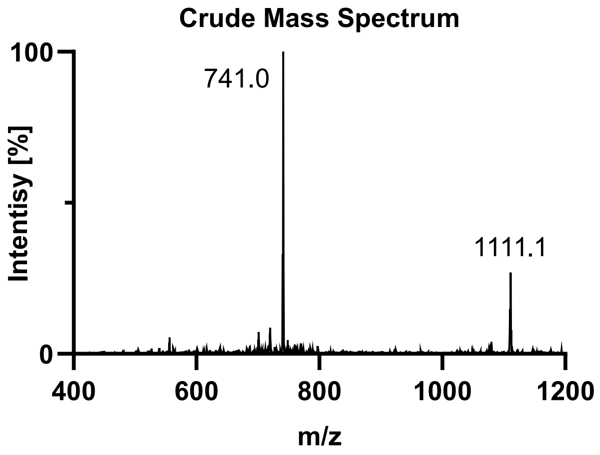

**Purification**: The material was purified over a semi-prep C4 column (10 x 250 mm, 10 µm, 300 Å, 5.0 mL/min) 18-45% MeCN/$H_2O$/0.1% TFA over 30 min. The fractions were lyophilized to afford **S3b** (80.8 mg, 7% yield).

**UPLC-MS:** Measured on a HALO C18 column (3.0 x 30 mm; 2.7 µm; 90 Å; 40 °C; 1.5 mL/min) with a gradient of 5% MeCN/$H_2O$/0.1% TFA until 0.3 min, 5-95% MeCN/$H_2O$/0.1% TFA until 3.0 min, 95% MeCN/$H_2O$/0.1% TFA until 4.0 min.
**Chemical Formula:** $C_{93}H_{146}N_{25}O_{31}PSe$
**Molecular Weight:** 2220.280 g/mol
**LRMS for [M+2H+]:** $C_{93}H_{148}N_{25}O_{31}PSe^{2+}$ 1111.5, found 1111.1.
**LRMS for [M+3H+]:** $C_{93}H_{149}N_{25}O_{31}PSe^{2+}$ 741.3, found 741.0.

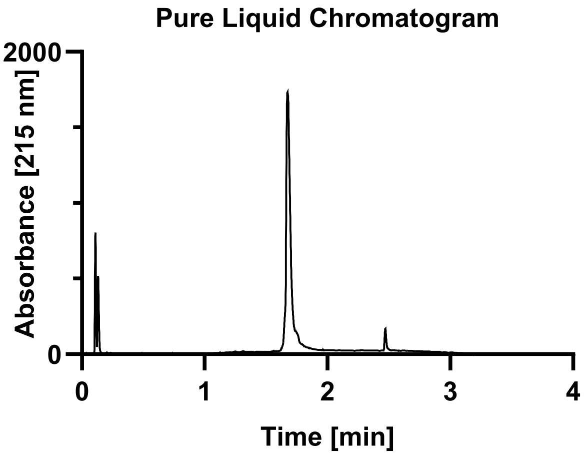

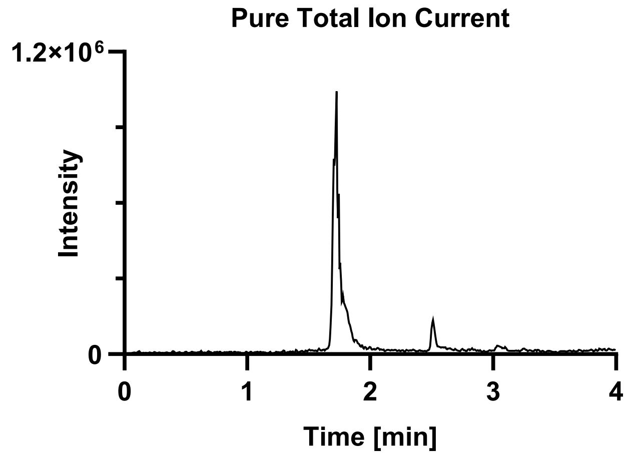

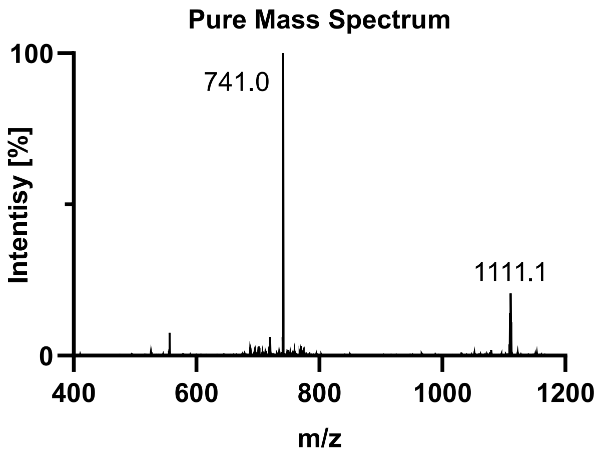

Tau(390-407) pS400 (**S3c**).

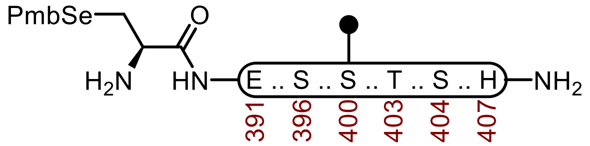

**S3c**

**Sequence:** H-C(SePmb)EIVYKSPVVS($OPO_3HBn$)GDTSPRH-$NH_2$

**Amino Acids:** Fmoc-L-SeC(Mob)-OH, Fmoc-L-Glu(O*t*Bu)-OH, Fmoc-Ile-OH, Fmoc-Val-OH, Fmoc-Tyr(*t*Bu)-OH, Fmoc-Lys(Boc)-OH, Fmoc-Ser(*t*Bu)-OH, Fmoc-Pro-OH, Fmoc-Val-OH, Fmoc-Ser($OPO_3HBn$)-OH, Fmoc-Gly-OH, Fmoc-Thr(*t*Bu)-OH, Fmoc-Arg(Pbf)-OH, Fmoc-His(Trt)-OH.

**Resin:** Fmoc-RinkAmide Protide resin from CEM (100-200 mesh, 0.6 mmol/g, 0.833 g, 0.50 mmol) was used.

**Coupling:** Amino acids were double coupled using a preactivated solution of Fmoc-AA-OH (0.25 M, 4 equiv.), HATU (0.5 M, 4 equiv.), and DIPEA (5 equiv.) at 23 °C for 1 h. The resin was washed with DMF (2 x 35 mL)

**Phosphoserine, phosphothreonine, and Selenocysteine Coupling.** Unique amino acids were single coupled using 0.25 M amino acid in DMF (2 equiv.), 0.5 M HATU (2 equiv.), oxyma (2 equiv.) and DIPEA (7 equiv.) at 23 °C for 16 h.

**Caping:** After double coupling, the resin was capped with $Ac_2O$ (5 equiv.) and DIPEA (10 equiv.) in DMF (20 mL) for 15 min. The resin was washed with DMF (3 x 20 mL).

**Deprotection:** 20% piperidine + 0.1 M HOBt in DMF (30 mL) at 23 ºC (2 x 15 min). The resin was washed with DMF (5 x 20 mL).

**Global Deprotection:** The dry resin in a 250 mL polypropylene vial was treated with TFA:$H_2O$:thioanisole:TIPSH (85:5:5:5, 15 mL) and mixed on an overhead stirrer. The mixture was stirred at 23 ºC for 3 h before filtering, and the resin was washed with an additional portion of TFA (3 x 4 mL). The pooled filtrate was cooled to 0°C and treated with $Et_2O$ to precipitate the peptide. The precipitate was centrifuged, and the resulting pellet was washed twice more with $Et_2O$ before it was dissolved in 40% MeCN and lyophilized.

**Crude UPLC-MS:** Measured on a HALO C18 column (3.0 x 30 mm; 2.7 µm; 90 Å; 40 °C; 1.5 mL/min) with a gradient of 5% MeCN/$H_2O$/0.1% TFA until 0.3 min, 5-95% MeCN/$H_2O$/0.1% TFA until 3.0 min, 95% MeCN/$H_2O$/0.1% TFA until 4.0 min.

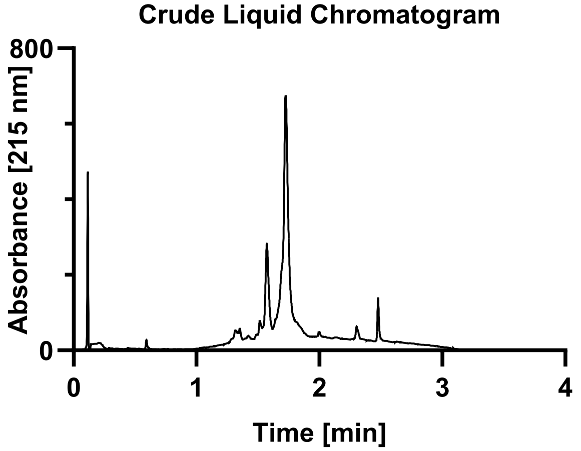

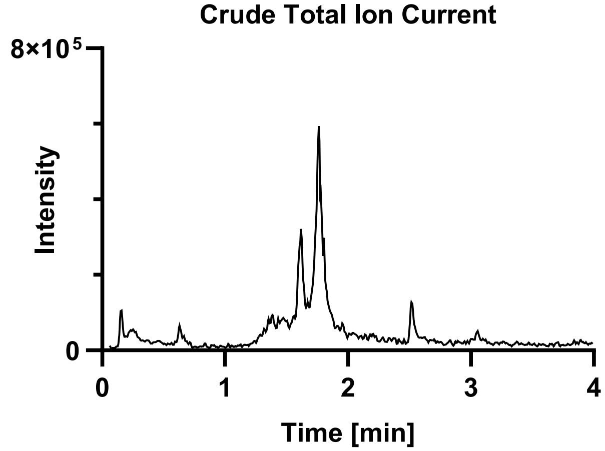

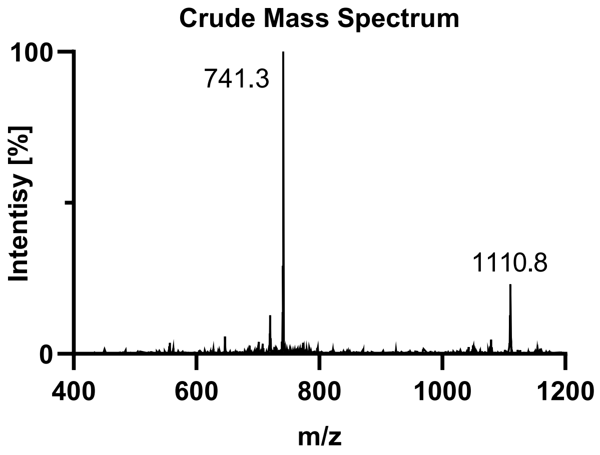

**Purification**: The material was purified over a semi-prep C4 column (10 x 250 mm, 10 µm, 300 Å, 5.0 mL/min) 18-45% MeCN/$H_2O$/0.1% TFA over 30 min. The fractions were lyophilized to afford **S3c** (86.1 mg, 8% yield).

**Pure UPLC-MS:** Measured on a HALO C18 column (3.0 x 30 mm; 2.7 µm; 90 Å; 40 °C; 1.5 mL/min) with a gradient of 5% MeCN/$H_2O$/0.1% TFA until 0.3 min, 5-95% MeCN/$H_2O$/0.1% TFA until 3.0 min, 95% MeCN/$H_2O$/0.1% TFA until 4.0 min.
**Chemical Formula**: $C_{93}H_{146}N_{25}O_{31}PSe$
**Molecular Weight:** 2220.280 g/mol
**LRMS for [M+2H+]:** $C_{93}H_{148}N_{25}O_{31}PSe^{2+}$ 1111.5, found 1110.8.
**LRMS for [M+3H+]:** $C_{93}H_{149}N_{25}O_{31}PSe^{2+}$ 741.3, found 741.0.

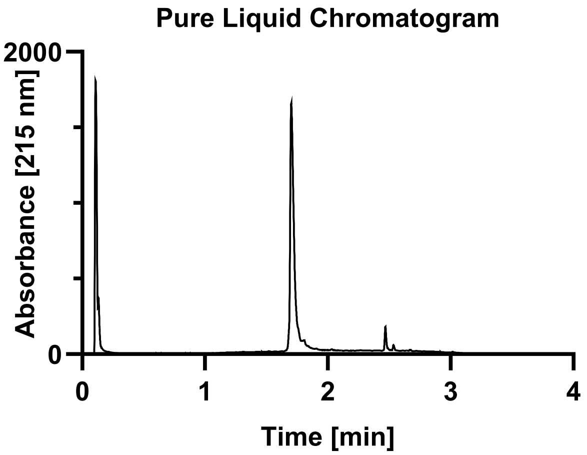

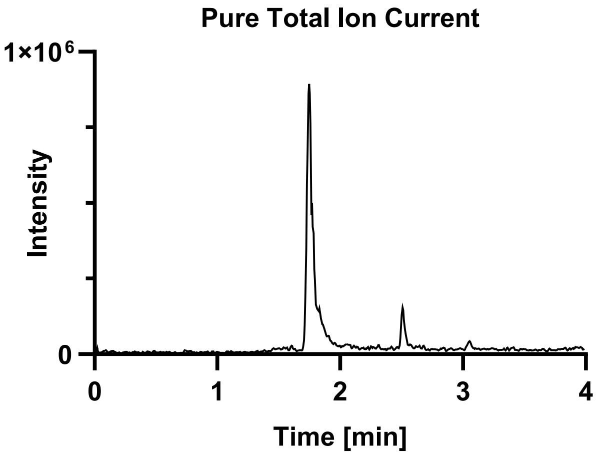

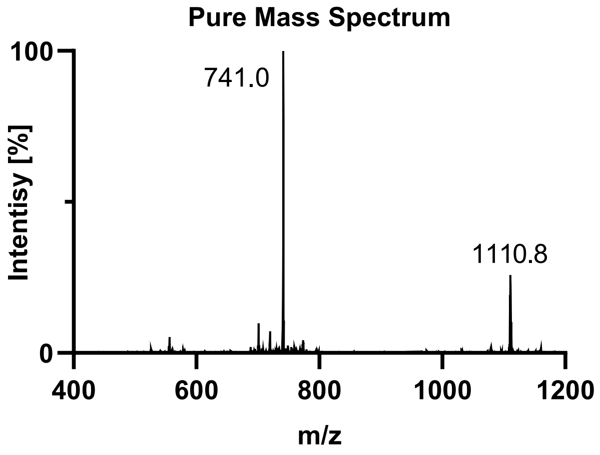

Tau(390-407) pT403 (**S3d**).

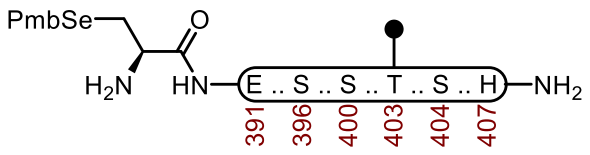

**S3d**

**Sequence:** H-C(SePmb)EIVYKSPVVSGDT($OPO_3HBn$)SPRH-$NH_2$

**Amino Acids:** Fmoc-L-SeC(Mob)-OH, Fmoc-L-Glu(O*t*Bu)-OH, Fmoc-Ile-OH, Fmoc-Val-OH, Fmoc-Tyr(*t*Bu)-OH, Fmoc-Lys(Boc)-OH, Fmoc-Ser(*t*Bu)-OH, Fmoc-Pro-OH, Fmoc-Val-OH, Fmoc-Ser($OPO_3HBn$)-OH, Fmoc-Gly-OH, Fmoc-Thr(*t*Bu)-OH, Fmoc-Arg(Pbf)-OH, Fmoc-His(Trt)-OH.

**Resin:** Fmoc-RinkAmide Protide resin from CEM (100-200 mesh, 0.6 mmol/g, 0.833 g, 0.50 mmol) was used.

**Coupling:** Amino acids were double coupled using a preactivated solution of Fmoc-AA-OH (0.25 M, 4 equiv.), HATU (0.5 M, 4 equiv.), and DIPEA (5 equiv.) at 23 °C for 1 h. The resin was washed with DMF (2 x 35 mL)

**Phosphoserine, phosphothreonine, and Selenocysteine Coupling.** Unique amino acids were single coupled using 0.25 M amino acid in DMF (2 equiv.), 0.5 M HATU (2 equiv.), oxyma (2 equiv.) and DIPEA (7 equiv.) at 23 °C for 16 h.

**Caping:** After double coupling, the resin was capped with $Ac_2O$ (5 equiv.) and DIPEA (10 equiv.) in DMF (20 mL) for 15 min. The resin was washed with DMF (3 x 20 mL).

**Deprotection:** 20% piperidine + 0.1 M HOBt in DMF (30 mL) at 23 ºC (2 x 15 min). The resin was washed with DMF (5 x 20 mL).

**Global Deprotection:** The dry resin in a 250 mL polypropylene vial was treated with TFA:$H_2O$:thioanisole:TIPSH (85:5:5:5, 15 mL) and mixed on an overhead stirrer. The mixture was stirred at 23 ºC for 3 h before filtering, and the resin was washed with an additional portion of TFA (3 x 4 mL). The pooled filtrate was cooled to 0°C and treated with $Et_2O$ to precipitate the peptide. The precipitate was centrifuged, and the resulting pellet was washed twice more with $Et_2O$ before it was dissolved in 40% MeCN and lyophilized.

**Crude UPLC-MS:** Measured on a HALO C18 column (3.0 x 30 mm; 2.7 µm; 90 Å; 40 °C; 1.5 mL/min) with a gradient of 5% MeCN/$H_2O$/0.1% TFA until 0.3 min, 5-95% MeCN/$H_2O$/0.1% TFA until 3.0 min, 95% MeCN/$H_2O$/0.1% TFA until 4.0 min.

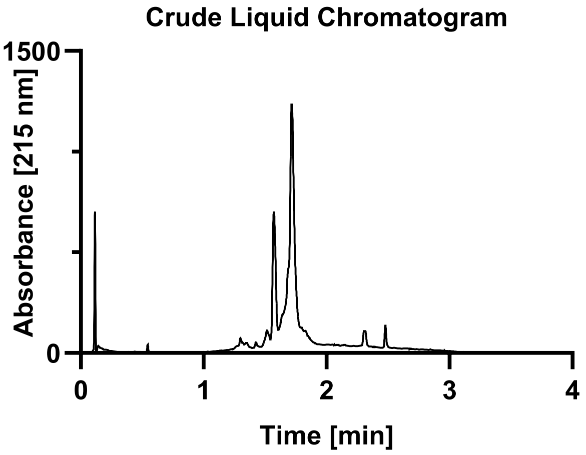

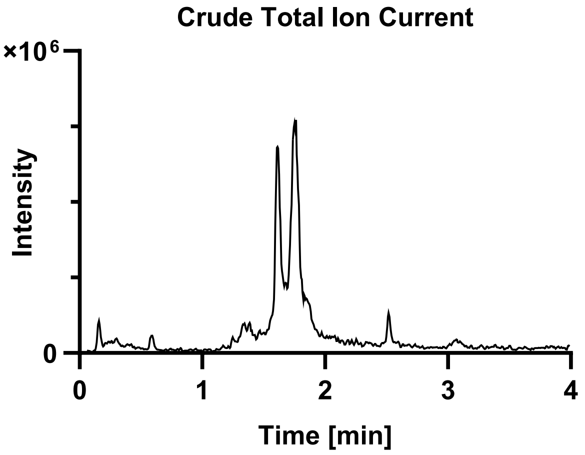

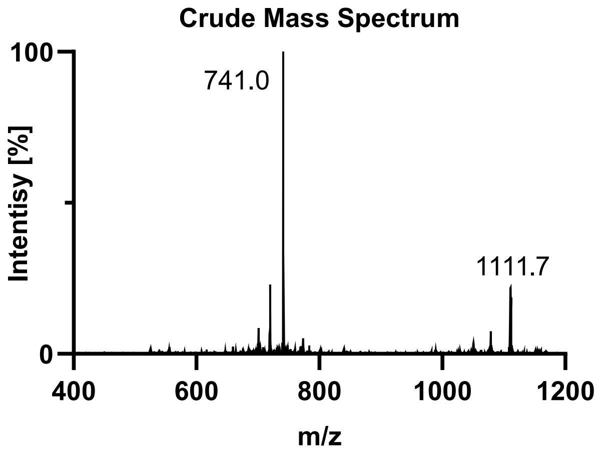

**Purification**: The material was purified over a semi-prep C4 column (10 x 250 mm, 10 µm, 300 Å, 5.0 mL/min) 18-45% MeCN/$H_2O$/0.1% TFA over 30 min. The fractions were lyophilized to afford **S3d** (93.6 mg, 8% yield).

**Pure UPLC-MS:** Measured on a HALO C18 column (3.0 x 30 mm; 2.7 µm; 90 Å; 40 °C; 1.5 mL/min) with a gradient of 5% MeCN/$H_2O$/0.1% TFA until 0.3 min, 5-95% MeCN/$H_2O$/0.1% TFA until 3.0 min, 95% MeCN/$H_2O$/0.1% TFA until 4.0 min.
**Chemical Formula:** $C_{93}H_{146}N_{25}O_{31}PSe$
**Molecular Weight:** 2220.280 g/mol
**LRMS for [M+2H+]:** $C_{93}H_{148}N_{25}O_{31}PSe^{2+}$ 1111.5, found 1110.8.
**LRMS for [M+3H+]:** $C_{93}H_{149}N_{25}O_{31}PSe^{2+}$ 741.3, found 741.0.

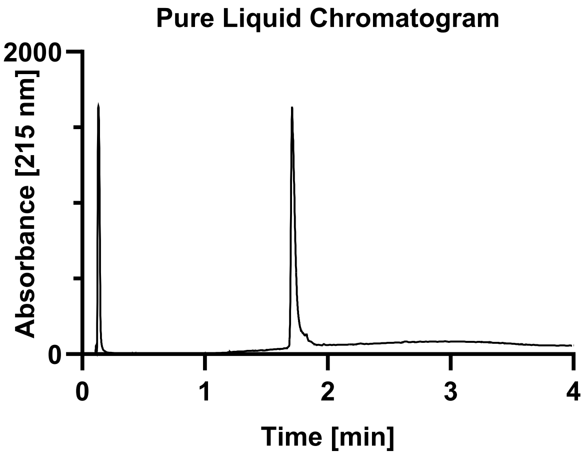

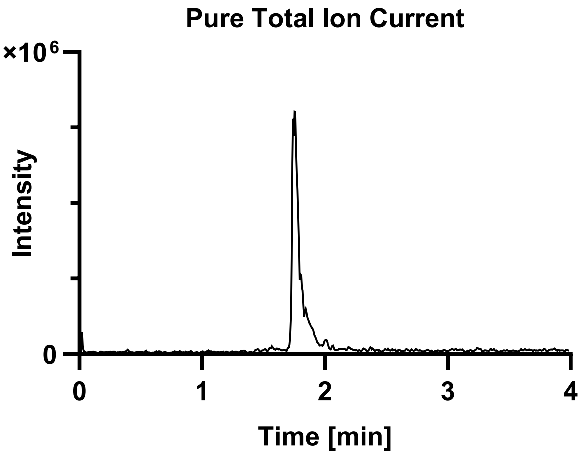

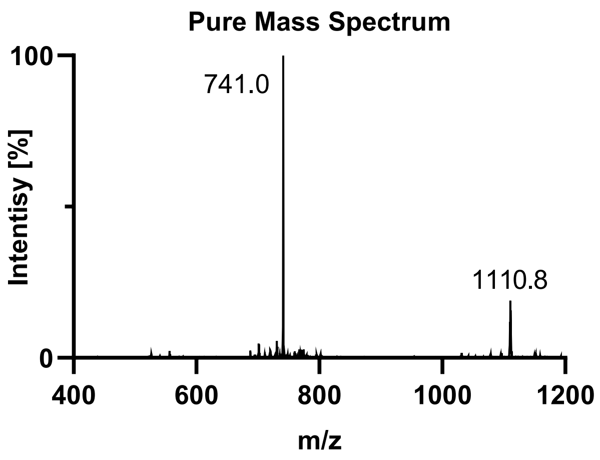

Tau(390-407) pS404 (**S3e**).

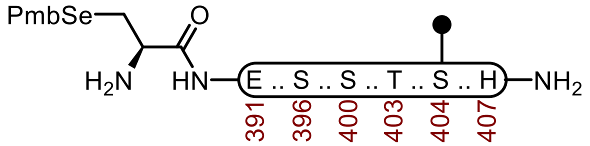

**S3e**

**Sequence:** H-SeC(Mob)EIVYKSPVVSGDTS($OPO_3HBn$)PRH-$NH_2$

**Amino Acids:** Fmoc-L-SeC(Mob)-OH, Fmoc-L-Glu(O*t*Bu)-OH, Fmoc-Ile-OH, Fmoc-Val-OH, Fmoc-Tyr(*t*Bu)-OH, Fmoc-Lys(Boc)-OH, Fmoc-Ser(*t*Bu)-OH, Fmoc-Pro-OH, Fmoc-Val-OH, Fmoc-Ser($OPO_3HBn$)-OH, Fmoc-Gly-OH, Fmoc-Thr(*t*Bu)-OH, Fmoc-Arg(Pbf)-OH, Fmoc-His(Trt)-OH.

**Resin:** Fmoc-RinkAmide Protide resin from CEM (100-200 mesh, 0.6 mmol/g, 0.833 g, 0.50 mmol) was used.

**Coupling:** Amino acids were double coupled using a preactivated solution of Fmoc-AA-OH (0.25 M, 4 equiv.), HATU (0.5 M, 4 equiv.), and DIPEA (5 equiv.) at 23 °C for 1 h. The resin was washed with DMF (2 x 35 mL)

**Phosphoserine, phosphothreonine, and Selenocysteine Coupling.** Unique amino acids were single coupled using 0.25 M amino acid in DMF (2 equiv.), 0.5 M HATU (2 equiv.), oxyma (2 equiv.) and DIPEA (7 equiv.) at 23 °C for 16 h.

**Caping:** After double coupling, the resin was capped with $Ac_2O$ (5 equiv.) and DIPEA (10 equiv.) in DMF (20 mL) for 15 min. The resin was washed with DMF (3 x 20 mL).

**Deprotection:** 20% piperidine + 0.1 M HOBt in DMF (30 mL) at 23 ºC (2 x 15 min). The resin was washed with DMF (5 x 20 mL).

**Global Deprotection:** The dry resin in a 250 mL polypropylene vial was treated with TFA:$H_2O$:thioanisole:TIPSH (85:5:5:5, 15 mL) and mixed on an overhead stirrer. The mixture was stirred at 23 ºC for 3 h before filtering, and the resin was washed with an additional portion of TFA (3 x 4 mL). The pooled filtrate was cooled to 0°C and treated with $Et_2O$ to precipitate the peptide. The precipitate was centrifuged, and the resulting pellet was washed twice more with $Et_2O$ before it was dissolved in 40% MeCN and lyophilized.

**Crude UPLC-MS:** Measured on a HALO C18 column (3.0 x 30 mm; 2.7 µm; 90 Å; 40 °C; 1.5 mL/min) with a gradient of 5% MeCN/$H_2O$/0.1% TFA until 0.3 min, 5-95% MeCN/$H_2O$/0.1% TFA until 3.0 min, 95% MeCN/$H_2O$/0.1% TFA until 4.0 min.

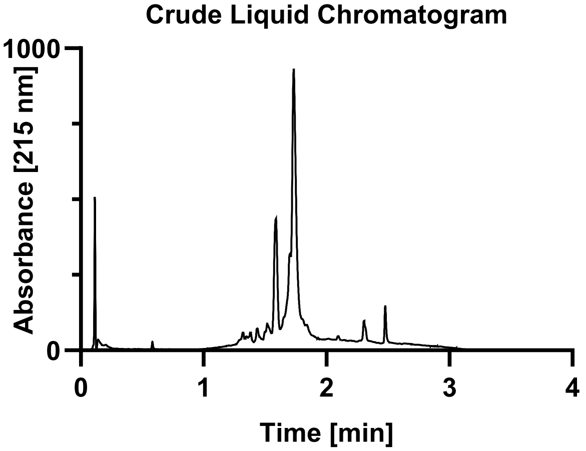

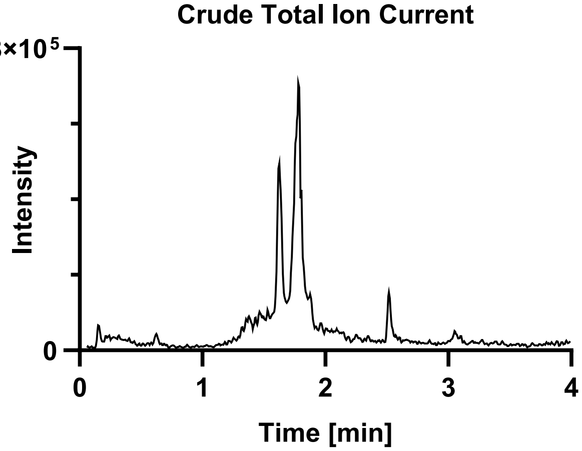

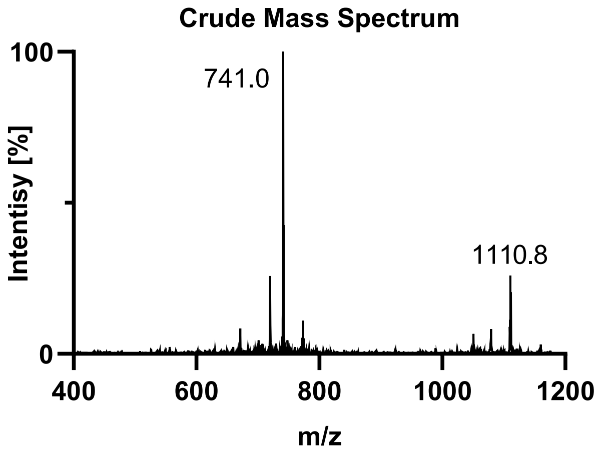

**Purification**: The material was purified over a semi-prep C4 column (10 x 250 mm, 10 µm, 300 Å, 5.0 mL/min) 18-45% MeCN/$H_2O$/0.1% TFA over 30 min. The fractions were lyophilized to afford **S3e** (74.8 mg, 33.7% yield).

**Pure UPLC-MS:** Measured on a HALO C18 column (3.0 x 30 mm; 2.7 µm; 90 Å; 40 °C; 1.5 mL/min) with a gradient of 5% MeCN/$H_2O$/0.1% TFA until 0.3 min, 5-95% MeCN/$H_2O$/0.1% TFA until 3.0 min, 95% MeCN/$H_2O$/0.1% TFA until 4.0 min.
**Chemical Formula:** $C_{93}H_{146}N_{25}O_{31}PSe$
**Molecular Weight:** 2220.280 g/mol
**LRMS for [M+2H+]:** $C_{93}H_{148}N_{25}O_{31}PSe^{2+}$ 1111.5, found 1110.9.
**LRMS for [M+3H+]:** $C_{93}H_{149}N_{25}O_{31}PSe^{2+}$ 741.3, found 741.3.

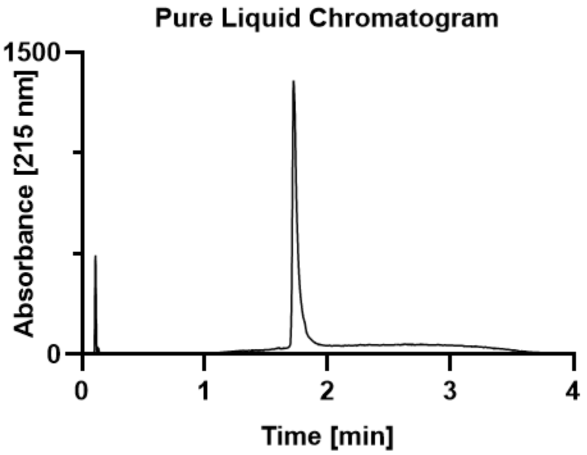

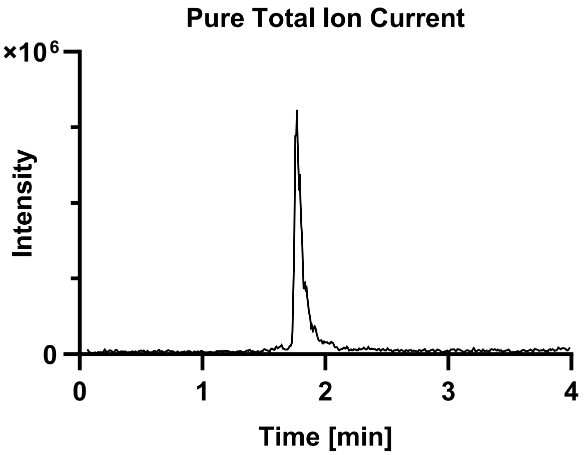

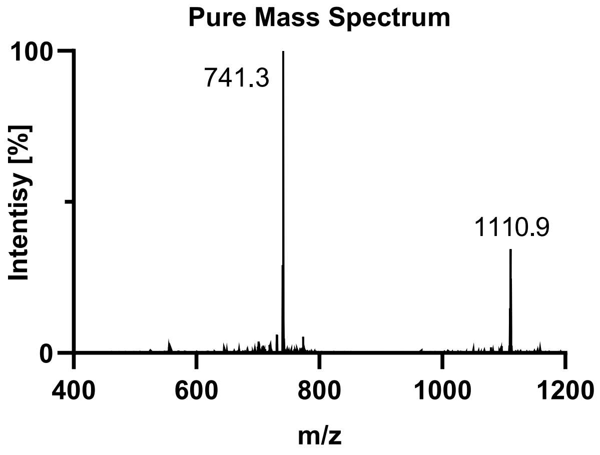

Tau(390-407) pS396 pS400 (**S3f**).

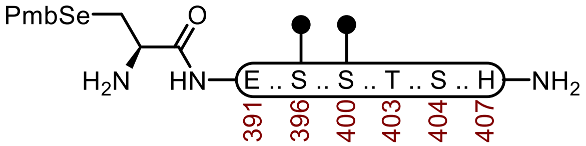

**S3f**

**Sequence:** H-C(SePmb)EIVYKS($OPO_3HBn$)PVVS($OPO_3HBn$)GDTSPRH-$NH_2$

**Amino Acids:** Fmoc-L-SeC(Mob)-OH, Fmoc-L-Glu(O*t*Bu)-OH, Fmoc-Ile-OH, Fmoc-Val-OH, Fmoc-Tyr(*t*Bu)-OH, Fmoc-Lys(Boc)-OH, Fmoc-Ser(*t*Bu)-OH, Fmoc-Pro-OH, Fmoc-Val-OH, Fmoc-Ser($OPO_3HBn$)-OH, Fmoc-Gly-OH, Fmoc-Thr(*t*Bu)-OH, Fmoc-Arg(Pbf)-OH, Fmoc-His(Trt)-OH.

**Resin:** Fmoc-RinkAmide Protide resin from CEM (100-200 mesh, 0.6 mmol/g, 0.833 g, 0.50 mmol) was used.

**Coupling:** Amino acids were double coupled using a preactivated solution of Fmoc-AA-OH (0.25 M, 4 equiv.), HATU (0.5 M, 4 equiv.), and DIPEA (5 equiv.) at 23 °C for 1 h. The resin was washed with DMF (2 x 35 mL)

**Phosphoserine, phosphothreonine, and Selenocysteine Coupling.** Unique amino acids were single coupled using 0.25 M amino acid in DMF (2 equiv.), 0.5 M HATU (2 equiv.), oxyma (2 equiv.) and DIPEA (7 equiv.) at 23 °C for 16 h.

**Caping:** After double coupling, the resin was capped with $Ac_2O$ (5 equiv.) and DIPEA (10 equiv.) in DMF (20 mL) for 15 min. The resin was washed with DMF (3 x 20 mL).

**Deprotection:** 20% piperidine + 0.1 M HOBt in DMF (30 mL) at 23 ºC (2 x 15 min). The resin was washed with DMF (5 x 20 mL).

**Global Deprotection:** The dry resin in a 250 mL polypropylene vial was treated with TFA:$H_2O$:thioanisole:TIPSH (85:5:5:5, 15 mL) and mixed on an overhead stirrer. The mixture was stirred at 23 ºC for 3 h before filtering, and the resin was washed with an additional portion of TFA (3 x 4 mL). The pooled filtrate was cooled to 0°C and treated with $Et_2O$ to precipitate the peptide. The precipitate was centrifuged, and the resulting pellet was washed twice more with $Et_2O$ before it was dissolved in 40% MeCN and lyophilized.

**Crude UPLC-MS:** Measured on a HALO C18 column (3.0 x 30 mm; 2.7 µm; 90 Å; 40 °C; 1.5 mL/min) with a gradient of 5% MeCN/$H_2O$/0.1% TFA until 0.3 min, 5-95% MeCN/$H_2O$/0.1% TFA until 3.0 min, 95% MeCN/$H_2O$/0.1% TFA until 4.0 min.

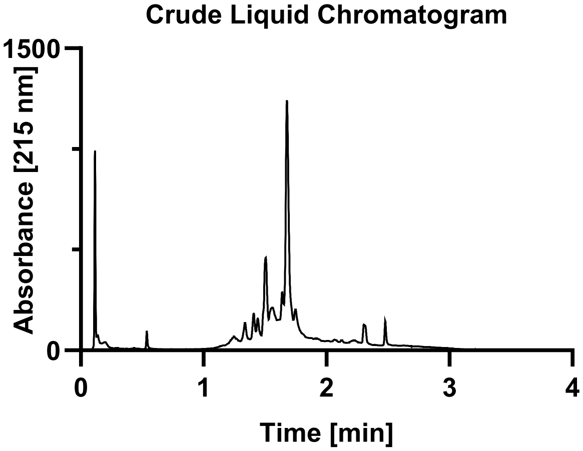

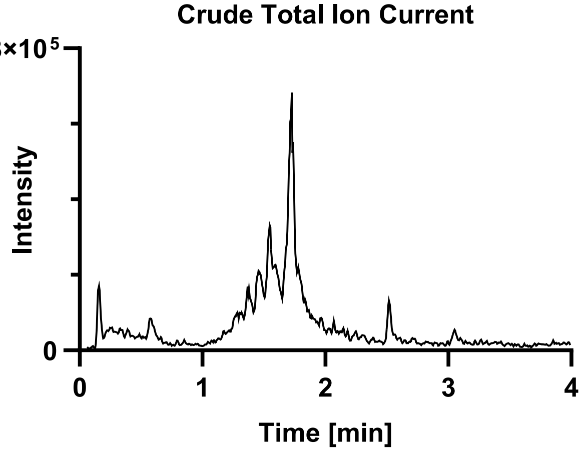

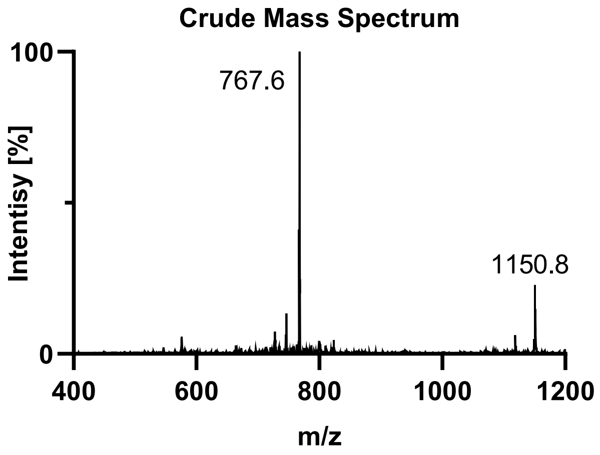

**Purification**: The material was purified over a semi-prep C4 column (10 x 250 mm, 10 µm, 300 Å, 5.0 mL/min) 18-45% MeCN/$H_2O$/0.1% TFA over 30 min. The fractions were lyophilized to afford **S3f** (76.9 mg, 7% yield).

**Pure UPLC-MS:** Measured on a HALO C18 column (3.0 x 30 mm; 2.7 µm; 90 Å; 40 °C; 1.5 mL/min) with a gradient of 5% MeCN/$H_2O$/0.1% TFA until 0.3 min, 5-95% MeCN/$H_2O$/0.1% TFA until 3.0 min, 95% MeCN/$H_2O$/0.1% TFA until 4.0 min.
**Chemical Formula:** $C_{93}H_{147}N_{25}O_{34}P_2Se$
**Molecular Weight:** 2300.259 g/mol
**LRMS for [M+2H+]:** $C_{93}H_{149}N_{25}O_{34}P_2Se^{2+}$ 1151.5, found 1151.2.
**LRMS for [M+3H+]:** $C_{93}H_{150}N_{25}O_{34}P_2Se^{3+}$ 768.0, found 767.6.

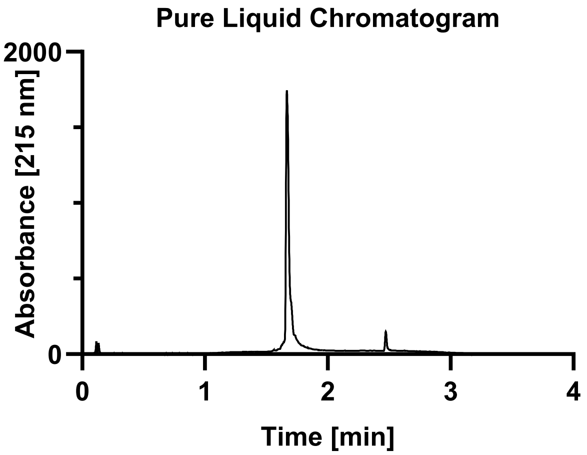

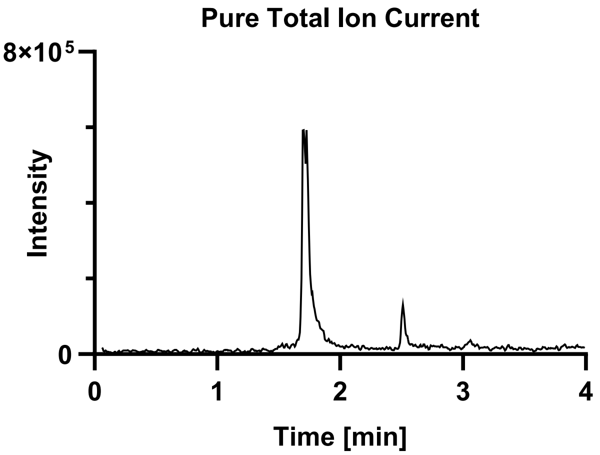

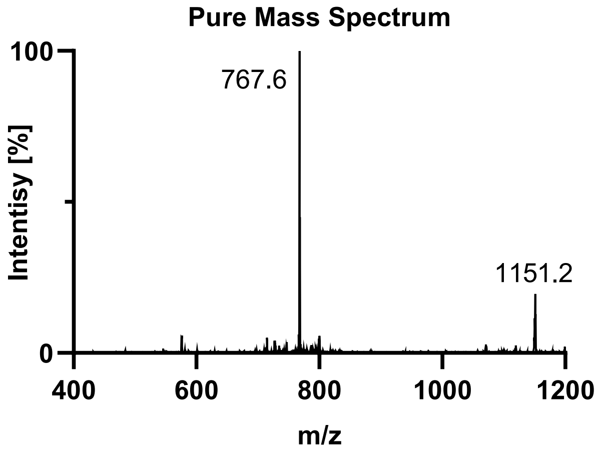

Tau(390-407) pS396 pT403 (**S3g**).

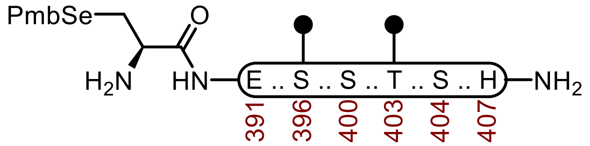

**S3g**

**Sequence:** H-C(SePmb)EIVYKS($OPO_3HBn$)PVVSGDT($OPO_3HBn$)SPRH-$NH_2$

**Amino Acids:** Fmoc-L-SeC(Mob)-OH, Fmoc-L-Glu(O*t*Bu)-OH, Fmoc-Ile-OH, Fmoc-Val-OH, Fmoc-Tyr(*t*Bu)-OH, Fmoc-Lys(Boc)-OH, Fmoc-Ser(*t*Bu)-OH, Fmoc-Pro-OH, Fmoc-Val-OH, Fmoc-Ser($OPO_3HBn$)-OH, Fmoc-Gly-OH, Fmoc-Thr(*t*Bu)-OH, Fmoc-Arg(Pbf)-OH, Fmoc-His(Trt)-OH.

**Resin:** Fmoc-RinkAmide Protide resin from CEM (100-200 mesh, 0.6 mmol/g, 0.833 g, 0.50 mmol) was used.

**Coupling:** Amino acids were double coupled using a preactivated solution of Fmoc-AA-OH (0.25 M, 4 equiv.), HATU (0.5 M, 4 equiv.), and DIPEA (5 equiv.) at 23 °C for 1 h. The resin was washed with DMF (2 x 35 mL)

**Phosphoserine, phosphothreonine, and Selenocysteine Coupling.** Unique amino acids were single coupled using 0.25 M amino acid in DMF (2 equiv.), 0.5 M HATU (2 equiv.), oxyma (2 equiv.) and DIPEA (7 equiv.) at 23 °C for 16 h.

**Caping:** After double coupling, the resin was capped with $Ac_2O$ (5 equiv.) and DIPEA (10 equiv.) in DMF (20 mL) for 15 min. The resin was washed with DMF (3 x 20 mL).

**Deprotection:** 20% piperidine + 0.1 M HOBt in DMF (30 mL) at 23 ºC (2 x 15 min). The resin was washed with DMF (5 x 20 mL).

**Global Deprotection:** The dry resin in a 250 mL polypropylene vial was treated with TFA:$H_2O$:thioanisole:TIPSH (85:5:5:5, 15 mL) and mixed on an overhead stirrer. The mixture was stirred at 23 ºC for 3 h before filtering, and the resin was washed with an additional portion of TFA (3 x 4 mL). The pooled filtrate was cooled to 0°C and treated with $Et_2O$ to precipitate the peptide. The precipitate was centrifuged, and the resulting pellet was washed twice more with $Et_2O$ before it was dissolved in 40% MeCN and lyophilized.

**Crude UPLC-MS:** Measured on a HALO C18 column (3.0 x 30 mm; 2.7 µm; 90 Å; 40 °C; 1.5 mL/min) with a gradient of 5% MeCN/$H_2O$/0.1% TFA until 0.3 min, 5-95% MeCN/$H_2O$/0.1% TFA until 3.0 min, 95% MeCN/$H_2O$/0.1% TFA until 4.0 min.

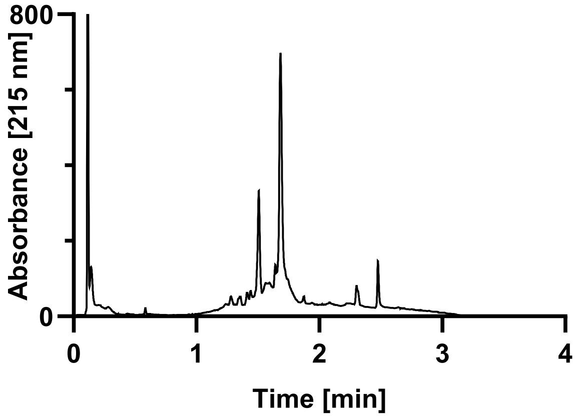

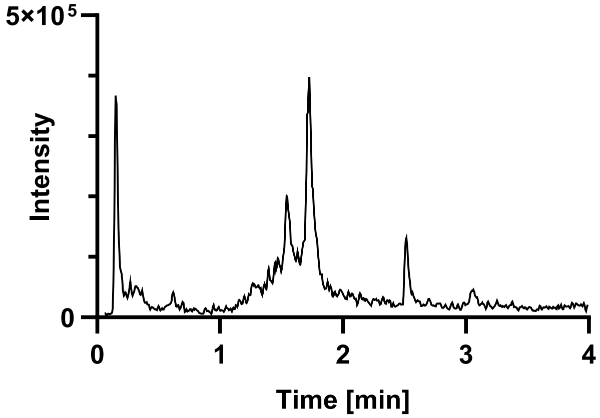

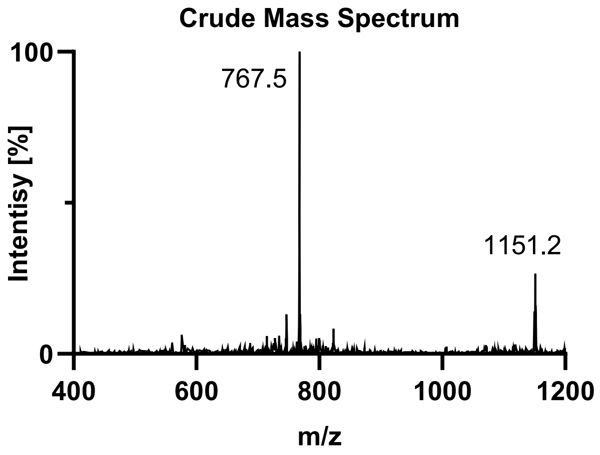

**Purification**: The material was purified over a semi-prep C4 column (10 x 250 mm, 10 µm, 300 Å, 5.0 mL/min) 18-45% MeCN/$H_2O$/0.1% TFA over 30 min. The fractions were lyophilized to afford **S3g** (45.7 mg, 4% yield).

**Pure UPLC-MS:** Measured on a HALO C18 column (3.0 x 30 mm; 2.7 µm; 90 Å; 40 °C; 1.5 mL/min) with a gradient of 5% MeCN/$H_2O$/0.1% TFA until 0.3 min, 5-95% MeCN/$H_2O$/0.1% TFA until 3.0 min, 95% MeCN/$H_2O$/0.1% TFA until 4.0 min.
**Chemical Formula:** $C_{93}H_{147}N_{25}O_{34}P_2Se$
**Molecular Weight:** 2300.259 g/mol
**LRMS for [M+2H+]:** $C_{93}H_{149}N_{25}O_{34}P_2Se^{2+}$ 1151.5, found 1151.2.
**LRMS for [M+3H+]:** $C_{93}H_{150}N_{25}O_{34}P_2Se^{3+}$ 768.0, found 767.8.

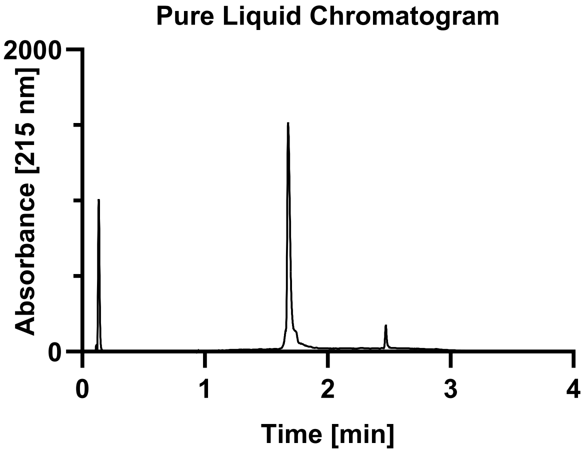

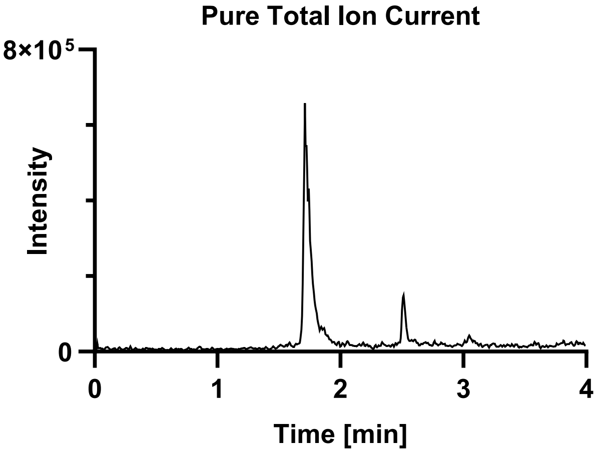

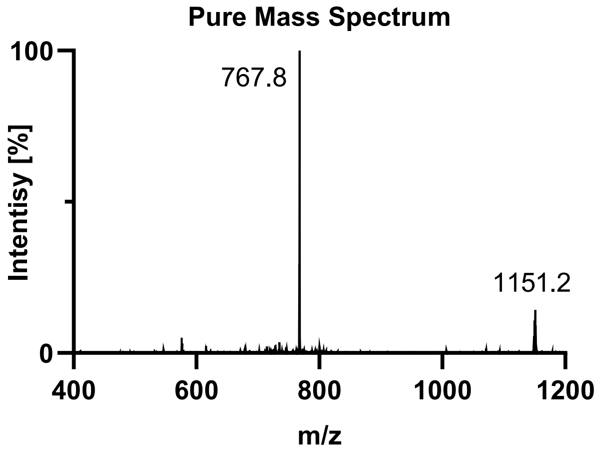

Tau(390-407) pS396 pS404 (**S3h**).

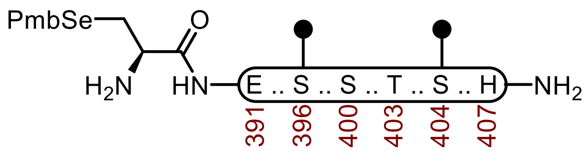

**S3h**

**Sequence:** H-C(SePmb)EIVYKS($OPO_3HBn$)PVVSGDTS($OPO_3HBn$)PRH-$NH_2$

**Amino Acids:** Fmoc-L-SeC(Mob)-OH, Fmoc-L-Glu(O*t*Bu)-OH, Fmoc-Ile-OH, Fmoc-Val-OH, Fmoc-Tyr(*t*Bu)-OH, Fmoc-Lys(Boc)-OH, Fmoc-Ser(*t*Bu)-OH, Fmoc-Pro-OH, Fmoc-Val-OH, Fmoc-Ser($OPO_3HBn$)-OH, Fmoc-Gly-OH, Fmoc-Thr(*t*Bu)-OH, Fmoc-Arg(Pbf)-OH, Fmoc-His(Trt)-OH.

**Resin:** Fmoc-RinkAmide Protide resin from CEM (100-200 mesh, 0.6 mmol/g, 0.833 g, 0.50 mmol) was used.

**Coupling:** Amino acids were double coupled using a preactivated solution of Fmoc-AA-OH (0.25 M, 4 equiv.), HATU (0.5 M, 4 equiv.), and DIPEA (5 equiv.) at 23 °C for 1 h. The resin was washed with DMF (2 x 35 mL)

**Phosphoserine, phosphothreonine, and Selenocysteine Coupling.** Unique amino acids were single coupled using 0.25 M amino acid in DMF (2 equiv.), 0.5 M HATU (2 equiv.), oxyma (2 equiv.) and DIPEA (7 equiv.) at 23 °C for 16 h.

**Caping:** After double coupling, the resin was capped with $Ac_2O$ (5 equiv.) and DIPEA (10 equiv.) in DMF (20 mL) for 15 min. The resin was washed with DMF (3 x 20 mL).

**Deprotection:** 20% piperidine + 0.1 M HOBt in DMF (30 mL) at 23 ºC (2 x 15 min). The resin was washed with DMF (5 x 20 mL).

**Global Deprotection:** The dry resin in a 250 mL polypropylene vial was treated with TFA:$H_2O$:thioanisole:TIPSH (85:5:5:5, 15 mL) and mixed on an overhead stirrer. The mixture was stirred at 23 ºC for 3 h before filtering, and the resin was washed with an additional portion of TFA (3 x 4 mL). The pooled filtrate was cooled to 0°C and treated with $Et_2O$ to precipitate the peptide. The precipitate was centrifuged, and the resulting pellet was washed twice more with $Et_2O$ before it was dissolved in 40% MeCN and lyophilized.

**Crude UPLC-MS:** Measured on a HALO C18 column (3.0 x 30 mm; 2.7 µm; 90 Å; 40 °C; 1.5 mL/min) with a gradient of 5% MeCN/$H_2O$/0.1% TFA until 0.3 min, 5-95% MeCN/$H_2O$/0.1% TFA until 3.0 min, 95% MeCN/$H_2O$/0.1% TFA until 4.0 min.

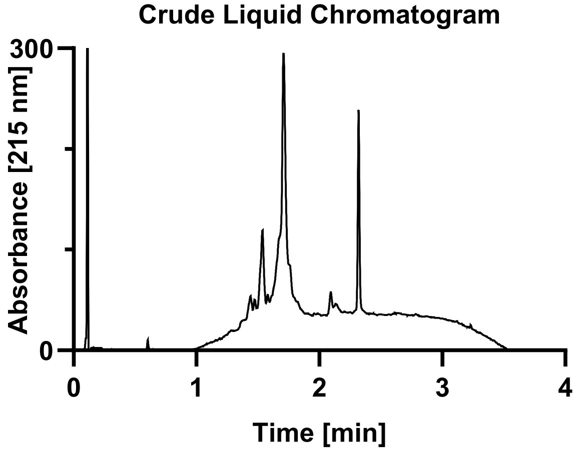

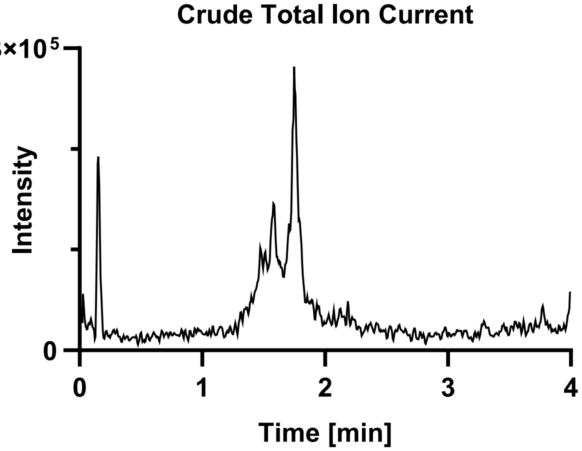

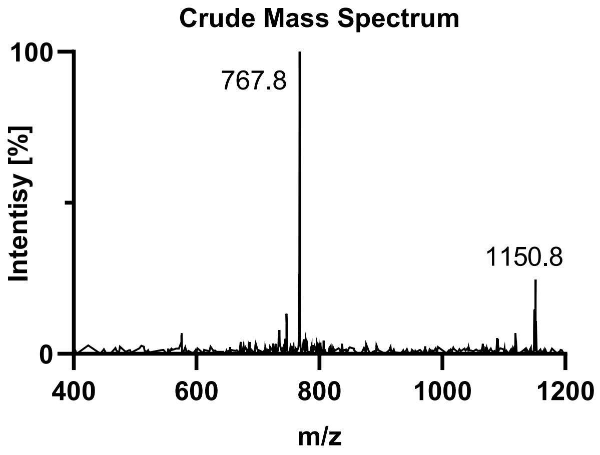

**Purification**: The material was purified over a semi-prep C4 column (10 x 250 mm, 10 µm, 300 Å, 5.0 mL/min) 18-45% MeCN/$H_2O$/0.1% TFA over 30 min. The fractions were lyophilized to afford **S3h** (51.9 mg, 5% yield).

**Pure UPLC-MS:** Measured on a HALO C18 column (3.0 x 30 mm; 2.7 µm; 90 Å; 40 °C; 1.5 mL/min) with a gradient of 5% MeCN/$H_2O$/0.1% TFA until 0.3 min, 5-95% MeCN/$H_2O$/0.1% TFA until 3.0 min, 95% MeCN/$H_2O$/0.1% TFA until 4.0 min.
**Chemical Formula:** $C_{93}H_{147}N_{25}O_{34}P_2Se$
**Molecular Weight:** 2300.259 g/mol
**LRMS for [M+2H+]:** $C_{93}H_{149}N_{25}O_{34}P_2Se^{2+}$ 1151.5, found 1150.9.
**LRMS for [M+3H+]:** $C_{93}H_{150}N_{25}O_{34}P_2Se^{3+}$ 768.0, found 767.8.

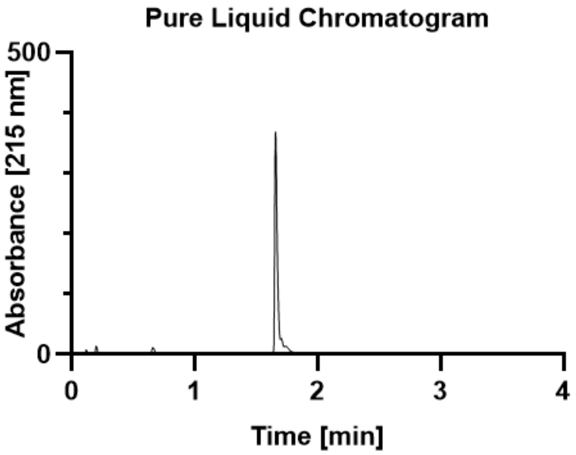

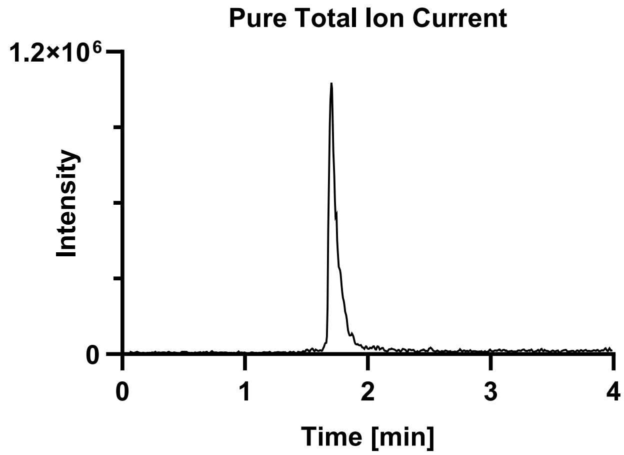

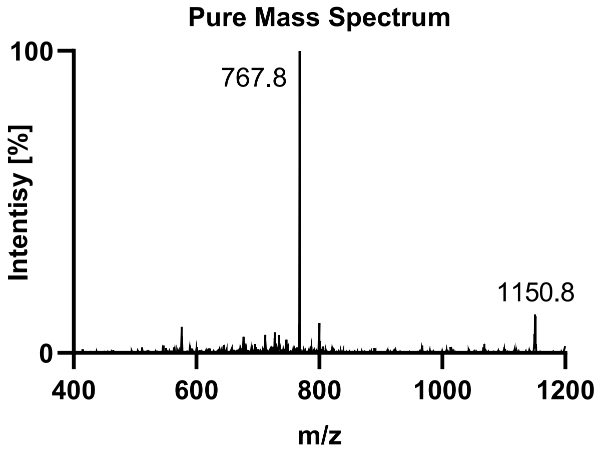

Tau(390-407) pS400 pT403 (**S3i**).

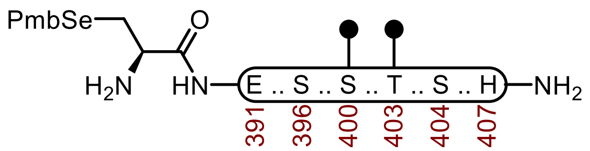

**S3i**

**Sequence:** H-C(SePmb)EIVYKSPVVS($OPO_3HBn$)GDT($OPO_3HBn$)SPRH-$NH_2$

**Amino Acids:** Fmoc-L-SeC(Mob)-OH, Fmoc-L-Glu(O*t*Bu)-OH, Fmoc-Ile-OH, Fmoc-Val-OH, Fmoc-Tyr(*t*Bu)-OH, Fmoc-Lys(Boc)-OH, Fmoc-Ser(*t*Bu)-OH, Fmoc-Pro-OH, Fmoc-Val-OH, Fmoc-Ser($OPO_3HBn$)-OH, Fmoc-Gly-OH, Fmoc-Thr(*t*Bu)-OH, Fmoc-Arg(Pbf)-OH, Fmoc-His(Trt)-OH.

**Resin:** Fmoc-RinkAmide Protide resin from CEM (100-200 mesh, 0.6 mmol/g, 0.833 g, 0.50 mmol) was used.

**Coupling:** Amino acids were double coupled using a preactivated solution of Fmoc-AA-OH (0.25 M, 4 equiv.), HATU (0.5 M, 4 equiv.), and DIPEA (5 equiv.) at 23 °C for 1 h. The resin was washed with DMF (2 x 35 mL)

**Phosphoserine, phosphothreonine, and Selenocysteine Coupling.** Unique amino acids were single coupled using 0.25 M amino acid in DMF (2 equiv.), 0.5 M HATU (2 equiv.), oxyma (2 equiv.) and DIPEA (7 equiv.) at 23 °C for 16 h.

**Caping:** After double coupling, the resin was capped with $Ac_2O$ (5 equiv.) and DIPEA (10 equiv.) in DMF (20 mL) for 15 min. The resin was washed with DMF (3 x 20 mL).

**Deprotection:** 20% piperidine + 0.1 M HOBt in DMF (30 mL) at 23 ºC (2 x 15 min). The resin was washed with DMF (5 x 20 mL).

**Global Deprotection:** The dry resin in a 250 mL polypropylene vial was treated with TFA:$H_2O$:thioanisole:TIPSH (85:5:5:5, 15 mL) and mixed on an overhead stirrer. The mixture was stirred at 23 ºC for 3 h before filtering, and the resin was washed with an additional portion of TFA (3 x 4 mL). The pooled filtrate was cooled to 0°C and treated with $Et_2O$ to precipitate the peptide. The precipitate was centrifuged, and the resulting pellet was washed twice more with $Et_2O$ before it was dissolved in 40% MeCN and lyophilized.

**Crude UPLC-MS:** Measured on a HALO C18 column (3.0 x 30 mm; 2.7 µm; 90 Å; 40 °C; 1.5 mL/min) with a gradient of 5% MeCN/$H_2O$/0.1% TFA until 0.3 min, 5-95% MeCN/$H_2O$/0.1% TFA until 3.0 min, 95% MeCN/$H_2O$/0.1% TFA until 4.0 min.

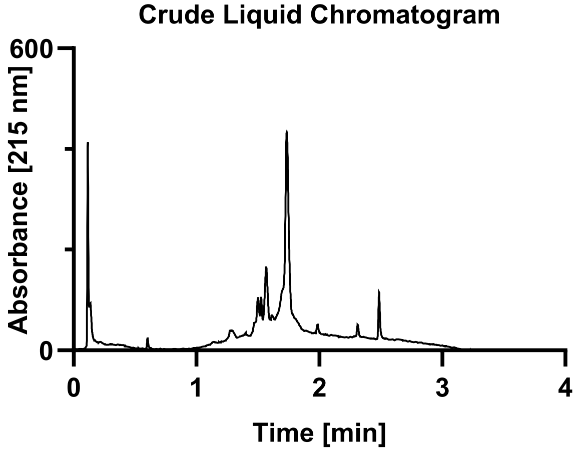

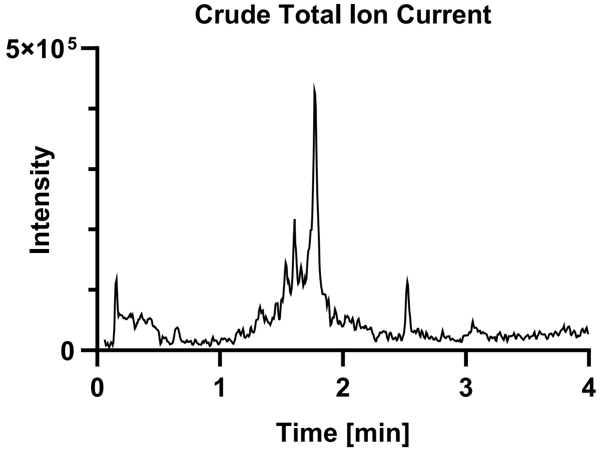

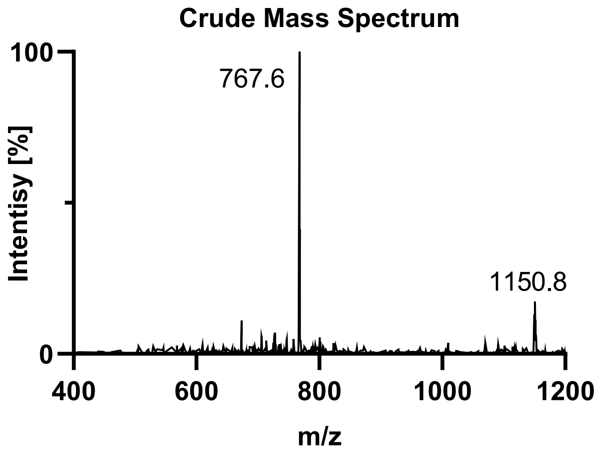

**Purification**: The material was purified over a semi-prep C4 column (10 x 250 mm, 10 µm, 300 Å, 5.0 mL/min) 18-45% MeCN/$H_2O$/0.1% TFA over 30 min. The fractions were lyophilized to afford **S3i** (56.7 mg, 4% yield).

**Pure UPLC-MS:** Measured on a HALO C18 column (3.0 x 30 mm; 2.7 µm; 90 Å; 40 °C; 1.5 mL/min) with a gradient of 5% MeCN/$H_2O$/0.1% TFA until 0.3 min, 5-95% MeCN/$H_2O$/0.1% TFA until 3.0 min, 95% MeCN/$H_2O$/0.1% TFA until 4.0 min.
**Chemical Formula:** $C_{93}H_{147}N_{25}O_{34}P_2Se$
**Molecular Weight:** 2300.259 g/mol
**LRMS for [M+2H+]:** $C_{93}H_{149}N_{25}O_{34}P_2Se^{2+}$ 1151.5, found 1150.8.
**LRMS for [M+3H+]:** $C_{93}H_{150}N_{25}O_{34}P_2Se^{3+}$ 768.0, found 767.8.

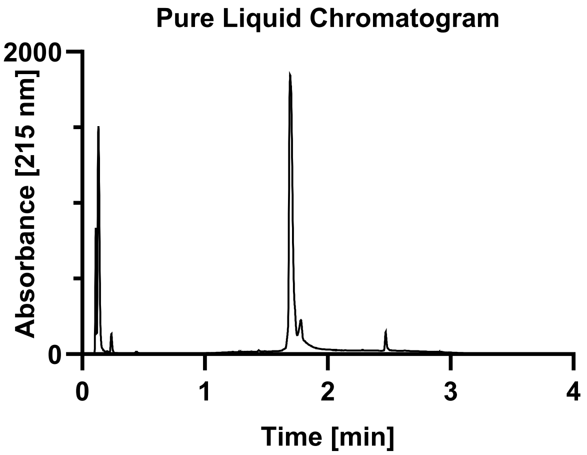

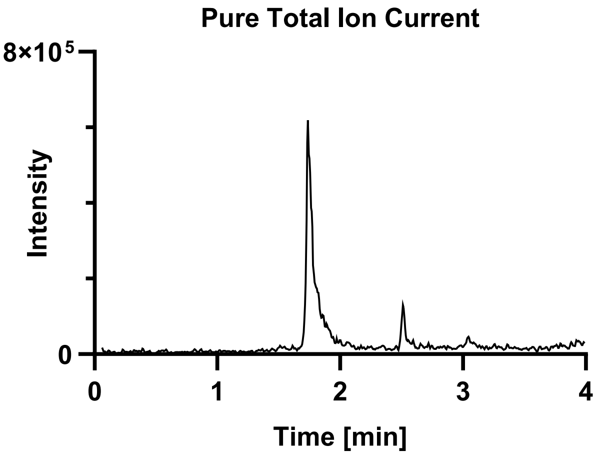

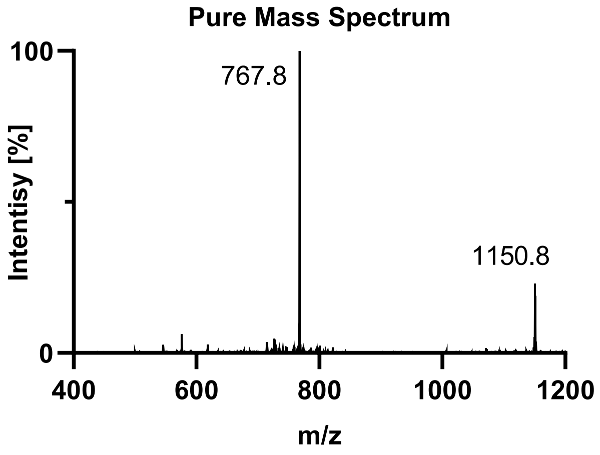

Tau(390-407) pS400 pS404 (**S3j**).

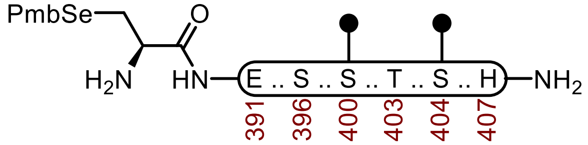

**S3j**

**Sequence:** H-C(SePmb)EIVYKSPVVS($OPO_3HBn$)GDTS($OPO_3HBn$)PRH-$NH_2$

**Amino Acids:** Fmoc-L-SeC(Mob)-OH, Fmoc-L-Glu(O*t*Bu)-OH, Fmoc-Ile-OH, Fmoc-Val-OH, Fmoc-Tyr(*t*Bu)-OH, Fmoc-Lys(Boc)-OH, Fmoc-Ser(*t*Bu)-OH, Fmoc-Pro-OH, Fmoc-Val-OH, Fmoc-Ser($OPO_3HBn$)-OH, Fmoc-Gly-OH, Fmoc-Thr(*t*Bu)-OH, Fmoc-Arg(Pbf)-OH, Fmoc-His(Trt)-OH.

**Resin:** Fmoc-RinkAmide Protide resin from CEM (100-200 mesh, 0.6 mmol/g, 0.833 g, 0.50 mmol) was used.

**Coupling:** Amino acids were double coupled using a preactivated solution of Fmoc-AA-OH (0.25 M, 4 equiv.), HATU (0.5 M, 4 equiv.), and DIPEA (5 equiv.) at 23 °C for 1 h. The resin was washed with DMF (2 x 35 mL)

**Phosphoserine, phosphothreonine, and Selenocysteine Coupling.** Unique amino acids were single coupled using 0.25 M amino acid in DMF (2 equiv.), 0.5 M HATU (2 equiv.), oxyma (2 equiv.) and DIPEA (7 equiv.) at 23 °C for 16 h.

**Caping:** After double coupling, the resin was capped with $Ac_2O$ (5 equiv.) and DIPEA (10 equiv.) in DMF (20 mL) for 15 min. The resin was washed with DMF (3 x 20 mL).

**Deprotection:** 20% piperidine + 0.1 M HOBt in DMF (30 mL) at 23 ºC (2 x 15 min). The resin was washed with DMF (5 x 20 mL).

**Global Deprotection:** The dry resin in a 250 mL polypropylene vial was treated with TFA:$H_2O$:thioanisole:TIPSH (85:5:5:5, 15 mL) and mixed on an overhead stirrer. The mixture was stirred at 23 ºC for 3 h before filtering, and the resin was washed with an additional portion of TFA (3 x 4 mL). The pooled filtrate was cooled to 0°C and treated with $Et_2O$ to precipitate the peptide. The precipitate was centrifuged, and the resulting pellet was washed twice more with $Et_2O$ before it was dissolved in 40% MeCN and lyophilized.

**Crude UPLC-MS:** Measured on a HALO C18 column (3.0 x 30 mm; 2.7 µm; 90 Å; 40 °C; 1.5 mL/min) with a gradient of 5% MeCN/$H_2O$/0.1% TFA until 0.3 min, 5-95% MeCN/$H_2O$/0.1% TFA until 3.0 min, 95% MeCN/$H_2O$/0.1% TFA until 4.0 min.

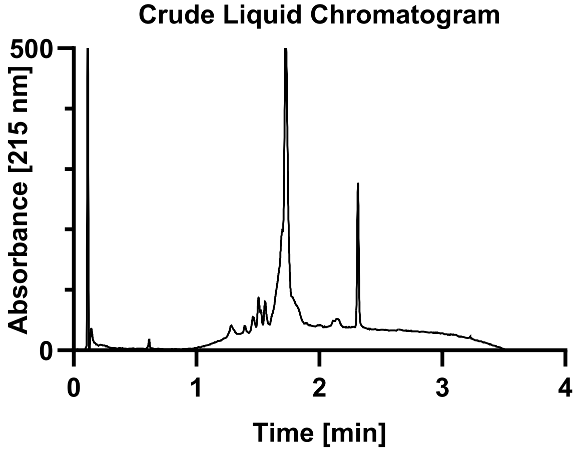

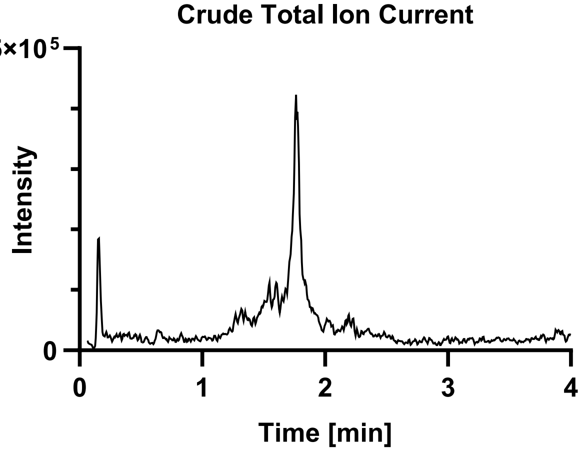

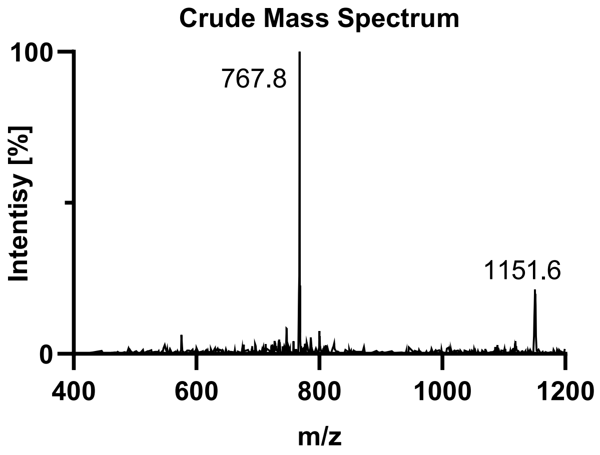

**Purification**: The material was purified over a semi-prep C4 column (10 x 250 mm, 10 µm, 300 Å, 5.0 mL/min) 18-45% MeCN/$H_2O$/0.1% TFA over 30 min. The fractions were lyophilized to afford **S3j** (54.2 mg, 5% yield).

**Pure UPLC-MS:** Measured on a HALO C18 column (3.0 x 30 mm; 2.7 µm; 90 Å; 40 °C; 1.5 mL/min) with a gradient of 5% MeCN/$H_2O$/0.1% TFA until 0.3 min, 5-95% MeCN/$H_2O$/0.1% TFA until 3.0 min, 95% MeCN/$H_2O$/0.1% TFA until 4.0 min.
**Chemical Formula:** $C_{93}H_{147}N_{25}O_{34}P_2Se$
**Molecular Weight:** 2300.259 g/mol
**LRMS for [M+2H+]:** $C_{93}H_{149}N_{25}O_{34}P_2Se^{2+}$ 1151.5, found 1151.6.
**LRMS for [M+3H+]:** $C_{93}H_{150}N_{25}O_{34}P_2Se^{3+}$ 768.0, found 767.8.

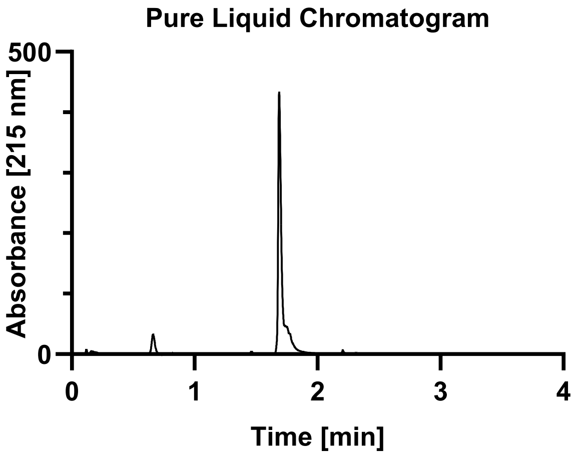

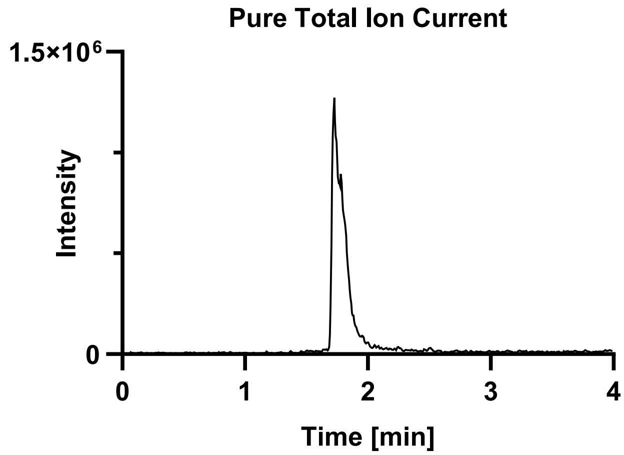

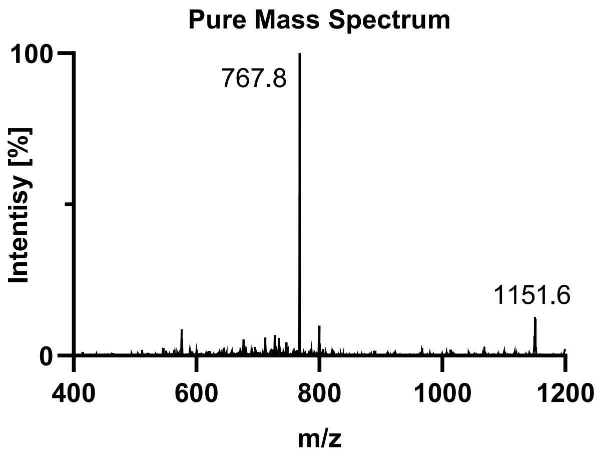

Tau(390-407) pT403 pS404 (**S3k**).

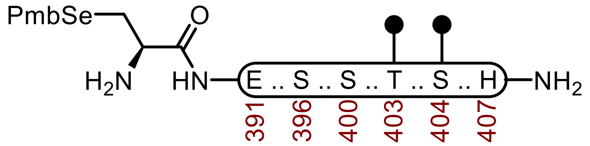

**S3k**

**Sequence:** H-C(SePmb)EIVYKSPVVSGDT($OPO_3HBn$)S($OPO_3HBn$)PRH-$NH_2$

**Amino Acids:** Fmoc-L-SeC(Mob)-OH, Fmoc-L-Glu(O*t*Bu)-OH, Fmoc-Ile-OH, Fmoc-Val-OH, Fmoc-Tyr(*t*Bu)-OH, Fmoc-Lys(Boc)-OH, Fmoc-Ser(*t*Bu)-OH, Fmoc-Pro-OH, Fmoc-Val-OH, Fmoc-Ser($OPO_3HBn$)-OH, Fmoc-Gly-OH, Fmoc-Thr(*t*Bu)-OH, Fmoc-Arg(Pbf)-OH, Fmoc-His(Trt)-OH.

**Resin:** Fmoc-RinkAmide Protide resin from CEM (100-200 mesh, 0.6 mmol/g, 0.833 g, 0.50 mmol) was used.

**Coupling:** Amino acids were double coupled using a preactivated solution of Fmoc-AA-OH (0.25 M, 4 equiv.), HATU (0.5 M, 4 equiv.), and DIPEA (5 equiv.) at 23 °C for 1 h. The resin was washed with DMF (2 x 35 mL)

**Phosphoserine, phosphothreonine, and Selenocysteine Coupling.** Unique amino acids were single coupled using 0.25 M amino acid in DMF (2 equiv.), 0.5 M HATU (2 equiv.), oxyma (2 equiv.) and DIPEA (7 equiv.) at 23 °C for 16 h.

**Caping:** After double coupling, the resin was capped with $Ac_2O$ (5 equiv.) and DIPEA (10 equiv.) in DMF (20 mL) for 15 min. The resin was washed with DMF (3 x 20 mL).

**Deprotection:** 20% piperidine + 0.1 M HOBt in DMF (30 mL) at 23 ºC (2 x 15 min). The resin was washed with DMF (5 x 20 mL).

**Global Deprotection:** The dry resin in a 250 mL polypropylene vial was treated with TFA:$H_2O$:thioanisole:TIPSH (85:5:5:5, 15 mL) and mixed on an overhead stirrer. The mixture was stirred at 23 ºC for 3 h before filtering, and the resin was washed with an additional portion of TFA (3 x 4 mL). The pooled filtrate was cooled to 0°C and treated with $Et_2O$ to precipitate the peptide. The precipitate was centrifuged, and the resulting pellet was washed twice more with $Et_2O$ before it was dissolved in 40% MeCN and lyophilized.

**Crude UPLC-MS:** Measured on a HALO C18 column (3.0 x 30 mm; 2.7 µm; 90 Å; 40 °C; 1.5 mL/min) with a gradient of 5% MeCN/$H_2O$/0.1% TFA until 0.3 min, 5-95% MeCN/$H_2O$/0.1% TFA until 3.0 min, 95% MeCN/$H_2O$/0.1% TFA until 4.0 min.

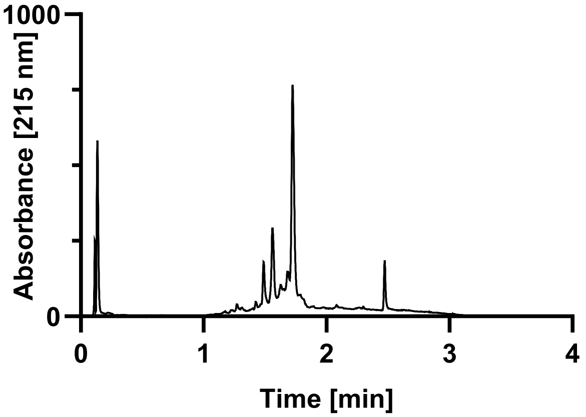

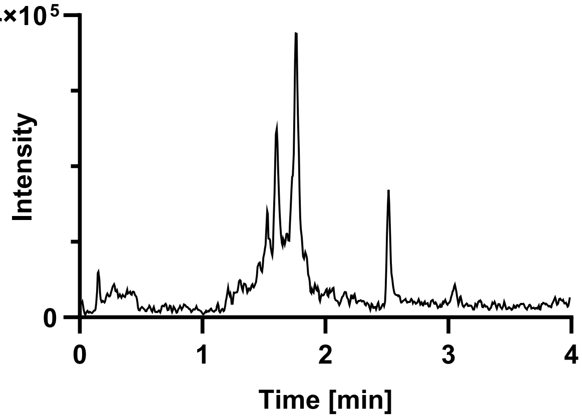

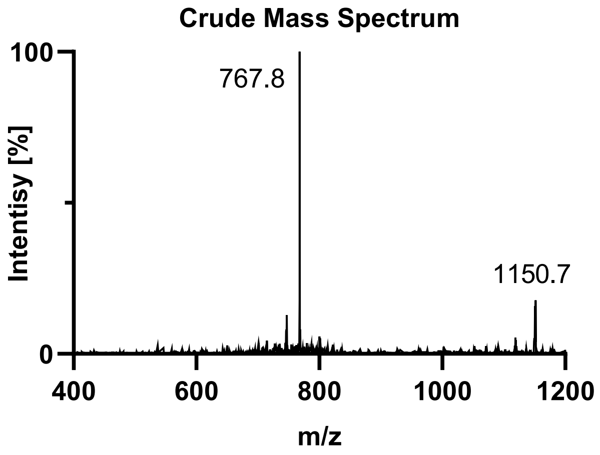

**Purification**: The material was purified over a semi-prep C4 column (10 x 250 mm, 10 µm, 300 Å, 5.0 mL/min) 18-45% MeCN/$H_2O$/0.1% TFA over 30 min. The fractions were lyophilized to afford **S3k** (110 mg, 10% yield).

**Pure UPLC-MS:** Measured on a HALO C18 column (3.0 x 30 mm; 2.7 µm; 90 Å; 40 °C; 1.5 mL/min) with a gradient of 5% MeCN/$H_2O$/0.1% TFA until 0.3 min, 5-95% MeCN/$H_2O$/0.1% TFA until 3.0 min, 95% MeCN/$H_2O$/0.1% TFA until 4.0 min.
**Chemical Formula:** $C_{93}H_{147}N_{25}O_{34}P_2Se$
**Molecular Weight:** 2300.259 g/mol
**LRMS for [M+2H+]:** $C_{93}H_{149}N_{25}O_{34}P_2Se^{2+}$ 1151.5, found 1151.2.
**LRMS for [M+3H+]:** $C_{93}H_{150}N_{25}O_{34}P_2Se^{3+}$ 768.0, found 767.7.

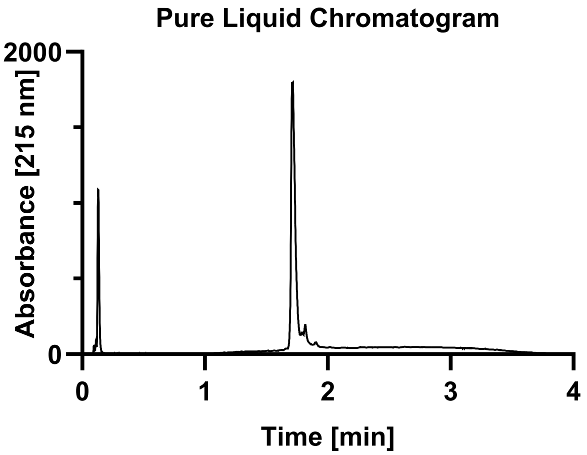

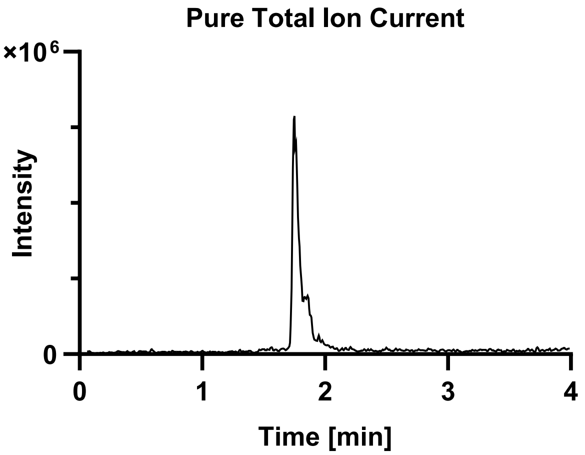

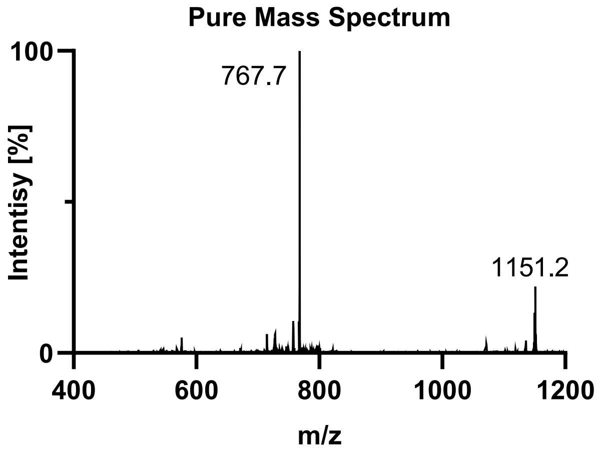

Tau(390-407) pS396 pS400 pT403 (**S3I**).

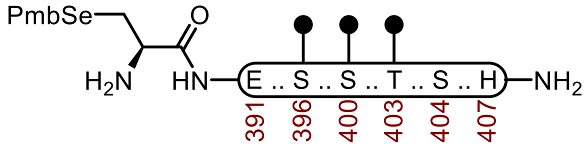

**S3I**

**Sequence:** H-C(SePmb)EIVYKS($OPO_3HBn$)PVVS($OPO_3HBn$)GDT($OPO_3HBn$)SPRH-$NH_2$

**Amino Acids:** Fmoc-L-SeC(Mob)-OH, Fmoc-L-Glu(O*t*Bu)-OH, Fmoc-Ile-OH, Fmoc-Val-OH, Fmoc-Tyr(*t*Bu)-OH, Fmoc-Lys(Boc)-OH, Fmoc-Ser(*t*Bu)-OH, Fmoc-Pro-OH, Fmoc-Val-OH, Fmoc-Ser($OPO_3HBn$)-OH, Fmoc-Gly-OH, Fmoc-Thr(*t*Bu)-OH, Fmoc-Arg(Pbf)-OH, Fmoc-His(Trt)-OH.

**Resin:** Fmoc-RinkAmide Protide resin from CEM (100-200 mesh, 0.6 mmol/g, 0.833 g, 0.50 mmol) was used.

**Coupling:** Amino acids were double coupled using a preactivated solution of Fmoc-AA-OH (0.25 M, 4 equiv.), HATU (0.5 M, 4 equiv.), and DIPEA (5 equiv.) at 23 °C for 1 h. The resin was washed with DMF (2 x 35 mL)

**Phosphoserine, phosphothreonine, and Selenocysteine Coupling.** Unique amino acids were single coupled using 0.25 M amino acid in DMF (2 equiv.), 0.5 M HATU (2 equiv.), oxyma (2 equiv.) and DIPEA (7 equiv.) at 23 °C for 16 h.

**Caping:** After double coupling, the resin was capped with $Ac_2O$ (5 equiv.) and DIPEA (10 equiv.) in DMF (20 mL) for 15 min. The resin was washed with DMF (3 x 20 mL).

**Deprotection:** 20% piperidine + 0.1 M HOBt in DMF (30 mL) at 23 ºC (2 x 15 min). The resin was washed with DMF (5 x 20 mL).

**Global Deprotection:** The dry resin in a 250 mL polypropylene vial was treated with TFA:$H_2O$:thioanisole:TIPSH (85:5:5:5, 15 mL) and mixed on an overhead stirrer. The mixture was stirred at 23 ºC for 3 h before filtering, and the resin was washed with an additional portion of TFA (3 x 4 mL). The pooled filtrate was cooled to 0°C and treated with $Et_2O$ to precipitate the peptide. The precipitate was centrifuged, and the resulting pellet was washed twice more with $Et_2O$ before it was dissolved in 40% MeCN and lyophilized.

**Crude UPLC-MS:** Measured on a HALO C18 column (3.0 x 30 mm; 2.7 µm; 90 Å; 40 °C; 1.5 mL/min) with a gradient of 5% MeCN/$H_2O$/0.1% TFA until 0.3 min, 5-95% MeCN/$H_2O$/0.1% TFA until 3.0 min, 95% MeCN/$H_2O$/0.1% TFA until 4.0 min.

**Crude Liquid Chromatogram**

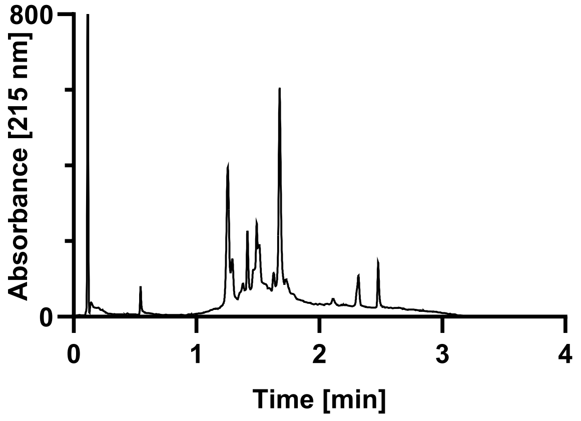

**Crude Total Ion Current**

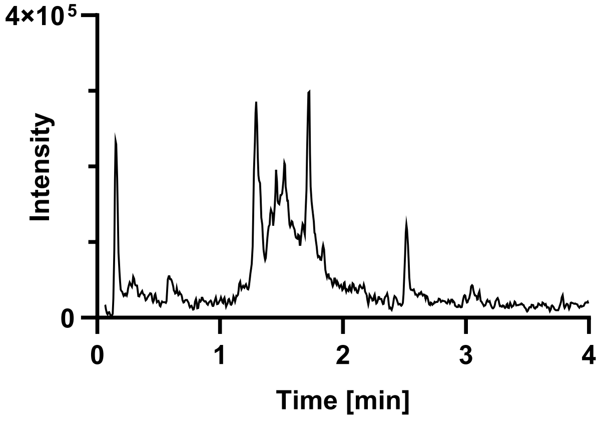

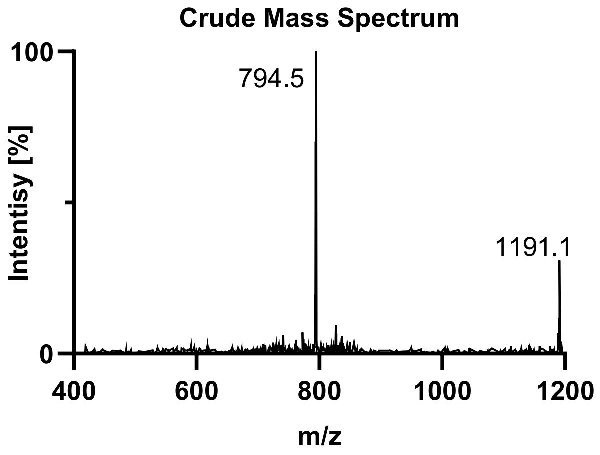

**Purification**: The material was purified over a semi-prep C4 column (10 x 250 mm, 10 µm, 300 Å, 5.0 mL/min) 18-45% MeCN/$H_2O$/0.1% TFA over 30 min. The fractions were lyophilized to afford **S3l** (38.9 mg, 3% yield).

**Pure UPLC-MS:** Measured on a HALO C18 column (3.0 x 30 mm; 2.7 µm; 90 Å; 40 °C; 1.5 mL/min) with a gradient of 5% MeCN/$H_2O$/0.1% TFA until 0.3 min, 5-95% MeCN/$H_2O$/0.1% TFA until 3.0 min, 95% MeCN/$H_2O$/0.1% TFA until 4.0 min.
**Chemical Formula:** $C_{93}H_{148}N_{25}O_{37}P_3Se$
**Molecular Weight:** 2380.237 g/mol
**LRMS for [M+2H+]:** $C_{93}H_{149}N_{25}O_{34}P_2Se^{2+}$ 1191.5, found 1190.4.
**LRMS for [M+3H+]:** $C_{93}H_{150}N_{25}O_{34}P_2Se^{3+}$ 794.6, found 794.5.

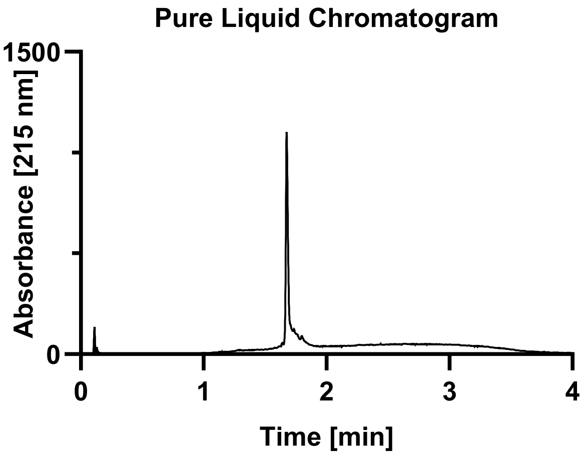

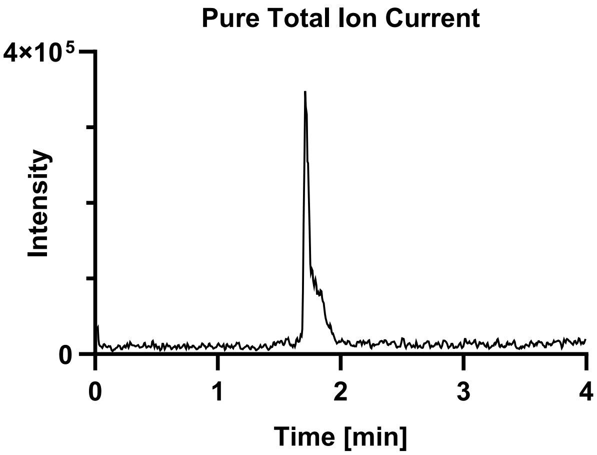

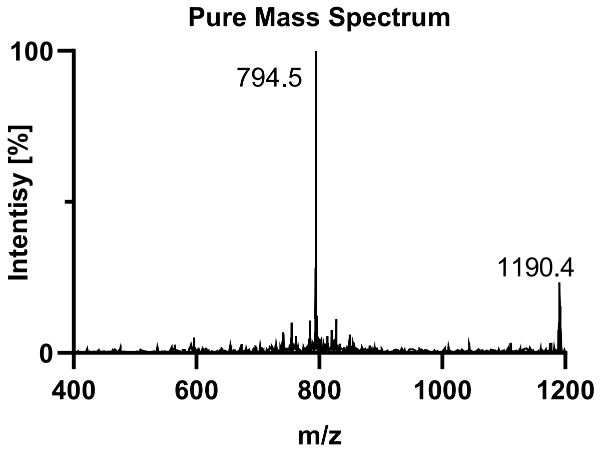

Tau(390-407) pS396 pS400 pS404 (**S3m**).

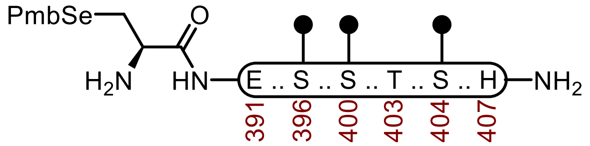

**S3m**

**Sequence:** H-C(SePmb)EIVYKS($OPO_3HBn$)PVVS($OPO_3HBn$)GDTS($OPO_3HBn$)PRH-$NH_2$

**Amino Acids:** Fmoc-L-SeC(Mob)-OH, Fmoc-L-Glu(O*t*Bu)-OH, Fmoc-Ile-OH, Fmoc-Val-OH, Fmoc-Tyr(*t*Bu)-OH, Fmoc-Lys(Boc)-OH, Fmoc-Ser(*t*Bu)-OH, Fmoc-Pro-OH, Fmoc-Val-OH, Fmoc-Ser($OPO_3HBn$)-OH, Fmoc-Gly-OH, Fmoc-Thr(*t*Bu)-OH, Fmoc-Arg(Pbf)-OH, Fmoc-His(Trt)-OH.

**Resin:** Fmoc-RinkAmide Protide resin from CEM (100-200 mesh, 0.6 mmol/g, 0.833 g, 0.50 mmol) was used.

**Coupling:** Amino acids were double coupled using a preactivated solution of Fmoc-AA-OH (0.25 M, 4 equiv.), HATU (0.5 M, 4 equiv.), and DIPEA (5 equiv.) at 23 °C for 1 h. The resin was washed with DMF (2 x 35 mL)

**Phosphoserine, phosphothreonine, and Selenocysteine Coupling.** Unique amino acids were single coupled using 0.25 M amino acid in DMF (2 equiv.), 0.5 M HATU (2 equiv.), oxyma (2 equiv.) and DIPEA (7 equiv.) at 23 °C for 16 h.

**Caping:** After double coupling, the resin was capped with $Ac_2O$ (5 equiv.) and DIPEA (10 equiv.) in DMF (20 mL) for 15 min. The resin was washed with DMF (3 x 20 mL).

**Deprotection:** 20% piperidine + 0.1 M HOBt in DMF (30 mL) at 23 ºC (2 x 15 min). The resin was washed with DMF (5 x 20 mL).

**Global Deprotection:** The dry resin in a 250 mL polypropylene vial was treated with TFA:$H_2O$:thioanisole:TIPSH (85:5:5:5, 15 mL) and mixed on an overhead stirrer. The mixture was stirred at 23 ºC for 3 h before filtering, and the resin was washed with an additional portion of TFA (3 x 4 mL). The pooled filtrate was cooled to 0°C and treated with $Et_2O$ to precipitate the peptide. The precipitate was centrifuged, and the resulting pellet was washed twice more with $Et_2O$ before it was dissolved in 40% MeCN and lyophilized.

**Crude UPLC-MS:** Measured on a HALO C18 column (3.0 x 30 mm; 2.7 µm; 90 Å; 40 °C; 1.5 mL/min) with a gradient of 5% MeCN/$H_2O$/0.1% TFA until 0.3 min, 5-95% MeCN/$H_2O$/0.1% TFA until 3.0 min, 95% MeCN/$H_2O$/0.1% TFA until 4.0 min.

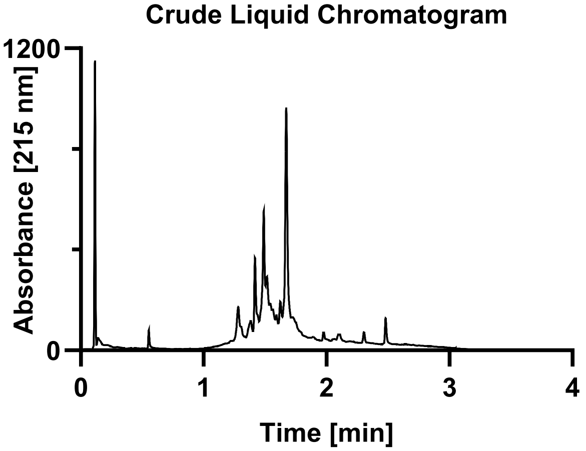

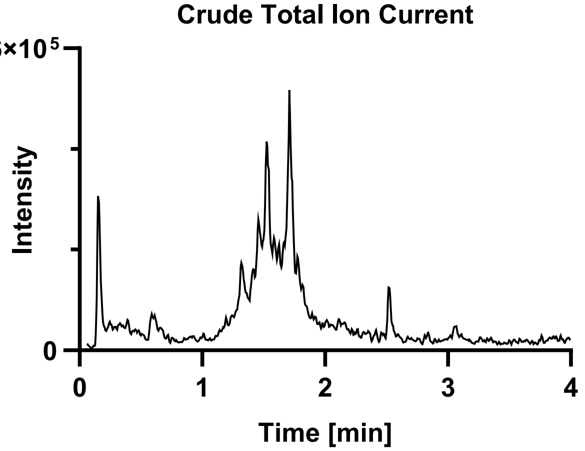

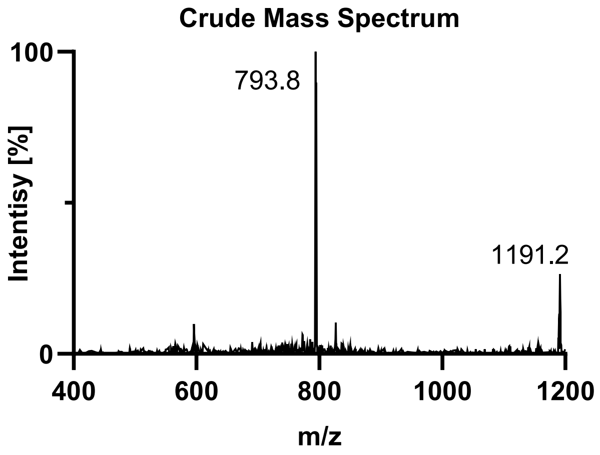

**Purification**: The material was purified over a semi-prep C4 column (10 x 250 mm, 10 µm, 300 Å, 5.0 mL/min) 18-45% MeCN/$H_2O$/0.1% TFA over 30 min. The fractions were lyophilized to afford **S3m** (52.5 mg, 5% yield).

**Pure UPLC-MS:** Measured on a HALO C18 column (3.0 x 30 mm; 2.7 µm; 90 Å; 40 °C; 1.5 mL/min) with a gradient of 5% MeCN/$H_2O$/0.1% TFA until 0.3 min, 5-95% MeCN/$H_2O$/0.1% TFA until 3.0 min, 95% MeCN/$H_2O$/0.1% TFA until 4.0 min.
**Chemical Formula:** $C_{93}H_{148}N_{25}O_{37}P_3Se$
**Molecular Weight:** 2380.237 g/mol
**LRMS for [M+2H+]:** $C_{93}H_{149}N_{25}O_{34}P_2Se^{2+}$ 1191.5, found 1191.3.
**LRMS for [M+3H+]:** $C_{93}H_{150}N_{25}O_{34}P_2Se^{3+}$ 794.6, found 794.5.

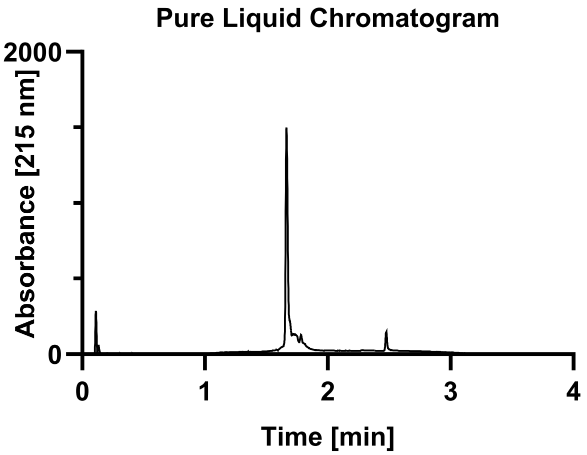

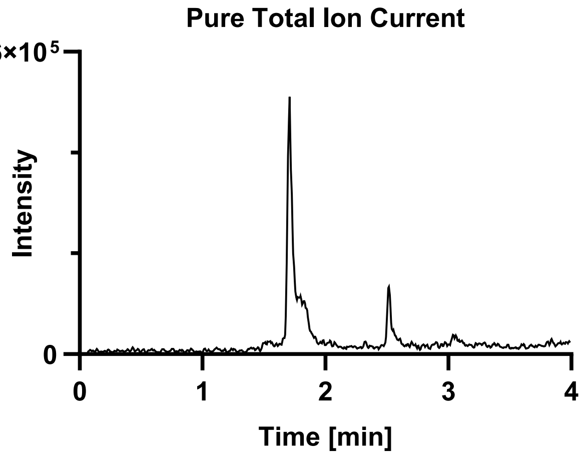

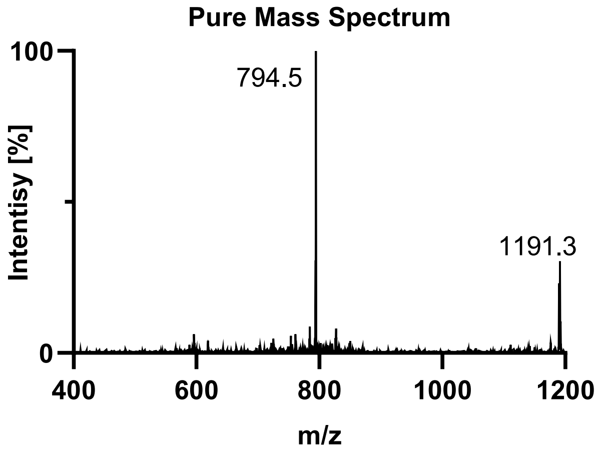

Tau(390-407) pS396 pT403 pS404 (**S3n**).

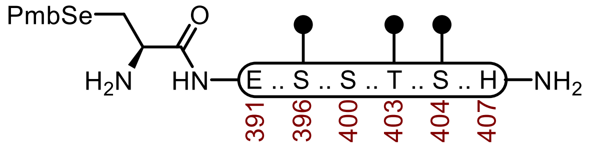

**S3n**

**Sequence:** H-C(SePmb)EIVYKS($OPO_3HBn$)PVVSGDT($OPO_3HBn$)S($OPO_3HBn$)PRH-$NH_2$

**Amino Acids:** Fmoc-L-SeC(Mob)-OH, Fmoc-L-Glu(O*t*Bu)-OH, Fmoc-Ile-OH, Fmoc-Val-OH, Fmoc-Tyr(*t*Bu)-OH, Fmoc-Lys(Boc)-OH, Fmoc-Ser(*t*Bu)-OH, Fmoc-Pro-OH, Fmoc-Val-OH, Fmoc-Ser($OPO_3HBn$)-OH, Fmoc-Gly-OH, Fmoc-Thr(*t*Bu)-OH, Fmoc-Arg(Pbf)-OH, Fmoc-His(Trt)-OH.

**Resin:** Fmoc-RinkAmide Protide resin from CEM (100-200 mesh, 0.6 mmol/g, 0.833 g, 0.50 mmol) was used.

**Coupling:** Amino acids were double coupled using a preactivated solution of Fmoc-AA-OH (0.25 M, 4 equiv.), HATU (0.5 M, 4 equiv.), and DIPEA (5 equiv.) at 23 °C for 1 h. The resin was washed with DMF (2 x 35 mL)

**Phosphoserine, phosphothreonine, and Selenocysteine Coupling.** Unique amino acids were single coupled using 0.25 M amino acid in DMF (2 equiv.), 0.5 M HATU (2 equiv.), oxyma (2 equiv.) and DIPEA (7 equiv.) at 23 °C for 16 h.

**Caping:** After double coupling, the resin was capped with $Ac_2O$ (5 equiv.) and DIPEA (10 equiv.) in DMF (20 mL) for 15 min. The resin was washed with DMF (3 x 20 mL).

**Deprotection:** 20% piperidine + 0.1 M HOBt in DMF (30 mL) at 23 ºC (2 x 15 min). The resin was washed with DMF (5 x 20 mL).

**Global Deprotection:** The dry resin in a 250 mL polypropylene vial was treated with TFA:$H_2O$:thioanisole:TIPSH (85:5:5:5, 15 mL) and mixed on an overhead stirrer. The mixture was stirred at 23 ºC for 3 h before filtering, and the resin was washed with an additional portion of TFA (3 x 4 mL). The pooled filtrate was cooled to 0°C and treated with $Et_2O$ to precipitate the peptide. The precipitate was centrifuged, and the resulting pellet was washed twice more with $Et_2O$ before it was dissolved in 40% MeCN and lyophilized.

**Crude UPLC-MS:** Measured on a HALO C18 column (3.0 x 30 mm; 2.7 µm; 90 Å; 40 °C; 1.5 mL/min) with a gradient of 5% MeCN/$H_2O$/0.1% TFA until 0.3 min, 5-95% MeCN/$H_2O$/0.1% TFA until 3.0 min, 95% MeCN/$H_2O$/0.1% TFA until 4.0 min.

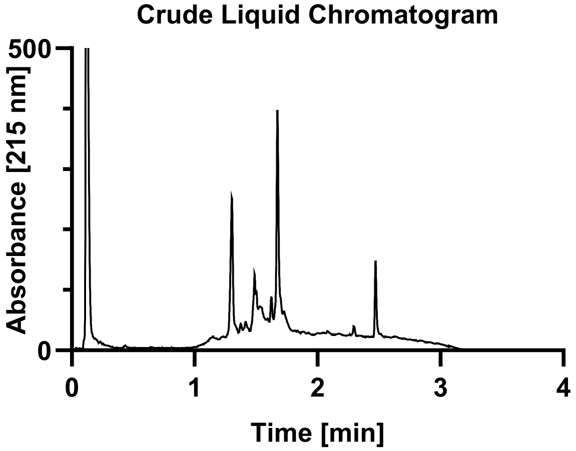

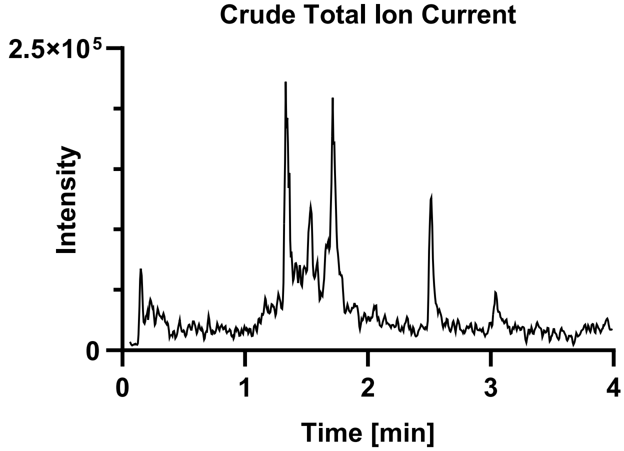

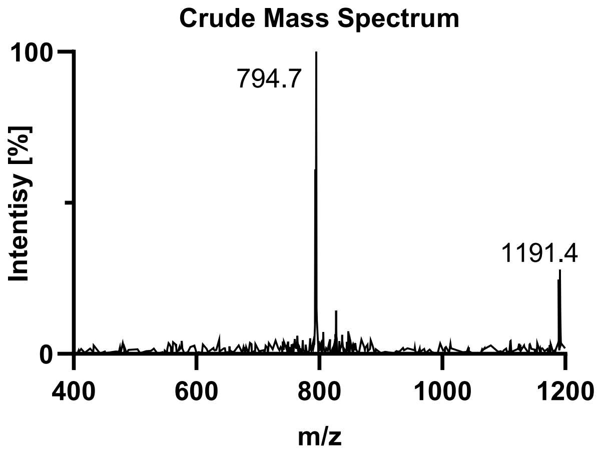

**Purification**: The material was purified over a semi-prep C4 column (10 x 250 mm, 10 µm, 300 Å, 5.0 mL/min) 18-45% MeCN/$H_2O$/0.1% TFA over 30 min. The fractions were lyophilized to afford **S3n** (61.2 mg, 5% yield).

**Pure UPLC-MS:** Measured on a HALO C18 column (3.0 x 30 mm; 2.7 µm; 90 Å; 40 °C; 1.5 mL/min) with a gradient of 5% MeCN/$H_2O$/0.1% TFA until 0.3 min, 5-95% MeCN/$H_2O$/0.1% TFA until 3.0 min, 95% MeCN/$H_2O$/0.1% TFA until 4.0 min.
**Chemical Formula**: $C_{93}H_{148}N_{25}O_{37}P_3Se$
**Molecular Weight:** 2380.237 g/mol
**LRMS for [M+2H+]:** $C_{93}H_{149}N_{25}O_{34}P_2Se^{2+}$ 1191.5, found 1191.3.
**LRMS for [M+3H+]:** $C_{93}H_{150}N_{25}O_{34}P_2Se^{3+}$ 794.6, found 794.3.

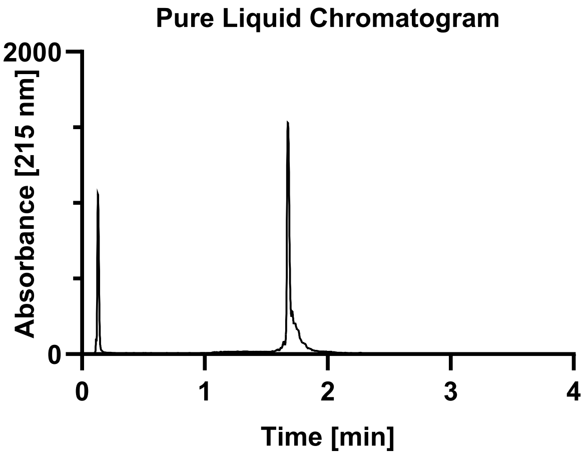

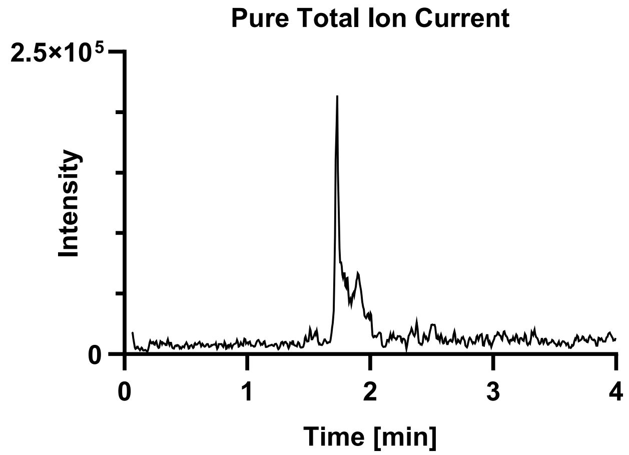

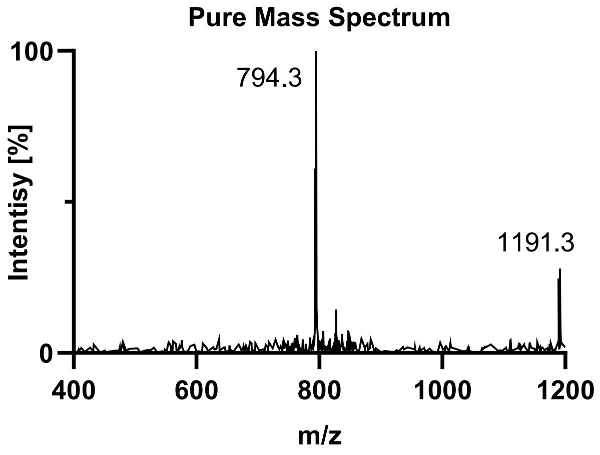

Tau(390-407) pS400 pT403 pS404 (**S3o**).

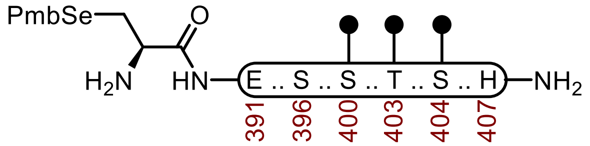

**S3o**

**Sequence:** H-C(SePmb)EIVYKSPVVS($OPO_3HBn$)GDT($OPO_3HBn$)S($OPO_3HBn$)PRH-$NH_2$

**Amino Acids:** Fmoc-L-SeC(Mob)-OH, Fmoc-L-Glu(O*t*Bu)-OH, Fmoc-Ile-OH, Fmoc-Val-OH, Fmoc-Tyr(*t*Bu)-OH, Fmoc-Lys(Boc)-OH, Fmoc-Ser(*t*Bu)-OH, Fmoc-Pro-OH, Fmoc-Val-OH, Fmoc-Ser($OPO_3HBn$)-OH, Fmoc-Gly-OH, Fmoc-Thr(*t*Bu)-OH, Fmoc-Arg(Pbf)-OH, Fmoc-His(Trt)-OH.

**Resin:** Fmoc-RinkAmide Protide resin from CEM (100-200 mesh, 0.6 mmol/g, 0.833 g, 0.50 mmol) was used.

**Coupling:** Amino acids were double coupled using a preactivated solution of Fmoc-AA-OH (0.25 M, 4 equiv.), HATU (0.5 M, 4 equiv.), and DIPEA (5 equiv.) at 23 °C for 1 h. The resin was washed with DMF (2 x 35 mL)

**Phosphoserine, phosphothreonine, and Selenocysteine Coupling.** Unique amino acids were single coupled using 0.25 M amino acid in DMF (2 equiv.), 0.5 M HATU (2 equiv.), oxyma (2 equiv.) and DIPEA (7 equiv.) at 23 °C for 16 h.

**Caping:** After double coupling, the resin was capped with $Ac_2O$ (5 equiv.) and DIPEA (10 equiv.) in DMF (20 mL) for 15 min. The resin was washed with DMF (3 x 20 mL).

**Deprotection:** 20% piperidine + 0.1 M HOBt in DMF (30 mL) at 23 ºC (2 x 15 min). The resin was washed with DMF (5 x 20 mL).

**Global Deprotection:** The dry resin in a 250 mL polypropylene vial was treated with TFA:$H_2O$:thioanisole:TIPSH (85:5:5:5, 15 mL) and mixed on an overhead stirrer. The mixture was stirred at 23 ºC for 3 h before filtering, and the resin was washed with an additional portion of TFA (3 x 4 mL). The pooled filtrate was cooled to 0°C and treated with $Et_2O$ to precipitate the peptide. The precipitate was centrifuged, and the resulting pellet was washed twice more with $Et_2O$ before it was dissolved in 40% MeCN and lyophilized.

**Crude UPLC-MS:** Measured on a HALO C18 column (3.0 x 30 mm; 2.7 µm; 90 Å; 40 °C; 1.5 mL/min) with a gradient of 5% MeCN/$H_2O$/0.1% TFA until 0.3 min, 5-95% MeCN/$H_2O$/0.1% TFA until 3.0 min, 95% MeCN/$H_2O$/0.1% TFA until 4.0 min.

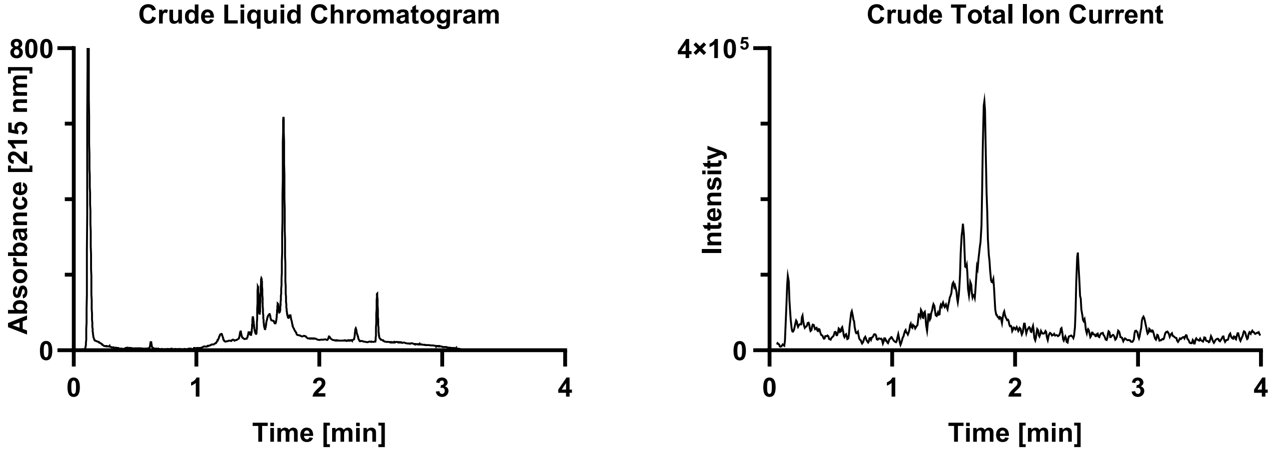

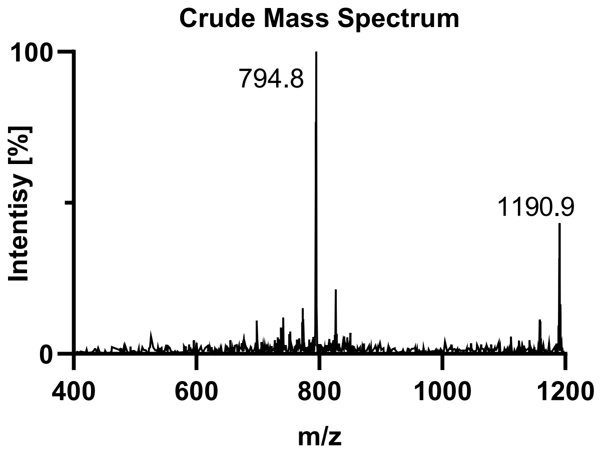

**Purification**: The material was purified over a semi-prep C4 column (10 x 250 mm, 10 µm, 300 Å, 5.0 mL/min) 18-45% MeCN/$H_2O$/0.1% TFA over 30 min. The fractions were lyophilized to afford **S3o** (62.1 mg, 5% yield).

**Pure UPLC-MS:** Measured on a HALO C18 column (3.0 x 30 mm; 2.7 µm; 90 Å; 40 °C; 1.5 mL/min) with a gradient of 5% MeCN/$H_2O$/0.1% TFA until 0.3 min, 5-95% MeCN/$H_2O$/0.1% TFA until 3.0 min, 95% MeCN/$H_2O$/0.1% TFA until 4.0 min.
**Chemical Formula:** $C_{93}H_{148}N_{25}O_{37}P_3Se$
**Molecular Weight:** 2380.237 g/mol
**LRMS for [M+2H+]:** $C_{93}H_{149}N_{25}O_{34}P_2Se^{2+}$ 1191.5, found 1191.8.
**LRMS for [M+3H+]:** $C_{93}H_{150}N_{25}O_{34}P_2Se^{3+}$ 794.6, found 794.3.

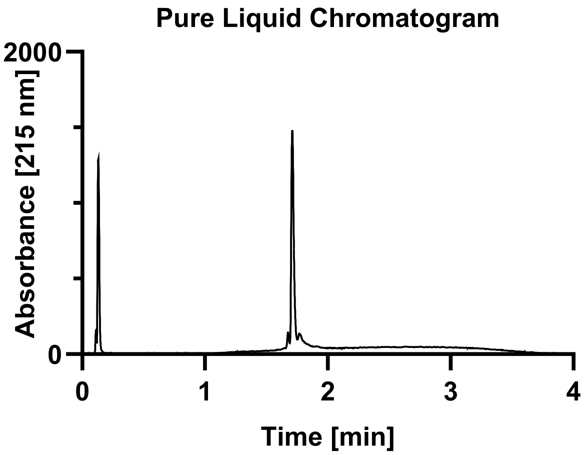

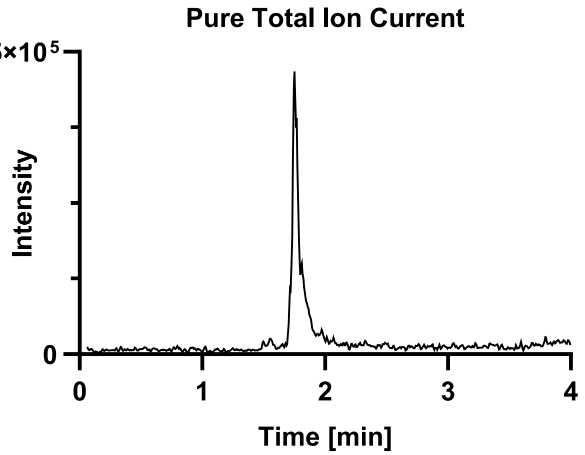

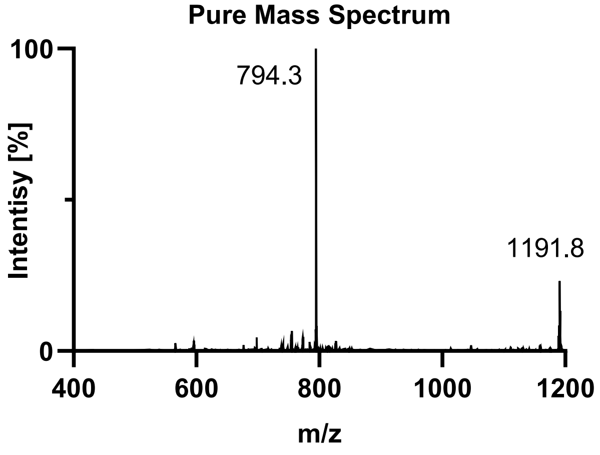

### DISELENIDE SYNTHESIS.

Tau(390-407) WT diselenide (**2a**).

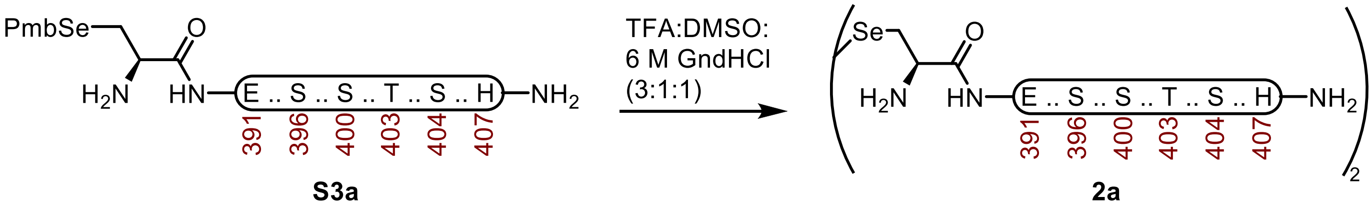

**Tau(390-407) diselenide (2a).** A 2-dram vial containing **S3a** (93.0 mg, 43.5 µmol) was dissolved in buffer (300 µL, 6 M Gnd·HCl, 0.1 M $Na_2HPO_4$, pH 7.2) and DMSO (300 µL). The mixture was treated with TFA (900 µL), wrapped with Al-Foil, and incubated at 23 ºC. After 45 min it was purified over a C4 column (10 x 250 mm, 10 µm, 100 Å, 4.7 mL/min) 20%-45% $MeCN/H_2O$/0.1% TFA over 30 min. The fractions were lyophilized to afford **2a** (78.7 mg, 90% yield).

**UPLC-MS:** Measured on an ACQUITY UPLC Protein BEH C4 column (2.1 x 50 mm; 1.7 µm; 300 Å; 50 °C; 0.6 mL/min) with a gradient of 5% $MeCN/H_2O$/0.1% FA until 0.1 min, 5-95% $MeCN/H_2O$/0.1% FA until 1.0 min, 95% $MeCN/H_2O$/0.1% FA until 1.1 min.

**Chemical Formula:** $C_{170}H_{272}N_{50}O_{54}Se_2$
**Molecular Weight:** 4038.2840 g/mol
**HRMS for [M+4H+]:** $C_{170}H_{276}N_{50}O_{54}Se_2^{4+}$ 1010.7, found 1011.1.
**HRMS for [M+5H+]:** $C_{170}H_{277}N_{50}O_{54}Se_2^{5+}$ 808.8, found 808.9.
**HRMS for [M+6H+]:** $C_{170}H_{278}N_{50}O_{54}Se_2^{6+}$ 674.1, found 674.0.

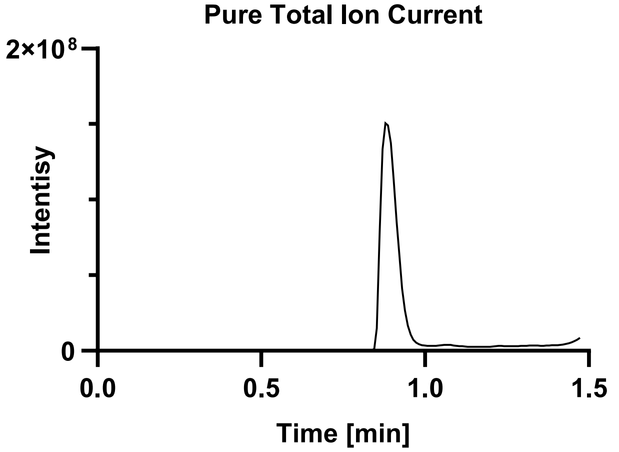

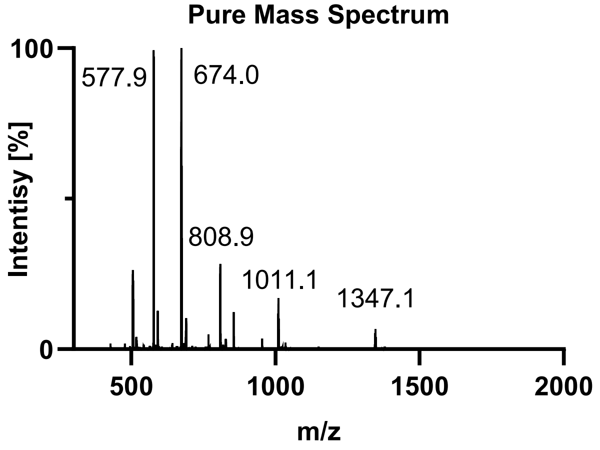

Tau(390-407) pS396 diselenide (**2b**).

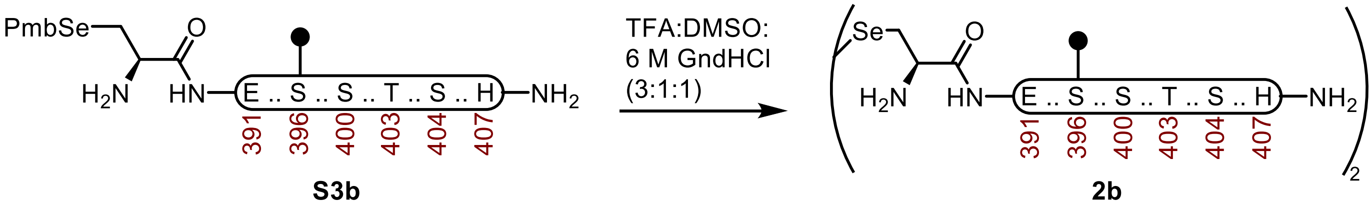

**Tau(390-407) pS396 diselenide (2b).** A 2-dram vial containing **S3b** (80.8 mg, 36.4 µmol) was dissolved in buffer (250 µL, 6 M Gnd·HCl, 0.1 M $Na_2HPO_4$, pH 7.2) and DMSO (250 µL). The mixture was treated with TFA (750 µL), wrapped with Al-Foil, and incubated at 23 ºC. After 40 min it was purified over a C4 column (10 x 250 mm, 10 µm, 100 Å, 4.7 mL/min) 7%-45% $MeCN/H_2O$/0.1% TFA over 30 min. The fractions were lyophilized to afford **2b** (58.0 mg, 76% yield).

**UPLC-MS:** Measured on an ACQUITY UPLC Protein BEH C4 column (2.1 x 50 mm; 1.7 µm; 300 Å; 50 °C; 0.6 mL/min) with a gradient of 5% $MeCN/H_2O$/0.1% FA until 0.1 min, 5-95% $MeCN/H_2O$/0.1% FA until 1.0 min, 95% $MeCN/H_2O$/0.1% FA until 1.1 min.

**Chemical Formula:** $C_{170}H_{274}N_{50}O_{60}P_2Se_2$
**Molecular Weight:** 4198.242 g/mol
**HRMS for [M+4H+]:** $C_{170}H_{278}N_{50}O_{60}P_2Se_2^{4+}$ 1050.7, found 1050.4.
**HRMS for [M+5H+]:** $C_{170}H_{279}N_{50}O_{60}P_2Se_2^{5+}$ 840.4, found 840.7.
**HRMS for [M+6H+]:** $C_{170}H_{280}N_{50}O_{60}P_2Se_2^{6+}$ 700.8, found 700.6.

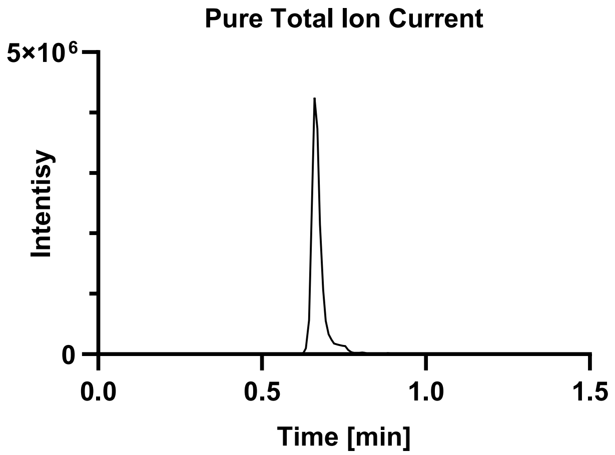

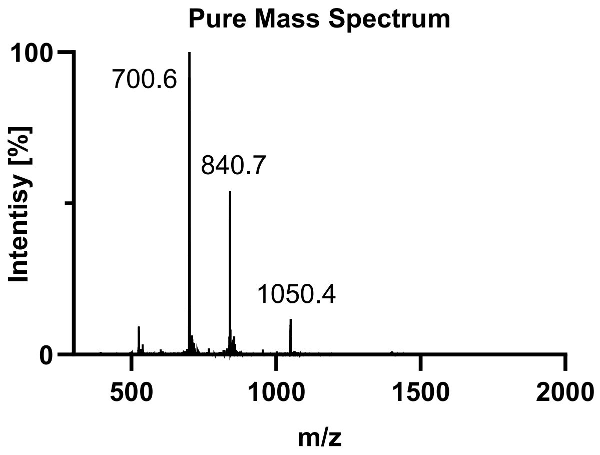

Tau(390-407) pS400 diselenide (**2c**).

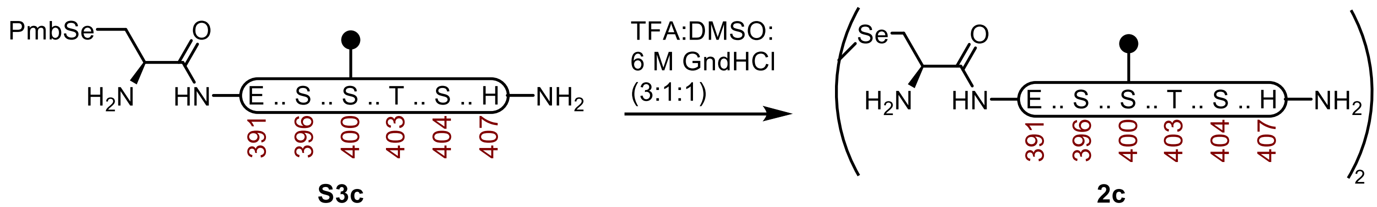

**S3c** **2c**

**Tau(390-407) pS400 diselenide (2c).** A 2-dram vial containing **S3c** (86.1 mg, 38.8 μmol) was dissolved in buffer (250 μL, 6 M Gnd·HCl, 0.1 M $Na_2HPO_4$, pH 7.2) and DMSO (250 μL). The mixture was treated with TFA (750 μL), wrapped with Al-Foil, and incubated at 23 ºC. After 60 min it was purified over a C4 column (10 x 250 mm, 10 μm, 100 Å, 4.7 mL/min) 7%-45% MeCN/$H_2O$/0.1% TFA over 30 min. The fractions were lyophilized to afford **2c** (68.7 mg, 84% yield).

**UPLC-MS:** Measured on an ACQUITY UPLC Protein BEH C4 column (2.1 x 50 mm; 1.7 μm; 300 Å; 50 °C; 0.6 mL/min) with a gradient of 5% MeCN/$H_2O$/0.1% FA until 0.1 min, 5-95% MeCN/$H_2O$/0.1% FA until 1.0 min, 95% MeCN/$H_2O$/0.1% FA until 1.1 min.

**Chemical Formula:** $C_{170}H_{274}N_{50}O_{60}P_2Se_2$
**Molecular Weight:** 4198.242 g/mol
**HRMS for [M+4H+]:** $C_{170}H_{278}N_{50}O_{60}P_2Se_2^{4+}$ 1050.7, found 1050.4.
**HRMS for [M+5H+]:** $C_{170}H_{279}N_{50}O_{60}P_2Se_2^{5+}$ 840.4, found 840.7.
**HRMS for [M+6H+]:** $C_{170}H_{280}N_{50}O_{60}P_2Se_2^{6+}$ 700.8, found 700.6.

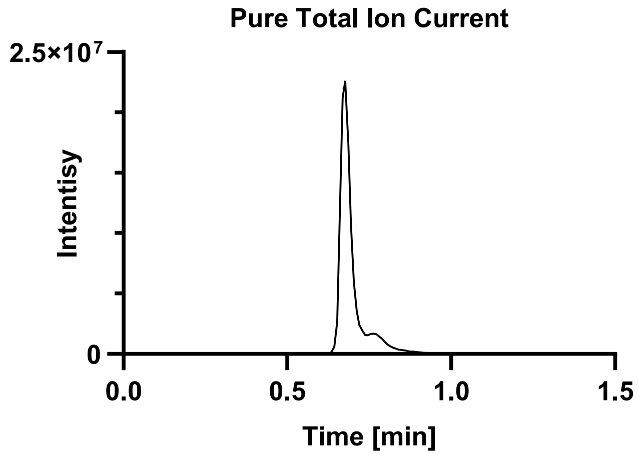

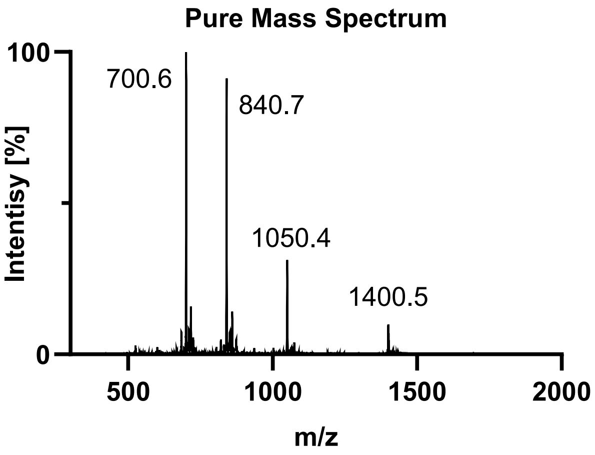

Tau(390-407) pT403 diselenide (**2d**).

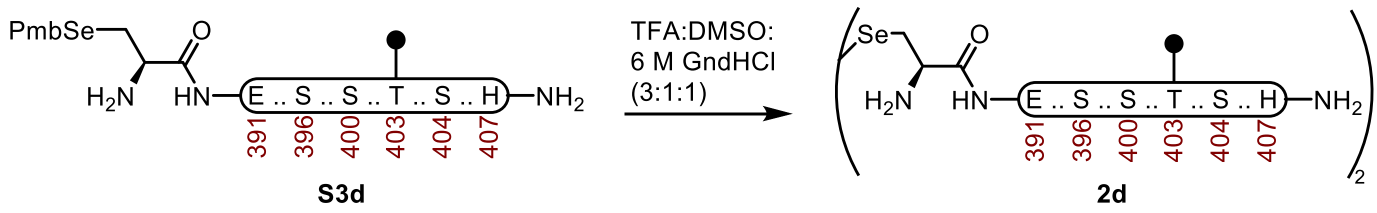

**Tau(390-407) pT403 diselenide (2d).** A 2-dram vial containing **S3d** (93.6 mg, 42.2 μmol) was dissolved in buffer (250 μL, 6 M Gnd·HCl, 0.1 M $Na_2HPO_4$, pH 7.2) and DMSO (250 μL). The mixture was treated with TFA (750 μL), wrapped with Al-Foil, and incubated at 23 ºC. After 30 min it was purified over a C4 column (10 x 250 mm, 10 μm, 100 Å, 4.7 mL/min) 7%-45% MeCN/$H_2O$/0.1% TFA over 30 min. The fractions were lyophilized to afford **2d** (81.2 mg, 92% yield).

**UPLC-MS:** Measured on an ACQUITY UPLC Protein BEH C4 column (2.1 x 50 mm; 1.7 μm; 300 Å; 50 °C; 0.6 mL/min) with a gradient of 5% MeCN/$H_2O$/0.1% FA until 0.1 min, 5-95% MeCN/$H_2O$/0.1% FA until 1.0 min, 95% MeCN/$H_2O$/0.1% FA until 1.1 min.

**Chemical Formula:** $C_{170}H_{274}N_{50}O_{60}P_2Se_2$
**Molecular Weight:** 4198.242 g/mol
**HRMS for [M+4H+]:** $C_{170}H_{278}N_{50}O_{60}P_2Se_2^{4+}$ 1050.7, found 1050.6.
**HRMS for [M+5H+]:** $C_{170}H_{279}N_{50}O_{60}P_2Se_2^{5+}$ 840.4, found 840.7.
**HRMS for [M+6H+]:** $C_{170}H_{280}N_{50}O_{60}P_2Se_2^{6+}$ 700.8, found 700.8.

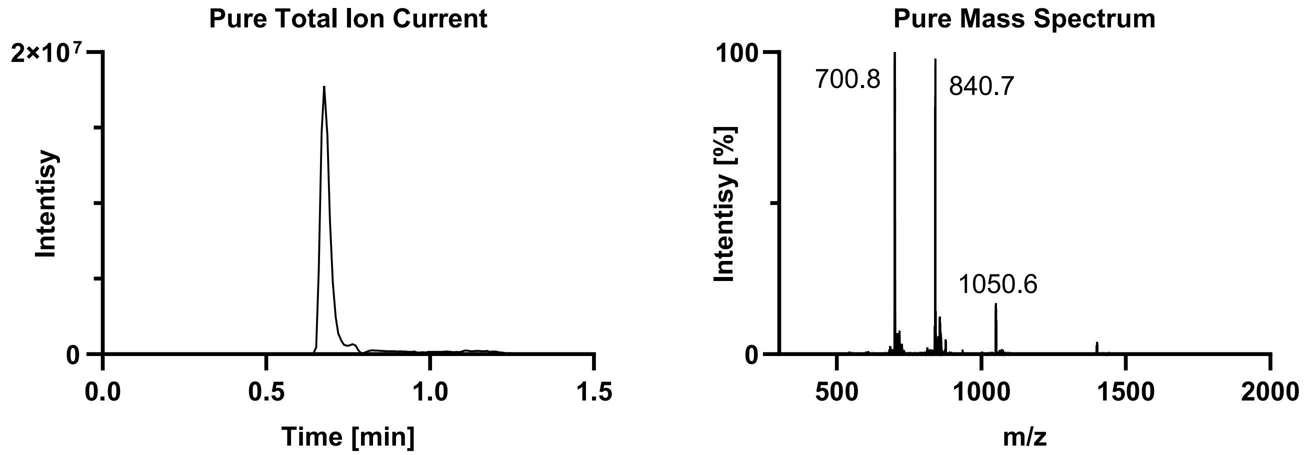

Tau(390-407) pS404 diselenide (**2e**).

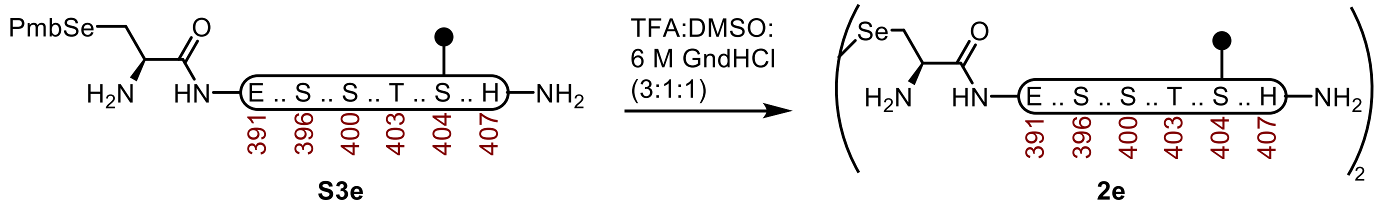

**Tau(390-407) pS404 diselenide (2e).** A 2-dram vial containing **S3e** (74.8 mg, 33.7 μmol) was dissolved in buffer (250 μL, 6 M Gnd·HCl, 0.1 M $Na_2HPO_4$, pH 7.2) and DMSO (250 μL). The mixture was treated with TFA (750 μL), wrapped with Al-Foil, and incubated at 23 ºC. After 30 min it was purified over a C4 column (10 x 250 mm, 10 μm, 100 Å, 4.7 mL/min) 7%-45% $MeCN/H_2O$/0.1% TFA over 30 min. The fractions were lyophilized to afford **2e** (61.5 mg, 87% yield).

**UPLC-MS:** Measured on an ACQUITY UPLC Protein BEH C4 column (2.1 x 50 mm; 1.7 μm; 300 Å; 50 °C; 0.6 mL/min) with a gradient of 5% $MeCN/H_2O$/0.1% FA until 0.1 min, 5-95% $MeCN/H_2O$/0.1% FA until 1.0 min, 95% $MeCN/H_2O$/0.1% FA until 1.1 min.

**Chemical Formula:** $C_{170}H_{274}N_{50}O_{60}P_2Se_2$
**Molecular Weight:** 4198.242 g/mol
**HRMS for [M+4H+]:** $C_{170}H_{278}N_{50}O_{60}P_2Se_2^{4+}$ 1050.7, found 1050.4.
**HRMS for [M+5H+]:** $C_{170}H_{279}N_{50}O_{60}P_2Se_2^{5+}$ 840.4, found 840.7.
**HRMS for [M+6H+]:** $C_{170}H_{280}N_{50}O_{60}P_2Se_2^{6+}$ 700.8, found 700.6.

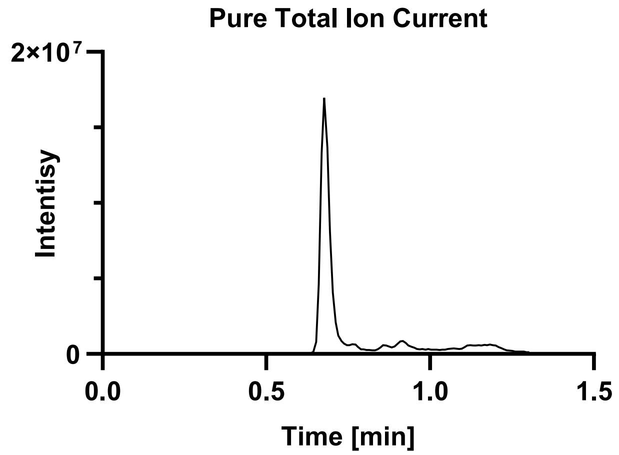

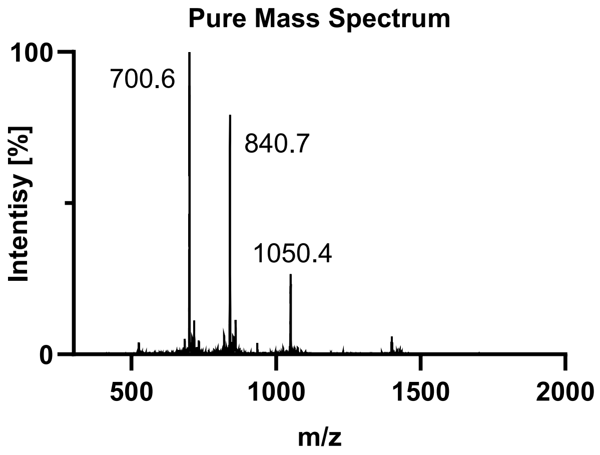

Tau(390-407) pS396 pS400 diselenide (**2f**).

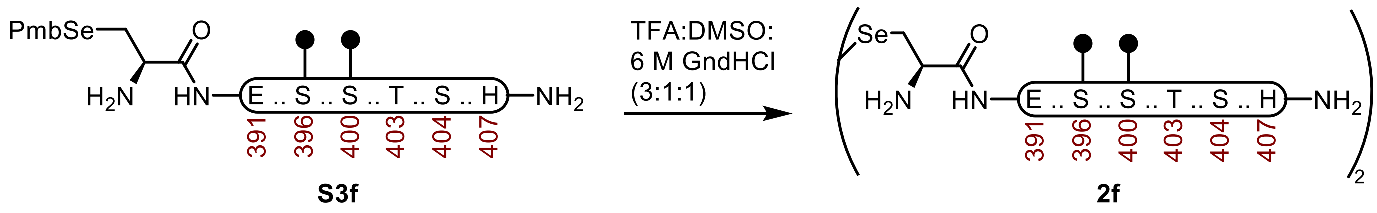

**Tau(390-407) pS396 pS400 diselenide (2f).** A 2-dram vial containing **S3f** (76.9 mg, 33.4 µmol) was dissolved in buffer (250 µL, 6 M Gnd·HCl, 0.1 M $Na_2HPO_4$, pH 7.2) and DMSO (250 µL). The mixture was treated with TFA (750 µL), wrapped with Al-Foil, and incubated at 23 ºC. After 30 min it was purified over a C4 column (10 x 250 mm, 10 µm, 100 Å, 4.7 mL/min) 7%-45% $MeCN/H_2O$/0.1% TFA over 30 min. The fractions were lyophilized to afford **2f** (56.6 mg, 78% yield).

**UPLC-MS:** Measured on an ACQUITY UPLC Protein BEH C4 column (2.1 x 50 mm; 1.7 µm; 300 Å; 50 °C; 0.6 mL/min) with a gradient of 5% $MeCN/H_2O$/0.1% FA until 0.1 min, 5-95% $MeCN/H_2O$/0.1% FA until 1.0 min, 95% $MeCN/H_2O$/0.1% FA until 1.1 min.

**Chemical Formula:** $C_{170}H_{276}N_{50}O_{66}P_4Se_2$
**Molecular Weight:** 4358.199 g/mol
**HRMS for [M+4H+]:** $C_{170}H_{280}N_{50}O_{66}P_4Se_2^{4+}$ 1090.7, found 1090.6.
**HRMS for [M+5H+]:** $C_{170}H_{281}N_{50}O_{66}P_4Se_2^{5+}$ 872.4, found 872.7.

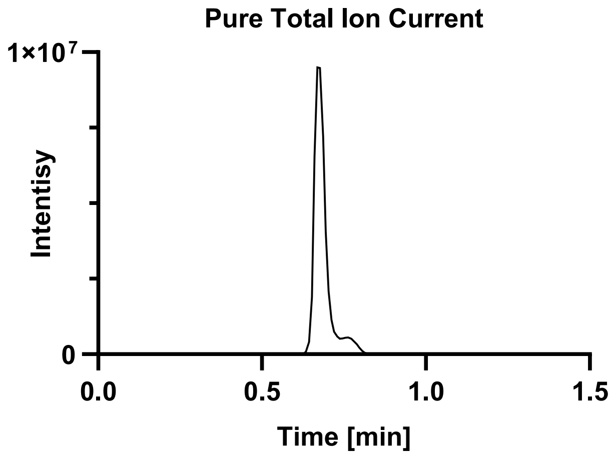

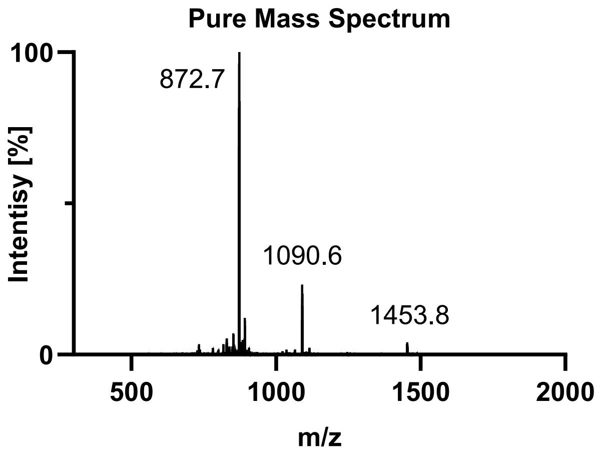

Tau(390-407) pS396 pT403 diselenide (**2g**).

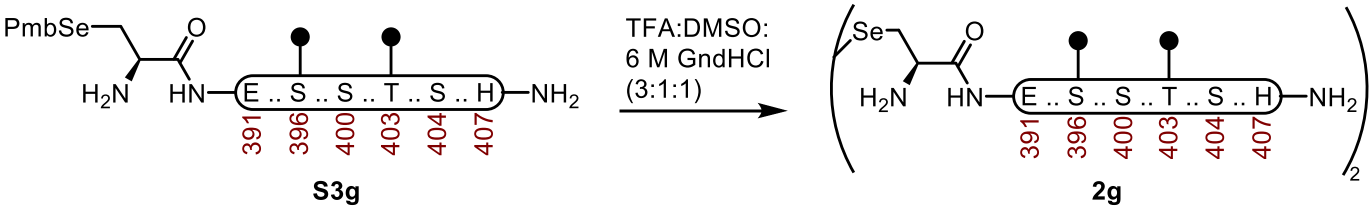

**Tau(390-407) pS396 pT403 diselenide (2g).** A 2-dram vial containing **S3g** (45.7 mg, 19.9 µmol) was dissolved in buffer (250 µL, 6 M Gnd·HCl, 0.1 M $Na_2HPO_4$, pH 7.2) and DMSO (250 µL). The mixture was treated with TFA (750 µL), wrapped with Al-Foil, and incubated at 23 ºC. After 30 min it was purified over a C4 column (10 x 250 mm, 10 µm, 100 Å, 4.7 mL/min) 7%-45% MeCN/$H_2O$/0.1% TFA over 30 min. The fractions were lyophilized to afford **2g** (38.5 mg, 89% yield).

**UPLC-MS:** Measured on an ACQUITY UPLC Protein BEH C4 column (2.1 x 50 mm; 1.7 µm; 300 Å; 50 °C; 0.6 mL/min) with a gradient of 5% MeCN/$H_2O$/0.1% FA until 0.1 min, 5-95% MeCN/$H_2O$/0.1% FA until 1.0 min, 95% MeCN/$H_2O$/0.1% FA until 1.1 min.

**Chemical Formula:** $C_{170}H_{276}N_{50}O_{66}P_4Se_2$
**Molecular Weight:** 4358.199 g/mol
**HRMS for [M+4H+]:** $C_{170}H_{280}N_{50}O_{66}P_4Se_2^{4+}$ 1090.7, found 1090.6.
**HRMS for [M+5H+]:** $C_{170}H_{281}N_{50}O_{66}P_4Se_2^{5+}$ 872.4, found 872.3.

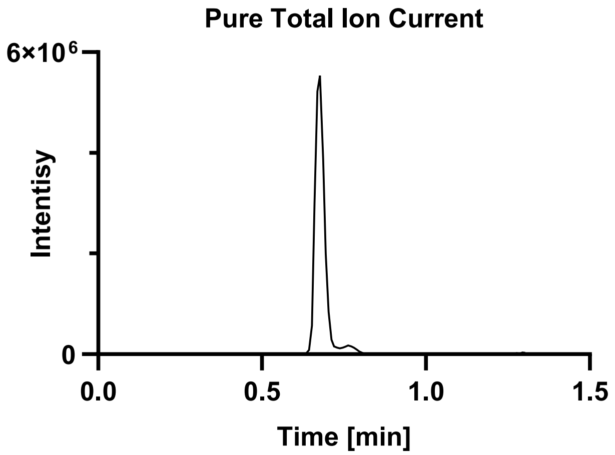

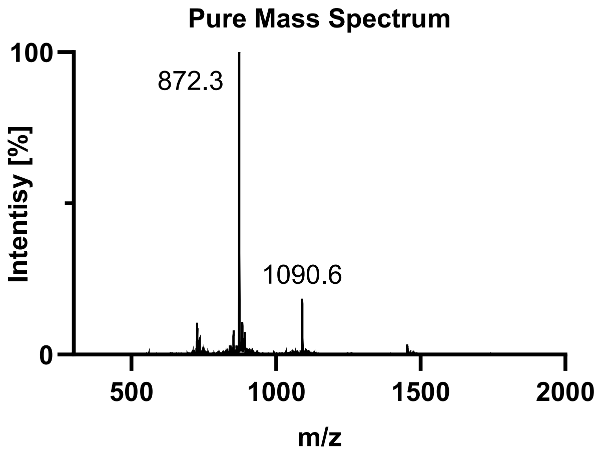

Tau(390-407) pS396 pS404 diselenide (**2h**).

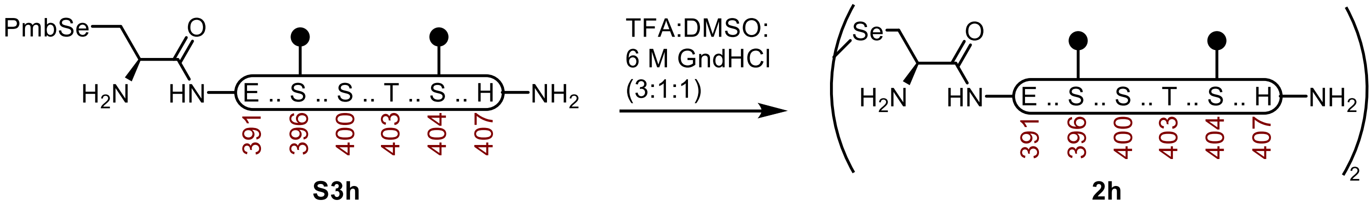

**Tau(390-407) pS396 pS404 diselenide (2h).** A 2-dram vial containing **S3h** (51.9 mg, 22.6 µmol) was dissolved in buffer (250 µL, 6 M Gnd·HCl, 0.1 M $Na_2HPO_4$, pH 7.2) and DMSO (250 µL). The mixture was treated with TFA (750 µL), wrapped with Al-Foil, and incubated at 23 ºC. After 40 min it was purified over a C4 column (10 x 250 mm, 10 µm, 100 Å, 4.7 mL/min) 7%-45% $MeCN/H_2O$/0.1% TFA over 30 min. The fractions were lyophilized to afford **2h** (45.1 mg, 92% yield).

**UPLC-MS:** Measured on an ACQUITY UPLC Protein BEH C4 column (2.1 x 50 mm; 1.7 µm; 300 Å; 50 °C; 0.6 mL/min) with a gradient of 5% $MeCN/H_2O$/0.1% FA until 0.1 min, 5-95% $MeCN/H_2O$/0.1% FA until 1.0 min, 95% $MeCN/H_2O$/0.1% FA until 1.1 min.

**Chemical Formula:** $C_{170}H_{276}N_{50}O_{66}P_4Se_2$
**Molecular Weight:** 4358.199 g/mol
**HRMS for [M+4H+]:** $C_{170}H_{280}N_{50}O_{66}P_4Se_2^{4+}$ 1090.7, found 1090.6.
**HRMS for [M+5H+]:** $C_{170}H_{281}N_{50}O_{66}P_4Se_2^{5+}$ 872.4, found 872.7.

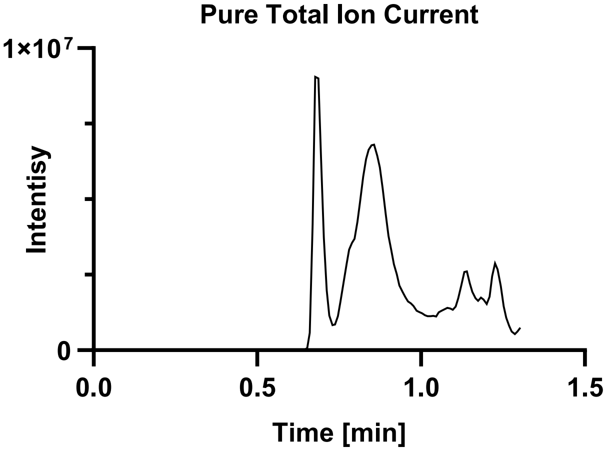

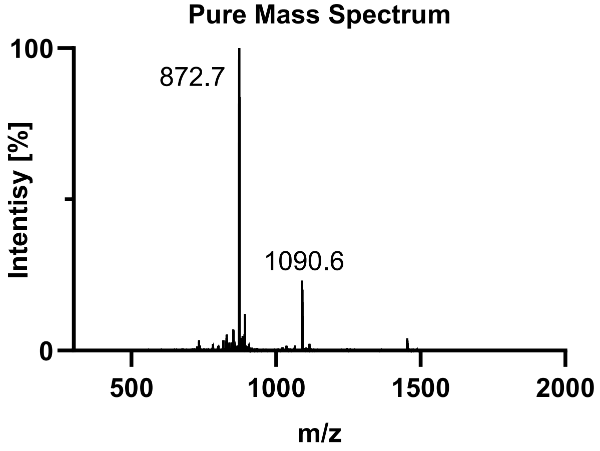

Tau(390-407) pS400 pT403 diselenide (**2i**).

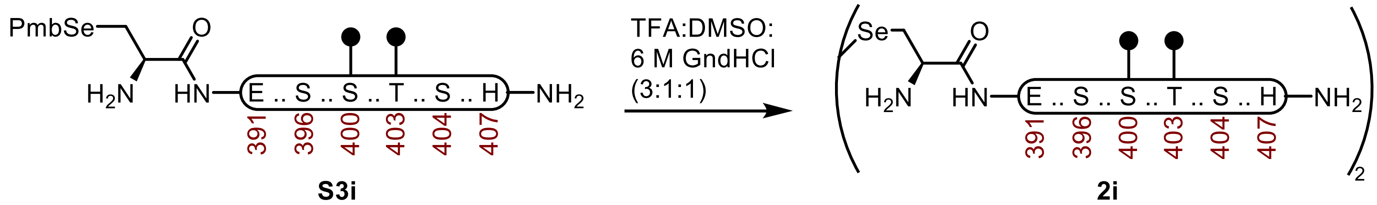

**Tau(390-407) pS400 pT403 diselenide (2i).** A 2-dram vial containing **S3i** (52.5 mg, 22.8 µmol) was dissolved in buffer (250 µL, 6 M Gnd·HCl, 0.1 M $Na_2HPO_4$, pH 7.2) and DMSO (250 µL). The mixture was treated with TFA (750 µL), wrapped with Al-Foil, and incubated at 23 ºC. After 30 min it was purified over a C4 column (10 x 250 mm, 10 µm, 100 Å, 4.7 mL/min) 7%-45% MeCN/$H_2O$/0.1% TFA over 30 min. The fractions were lyophilized to afford **2i** (44.1 mg, 89% yield).

**UPLC-MS:** Measured on an ACQUITY UPLC Protein BEH C4 column (2.1 x 50 mm; 1.7 µm; 300 Å; 50 °C; 0.6 mL/min) with a gradient of 5% MeCN/$H_2O$/0.1% FA until 0.1 min, 5-95% MeCN/$H_2O$/0.1% FA until 1.0 min, 95% MeCN/$H_2O$/0.1% FA until 1.1 min.

**Chemical Formula:** $C_{170}H_{276}N_{50}O_{66}P_4Se_2$
**Molecular Weight:** 4358.199 g/mol
**HRMS for [M+4H+]:** $C_{170}H_{280}N_{50}O_{66}P_4Se_2^{4+}$ 1090.7, found 1090.4.
**HRMS for [M+5H+]:** $C_{170}H_{281}N_{50}O_{66}P_4Se_2^{5+}$ 872.4, found 872.3.

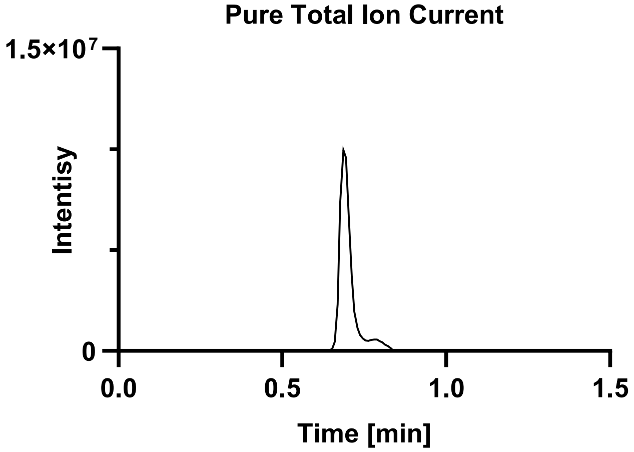

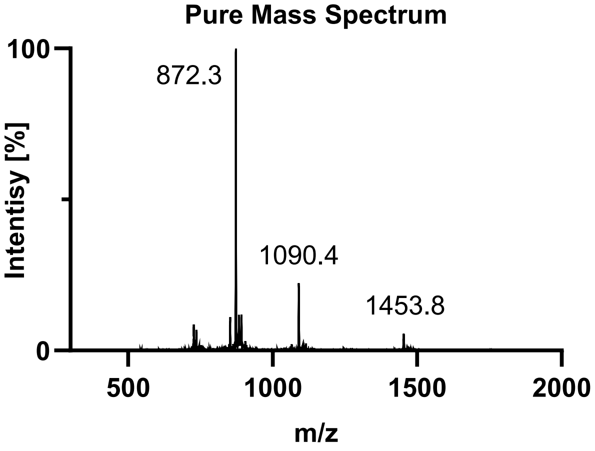

Tau(390-407) pS400 pS404 diselenide (**2j**).

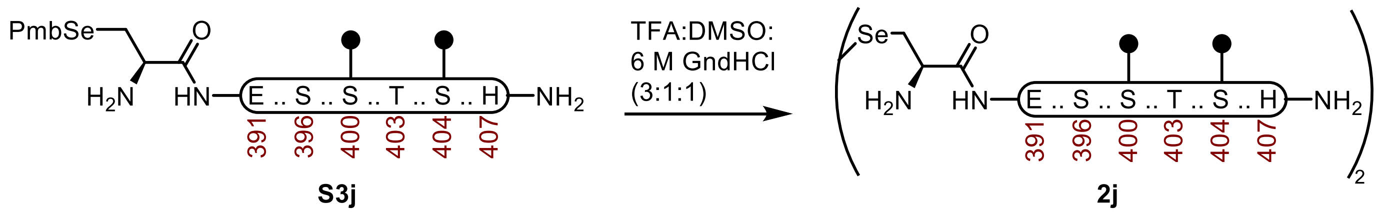

**Tau(390-407) pS400 pS404 diselenide (2j).** A 2-dram vial containing **S3j** (54.2 mg, 23.6 µmol) was dissolved in buffer (250 µL, 6 M Gnd·HCl, 0.1 M $Na_2HPO_4$, pH 7.2) and DMSO (250 µL). The mixture was treated with TFA (750 µL), wrapped with Al-Foil, and incubated at 23 ºC. After 30 min it was purified over a C4 column (10 x 250 mm, 10 µm, 100 Å, 4.7 mL/min) 7%-45% MeCN/$H_2O$/0.1% TFA over 30 min. The fractions were lyophilized to afford **2j** (15 mg, 15% yield).

**UPLC-MS:** Measured on an ACQUITY UPLC Protein BEH C4 column (2.1 x 50 mm; 1.7 µm; 300 Å; 50 °C; 0.6 mL/min) with a gradient of 5% MeCN/$H_2O$/0.1% FA until 0.1 min, 5-95% MeCN/$H_2O$/0.1% FA until 1.0 min, 95% MeCN/$H_2O$/0.1% FA until 1.1 min.

**Chemical Formula:** $C_{170}H_{276}N_{50}O_{66}P_4Se_2$
**Molecular Weight:** 4358.199 g/mol
**HRMS for [M+4H+]:** $C_{170}H_{280}N_{50}O_{66}P_4Se_2^{4+}$ 1090.7, found 1090.9.
**HRMS for [M+5H+]:** $C_{170}H_{281}N_{50}O_{66}P_4Se_2^{5+}$ 872.4, found 872.6.

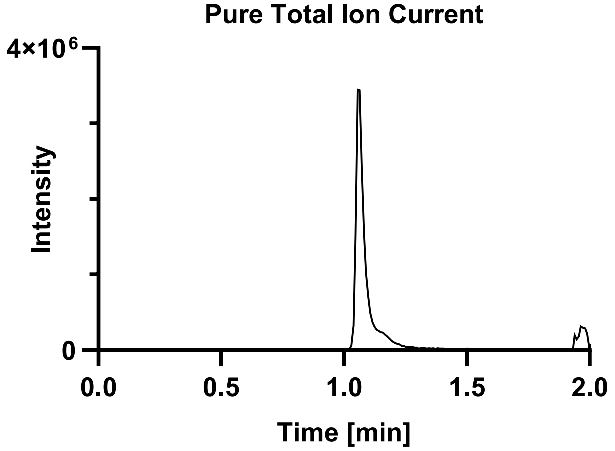

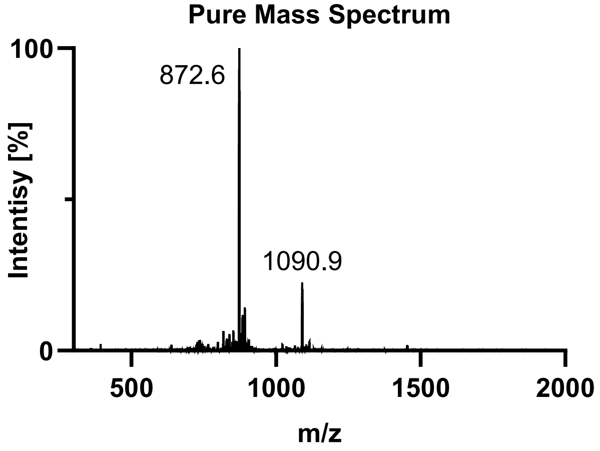

Tau(390-407) pT403 pS404 diselenide (**2k**).

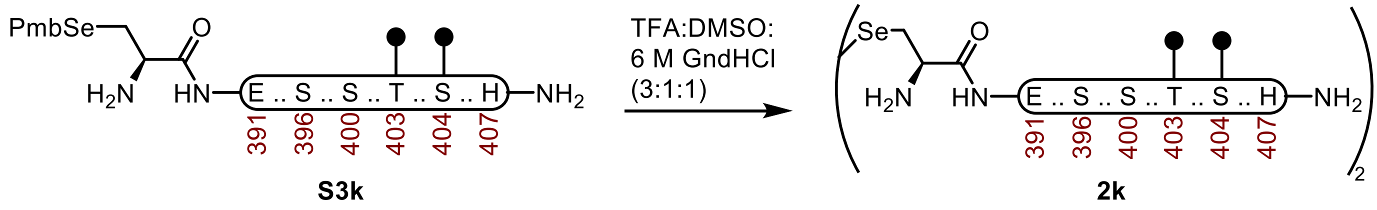

**Tau(390-407) pT403 pS404 diselenide (2k).** A 2-dram vial containing **S3k** (110 mg, 47.8 µmol) was dissolved in buffer (250 µL, 6 M Gnd·HCl, 0.1 M $Na_2HPO_4$, pH 7.2) and DMSO (250 µL). The mixture was treated with TFA (750 µL), wrapped with Al-Foil, and incubated at 23 ºC. After 30 min it was purified over a C4 column (10 x 250 mm, 10 µm, 100 Å, 4.7 mL/min) 7%-45% MeCN/$H_2O$/0.1% TFA over 30 min. The fractions were lyophilized to afford **2k** (82.3 mg, 79% yield).

**UPLC-MS:** Measured on an ACQUITY UPLC Protein BEH C4 column (2.1 x 50 mm; 1.7 µm; 300 Å; 50 °C; 0.6 mL/min) with a gradient of 5% MeCN/$H_2O$/0.1% FA until 0.1 min, 5-95% MeCN/$H_2O$/0.1% FA until 1.0 min, 95% MeCN/$H_2O$/0.1% FA until 1.1 min.

**Chemical Formula:** $C_{170}H_{276}N_{50}O_{66}P_4Se_2$
**Molecular Weight:** 4358.199 g/mol
**HRMS for [M+4H+]:** $C_{170}H_{280}N_{50}O_{66}P_4Se_2^{4+}$ 1090.7, found 1090.4.
**HRMS for [M+5H+]:** $C_{170}H_{281}N_{50}O_{66}P_4Se_2^{5+}$ 872.4, found 872.7.

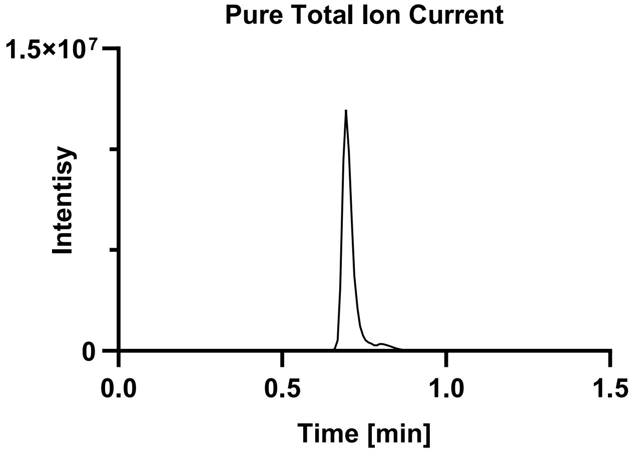

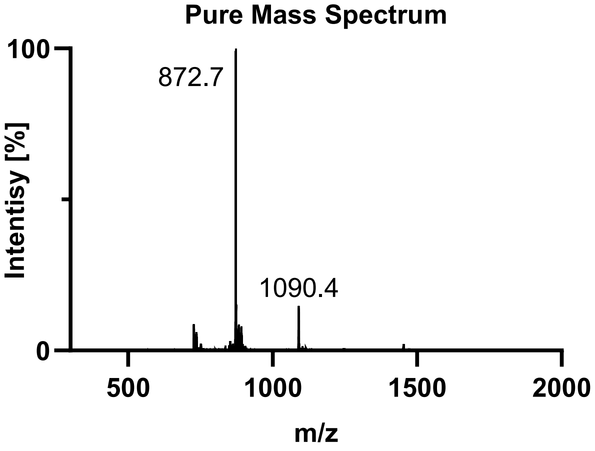

Tau(390-407) pS396 pS400 pT403 diselenide (**2l**).

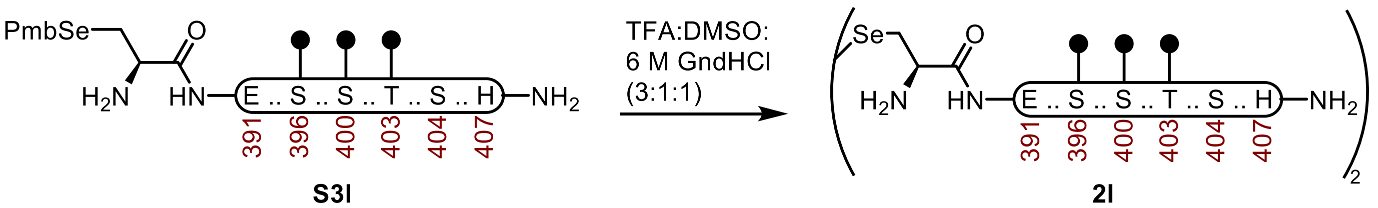

**Tau(390-407) pS396 pS400 pT403 diselenide (2l).** A 2-dram vial containing **S3l** (38.9 mg, 16.3 µmol) was dissolved in buffer (250 µL, 6 M Gnd·HCl, 0.1 M $Na_2HPO_4$, pH 7.2) and DMSO (250 µL). The mixture was treated with TFA (750 µL), wrapped with Al-Foil, and incubated at 23 ºC. After 40 min it was purified over a C4 column (10 x 250 mm, 10 µm, 100 Å, 4.7 mL/min) 7%-45% MeCN/$H_2O$/0.1% TFA over 30 min. The fractions were lyophilized to afford **2l** (33.1 mg, 90% yield).

**UPLC-MS:** Measured on an ACQUITY UPLC Protein BEH C4 column (2.1 x 50 mm; 1.7 µm; 300 Å; 50 °C; 0.6 mL/min) with a gradient of 5% MeCN/$H_2O$/0.1% FA until 0.1 min, 5-95% MeCN/$H_2O$/0.1% FA until 1.0 min, 95% MeCN/$H_2O$/0.1% FA until 1.1 min.

**Chemical Formula:** $C_{170}H_{278}N_{50}O_{72}P_6Se_2$
**Molecular Weight:** 4518.157 g/mol
**HRMS for [M+4H+]:** $C_{170}H_{282}N_{50}O_{72}P_6Se_2^{4+}$ 1130.2, found 1130.6.
**HRMS for [M+5H+]:** $C_{170}H_{283}N_{50}O_{72}P_6Se_2^{5+}$ 904.7, found 904.7.
**HRMS for [M+6H+]:** $C_{170}H_{284}N_{50}O_{72}P_6Se_2^{6+}$ 754.1, found 753.6.

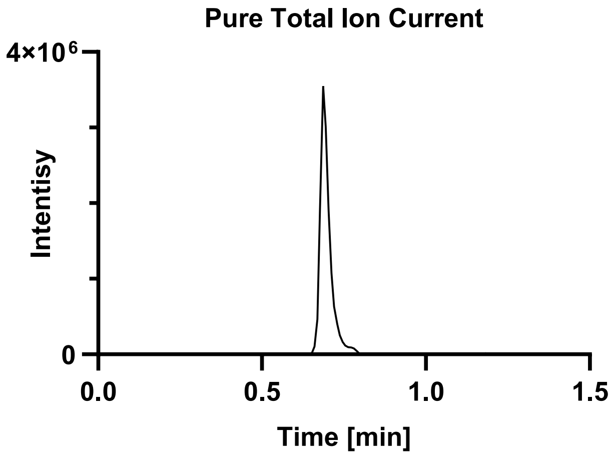

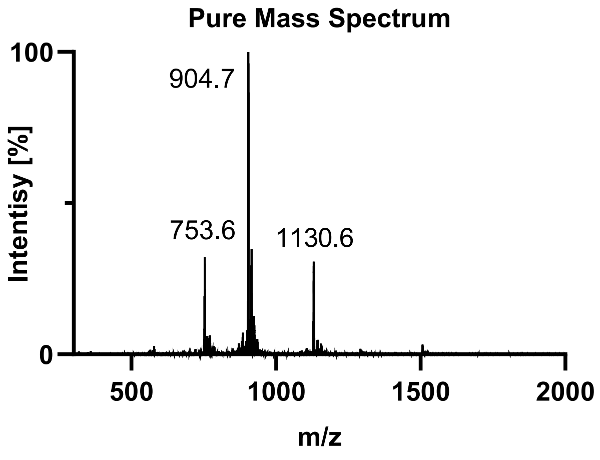

Tau(390-407) pS396 pS400 pS404 diselenide (**2m**).

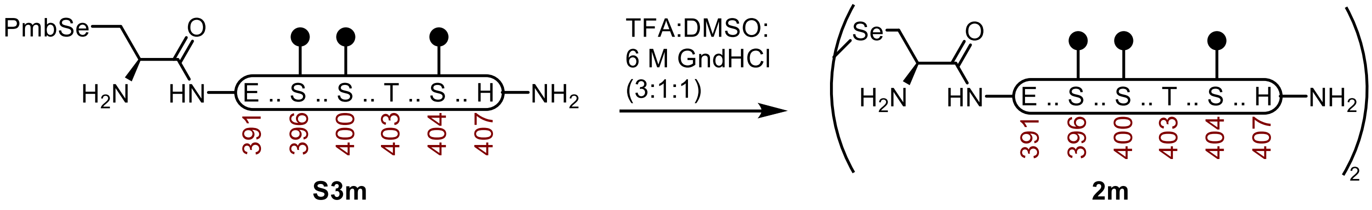

**Tau(390-407) pS396 pS400 pS404 diselenide (2m).** A 2-dram vial containing **S3m** (52.5 mg, 22.1 µmol) was dissolved in buffer (250 µL, 6 M Gnd·HCl, 0.1 M $Na_2HPO_4$, pH 7.2) and DMSO (250 µL). The mixture was treated with TFA (750 µL), wrapped with Al-Foil, and incubated at 23 ºC. After 30 min it was purified over a C4 column (10 x 250 mm, 10 µm, 100 Å, 4.7 mL/min) 7%-45% MeCN/$H_2O$/0.1% TFA over 30 min. The fractions were lyophilized to afford **2m** (39.2 mg, 78% yield).

**UPLC-MS:** Measured on an ACQUITY UPLC Protein BEH C4 column (2.1 x 50 mm; 1.7 µm; 300 Å; 50 °C; 0.6 mL/min) with a gradient of 5% MeCN/$H_2O$/0.1% FA until 0.1 min, 5-95% MeCN/$H_2O$/0.1% FA until 1.0 min, 95% MeCN/$H_2O$/0.1% FA until 1.1 min.

**Chemical Formula:** $C_{170}H_{278}N_{50}O_{72}P_6Se_2$
**Molecular Weight:** 4518.157 g/mol
**HRMS for [M+4H+]:** $C_{170}H_{282}N_{50}O_{72}P_6Se_2^{4+}$ 1130.2, found 1130.3.
**HRMS for [M+5H+]:** $C_{170}H_{283}N_{50}O_{72}P_6Se_2^{5+}$ 904.7, found 904.7.

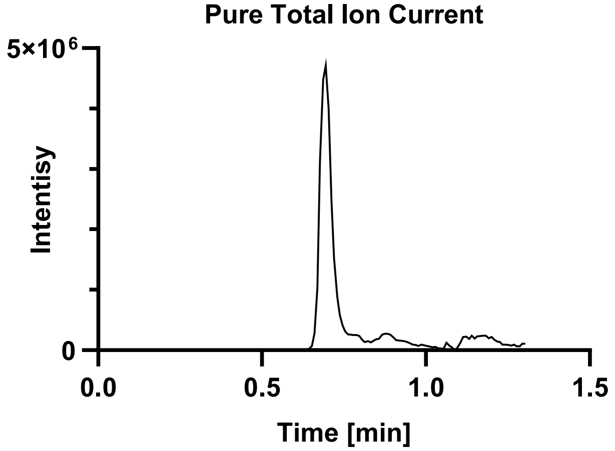

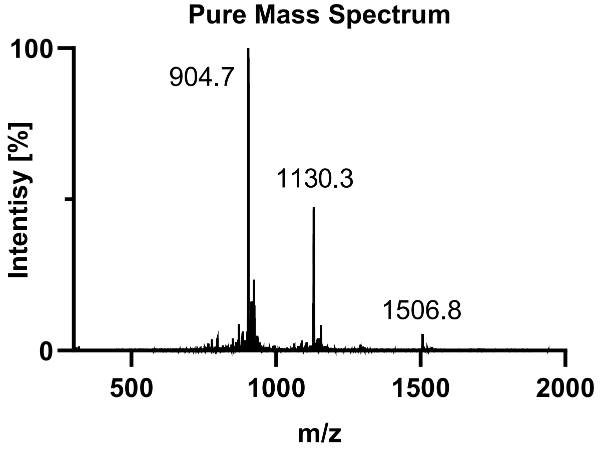

Tau(390-407) pS396 pT403 pS404 diselenide (**2n**).

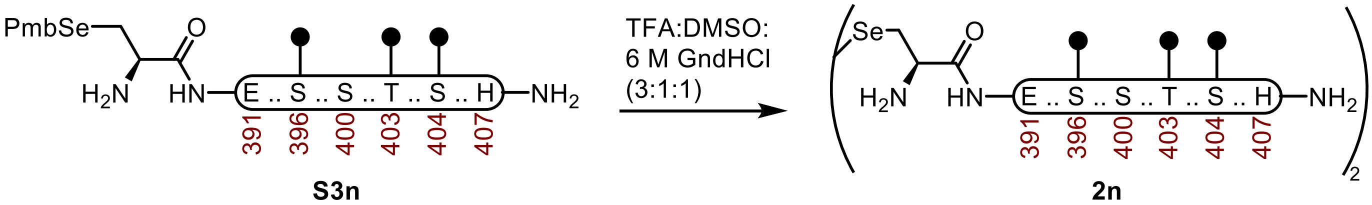

**Tau(390-407) pS396 pT403 pS404 diselenide (2n).** A 2-dram vial containing **S3n** (61.2 mg, 25.7 µmol) was dissolved in buffer (250 µL, 6 M Gnd·HCl, 0.1 M $Na_2HPO_4$, pH 7.2) and DMSO (250 µL). The mixture was treated with TFA (750 µL), wrapped with Al-Foil, and incubated at 23 ºC. After 40 min it was purified over a C4 column (10 x 250 mm, 10 µm, 100 Å, 4.7 mL/min) 7%-45% MeCN/$H_2O$/0.1% TFA over 30 min. The fractions were lyophilized to afford **2n** (48.8 mg, 84% yield).

**UPLC-MS:** Measured on an ACQUITY UPLC Protein BEH C4 column (2.1 x 50 mm; 1.7 µm; 300 Å; 50 °C; 0.6 mL/min) with a gradient of 5% MeCN/$H_2O$/0.1% FA until 0.1 min, 5-95% MeCN/$H_2O$/0.1% FA until 1.0 min, 95% MeCN/$H_2O$/0.1% FA until 1.1 min.

**Chemical Formula:** $C_{170}H_{278}N_{50}O_{72}P_6Se_2$
**Molecular Weight:** 4518.157 g/mol
**HRMS for [M+4H+]:** $C_{170}H_{282}N_{50}O_{72}P_6Se_2^{4+}$ 1130.2, found 1130.6.
**HRMS for [M+5H+]:** $C_{170}H_{283}N_{50}O_{72}P_6Se_2^{5+}$ 904.7, found 904.7.

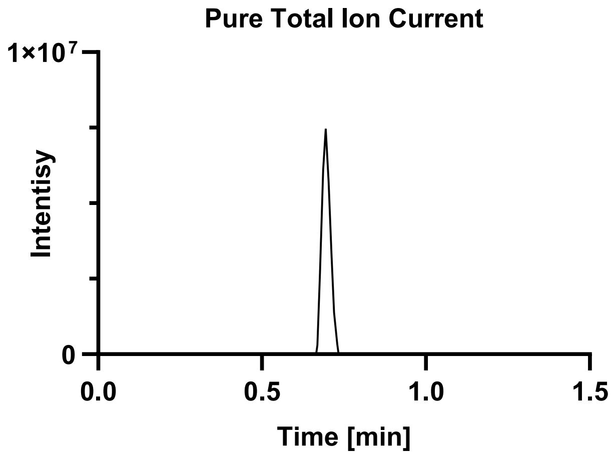

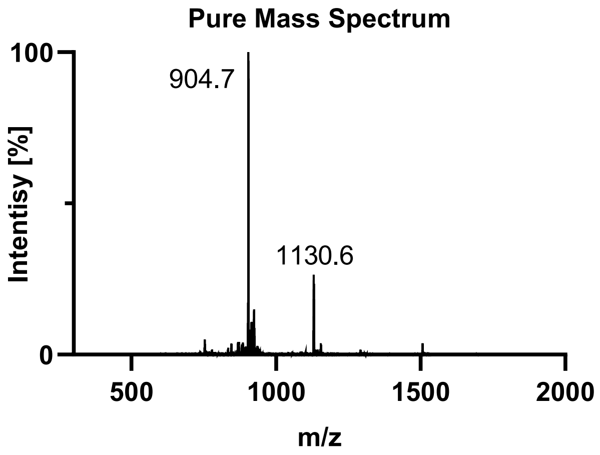

Tau(390-407) pS400 pT403 pS404 diselenide (**2o**).

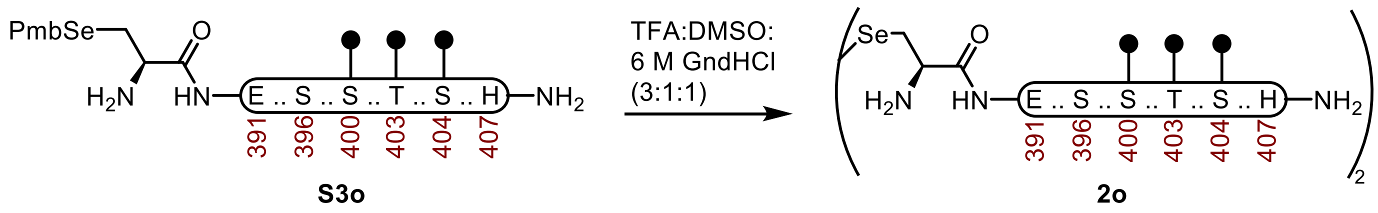

**Tau(390-407) pS400 pT403 pS404 diselenide (2o).** A 2-dram vial containing **S3o** (62.1 mg, 26.1 µmol) was dissolved in buffer (250 µL, 6 M Gnd·HCl, 0.1 M $Na_2HPO_4$, pH 7.2) and DMSO (250 µL). The mixture was treated with TFA (750 µL), wrapped with Al-Foil, and incubated at 23 ºC. After 30 min it was purified over a C4 column (10 x 250 mm, 10 µm, 100 Å, 4.7 mL/min) 7%-45% $MeCN/H_2O$/0.1% TFA over 30 min. The fractions were lyophilized to afford **2o** (53.6 mg, 91% yield).

**UPLC-MS:** Measured on an ACQUITY UPLC Protein BEH C4 column (2.1 x 50 mm; 1.7 µm; 300 Å; 50 °C; 0.6 mL/min) with a gradient of 5% $MeCN/H_2O$/0.1% FA until 0.1 min, 5-95% $MeCN/H_2O$/0.1% FA until 1.0 min, 95% $MeCN/H_2O$/0.1% FA until 1.1 min.

**Chemical Formula:** $C_{170}H_{278}N_{50}O_{72}P_6Se_2$
**Molecular Weight:** 4518.157 g/mol
**HRMS for [M+4H+]:** $C_{170}H_{282}N_{50}O_{72}P_6Se_2^{4+}$ 1130.2, found 1130.3.
**HRMS for [M+5H+]:** $C_{170}H_{283}N_{50}O_{72}P_6Se_2^{5+}$ 904.7, found 904.7.
**HRMS for [M+6H+]:** $C_{170}H_{284}N_{50}O_{72}P_6Se_2^{6+}$ 754.1, found 753.6.

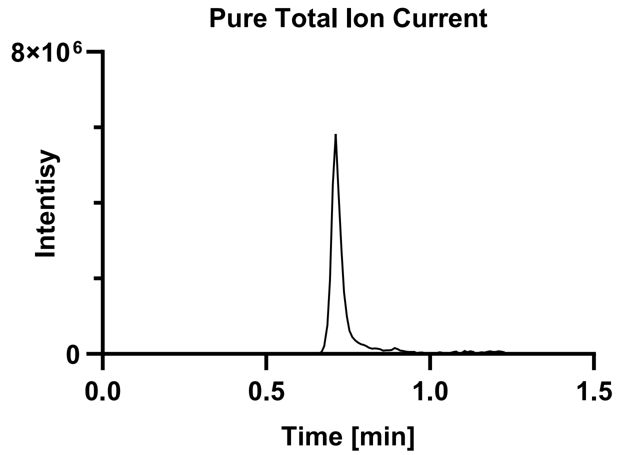

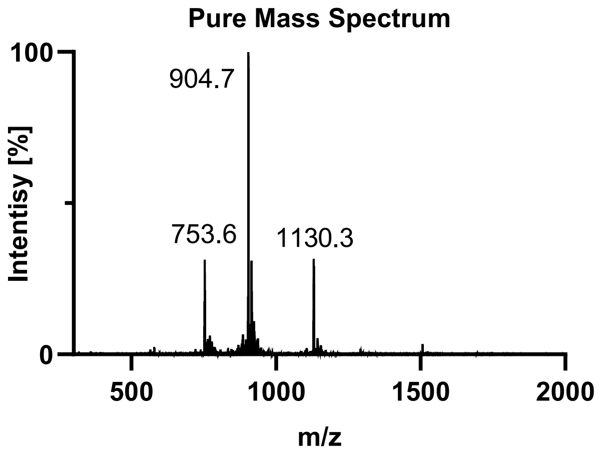

### S3. SEMI-SYNTHESIS OF PHOSPHORYLATED TAU(297-407).

#### Tau(297-407) WT (**3a**).

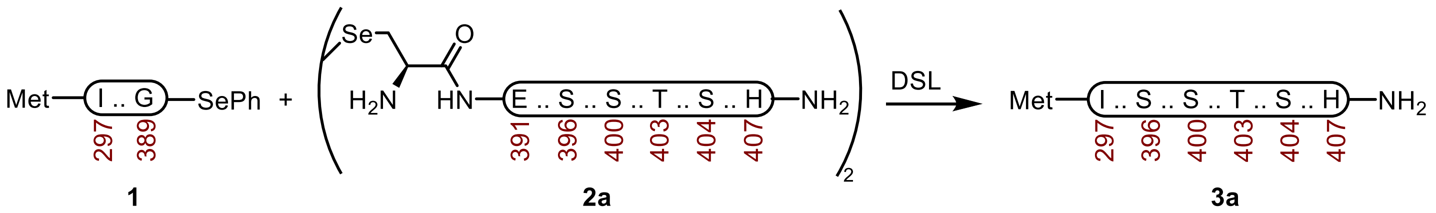

**Tau(297-407) WT (3a).** A 20 mL vial with a stir-bar containing a selenoester **1** (42.2 mg, 4.12 μmol) and diselenide **2a** (17.1 mg, 4.23 μmol) was treated with Ar-purged DSL buffer (1.0 mL, 6 M Gnd·HCl, 0.1 M $Na_2HPO_4$, pH 6.8), and the vial was carefully sealed under a stream of Ar. After the reaction was mixed at 23 ºC for 2 h, the DPDS was extracted with Ar-purged hexanes (5 x 1 mL) under a stream of Ar. The reaction was treated with Ar-purged deselenization buffer (2.0 mL, 200 mM TCEP·HCl, 100 mM DTT, 6 M Gnd·HCl, 100 mM HEPES, pH 6.9) and it was mixed at 23 ºC. After 24 h, it was purified by sample displacement mode chromatography (Microsorb C18, 4.6 x 250 mm, 5 μm, 100 Å, 50 ºC, 1 mL/min, 5%-65% MeCN/$H_2O$/0.1% TFA over 60 min), and the fractions were lyophilized to afford **3a** (30.5 mg, 62% yield).

**UPLC-HRMS:** Measured on an ACQUITY UPLC Protein BEH C4 column (2.1 x 50 mm; 1.7 μm; 300 Å; 55 °C; 0.6 mL/min) with a gradient of 5% MeCN/$H_2O$/0.1% FA until 0.1 min, 5-95% MeCN/$H_2O$/0.1% FA until 1.0 min, 95% MeCN/$H_2O$/0.1% FA until 1.1 min.

**Calculated Mass:** $C_{527}H_{862}N_{158}O_{159}S_2$ 12019.8, 12178.5 found.

**Crude UPLC-HRMS:**

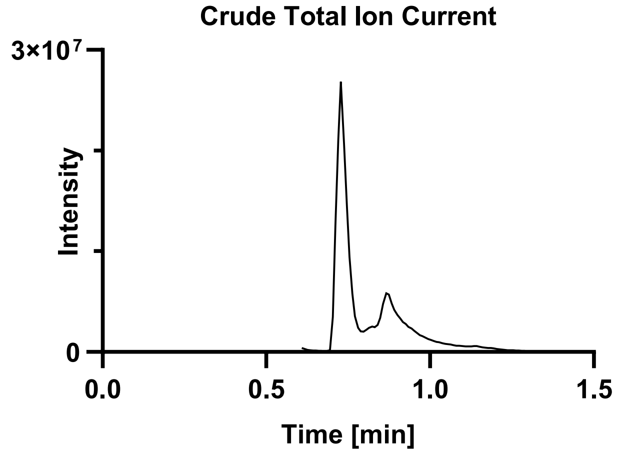

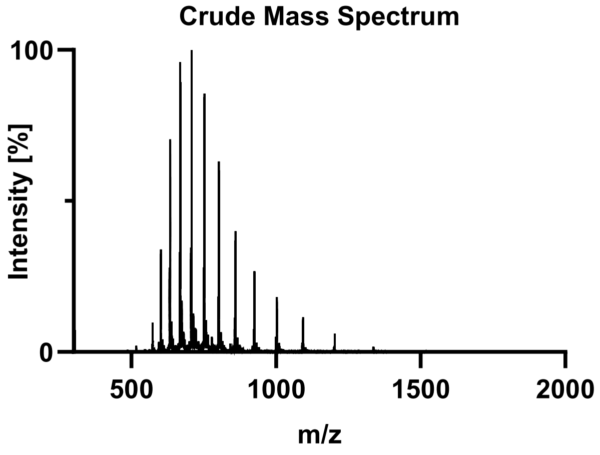

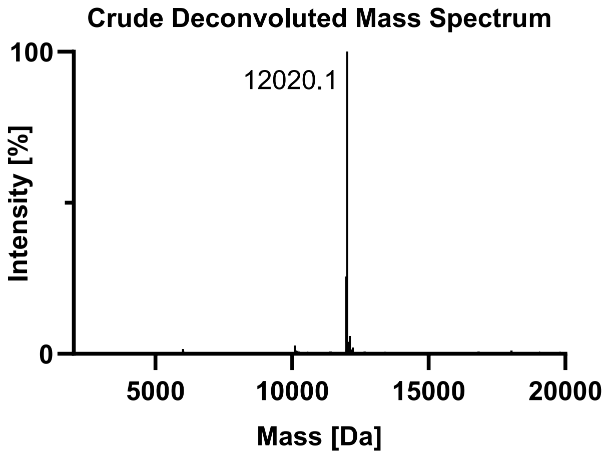

**Pure UPLC-HRMS:**

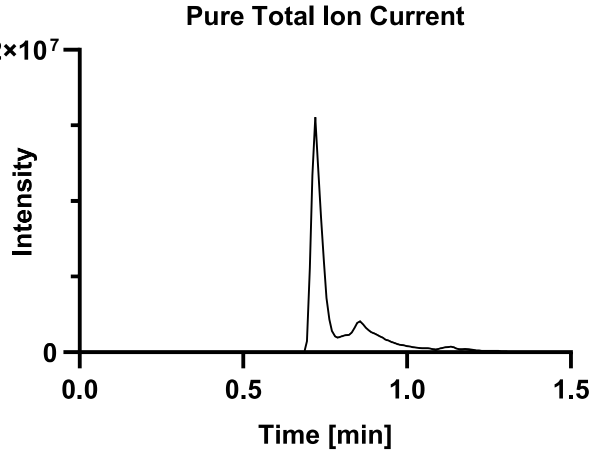

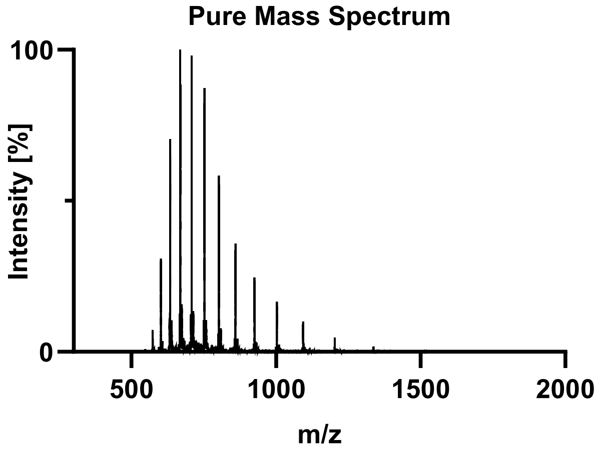

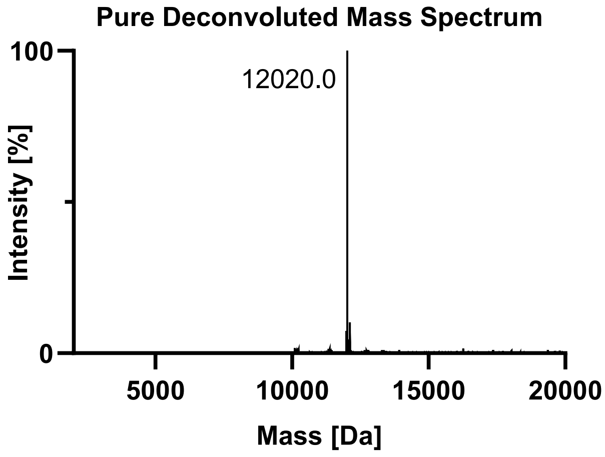

Tau(297-407) pS396 (**3b**).

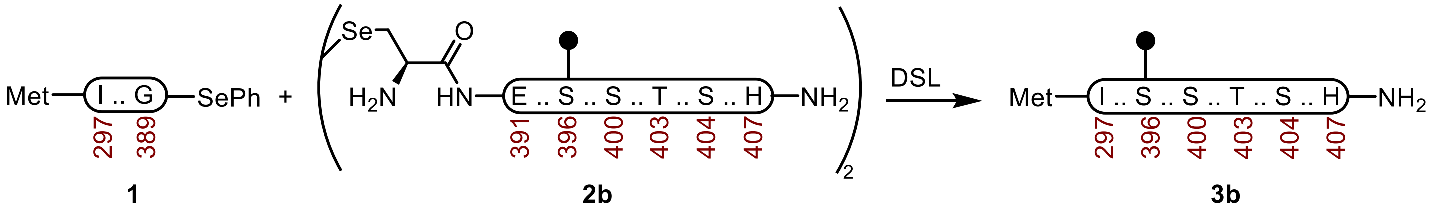

**Tau(297-407) pS396 (3b).** A 20 mL vial with a stir-bar containing a selenoester **1** (37.9 mg, 3.70 µmol) and diselenide **2b** (8.9 mg, 2.12 µmol) was treated with Ar-purged DSL buffer (1.0 mL, 6 M Gnd·HCl, 0.1 M $Na_2HPO_4$, pH 6.9), and the vial was carefully sealed under a stream of Ar. After the reaction was mixed at 23 ºC for 2.5 h, the DPDS was extracted with Ar-purged hexanes (5 x 1 mL) under a stream of Ar. The reaction was treated with Ar-purged deselenization buffer (2.0 mL, 200 mM TCEP·HCl, 100 mM DTT, 6 M Gnd·HCl, 100 mM HEPES, pH 6.8) and it was mixed at 23 ºC. After 24 h, it was purified over a C4 column (10 x 250 mm, 10 µm, 100 Å, 4.7 mL/min) 10%-45% MeCN/$H_2O$/0.1% TFA over 30 min. The material was further purified by sample displacement mode chromatography (Microsorb C18, 4.6 x 250 mm, 5 µm, 100 Å, 50 ºC, 1 mL/min, 5%-65% MeCN/$H_2O$/0.1% TFA over 60 min), and the fractions were lyophilized to afford **3b** (18.8 mg, 42% yield).

**UPLC-HRMS:** Measured on an ACQUITY UPLC Protein BEH C4 column (2.1 x 50 mm; 1.7 µm; 300 Å; 55 °C; 0.6 mL/min) with a gradient of 5% MeCN/$H_2O$/0.1% FA until 0.1 min, 5-95% MeCN/$H_2O$/0.1% FA until 1.0 min, 95% MeCN/$H_2O$/0.1% FA until 1.1 min.

**Calculated Mass:** $C_{527}H_{863}N_{158}O_{162}PS_2$ 12099.7, 12099.5 found.

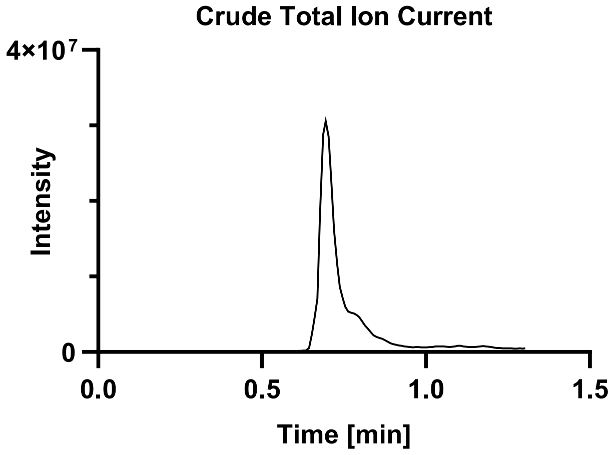

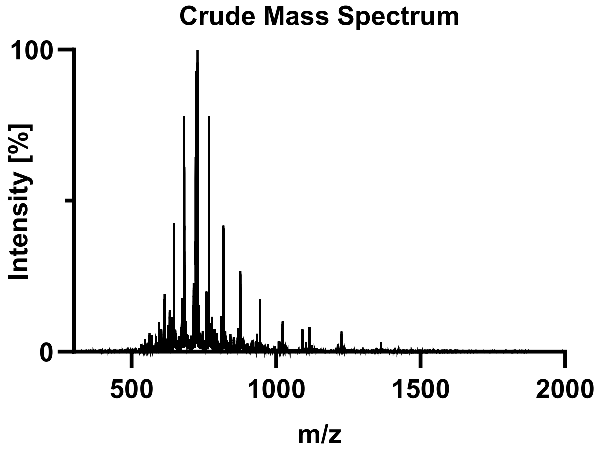

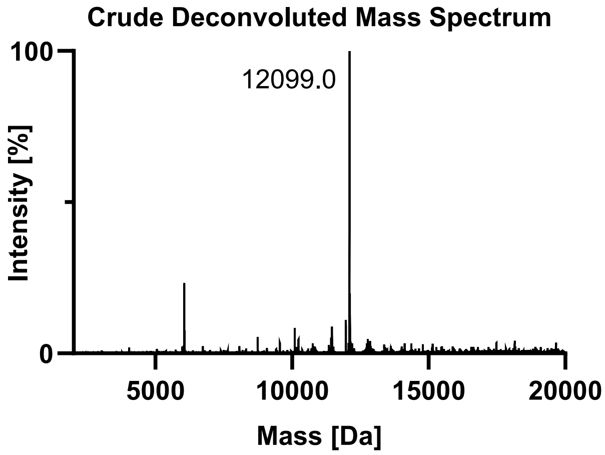

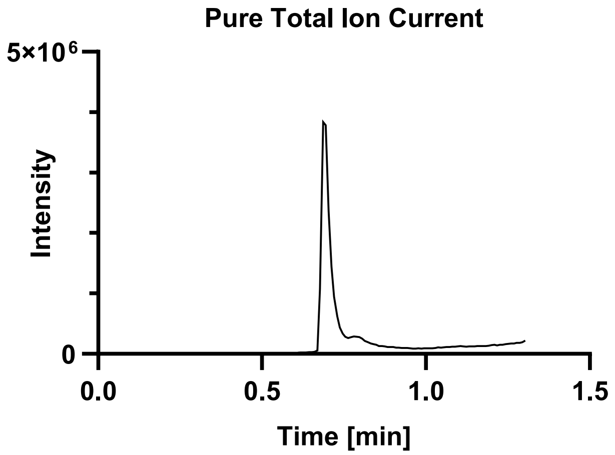
Pure Total Ion Current
5×10⁶
0
Intensity
0.0
0.5
1.0
1.5
Time [min]

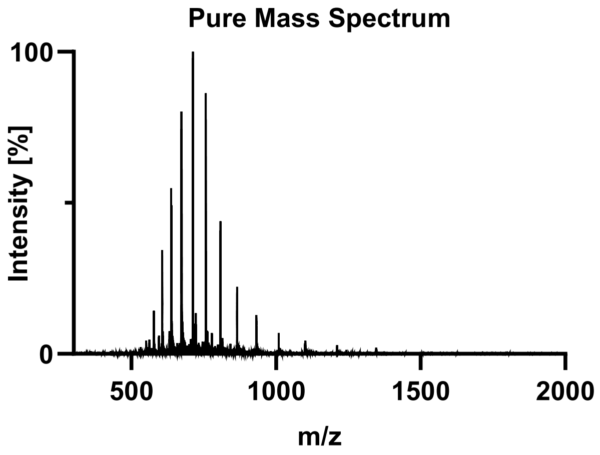
Pure Mass Spectrum
100
0
Intensity [%]
500
1000
1500
2000
m/z

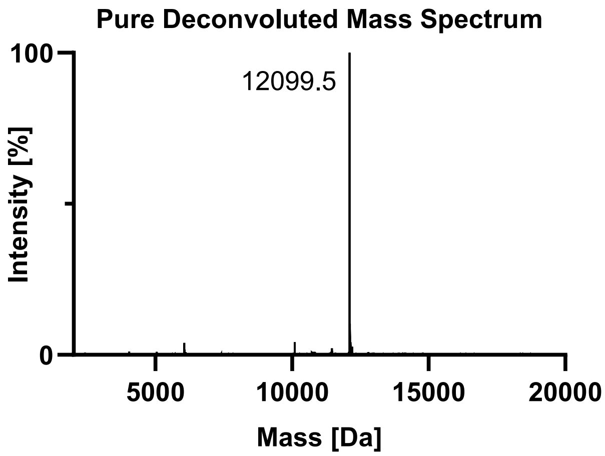
Pure Deconvoluted Mass Spectrum
100
0
12099.5
Intensity [%]
5000
10000
15000
20000
Mass [Da]

Tau(297-407) pS400 (**3c**).

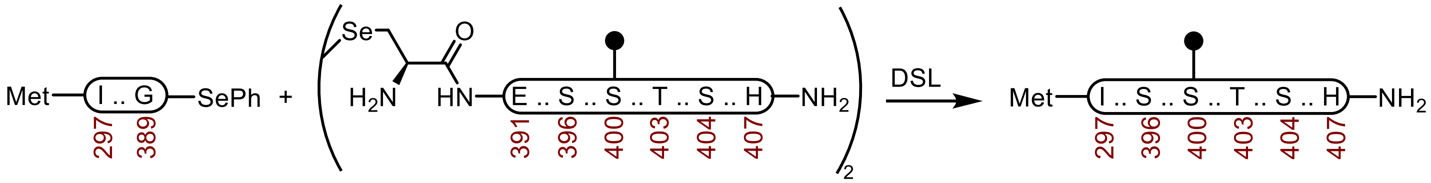

**1** **2c** **3c**

**Tau(297-407) pS400 (3c).** A 20 mL vial with a stir-bar containing a selenoester **1** (38.8 mg, 3.79 μmol) and diselenide **2c** (9.5 mg, 2.26 μmol) was treated with Ar-purged DSL buffer (1.0 mL, 6 M Gnd·HCl, 0.1 M $Na_2HPO_4$, pH 6.9), and the vial was carefully sealed under a stream of Ar. After the reaction was mixed at 23 ºC for 2.5 h, the DPDS was extracted with Ar-purged hexanes (5 x 1 mL) under a stream of Ar. The reaction was treated with Ar-purged deselenization buffer (2.0 mL, 200 mM TCEP·HCl, 100 mM DTT, 6 M Gnd·HCl, 100 mM HEPES, pH 6.8) and it was mixed at 23 ºC. After 24 h, it was purified over a C4 column (10 x 250 mm, 10 μm, 100 Å, 4.7 mL/min) 10%-45% MeCN/$H_2O$/0.1% TFA over 30 min. The material was further purified by sample displacement mode chromatography (Microsorb C18, 4.6 x 250 mm, 5 μm, 100 Å, 50 ºC, 1 mL/min, 5%-65% MeCN/$H_2O$/0.1% TFA over 60 min), and the fractions were lyophilized to afford **3c** (16.7 mg, 36% yield).

**UPLC-HRMS:** Measured on an ACQUITY UPLC Protein BEH C4 column (2.1 x 50 mm; 1.7 μm; 300 Å; 55 °C; 0.6 mL/min) with a gradient of 5% MeCN/$H_2O$/0.1% FA until 0.1 min, 5-95% MeCN/$H_2O$/0.1% FA until 1.0 min, 95% MeCN/$H_2O$/0.1% FA until 1.1 min.

**Calculated Mass:** $C_{527}H_{863}N_{158}O_{162}PS_2$ 12099.7, 12099.5 found.

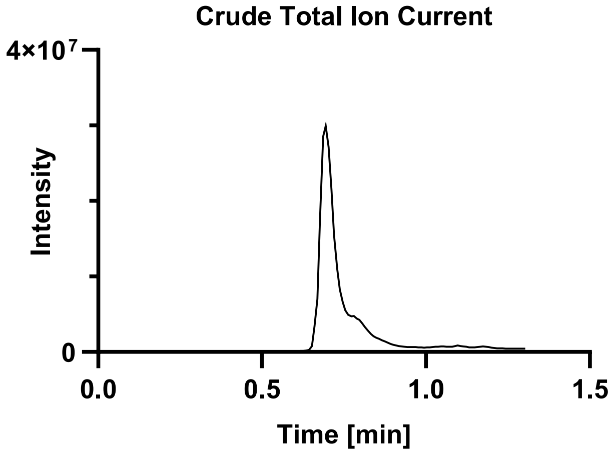

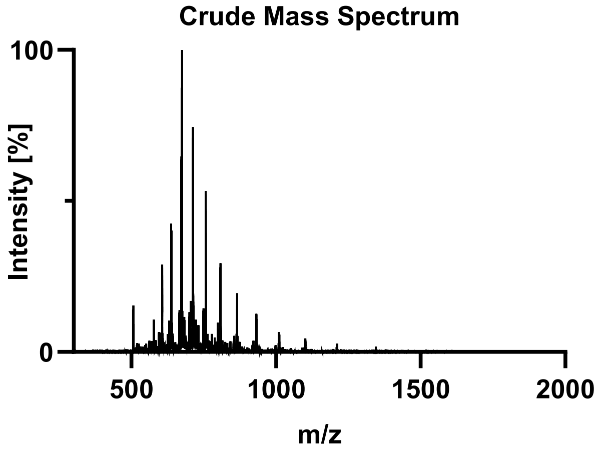

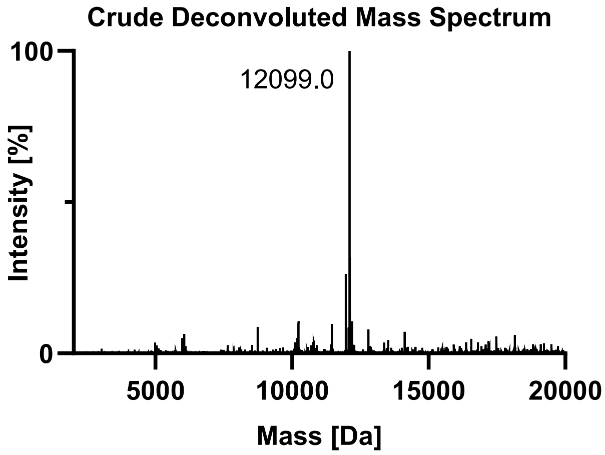

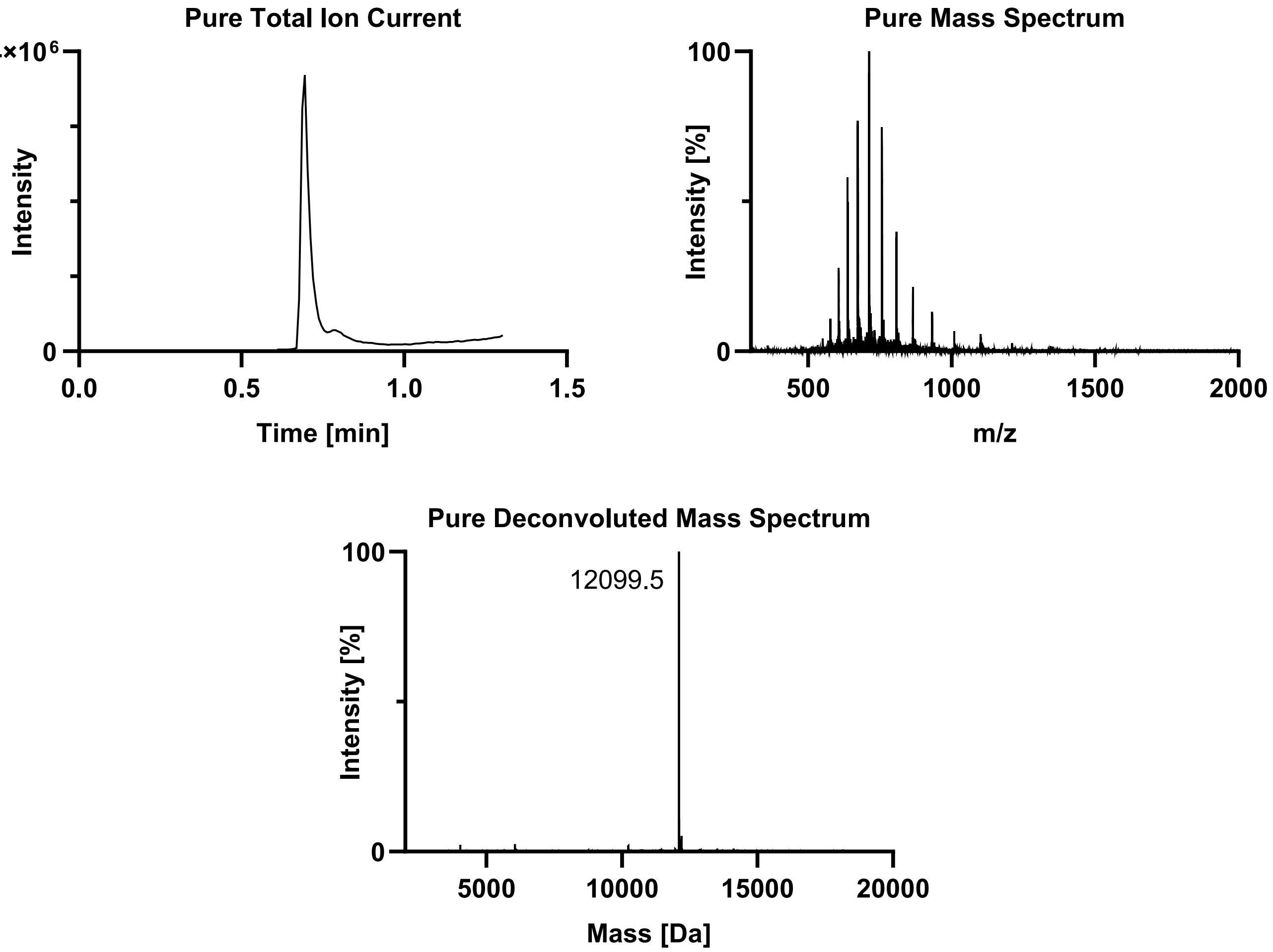
Pure Total Ion Current
4×10^6
Intensity
0
0.0
0.5
1.0
1.5
Time [min]
Pure Mass Spectrum
100
Intensity [%]
0
500
1000
1500
2000
m/z
Pure Deconvoluted Mass Spectrum
100
12099.5
Intensity [%]
0
5000
10000
15000
20000
Mass [Da]

Tau(297-407) pT403 (**3d**).

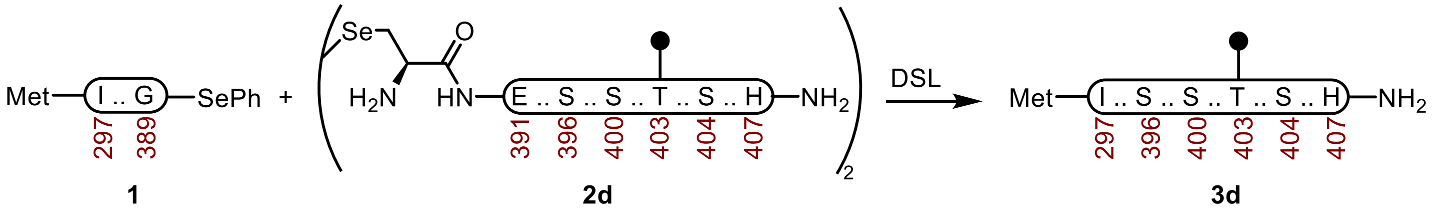

**Tau(297-407) pT403 (3d).** A 20 mL vial with a stir-bar containing a selenoester **1** (41.6 mg, 4.06 μmol) and diselenide **2d** (10.8 mg, 2.57 μmol) was treated with Ar-purged DSL buffer (1.0 mL, 6 M Gnd·HCl, 0.1 M $Na_2HPO_4$, pH 7.0), and the vial was carefully sealed under a stream of Ar. After the reaction was mixed at 23 ºC for 1.5 h, the DPDS was extracted with Ar-purged hexanes (5 x 1 mL) under a stream of Ar. The reaction was treated with Ar-purged deselenization buffer (2.0 mL, 200 mM TCEP·HCl, 100 mM DTT, 6 M Gnd·HCl, 100 mM HEPES, pH 7.2) and it was mixed at 23 ºC. After 24 h, it was purified by sample displacement mode chromatography (Microsorb C18, 4.6 x 250 mm, 5 μm, 100 Å, 50 ºC, 1 mL/min, 5%-65% MeCN/$H_2O$/0.1% TFA over 60 min), and the fractions were lyophilized to afford **3d** (24.1 mg, 49% yield).

**UPLC-HRMS:** Measured on an ACQUITY UPLC Protein BEH C4 column (2.1 x 50 mm; 1.7 μm; 300 Å; 55 °C; 0.6 mL/min) with a gradient of 5% MeCN/$H_2O$/0.1% FA until 0.1 min, 5-95% MeCN/$H_2O$/0.1% FA until 1.0 min, 95% MeCN/$H_2O$/0.1% FA until 1.1 min.

**Calculated Mass:** $C_{527}H_{863}N_{158}O_{162}PS_2$ 12099.7, 12099.0 found.

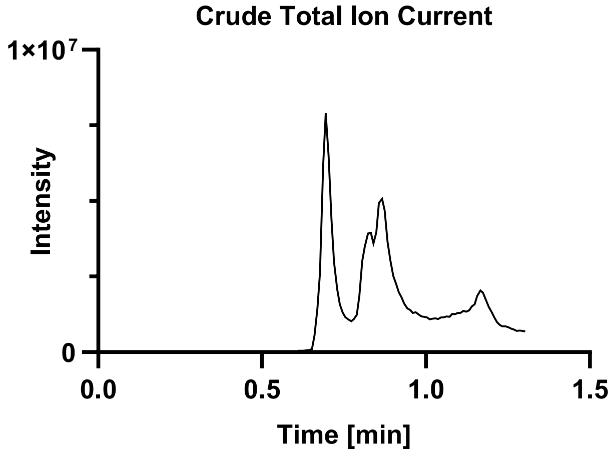

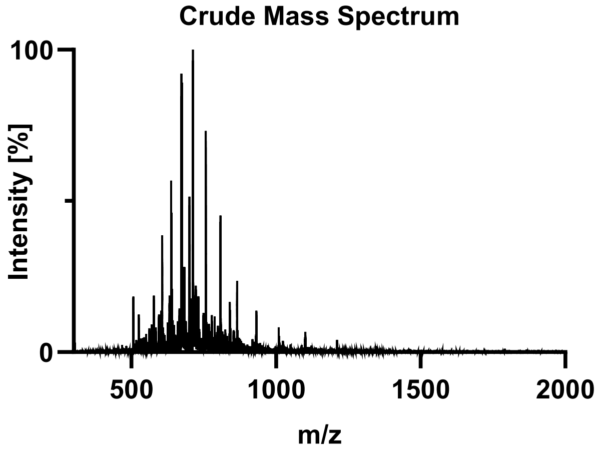

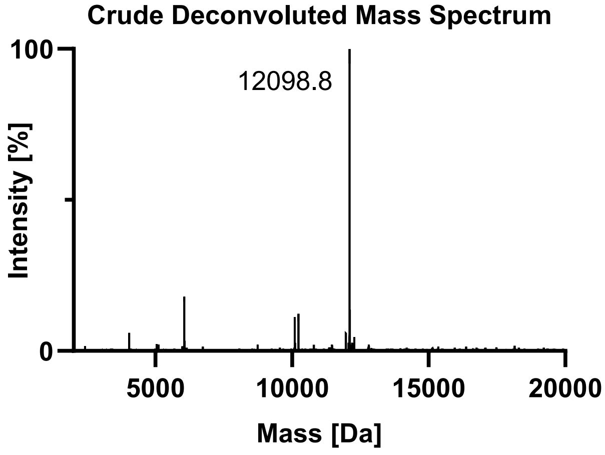

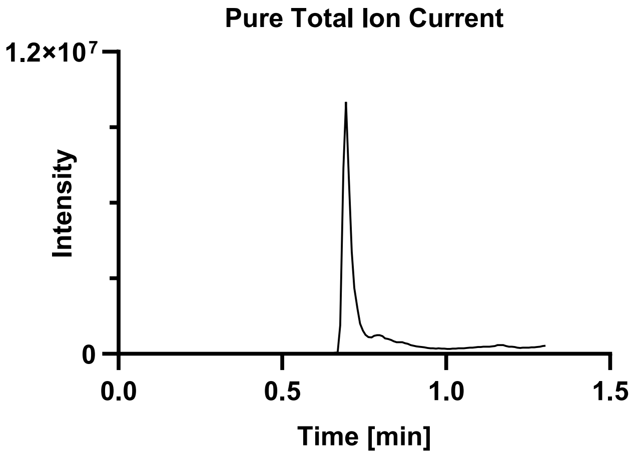
Pure Total Ion Current
1.2×10^7
0
Intensity
0.0
0.5
1.0
1.5
Time [min]

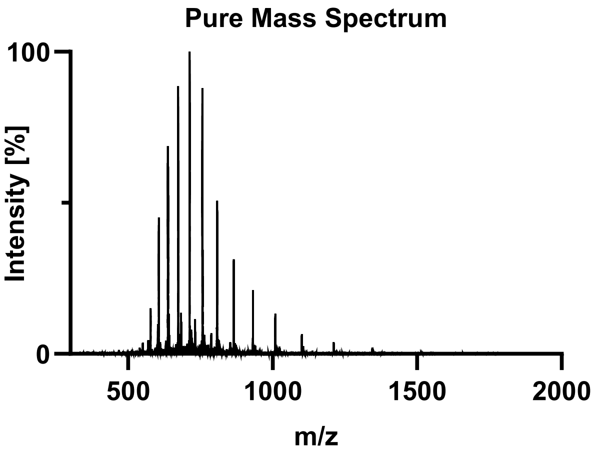
Pure Mass Spectrum
100
0
Intensity [%]
500
1000
1500
2000
m/z

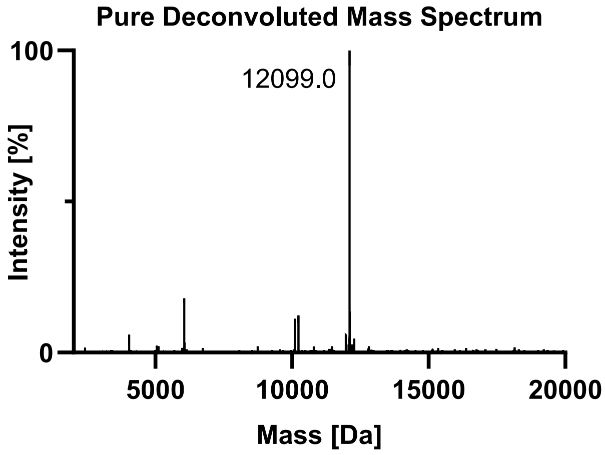
Pure Deconvoluted Mass Spectrum
100
0
12099.0
Intensity [%]
5000
10000
15000
20000
Mass [Da]

Tau(297-407) pS404 (**3e**).

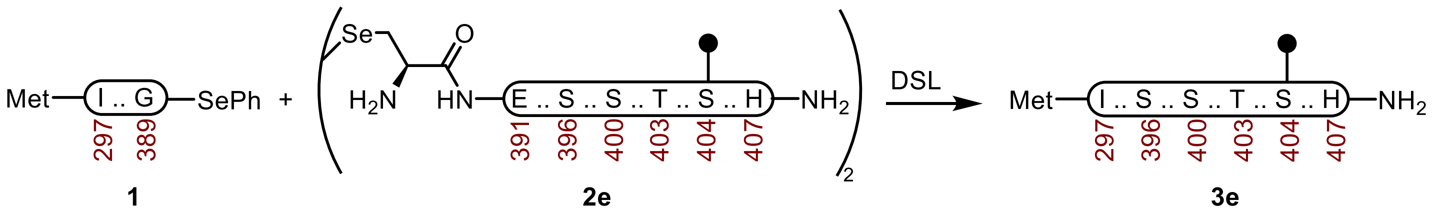

**Tau(297-407) pS404 (3e).** A 20 mL vial with a stir-bar containing a selenoester **1** (40.4 mg, 3.95 µmol) and diselenide **2e** (10.1 mg, 2.41 µmol) was treated with Ar-purged DSL buffer (1.0 mL, 6 M Gnd·HCl, 0.1 M $Na_2HPO_4$, pH 7.0), and the vial was carefully sealed under a stream of Ar. After the reaction was mixed at 23 ºC for 2 h, the DPDS was extracted with Ar-purged hexanes (5 x 1 mL) under a stream of Ar. The reaction was treated with Ar-purged deselenization buffer (2.0 mL, 200 mM TCEP·HCl, 100 mM DTT, 6 M Gnd·HCl, 100 mM HEPES, pH 7.2) and it was mixed at 23 ºC. After 24 h, it was purified by sample displacement mode chromatography (Microsorb C18, 4.6 x 250 mm, 5 µm, 100 Å, 50 ºC, 1 mL/min, 5%-65% MeCN/$H_2O$/0.1% TFA over 60 min), and the fractions were lyophilized to afford **3e** (20.6 mg, 43% yield).

**UPLC-HRMS:** Measured on an ACQUITY UPLC Protein BEH C4 column (2.1 x 50 mm; 1.7 µm; 300 Å; 55 °C; 0.6 mL/min) with a gradient of 5% MeCN/$H_2O$/0.1% FA until 0.1 min, 5-95% MeCN/$H_2O$/0.1% FA until 1.0 min, 95% MeCN/$H_2O$/0.1% FA until 1.1 min.

**Calculated Mass:** $C_{527}H_{863}N_{158}O_{162}PS_2$ 12099.7, 12099.0 found.

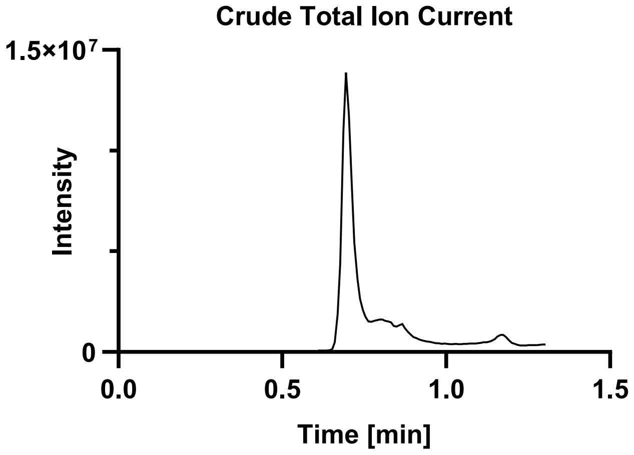

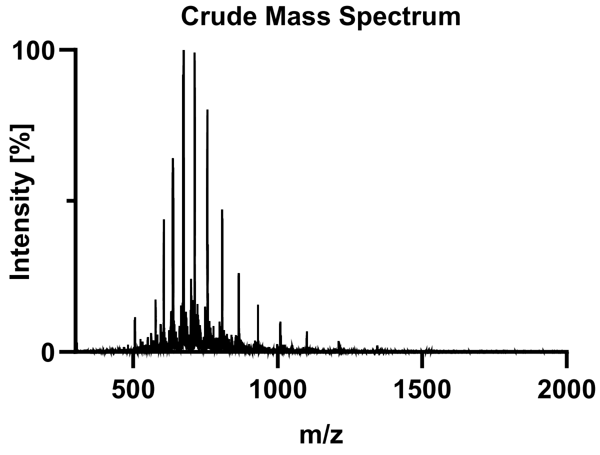

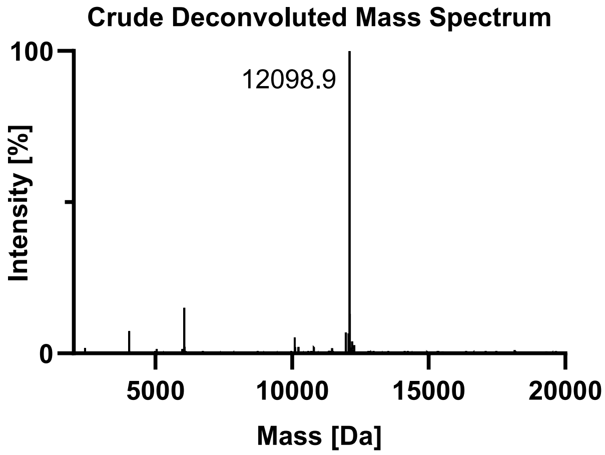

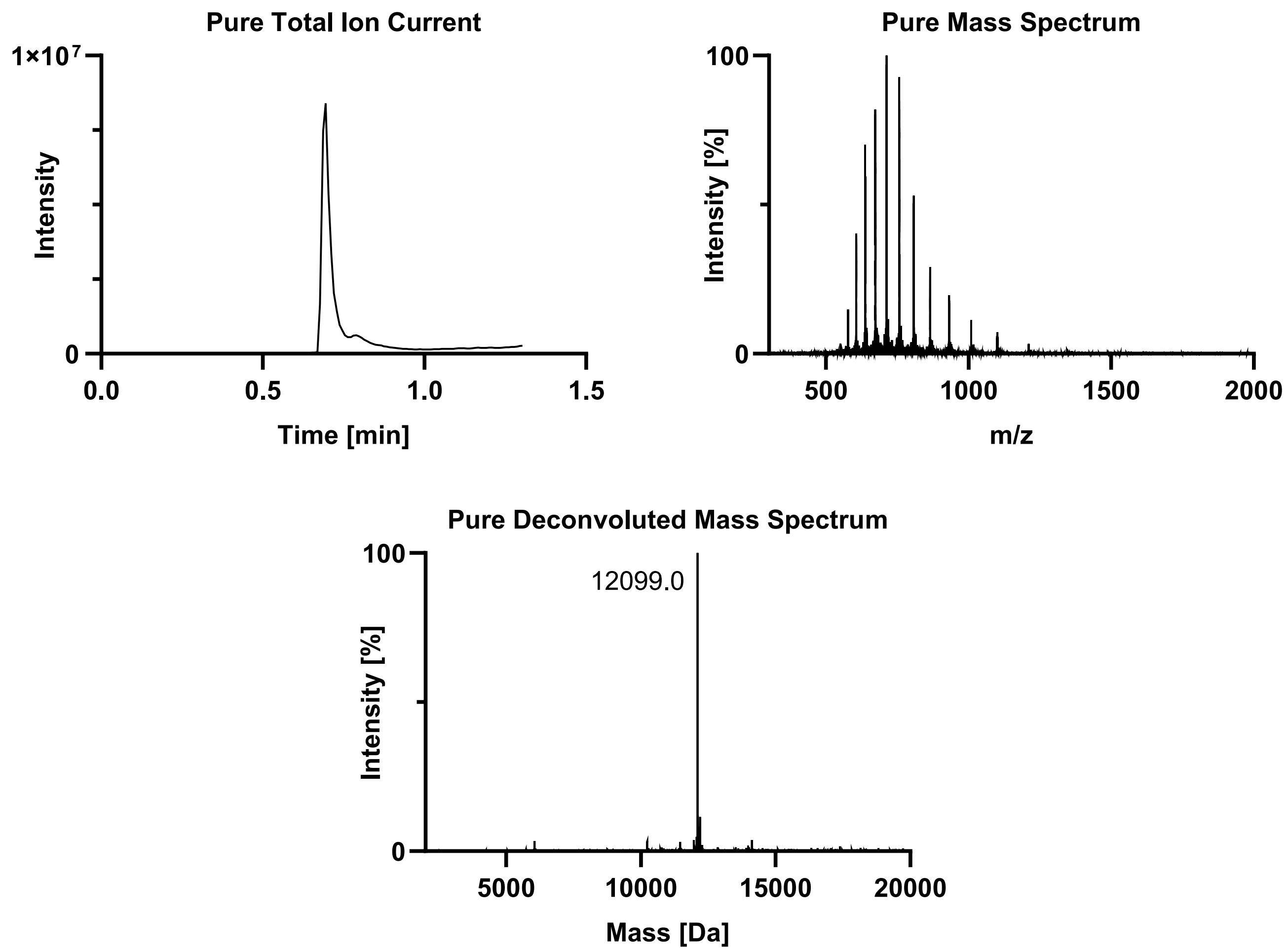
Pure Total Ion Current
1×10^7
0
Intensity
0.0
0.5
1.0
1.5
Time [min]
Pure Mass Spectrum
100
0
Intensity [%]
500
1000
1500
2000
m/z
Pure Deconvoluted Mass Spectrum
100
12099.0
0
Intensity [%]
5000
10000
15000
20000
Mass [Da]

Tau(297-407) pS396 + pS400 (**3f**).

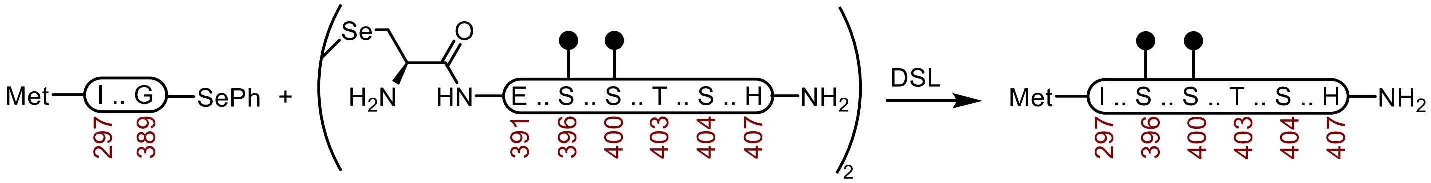

**Tau(297-407) pS396 pS400 (3f).** A 1-dram vial with a stir-bar containing a selenoester **1** (39.8 mg, 3.89 µmol) and diselenide **2f** (14.1 mg, 3.23 µmol) was treated with Ar-purged DSL buffer (1.0 mL, 6 M Gnd·HCl, 0.1 M $Na_2HPO_4$, pH 6.8), and the vial was carefully sealed under a stream of Ar. After the reaction was mixed at 23 ºC for 2 h, the DPDS was extracted with Ar-purged hexanes (5 x 1 mL) under a stream of Ar. The reaction was treated with Ar-purged deselenization buffer (2.0 mL, 200 mM TCEP·HCl, 100 mM DTT, 6 M Gnd·HCl, 100 mM HEPES, pH 7.0) and it was mixed at 23 ºC. After 24 h, it was purified by sample displacement mode chromatography (Microsorb C18, 4.6 x 250 mm, 5 µm, 100 Å, 50 ºC, 1 mL/min, 5%-65% MeCN/$H_2O$/0.1% TFA over 60 min), and the fractions were lyophilized to afford **3f** (26.4 mg, 56% yield).

**UPLC-HRMS:** Measured on an ACQUITY UPLC Protein BEH C4 column (2.1 x 50 mm; 1.7 µm; 300 Å; 55 °C; 0.6 mL/min) with a gradient of 5% MeCN/$H_2O$/0.1% FA until 0.1 min, 5-95% MeCN/$H_2O$/0.1% FA until 1.0 min, 95% MeCN/$H_2O$/0.1% FA until 1.1 min.

**Calculated Mass:** $C_{527}H_{864}N_{158}O_{165}P_2S_2$ 12179.7, 12180.2 found.

**Crude UPLC-HRMS:**

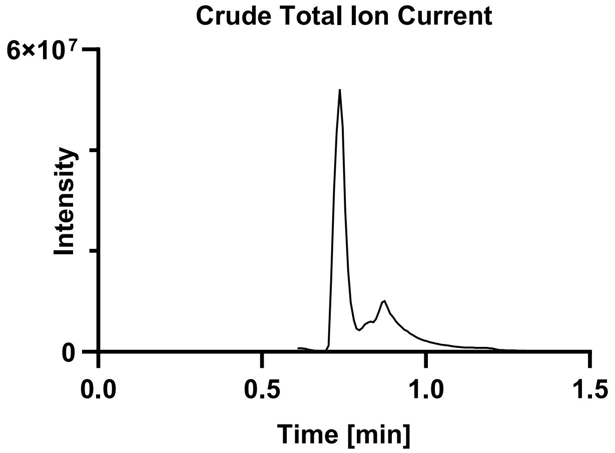

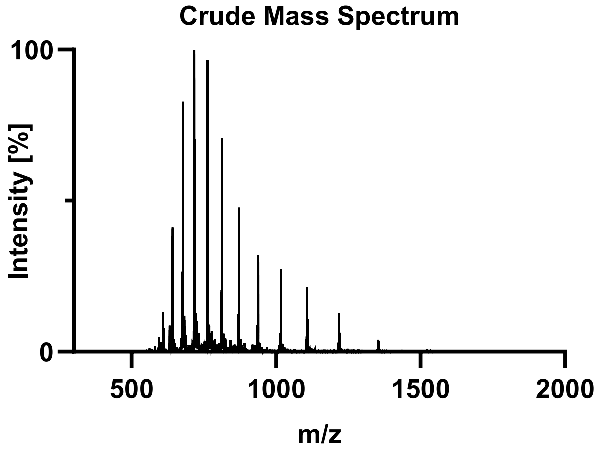

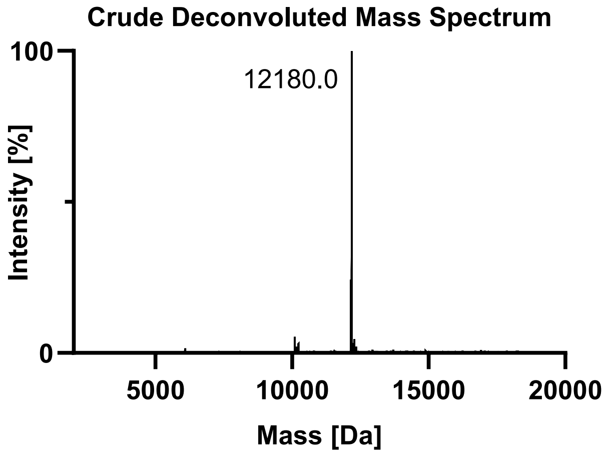

**Pure UPLC-HRMS:**

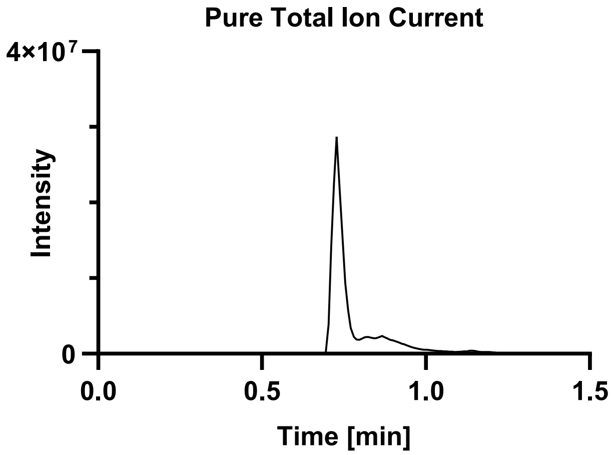

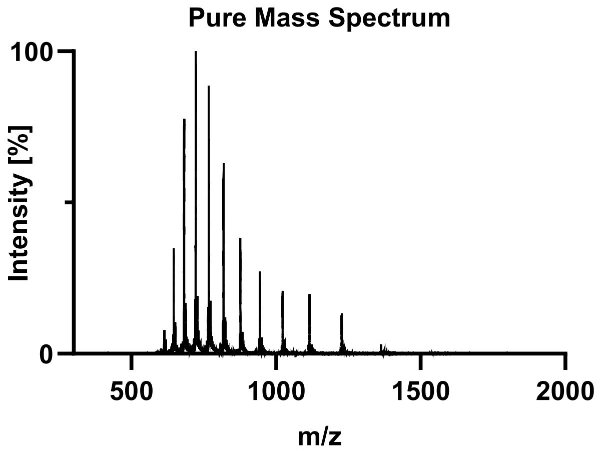

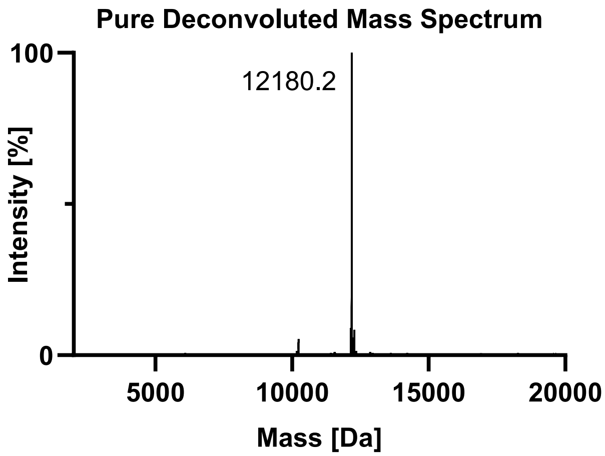

Tau(297-407) pS396 pT403 (**3g**).

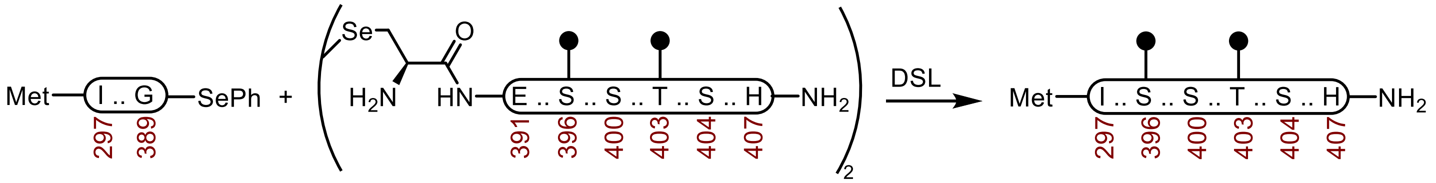

**Tau(297-407) pS396 pT403 (3g).** A 20 mL vial with a stir-bar containing a selenoester **1** (38.6 mg, 3.77 µmol) and diselenide **2g** (10.1 mg, 2.32 µmol) was treated with Ar-purged DSL buffer (0.75 mL, 6 M Gnd·HCl, 0.1 M $Na_2HPO_4$, pH 6.8), and the vial was carefully sealed under a stream of Ar. After the reaction was mixed at 23 °C for 2 h, the DPDS was extracted with Ar-purged hexanes (5 × 1 mL) under a stream of Ar. The reaction was treated with Ar-purged deselenization buffer (1.5 mL, 200 mM TCEP·HCl, 100 mM DTT, 6 M Gnd·HCl, 100 mM HEPES, pH 7.0) and it was mixed at 23 ºC. After 24 h, it was purified over a C4 column (10 x 250 mm, 10 µm, 100 Å, 4.7 mL/min) 10%-45% MeCN/$H_2O$/0.1% TFA over 30 min. The material was further purified by sample displacement mode chromatography (Microsorb C18, 4.6 x 250 mm, 5 µm, 100 Å, 50 ºC, 1 mL/min, 5%-65% MeCN/$H_2O$/0.1% TFA over 60 min), and the fractions were lyophilized to afford **3g** (24.5 mg, 53% yield).

**UPLC-HRMS:** Measured on an ACQUITY UPLC Protein BEH C4 column (2.1 x 50 mm; 1.7 µm; 300 Å; 55 °C; 0.6 mL/min) with a gradient of 5% MeCN/$H_2O$/0.1% FA until 0.1 min, 5-95% MeCN/$H_2O$/0.1% FA until 1.0 min, 95% MeCN/$H_2O$/0.1% FA until 1.1 min.

**Calculated Mass:** $C_{527}H_{864}N_{158}O_{165}P_2S_2$ 12179.7, 12178.5 found.

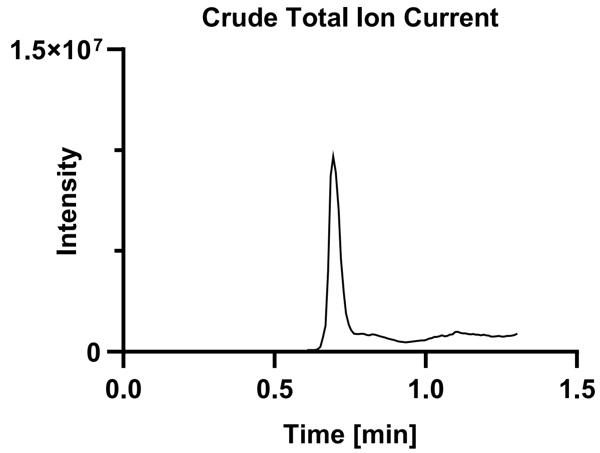

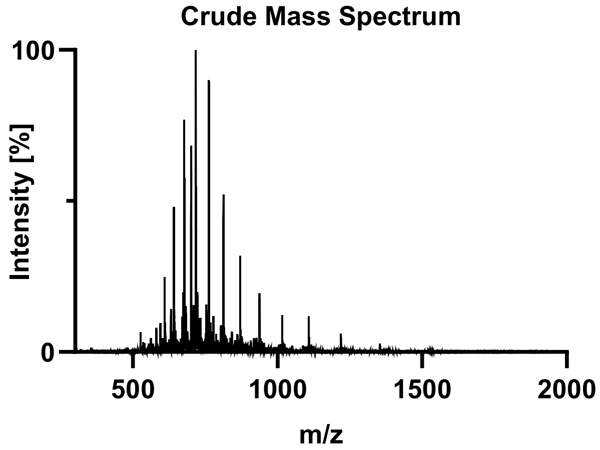

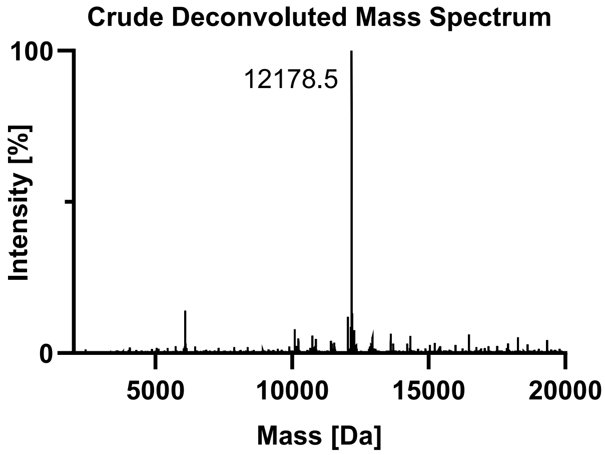

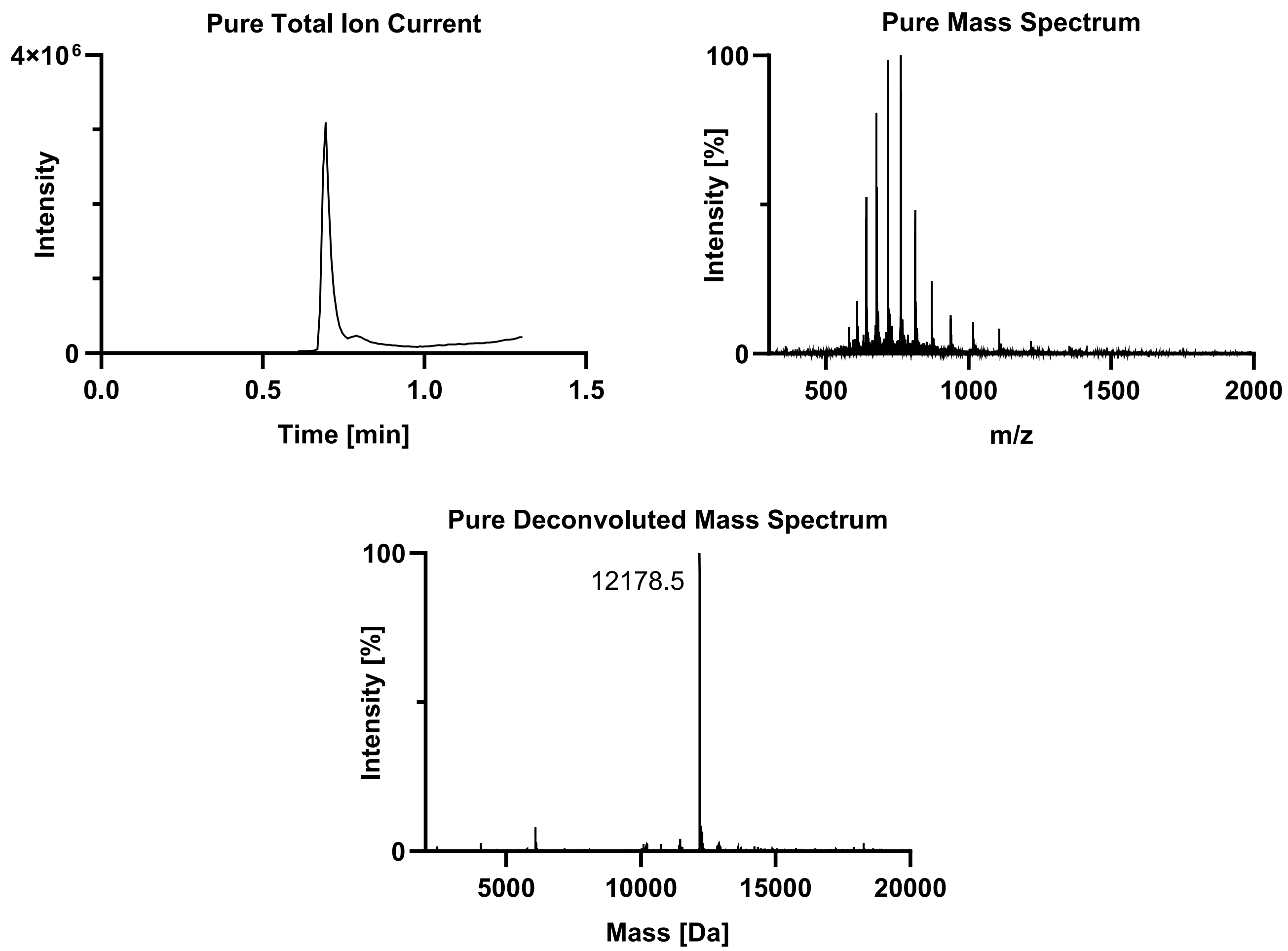
Pure Total Ion Current
4×10^6
0
Intensity
0.0
0.5
1.0
1.5
Time [min]
Pure Mass Spectrum
100
0
Intensity [%]
500
1000
1500
2000
m/z
Pure Deconvoluted Mass Spectrum
100
0
Intensity [%]
12178.5
5000
10000
15000
20000
Mass [Da]

Tau(297-407) pS396 pS404 (**3h**).

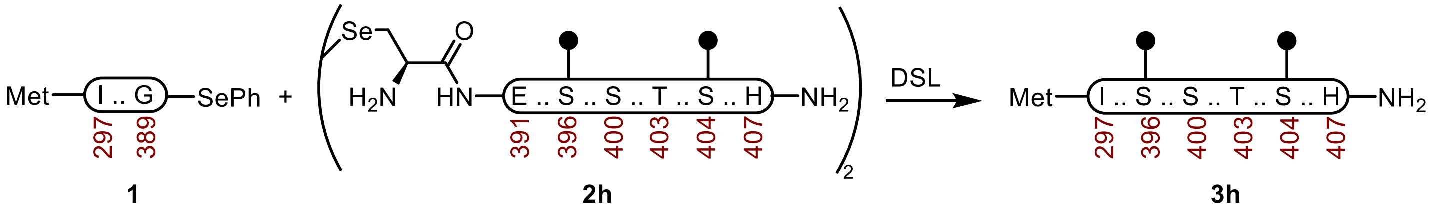

**Tau(297-407) pS396 pS404 (3h).** A 20 mL vial with a stir-bar containing a selenoester **1** (33.2 mg, 3.24 µmol) and diselenide **2h** (7.0 mg, 1.61 µmol) was treated with Ar-purged DSL buffer (0.75 mL, 6 M Gnd·HCl, 0.1 M $Na_2HPO_4$, pH 6.8), and the vial was carefully sealed under a stream of Ar. After the reaction was mixed at 23 ºC for 2 h, the DPDS was extracted with Ar-purged hexanes (5 x 1 mL) under a stream of Ar. The reaction was treated with Ar-purged deselenization buffer (1.5 mL, 200 mM TCEP·HCl, 100 mM DTT, 6 M Gnd·HCl, 100 mM HEPES, pH X) and it was mixed at 23 ºC. After 24 h, it was purified over a C4 column (10 x 250 mm, 10 µm, 100 Å, 4.7 mL/min) 10%-45% MeCN/$H_2O$/0.1% TFA over 30 min. The material was further purified by sample displacement mode chromatography (Microsorb C18, 4.6 x 250 mm, 5 µm, 100 Å, 50 ºC, 1 mL/min, 5%-65% MeCN/$H_2O$/0.1% TFA over 60 min), and the fractions were lyophilized to afford **3h** (16.2 mg, 41% yield).

**UPLC-HRMS:** Measured on an ACQUITY UPLC Protein BEH C4 column (2.1 x 50 mm; 1.7 µm; 300 Å; 55 °C; 0.6 mL/min) with a gradient of 5% MeCN/$H_2O$/0.1% FA until 0.1 min, 5-95% MeCN/$H_2O$/0.1% FA until 1.0 min, 95% MeCN/$H_2O$/0.1% FA until 1.1 min.

**Calculated Mass:** $C_{527}H_{864}N_{158}O_{165}P_2S_2$ 12179.7, 12179.0 found.

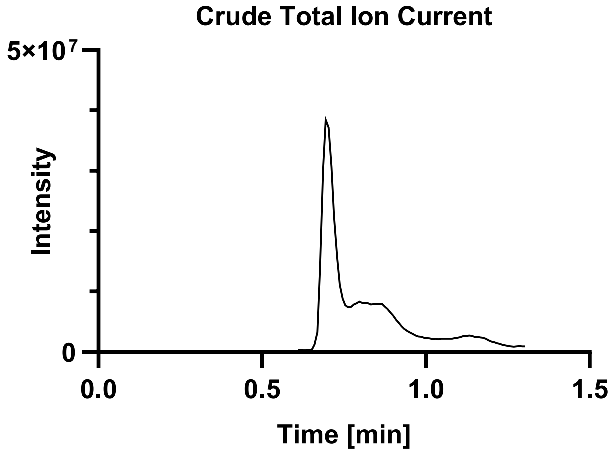

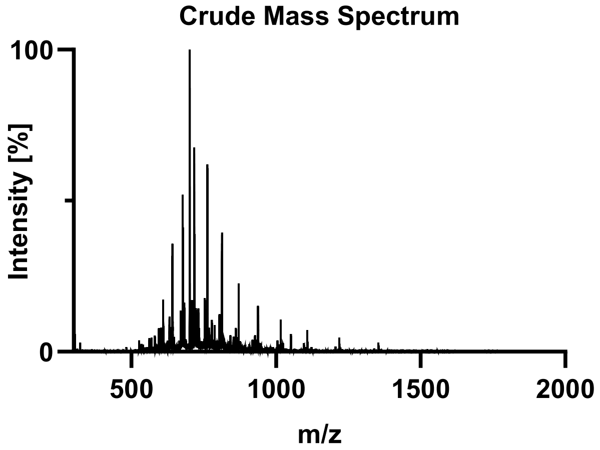

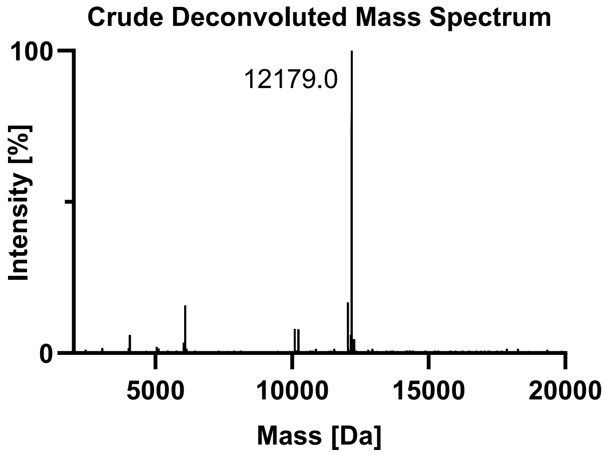

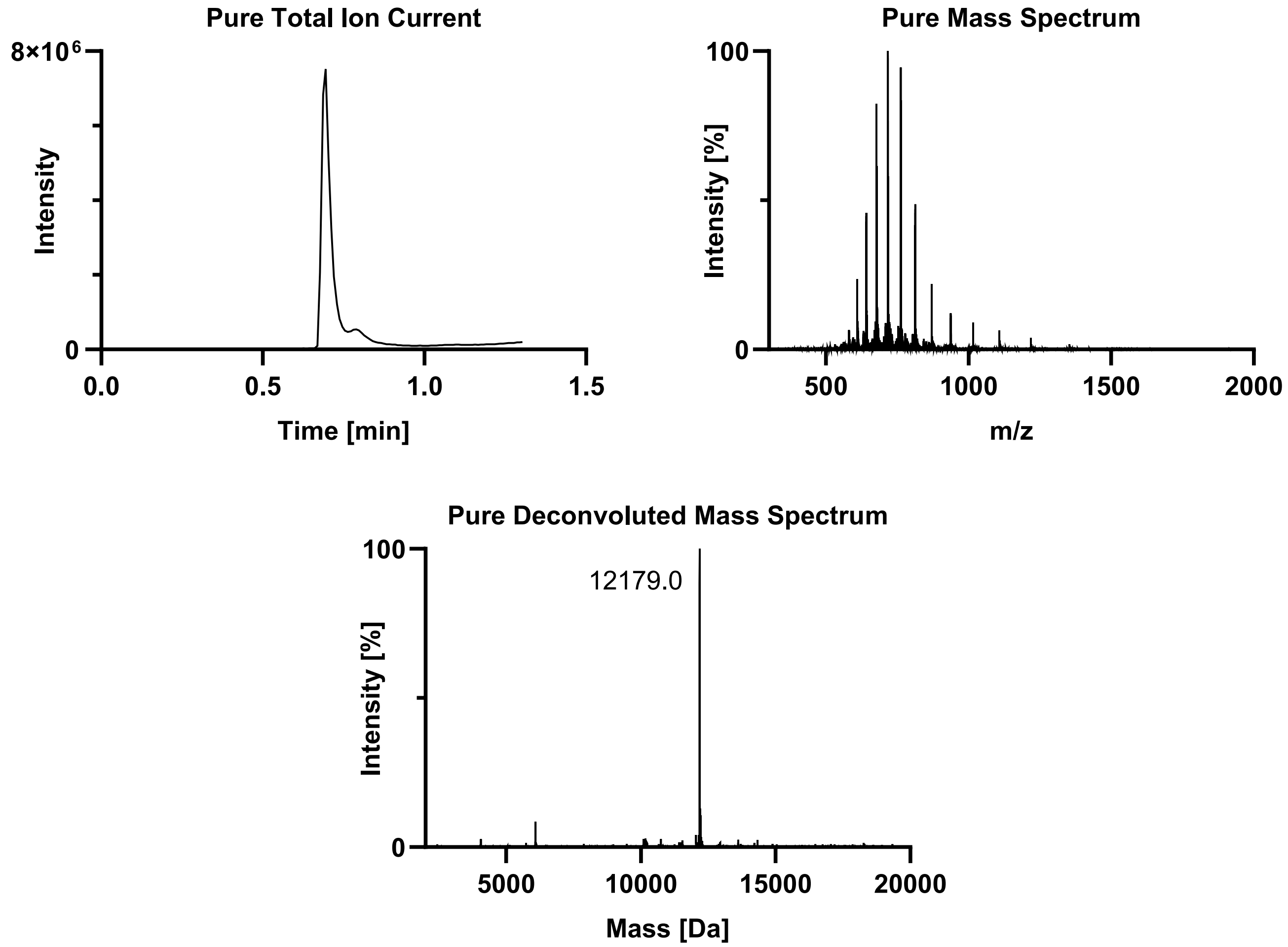
Pure Total Ion Current
8×10^6
0
Intensity
0.0
0.5
1.0
1.5
Time [min]
Pure Mass Spectrum
100
0
Intensity [%]
500
1000
1500
2000
m/z
Pure Deconvoluted Mass Spectrum
100
0
12179.0
Intensity [%]
5000
10000
15000
20000
Mass [Da]

Tau(297-407) pS400 pT403 (**3i**).

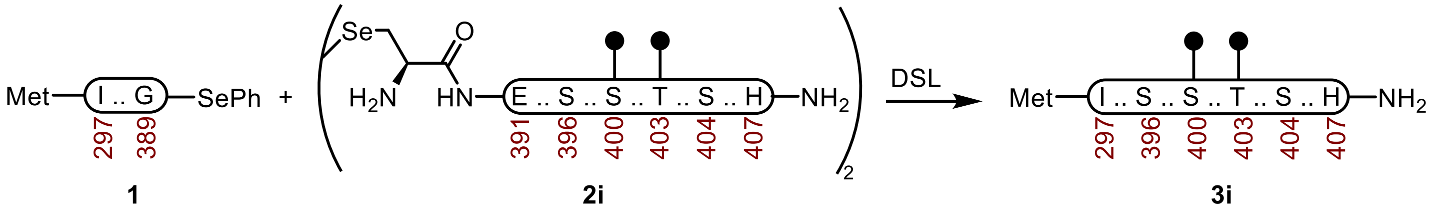

**Tau(297-407) pS400 pT403 (3i).** A 20 mL vial with a stir-bar containing a selenoester **1** (32.8 mg, 3.20 μmol) and diselenide **2i** (7.6 mg, 1.74 μmol) was treated with Ar-purged DSL buffer (0.75 mL, 6 M Gnd·HCl, 0.1 M $Na_2HPO_4$, pH 6.8), and the vial was carefully sealed under a stream of Ar. After the reaction was mixed at 23 °C for 2 h, the DPDS was extracted with Ar-purged hexanes (5 × 1 mL) under a stream of Ar. The reaction was treated with Ar-purged deselenization buffer (1.5 mL, 200 mM TCEP·HCl, 100 mM DTT, 6 M Gnd·HCl, 100 mM HEPES, pH 6.8) and it was mixed at 23 ºC. After 24 h, it was purified over a C4 column (10 x 250 mm, 10 μm, 100 Å, 4.7 mL/min) 10%-45% MeCN/$H_2O$/0.1% TFA over 30 min. The material was further purified by sample displacement mode chromatography (Microsorb C18, 4.6 x 250 mm, 5 μm, 100 Å, 50 ºC, 1 mL/min, 5%-65% MeCN/$H_2O$/0.1% TFA over 60 min), and the fractions were lyophilized to afford **3i** (13.4 mg, 34% yield).

**UPLC-HRMS:** Measured on an ACQUITY UPLC Protein BEH C4 column (2.1 x 50 mm; 1.7 μm; 300 Å; 55 °C; 0.6 mL/min) with a gradient of 5% MeCN/$H_2O$/0.1% FA until 0.1 min, 5-95% MeCN/$H_2O$/0.1% FA until 1.0 min, 95% MeCN/$H_2O$/0.1% FA until 1.1 min.

**Calculated Mass:** $C_{527}H_{864}N_{158}O_{165}P_2S_2$ 12179.7, 12178.5 found.

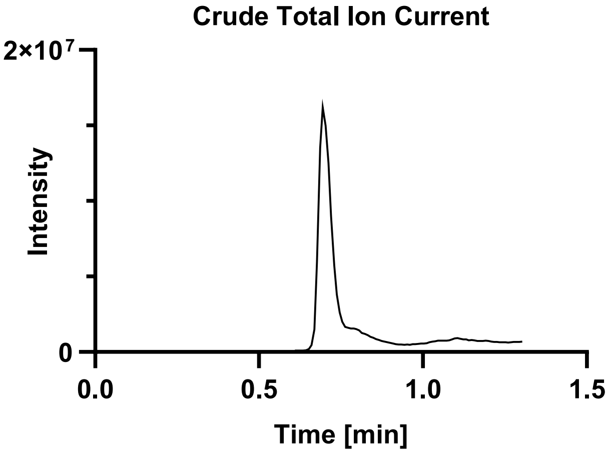

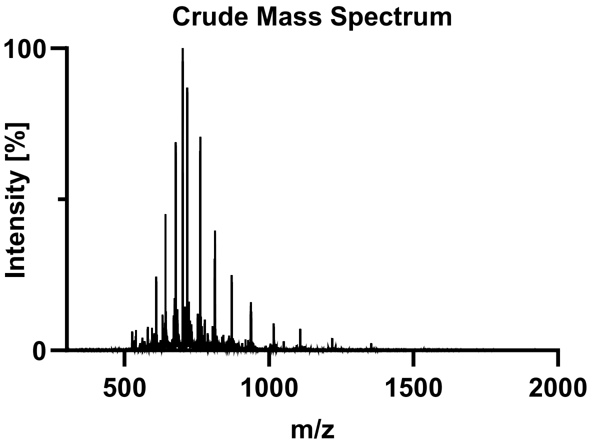

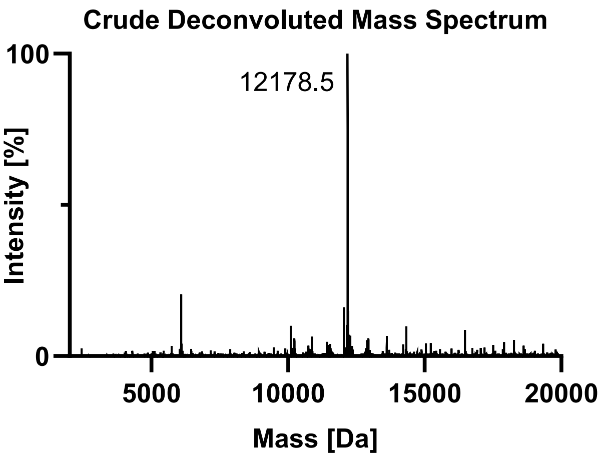

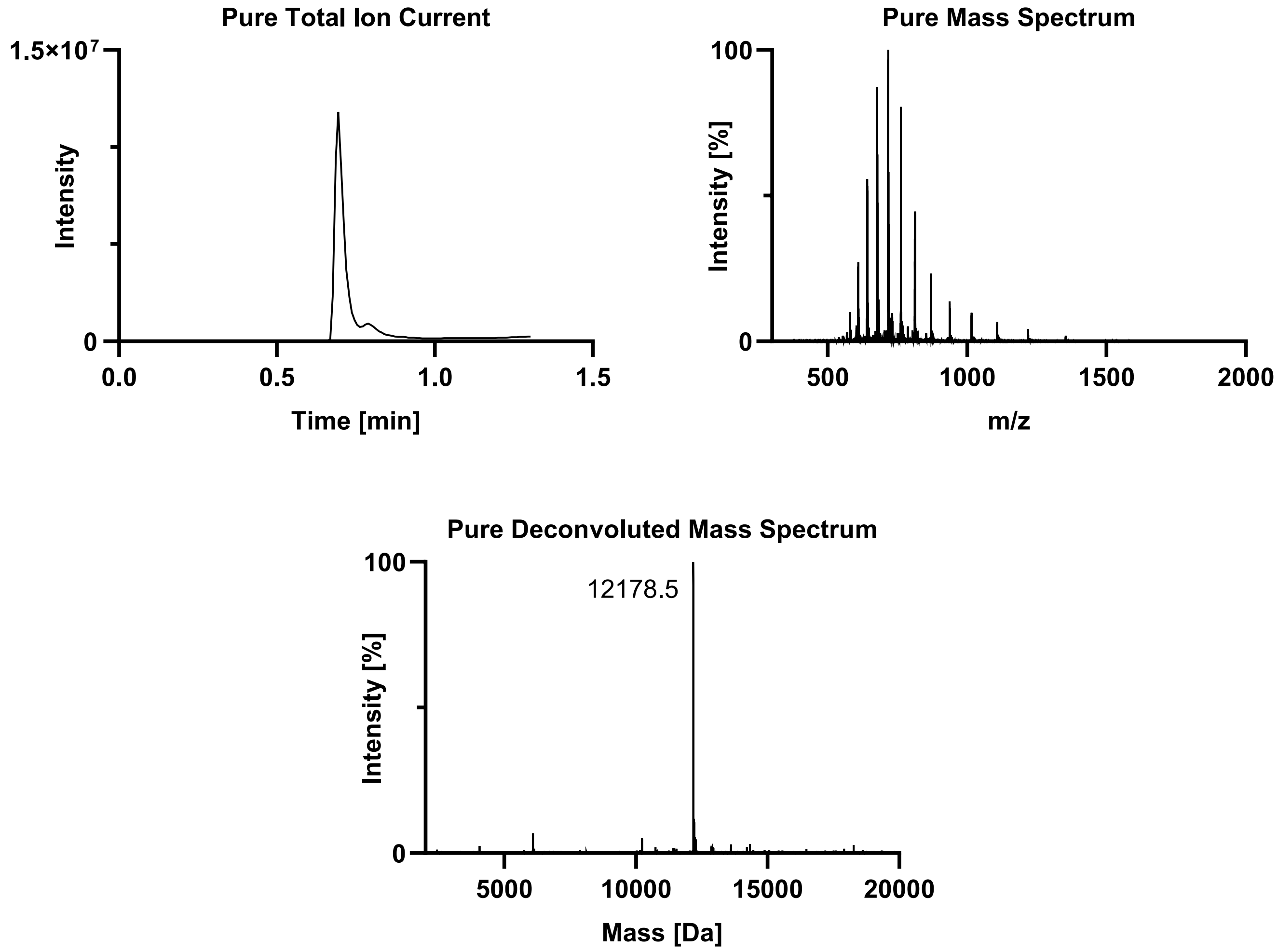
Pure Total Ion Current
1.5×10^7
Intensity
0
0.0
0.5
1.0
1.5
Time [min]
Pure Mass Spectrum
100
Intensity [%]
0
500
1000
1500
2000
m/z
Pure Deconvoluted Mass Spectrum
100
12178.5
Intensity [%]
0
5000
10000
15000
20000
Mass [Da]

#### Tau(297-407) pS400 pS404 (**3j**).

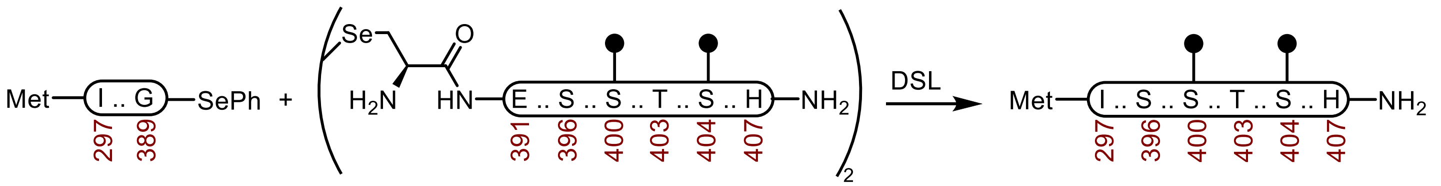

**1** **2j** **3j**

**Tau(297-407) pS400 pS404 (3j).** A 20 mL vial with a stir-bar containing a selenoester **1** (44.1 mg, 4.31 μmol) and diselenide **2j** (10.9 mg, 2.50 μmol) was treated with Ar-purged DSL buffer (1.0 mL, 6 M Gnd·HCl, 0.1 M $Na_2HPO_4$, pH 6.9), and the vial was carefully sealed under a stream of Ar. After the reaction was mixed at 23 °C for 2 h, the DPDS was extracted with Ar-purged hexanes (5 × 1 mL) under a stream of Ar. The reaction was treated with Ar-purged deselenization buffer (2.0 mL, 200 mM TCEP·HCl, 100 mM DTT, 6 M Gnd·HCl, 100 mM HEPES, pH 6.9) and it was mixed at 23 ºC. After 24 h, it was purified by sample displacement mode chromatography (Microsorb C18, 4.6 x 250 mm, 5 μm, 100 Å, 50 ºC, 1 mL/min, 5%-65% MeCN/$H_2O$/0.1% TFA over 60 min), and the fractions were lyophilized to afford **3j** (30.8 mg, 59% yield).

**UPLC-HRMS:** Measured on an ACQUITY UPLC Protein BEH C4 column (2.1 x 50 mm; 1.7 μm; 300 Å; 55 °C; 0.6 mL/min) with a gradient of 5% MeCN/$H_2O$/0.1% FA until 0.1 min, 5-95% MeCN/$H_2O$/0.1% FA until 1.0 min, 95% MeCN/$H_2O$/0.1% FA until 1.1 min.

**Calculated Mass:** $C_{527}H_{864}N_{158}O_{165}P_2S_2$ 12179.7, 12181.2 found.

**Crude UPLC-HRMS:**

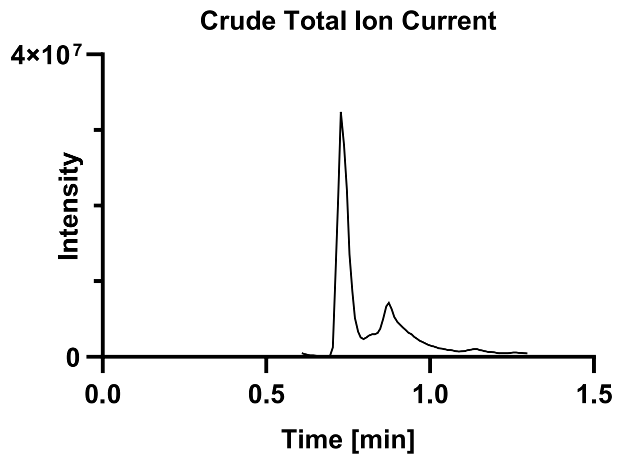

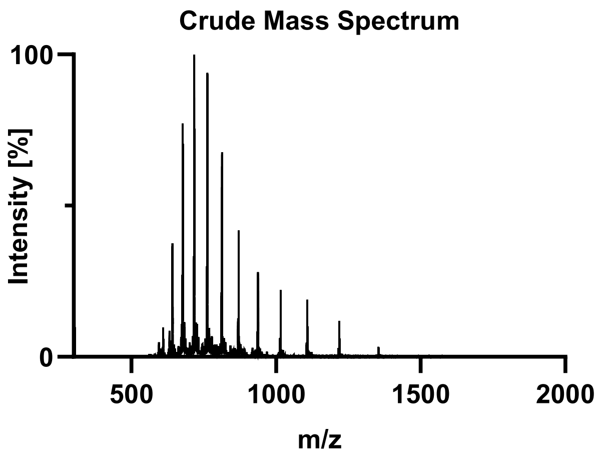

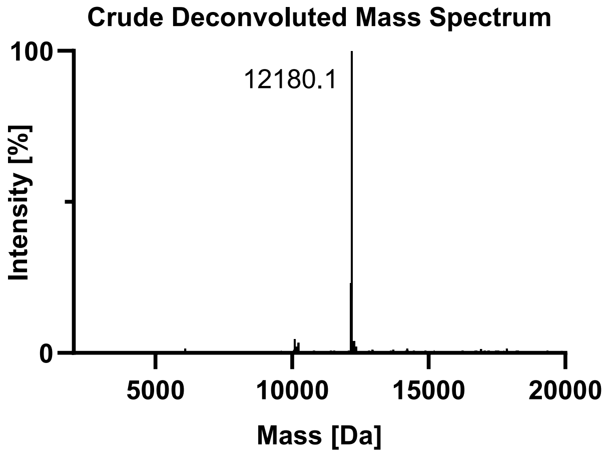

**Pure UPLC-HRMS:**

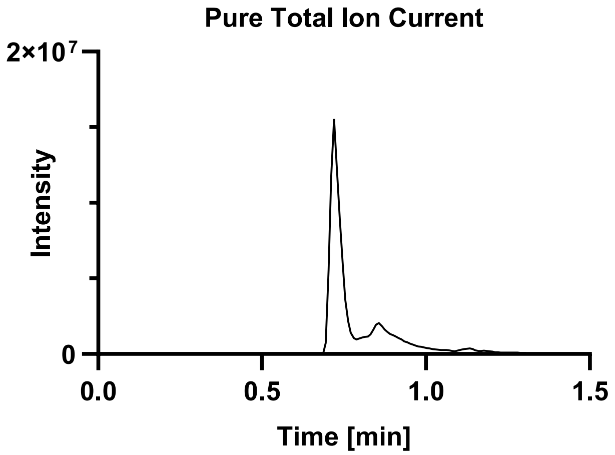

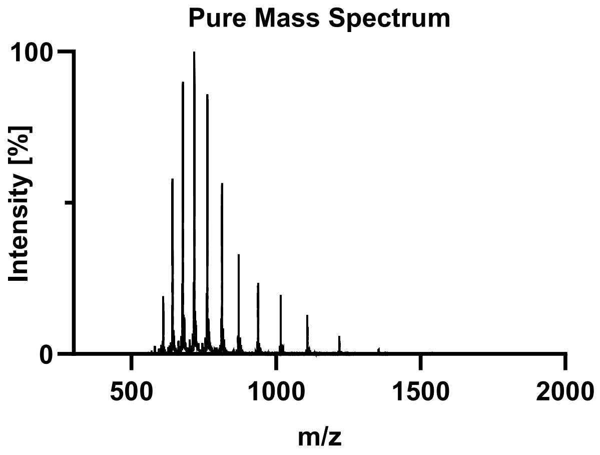

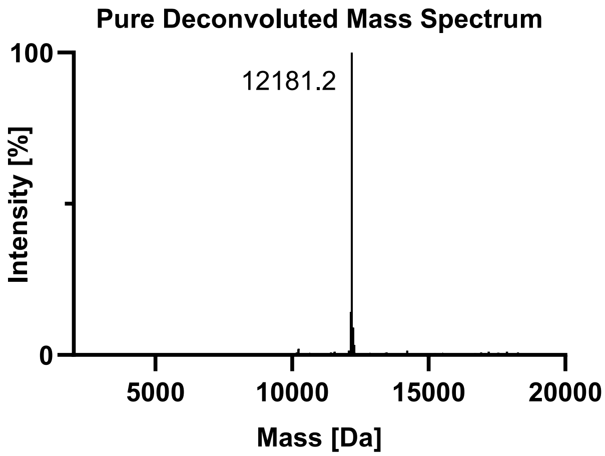

Tau(297-407) pT403 pS404 (**3k**).

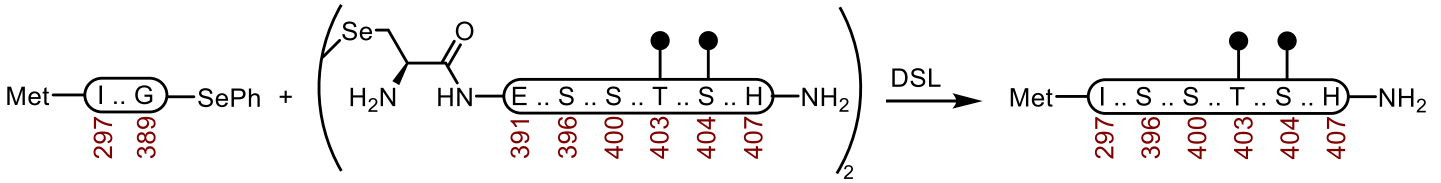

**Tau(297-407) pT403 pS404 (3k).** A 20 mL vial with a stir-bar containing a selenoester **1** (33.3 mg, 3.25 µmol) and diselenide **2k** (8.8 mg, 2.02 µmol) was treated with Ar-purged DSL buffer (0.75 mL, 6 M Gnd·HCl, 0.1 M $Na_2HPO_4$, pH 6.9), and the vial was carefully sealed under a stream of Ar. After the reaction was mixed at 23 °C for 2.5 h, the DPDS was extracted with Ar-purged hexanes (5 × 1 mL) under a stream of Ar. The reaction was treated with Ar-purged deselenization buffer (1.5 mL, 200 mM TCEP·HCl, 100 mM DTT, 6 M Gnd·HCl, 100 mM HEPES, pH 6.8) and it was mixed at 23 ºC. After 24 h, it was purified over a C4 column (10 x 250 mm, 10 µm, 100 Å, 4.7 mL/min) 10%-45% MeCN/$H_2O$/0.1% TFA over 30 min. The material was further purified by sample displacement mode chromatography (Microsorb C18, 4.6 x 250 mm, 5 µm, 100 Å, 50 ºC, 1 mL/min, 5%-65% MeCN/$H_2O$/0.1% TFA over 60 min), and the fractions were lyophilized to afford **3k** (13.6 mg, 34% yield).

**UPLC-HRMS:** Measured on an ACQUITY UPLC Protein BEH C4 column (2.1 x 50 mm; 1.7 µm; 300 Å; 55 °C; 0.6 mL/min) with a gradient of 5% MeCN/$H_2O$/0.1% FA until 0.1 min, 5-95% MeCN/$H_2O$/0.1% FA until 1.0 min, 95% MeCN/$H_2O$/0.1% FA until 1.1 min.

**Calculated Mass:** $C_{527}H_{864}N_{158}O_{165}P_2S_2$ 12179.7, 12179.0 found.

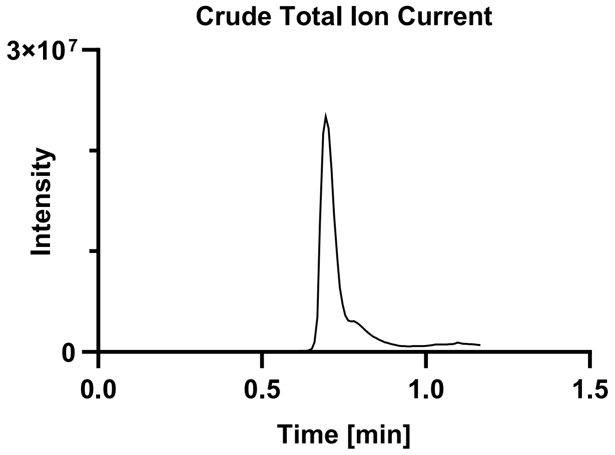

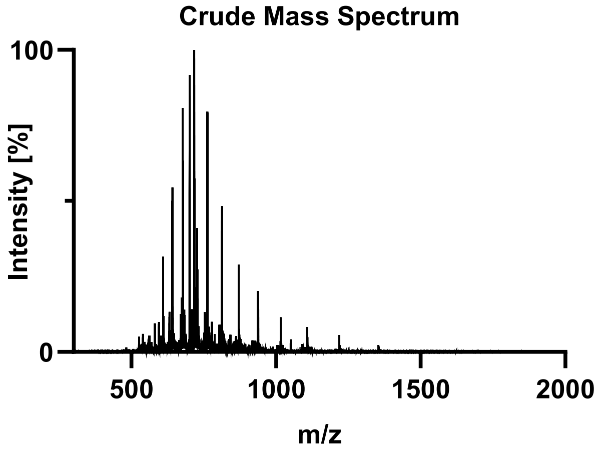

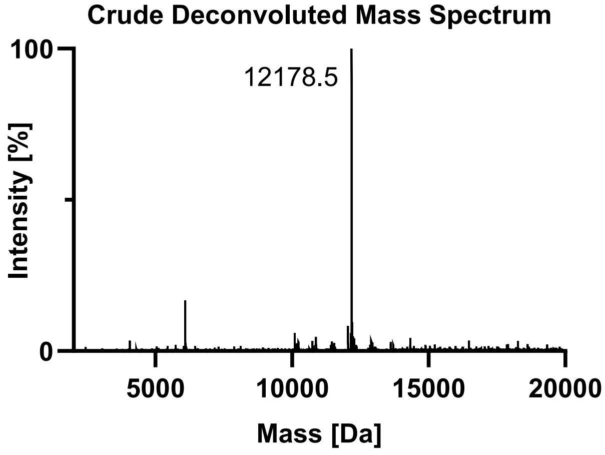

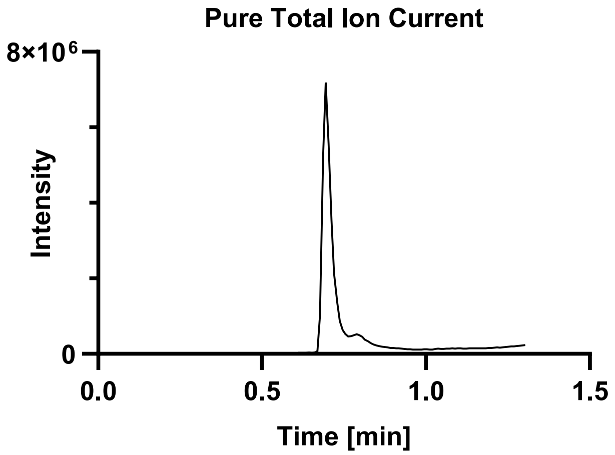
Pure Total Ion Current
8×10^6
0
Intensity
0.0
0.5
1.0
1.5
Time [min]

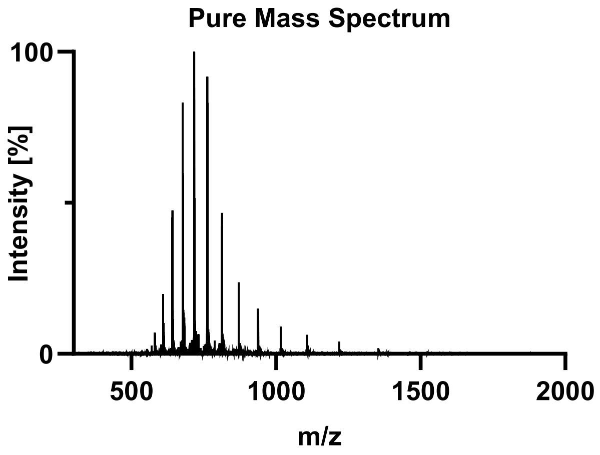
Pure Mass Spectrum
100
0
Intensity [%]
500
1000
1500
2000
m/z

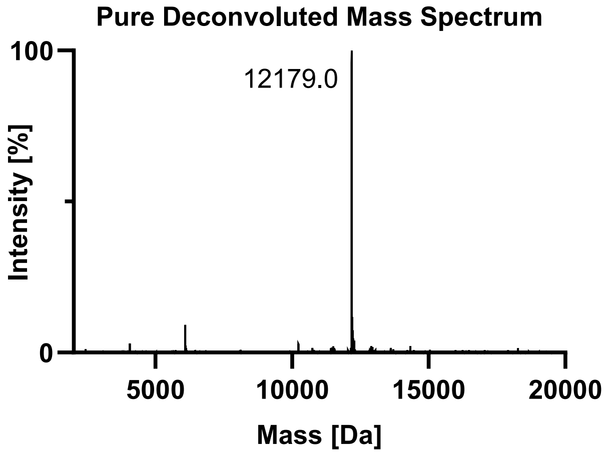
Pure Deconvoluted Mass Spectrum
100
0
12179.0
Intensity [%]
5000
10000
15000
20000
Mass [Da]

Tau(297-407) pS396 pS400 pT403 (**3l**).

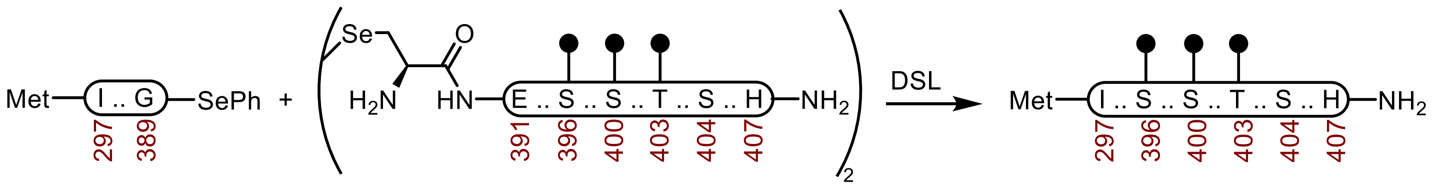

**1** **2l** **3l**

**Tau(297-407) pS396 pS400 pT403 (3l).** A 20 mL vial with a stir-bar containing a selenoester **1** (39.7 mg, 3.88 μmol) and diselenide **2l** (10.5 mg, 2.32 μmol) was treated with Ar-purged DSL buffer (0.75 mL, 6 M Gnd·HCl, 0.1 M $Na_2HPO_4$, pH 6.8), and the vial was carefully sealed under a stream of Ar. After the reaction was mixed at 23 °C for 2 h, the DPDS was extracted with Ar-purged hexanes (5 × 1 mL) under a stream of Ar. The reaction was treated with Ar-purged deselenization buffer (1.5 mL, 200 mM TCEP·HCl, 100 mM DTT, 6 M Gnd·HCl, 100 mM HEPES, pH 7.0) and it was mixed at 23 ºC. After 24 h, it was purified by sample displacement mode chromatography (Microsorb C18, 4.6 x 250 mm, 5 μm, 100 Å, 50 ºC, 1 mL/min, 5%-65% MeCN/$H_2O$/0.1% TFA over 60 min), and the fractions were lyophilized to afford **3l** (26.1 mg, 55% yield).

**UPLC-HRMS:** Measured on an ACQUITY UPLC Protein BEH C4 column (2.1 x 50 mm; 1.7 μm; 300 Å; 55 °C; 0.6 mL/min) with a gradient of 5% MeCN/$H_2O$/0.1% FA until 0.1 min, 5-95% MeCN/$H_2O$/0.1% FA until 1.0 min, 95% MeCN/$H_2O$/0.1% FA until 1.1 min.

**Calculated Mass:** $C_{527}H_{865}N_{158}O_{168}P_3S_2$ 12259.7, 12260.0 found.

**Crude UPLC-HRMS:**

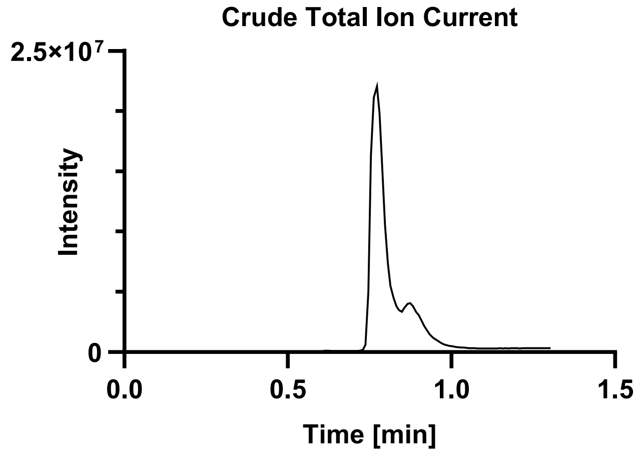

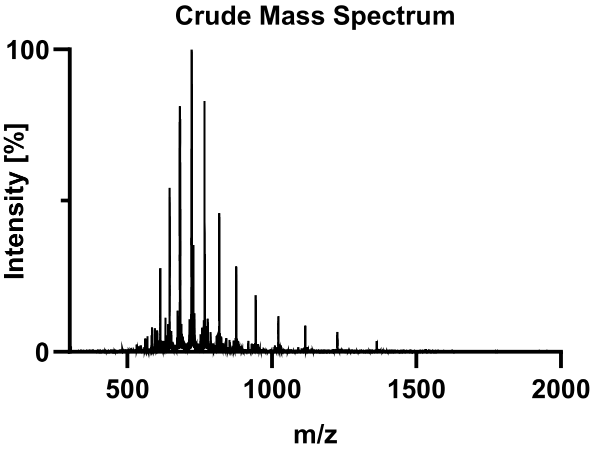

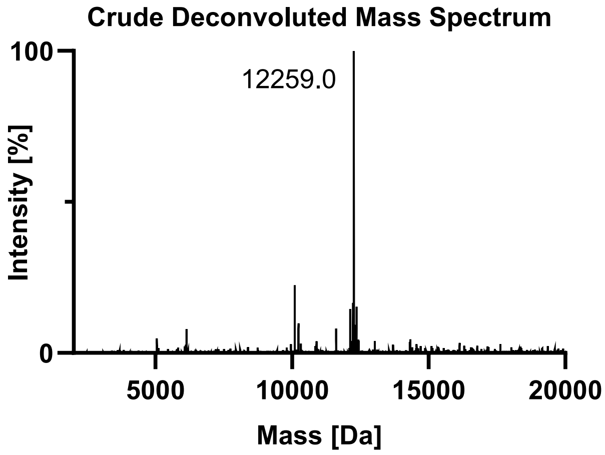

**Pure UPLC-HRMS:**

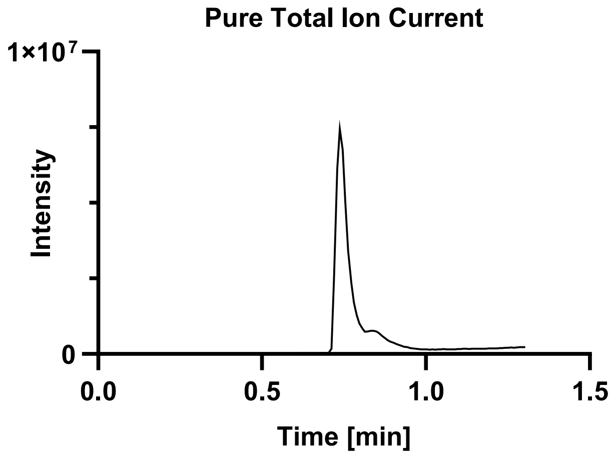

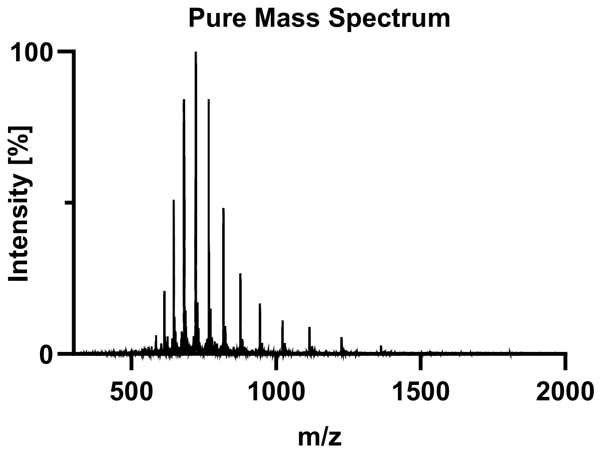

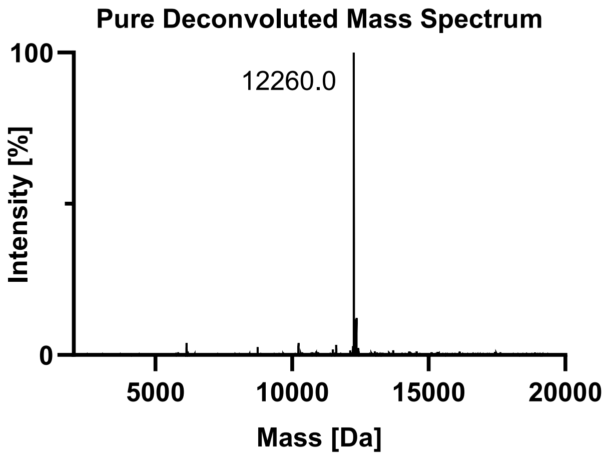

Tau(297-407) pS396 pS400 pS404 (**3m**).

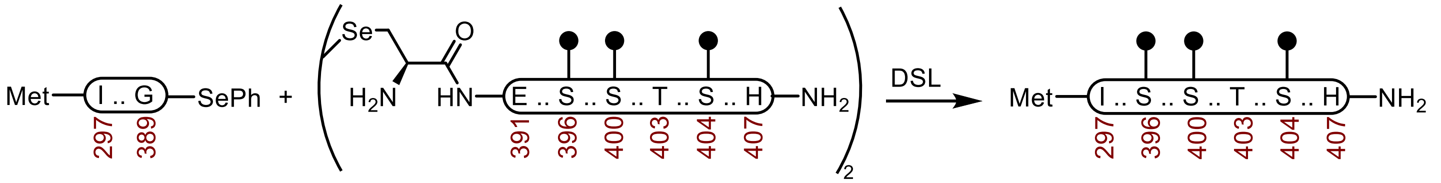

**1** **2m** **3m**

**Tau(297-407) pS396 pS400 pS404 (3m).** A 20 mL vial with a stir-bar containing a selenoester **1** (39.3 mg, 3.84 μmol) and diselenide **2m** (9.5 mg, 2.03 μmol) was treated with Ar-purged DSL buffer (1.0 mL, 6 M Gnd·HCl, 0.1 M $Na_2HPO_4$, pH 6.8), and the vial was carefully sealed under a stream of Ar. After the reaction was mixed at 23 °C for 2 h, the DPDS was extracted with Ar-purged hexanes (5 × 1 mL) under a stream of Ar. The reaction was treated with Ar-purged deselenization buffer (2.0 mL, 200 mM TCEP·HCl, 100 mM DTT, 6 M Gnd·HCl, 100 mM HEPES, pH 6.8) and it was mixed at 23 ºC. After 24 h, it was purified over a C4 column (10 x 250 mm, 10 μm, 100 Å, 4.7 mL/min) 10%-45% MeCN/$H_2O$/0.1% TFA over 30 min. The material was further purified by sample displacement mode chromatography (Microsorb C18, 4.6 x 250 mm, 5 μm, 100 Å, 50 ºC, 1 mL/min, 5%-65% MeCN/$H_2O$/0.1% TFA over 60 min), and the fractions were lyophilized to afford **3m** (16.2 mg, 34% yield).

**UPLC-HRMS:** Measured on an ACQUITY UPLC Protein BEH C4 column (2.1 x 50 mm; 1.7 μm; 300 Å; 55 °C; 0.6 mL/min) with a gradient of 5% MeCN/$H_2O$/0.1% FA until 0.1 min, 5-95% MeCN/$H_2O$/0.1% FA until 1.0 min, 95% MeCN/$H_2O$/0.1% FA until 1.1 min.

**Calculated Mass:** $C_{527}H_{865}N_{158}O_{168}P_3S_2$ 12259.7, 12259.0 found.

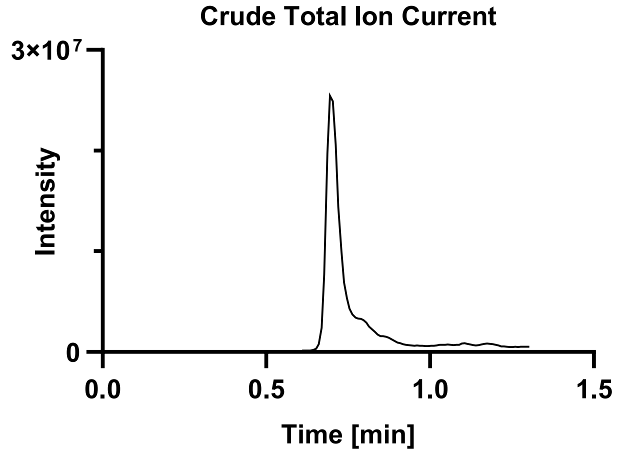

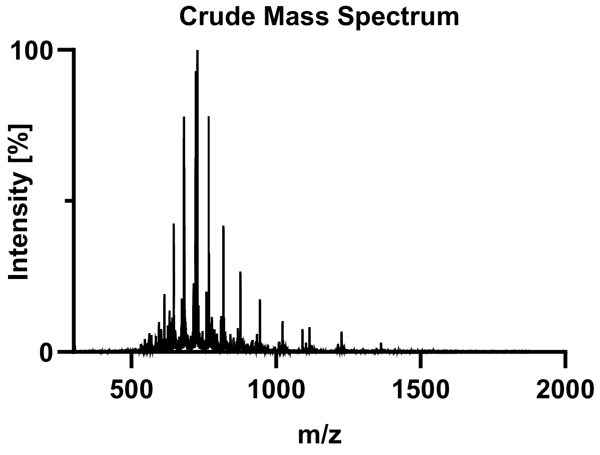

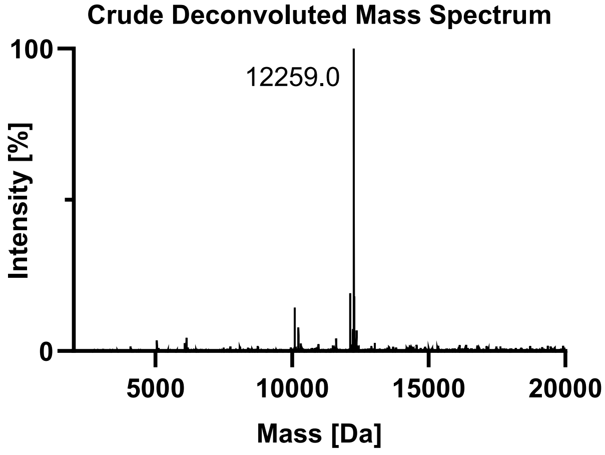

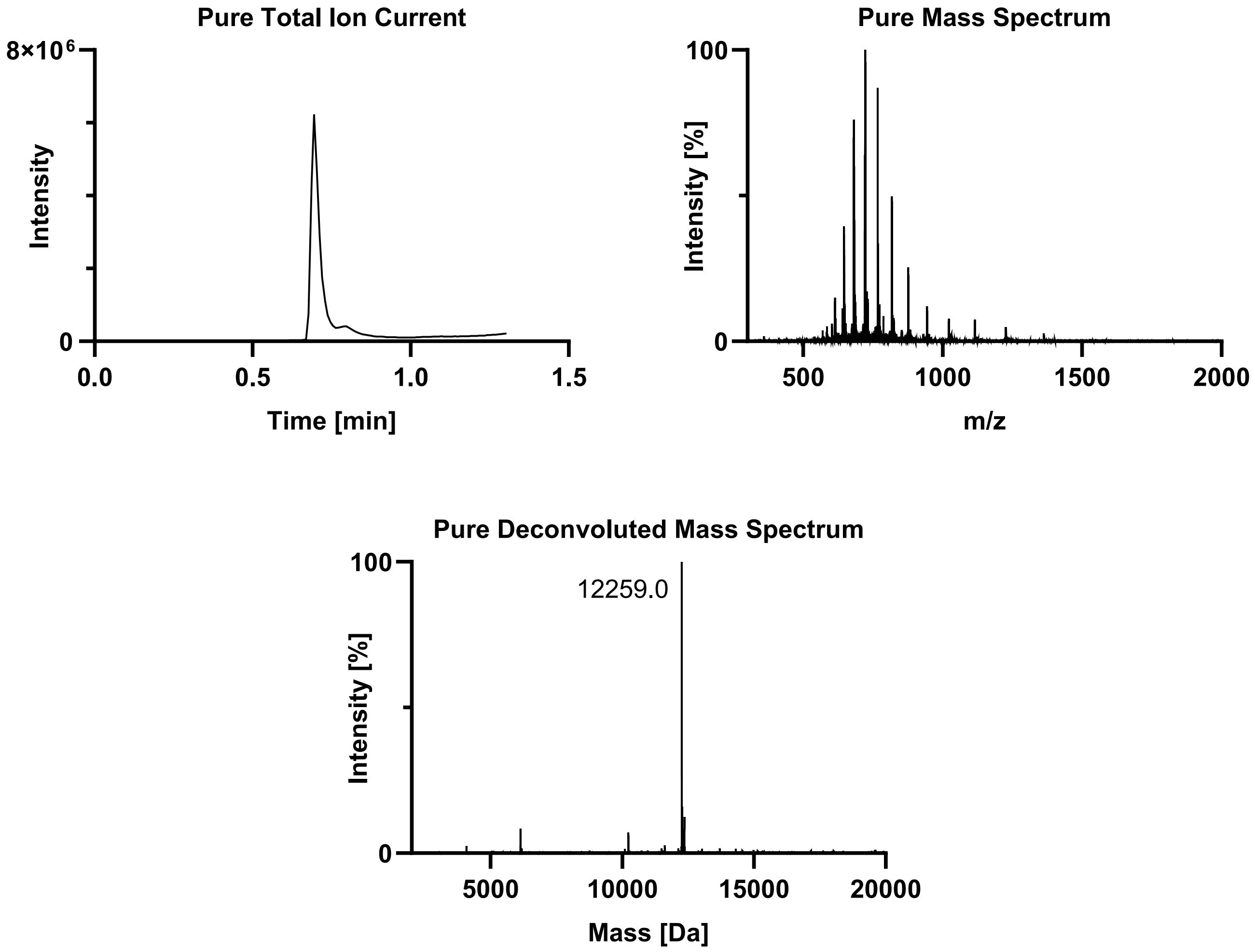
Pure Total Ion Current
8×10^6
0
Intensity
0.0
0.5
1.0
1.5
Time [min]
Pure Mass Spectrum
100
0
Intensity [%]
500
1000
1500
2000
m/z
Pure Deconvoluted Mass Spectrum
100
0
Intensity [%]
12259.0
5000
10000
15000
20000
Mass [Da]

Tau(297-407) pS396 pT403 pS404 (**3n**).

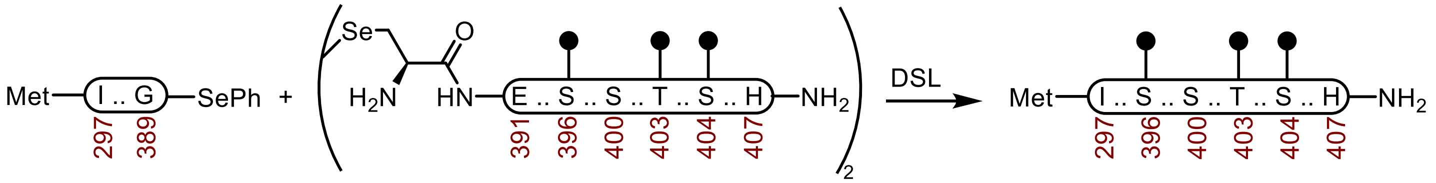

**1** **2n** **3n**

**Tau(297-407) pS396 pT403 pS404 (3n).** A 1-dram vial with a stir-bar containing a selenoester **1** (39.9 mg, 3.90 µmol) and diselenide **2n** (12.1 mg, 2.68 µmol) was treated with Ar-purged DSL buffer (1.0 mL, 6 M Gnd·HCl, 0.1 M $Na_2HPO_4$, pH 6.8), and the vial was carefully sealed under a stream of Ar. After the reaction was mixed at 23 °C for 2 h, the DPDS was extracted with Ar-purged hexanes (5 × 1 mL) under a stream of Ar. The reaction was treated with Ar-purged deselenization buffer (2.0 mL, 200 mM TCEP·HCl, 100 mM DTT, 6 M Gnd·HCl, 100 mM HEPES, pH 7.0) and it was mixed at 23 ºC. After 24 h, it was purified by sample displacement mode chromatography (Microsorb C18, 4.6 x 250 mm, 5 µm, 100 Å, 50 ºC, 1 mL/min, 5%-65% MeCN/$H_2O$/0.1% TFA over 60 min), and the fractions were lyophilized to afford **3n** (25.2 mg, 53% yield).

**UPLC-HRMS:** Measured on an ACQUITY UPLC Protein BEH C4 column (2.1 x 50 mm; 1.7 µm; 300 Å; 55 °C; 0.6 mL/min) with a gradient of 5% MeCN/$H_2O$/0.1% FA until 0.1 min, 5-95% MeCN/$H_2O$/0.1% FA until 1.0 min, 95% MeCN/$H_2O$/0.1% FA until 1.1 min.

**Calculated Mass:** $C_{527}H_{865}N_{158}O_{168}P_3S_2$ 12259.7, 12260.3 found.

**Crude UPLC-HRMS:**

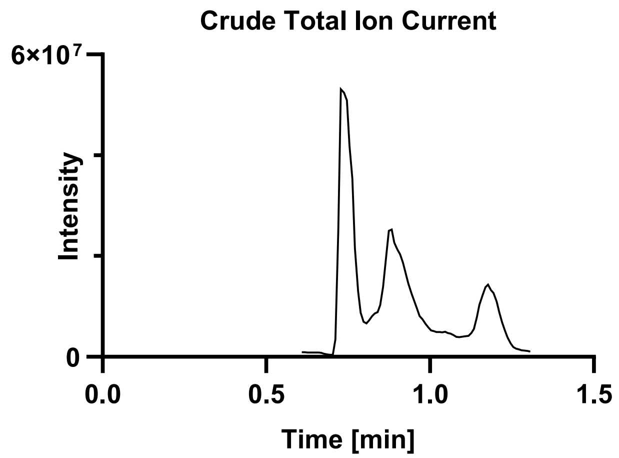

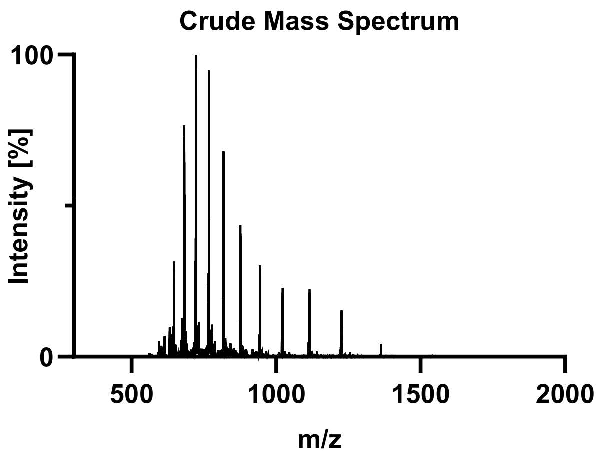

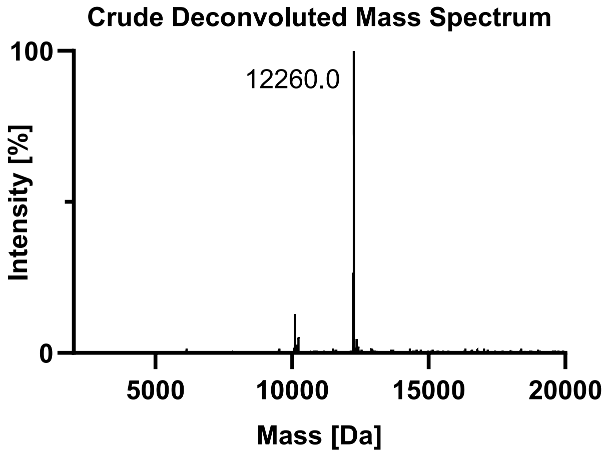

**Pure UPLC-HRMS:**

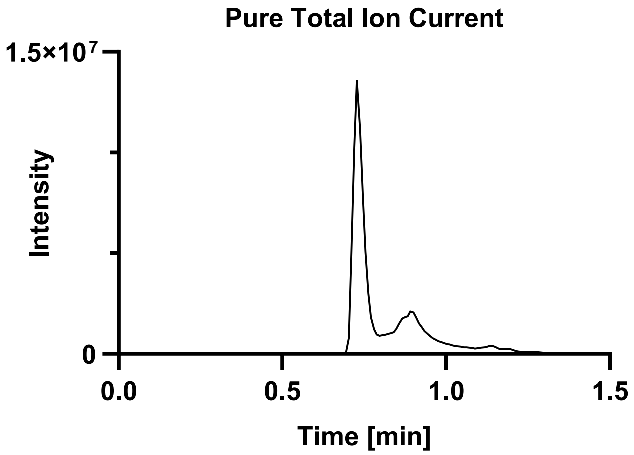

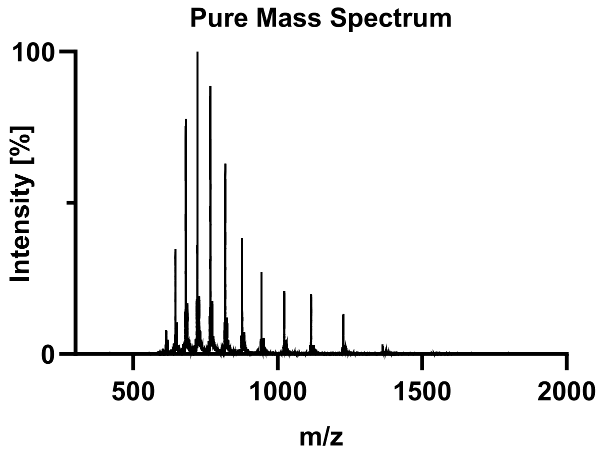

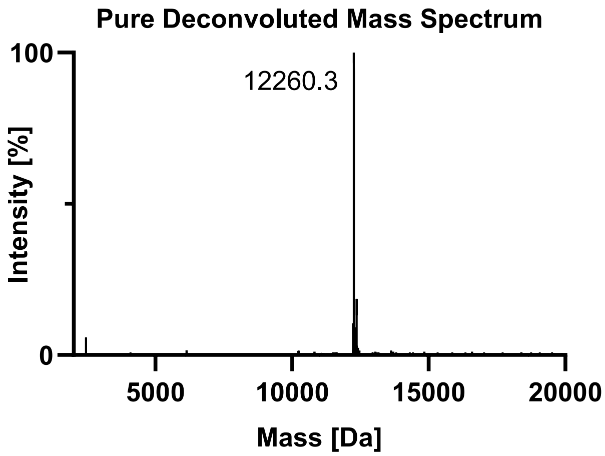

Tau(297-407) pS400 pT403 pS404 (**3o**).

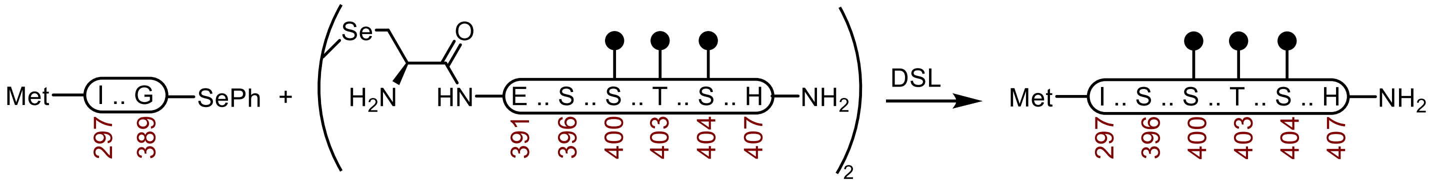

**1** **2o** **3o**

**Tau(297-407) pS400 pT403 pS404 (3o).** A 20 mL vial with a stir-bar containing a selenoester **1** (41.1 mg, 4.02 μmol) and diselenide **2o** (12.1 mg, 2.68 μmol) was treated with Ar-purged DSL buffer (0.75 mL, 6 M Gnd·HCl, 0.1 M $Na_2HPO_4$, pH 6.8), and the vial was carefully sealed under a stream of Ar. After the reaction was mixed at 23 °C for 2 h, the DPDS was extracted with Ar-purged hexanes (5 × 1 mL) under a stream of Ar. The reaction was treated with Ar-purged deselenization buffer (1.5 mL, 200 mM TCEP·HCl, 100 mM DTT, 6 M Gnd·HCl, 100 mM HEPES, pH 7.0) and it was mixed at 23 ºC. After 24 h, it was purified by sample displacement mode chromatography (Microsorb C18, 4.6 x 250 mm, 5 μm, 100 Å, 50 ºC, 1 mL/min, 5%-65% MeCN/$H_2O$/0.1% TFA over 60 min), and the fractions were lyophilized to afford **3o** (20.3 mg, 41% yield).

**UPLC-HRMS:** Measured on an ACQUITY UPLC Protein BEH C4 column (2.1 x 50 mm; 1.7 μm; 300 Å; 55 °C; 0.6 mL/min) with a gradient of 5% MeCN/$H_2O$/0.1% FA until 0.1 min, 5-95% MeCN/$H_2O$/0.1% FA until 1.0 min, 95% MeCN/$H_2O$/0.1% FA until 1.1 min.

**Calculated Mass:** $C_{527}H_{865}N_{158}O_{168}P_3S_2$ 12259.7, 12259.0 found.

**Crude UPLC-HRMS:**

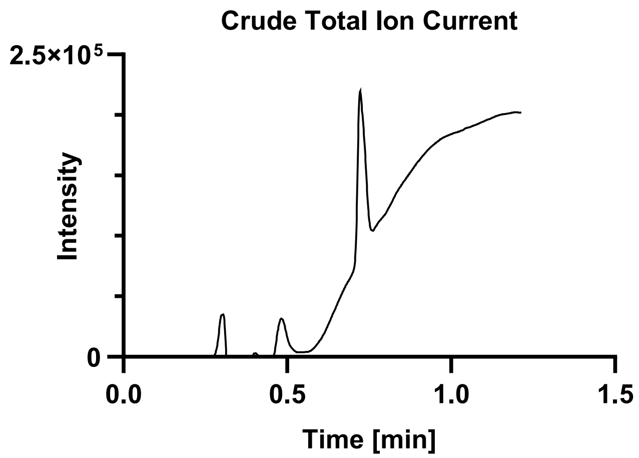

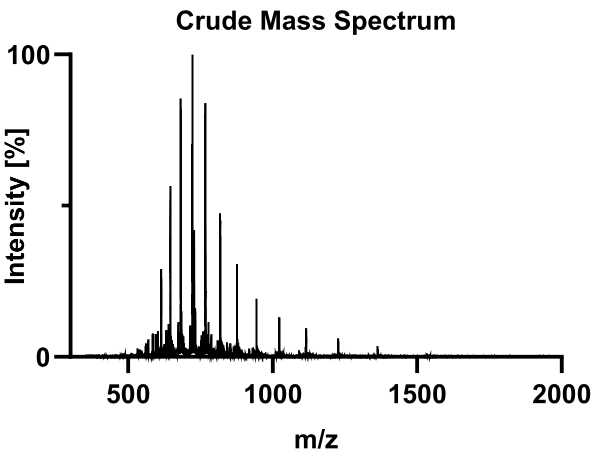

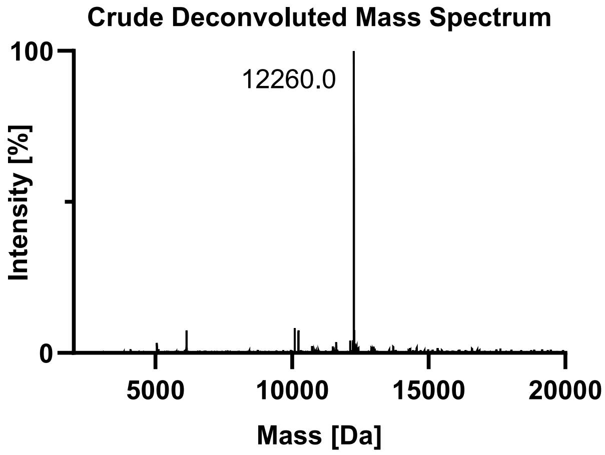

**Pure UPLC-HRMS:**

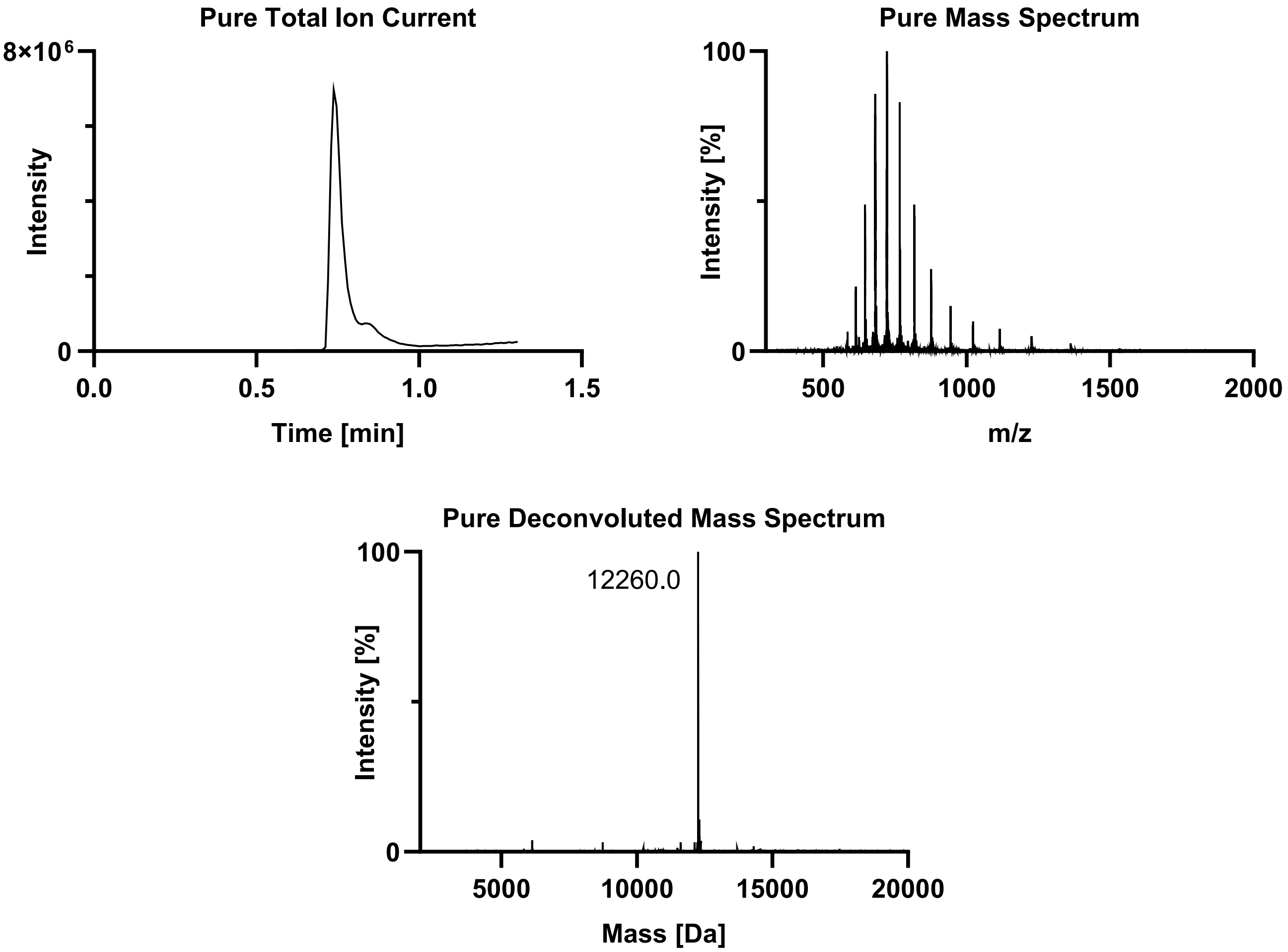

### S4. PRIMARY NUCLEATION KINETICS.

**Assembly conditions.** 200 µM Tau, 10 mM KPB-KOH pH 6.5, 200 mM potassium citrate tribasic, 10 mM DTT, 2 µM ThT, 37 ºC, 7 d.

**Stock solutions.**

- ~1000 µM Tau(297-407) proteoforms in $H_2O$ ($\varepsilon_{280}$ = 2980 $M^{-1}cm^{-1}$).
- 100 mM KPB-KOH pH 6.5.
- 100 mM DTT in $H_2O$.
- 200 µM ThT in $H_2O$.
- 1 M potassium citrate tribasic in $H_2O$.

**Assembly method.** The assembly reactions were performed in black, non-treated 96-well microplates (Corning, 3915), sealed with adhesive microplate film (VWR, 7659). Each well was directly filled with 110 µL of reaction mixture, and the plate was incubated at 37 ºC in the microplate reader without shaking. The final pH was about 7.5 after mixing the buffer components. The reactions were performed in triplicate, with 6 technical replicates per reaction. The wells that evaporated were removed from the analysis. The final pH was about 7.5.

**Microplate settings.** The experiment was performed in a Molecular Devices M5 plate reader. The ThT fluorescence was measured every 10 minutes (Ex = 440 nm, EM = 480 nm, PMT gain = medium, 3 flashes per read).

**Data processing.** The assembly kinetics are fitted to a Boltzmann sigmoidal function to obtain $t_{1/2}$ values with their standard errors. The data points for each condition are shown as circles, and the error is represented as the standard deviation. The Boltzmann fitting is overlayed as a solid black line.

**Primary nucleation ThT traces.**

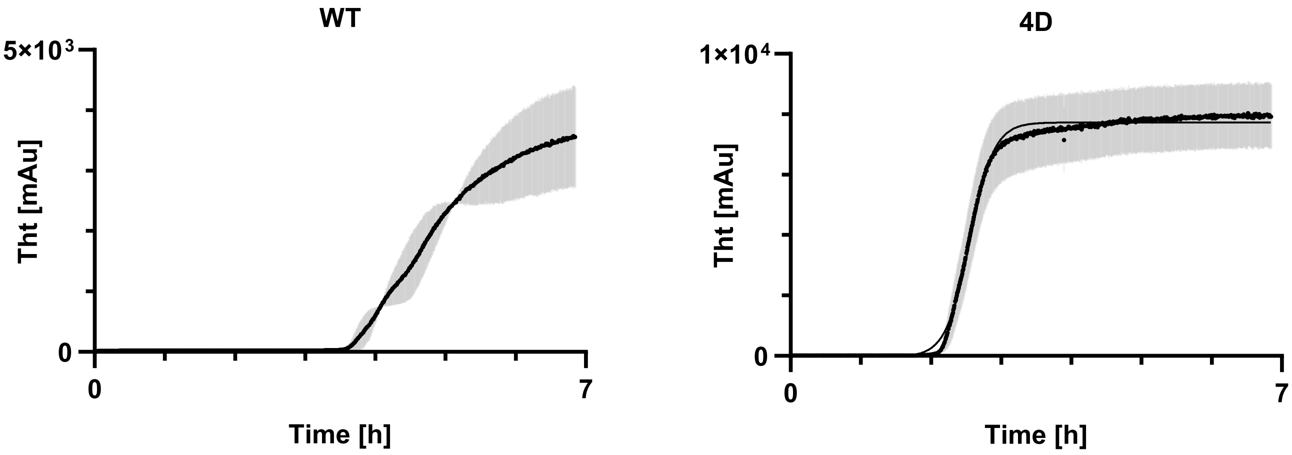

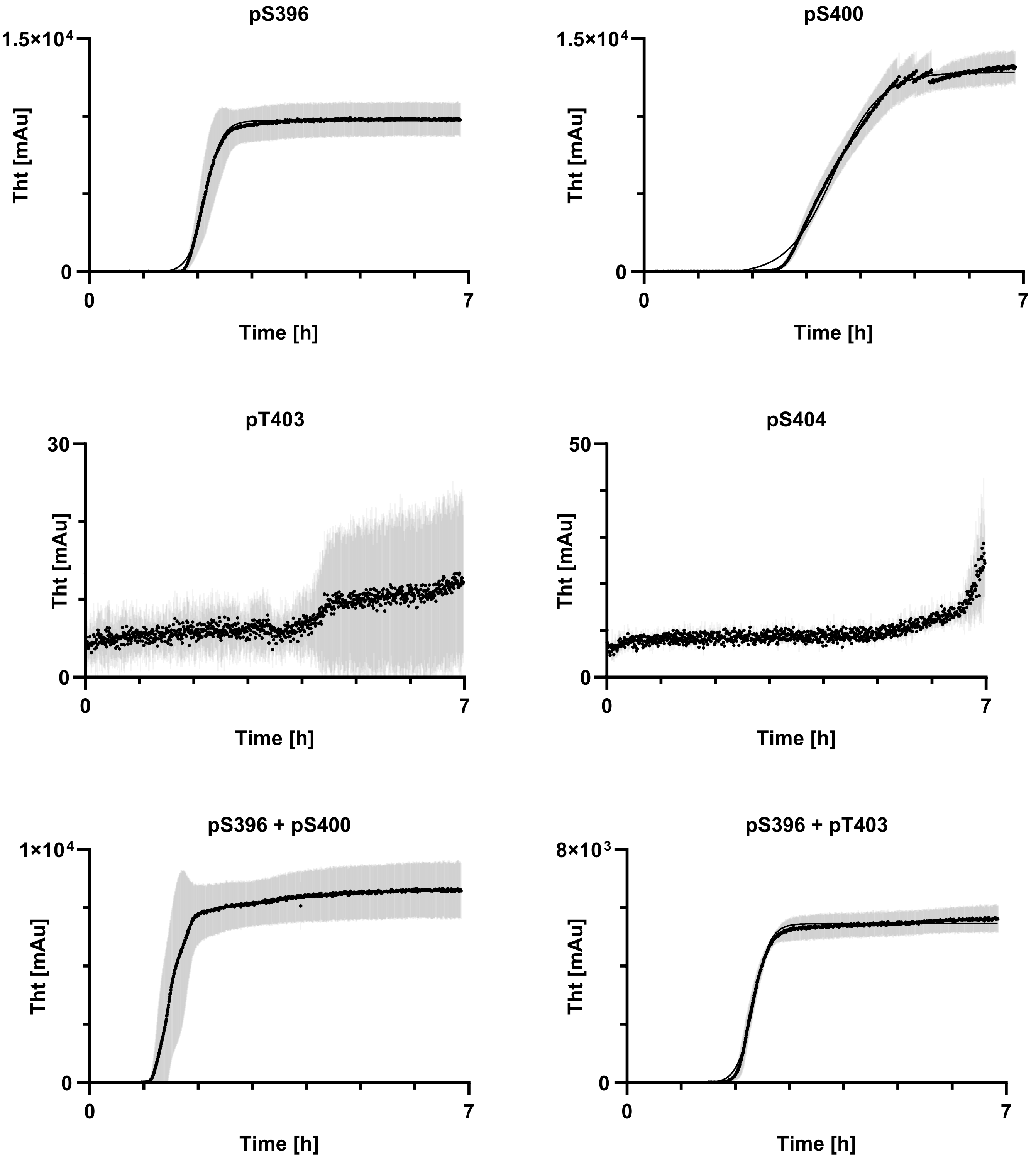
pS396
1.5×10⁴
0
Tht [mAu]
0
7
Time [h]
pS400
1.5×10⁴
0
Tht [mAu]
0
7
Time [h]
pT403
30
0
Tht [mAu]
0
7
Time [h]
pS404
50
0
Tht [mAu]
0
7
Time [h]
pS396 + pS400
1×10⁴
0
Tht [mAu]
0
7
Time [h]
pS396 + pT403
8×10³
0
Tht [mAu]
0
7
Time [h]

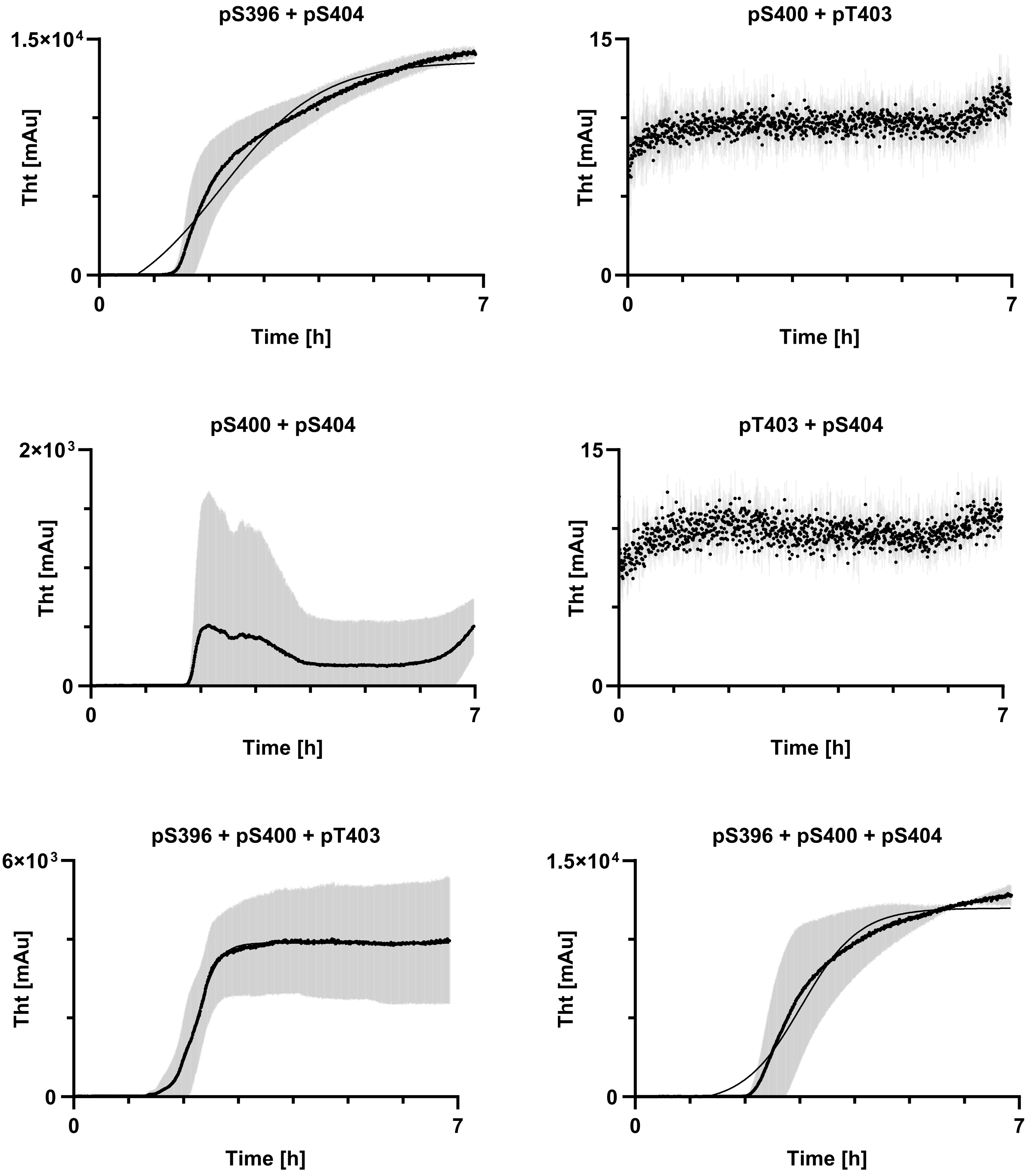
pS396 + pS404
1.5×10⁴
0
Tht [mAu]
0
7
Time [h]
pS400 + pT403
15
0
Tht [mAu]
0
7
Time [h]
pS400 + pS404
2×10³
0
Tht [mAu]
0
7
Time [h]
pT403 + pS404
15
0
Tht [mAu]
0
7
Time [h]
pS396 + pS400 + pT403
6×10³
0
Tht [mAu]
0
7
Time [h]
pS396 + pS400 + pS404
1.5×10⁴
0
Tht [mAu]
0
7
Time [h]

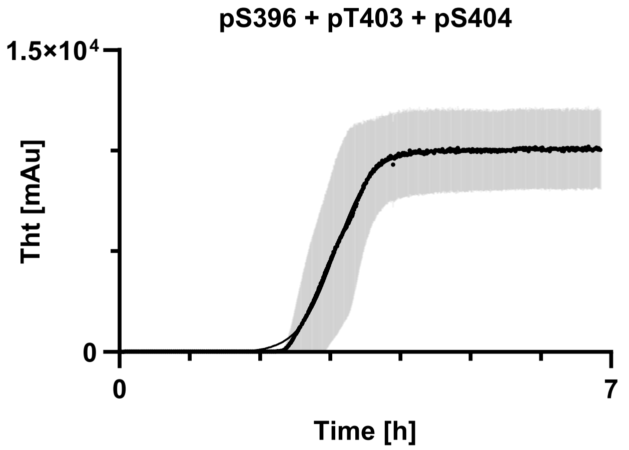

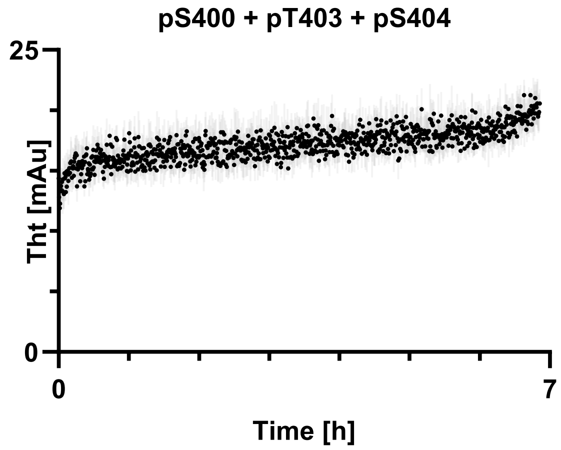

**Table of $t_{1/2}$ values and standard error after primary nucleation.**

| tau proteoform | $t_{1/2}$ [d] | error [d] |
|---|---|---|
| WT | 4.698 | 0.011 |
| 4D | 2.558 | 0.004 |
| pS396 | 2.144 | 0.004 |
| pS400 | 3.508 | 0.005 |
| pS396 + pS400 | 1.534 | 0.007 |
| pS396 + pT403 | 2.339 | 0.003 |
| pS396 + pS404 | 2.222 | 0.026 |
| pS396 + pS400 + pT403 | 2.242 | 0.012 |
| pS396 + pS400 + pS404 | 3.030 | 0.016 |
| pS396 + pT403 + pS404 | 3.074 | 0.012 |

### S5. SEEDED ASSEMBLY KINETICS.

**Assembly conditions.** 50 μM Tau, 5 μM Tau(297-407)-4D seeds, 10 mM KPB-KOH pH 6.5, 200 mM potassium citrate tribasic, 10 mM DTT, 3 μM ThT, 37 °C, 4 d.

**Stock solutions.**

- ~1000 μM Tau(297-407) proteoforms in $H_2O$ ($\varepsilon_{280}$ = 2980 $M^{-1}cm^{-1}$).
- 200 μM Tau(297-407)-4D fibrils (quiescent) in 10 mM KPB-KOH pH 6.5, 200 mM potassium citrate, 10 mM DTT, 1 μM ThT.
- 100 mM KPB-KOH pH 6.5.
- 100 mM DTT in $H_2O$.
- 200 μM ThT in $H_2O$.
- 1 M potassium citrate tribasic in $H_2O$.

**Seeded quiescent assembly method, with 5 μM seeds.** The assembly reactions were performed in black, non-binding, low-volume 384-well microplates (Greiner 784900), sealed with adhesive microplate film (VWR 7659). The plate was incubated at 37 °C in the microplate reader. Reads were conducted every 10 minutes, without shaking. The reactions were performed in triplicate. A 1.6 mL Eppendorf tube was filled with 70 μL of master mix, and 20 μL was pipetted into three wells.

**Microplate settings.** The experiment was performed in a Molecular Devices M5 plate reader. The ThT fluorescence was measured every 10 minutes (Ex = 440 nm, EM = 480 nm, PMT gain = medium, 3 flashes per read). The kinetic traces are reported as averages, and the error bars represent standard deviations. The $t_{1/2}$ values were manually determined from the kinetic traces, averaged, and the error bars represent the standard deviation.

**ThT traces after seeding with Tau(297-407)-4D seeds.**

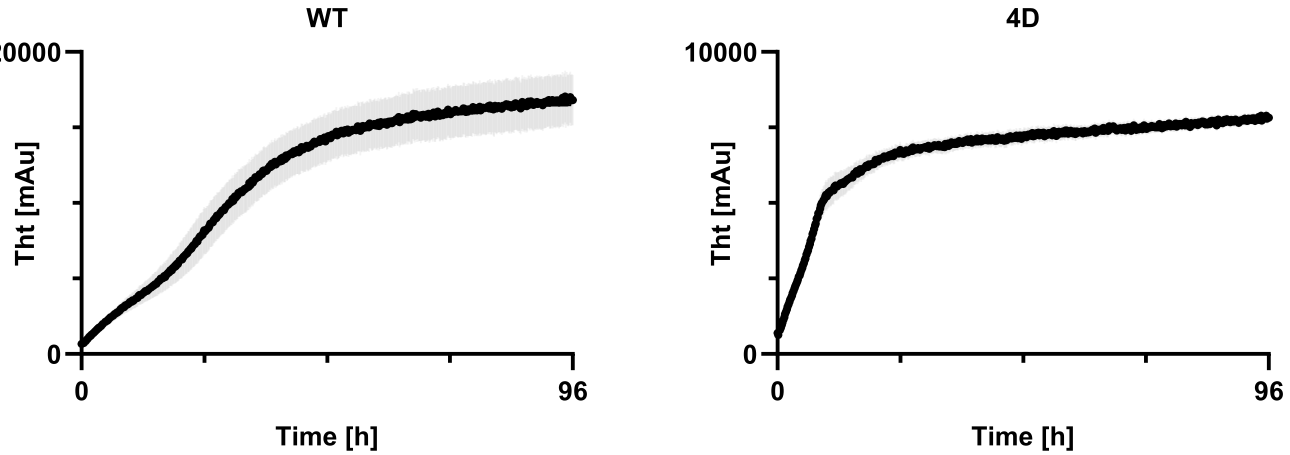

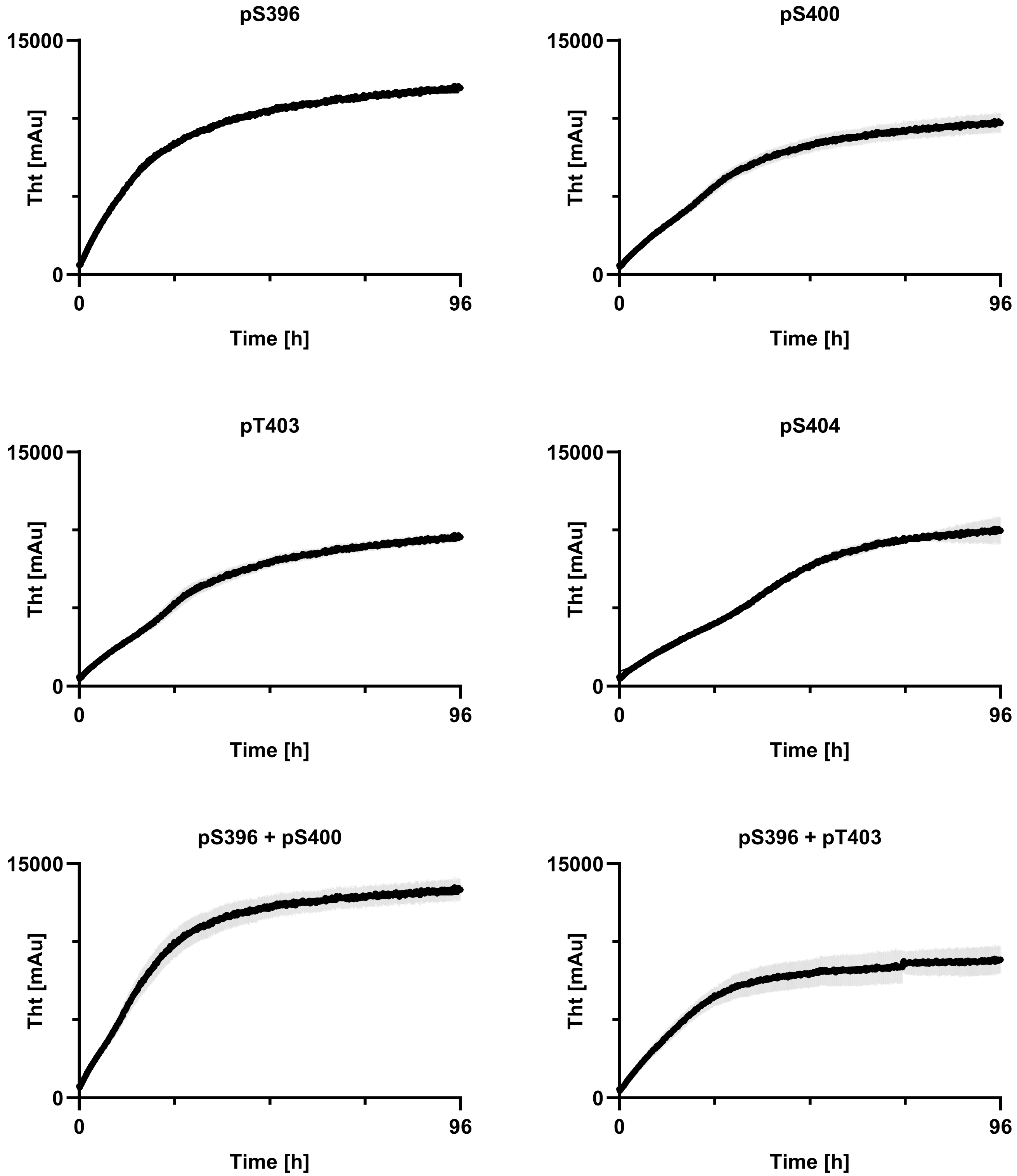
pS396
pS400
pT403
pS404
pS396 + pS400
pS396 + pT403
15000
0
Tht [mAu]
0
96
Time [h]

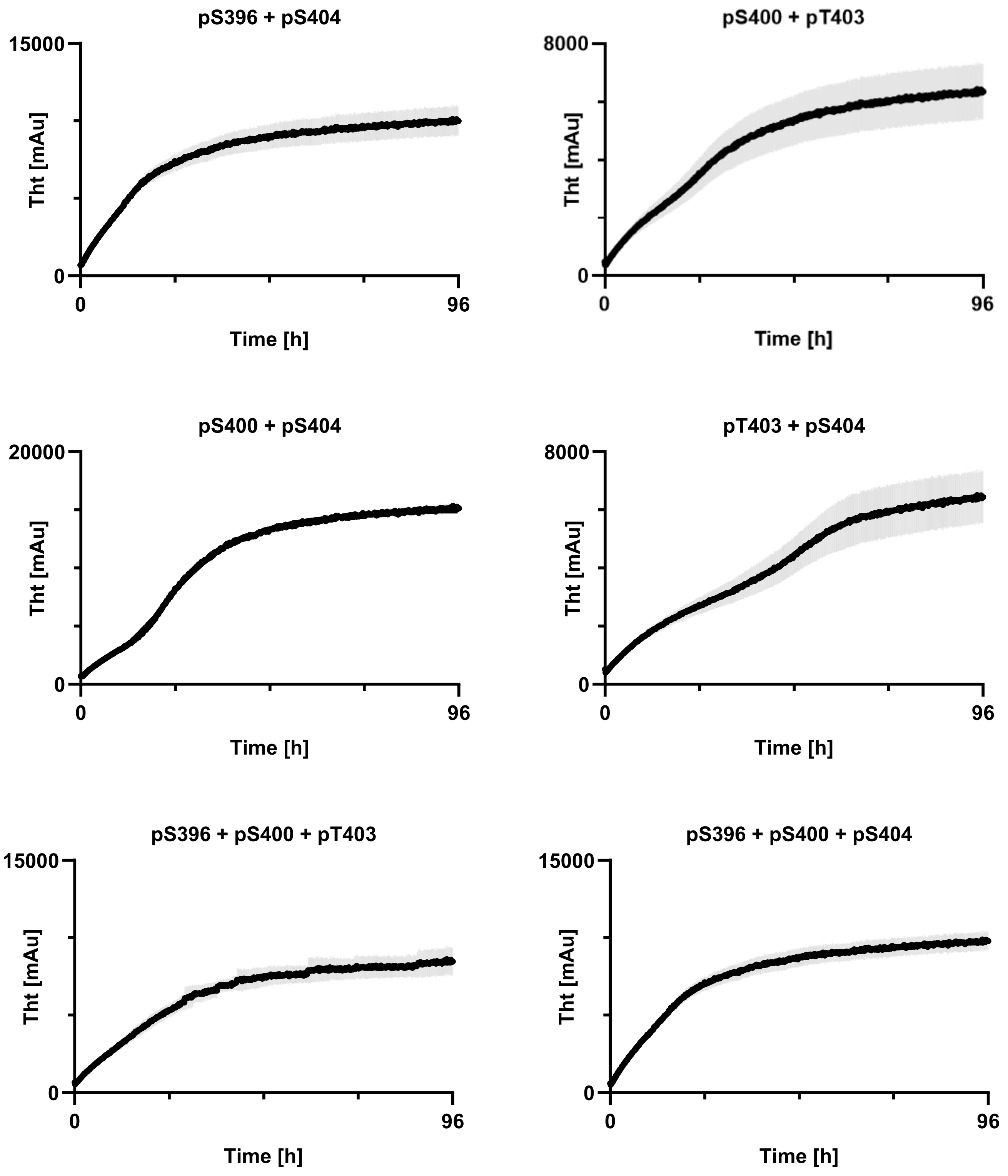
pS396 + pS404
15000
0
Tht [mAu]
0
96
Time [h]
pS400 + pT403
8000
0
Tht [mAu]
0
96
Time [h]
pS400 + pS404
20000
0
Tht [mAu]
0
96
Time [h]
pT403 + pS404
8000
0
Tht [mAu]
0
96
Time [h]
pS396 + pS400 + pT403
15000
0
Tht [mAu]
0
96
Time [h]
pS396 + pS400 + pS404
15000
0
Tht [mAu]
0
96
Time [h]

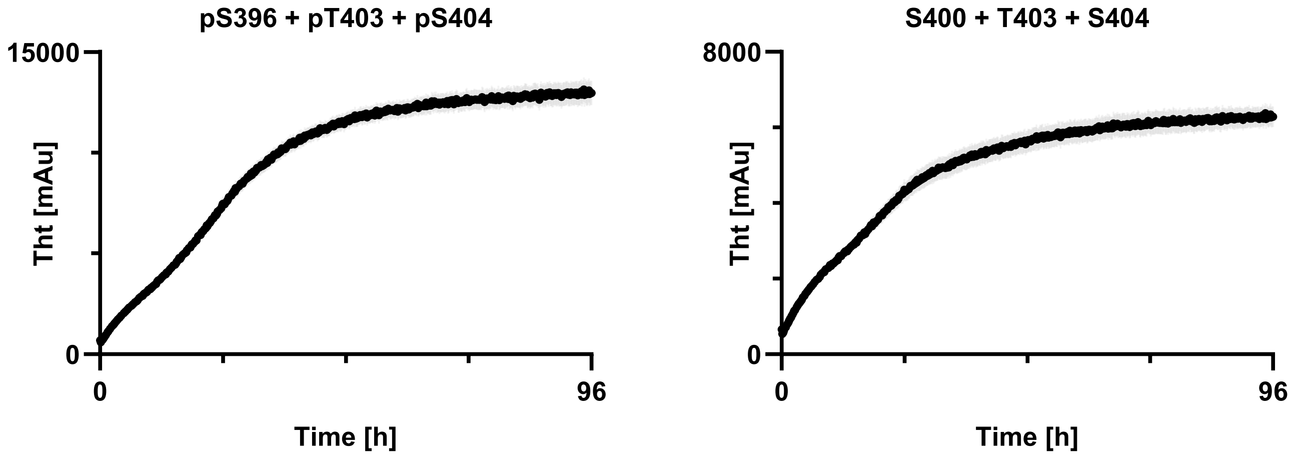

Graphical representation of $t_{1/2}$ values.

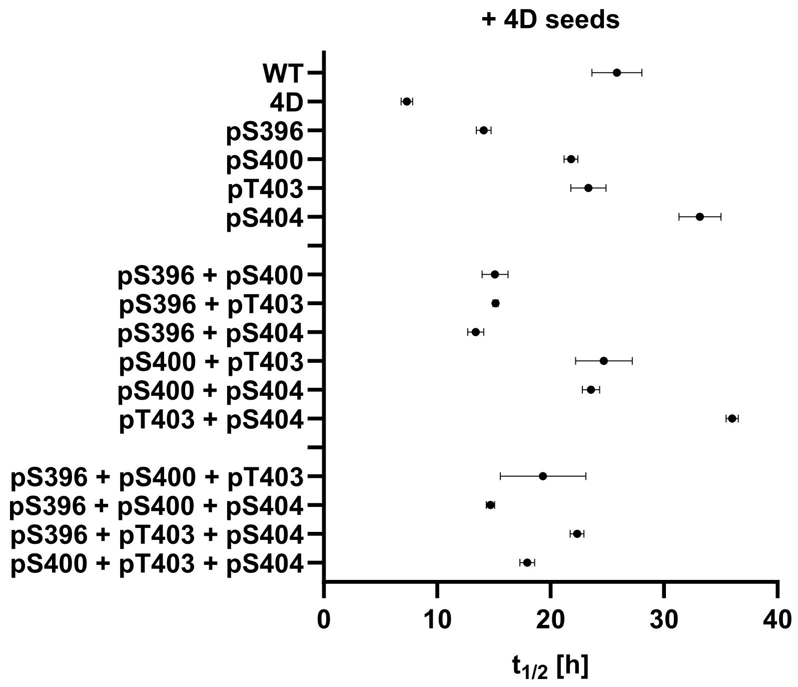

**Table of $t_{1/2}$ values and standard deviation after seeding with Tau(297-407)-4D seeds.**

| tau proteoform | 1 [h] | 2 [h] | 3 [h] | average [h] | STDEV [h] |
|---|---|---|---|---|---|
| WT | 23.7 | 28.1 | 25.7 | 25.8 | 2.2 |
| 4D | 6.9 | 7.2 | 7.9 | 7.3 | 0.5 |
| pS396 | 14.2 | 13.4 | 14.7 | 14.1 | 0.7 |
| pS400 | 22.2 | 21.1 | 22.1 | 21.8 | 0.6 |
| pT403 | 22.7 | 22.2 | 25.1 | 23.3 | 1.6 |
| pS404 | 33.7 | 31.1 | 34.7 | 33.2 | 1.9 |
| pS396 + pS400 | 14.2 | 14.7 | 16.4 | 15.1 | 1.2 |
| pS396 + pT403 | 15.2 | 14.9 | 15.3 | 15.1 | 0.2 |
| pS396 + pS404 | 14.2 | 13.1 | 12.9 | 13.4 | 0.7 |
| pS400 + pT403 | 25.5 | 21.9 | 26.7 | 24.7 | 2.5 |
| pS400 + pS404 | 22.9 | 23.4 | 24.4 | 23.6 | 0.8 |
| pT403 + pS404 | 36.4 | 36.2 | 35.4 | 36.0 | 0.5 |
| pS396 + pS400 + pT403 | 15.4 | 19.7 | 22.9 | 19.3 | 3.8 |
| pS396 + pS400 + pS404 | 14.6 | 15.1 | 14.4 | 14.7 | 0.4 |
| pS396 + pT403 + pS404 | 21.7 | 22.9 | 22.4 | 22.3 | 0.6 |
| pS400 + pT403 + pS404 | 17.2 | 18.2 | 18.4 | 17.9 | 0.6 |

ThT traces after seeding with Tau(297-407) pS396 + pS400 seeds.

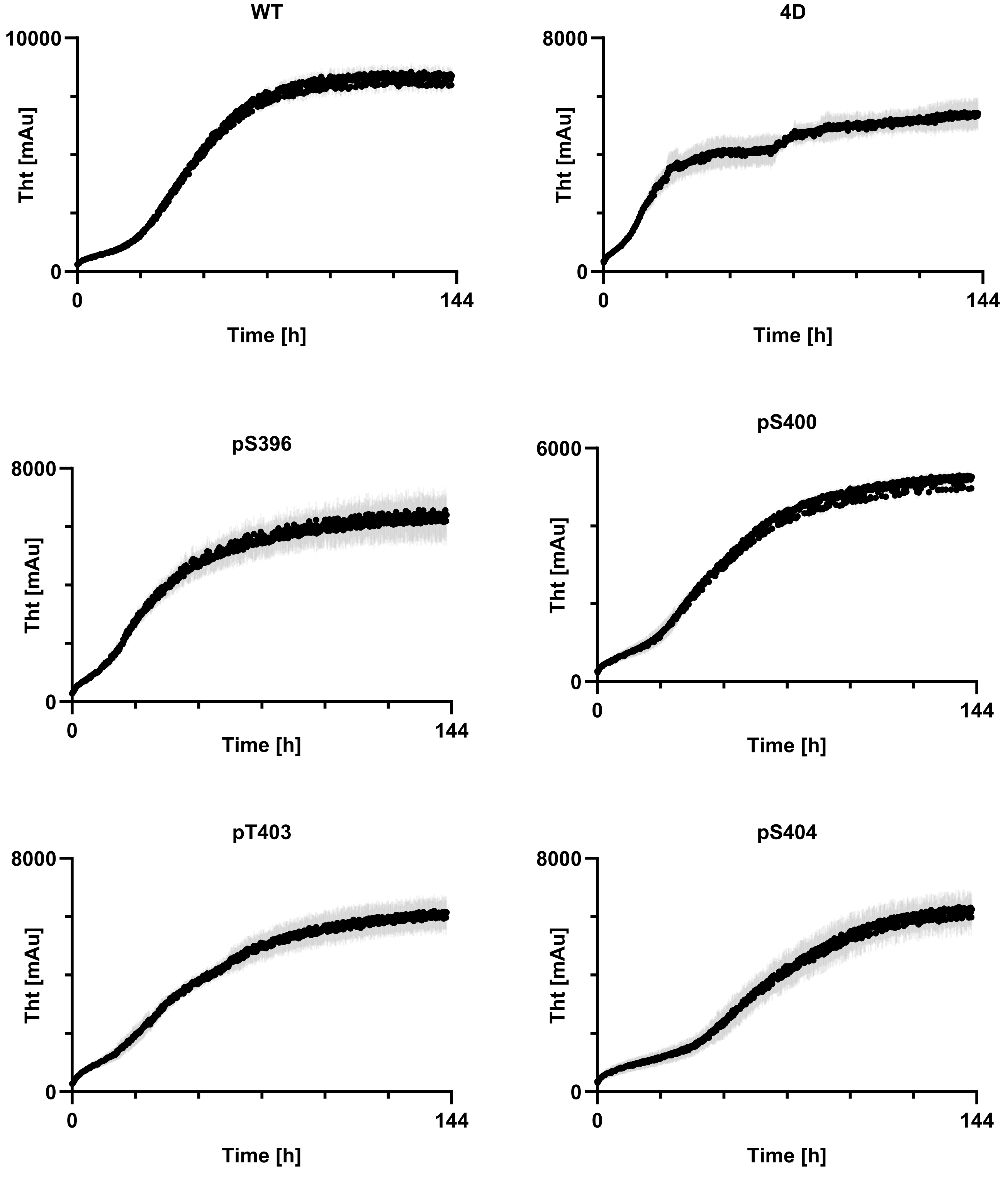

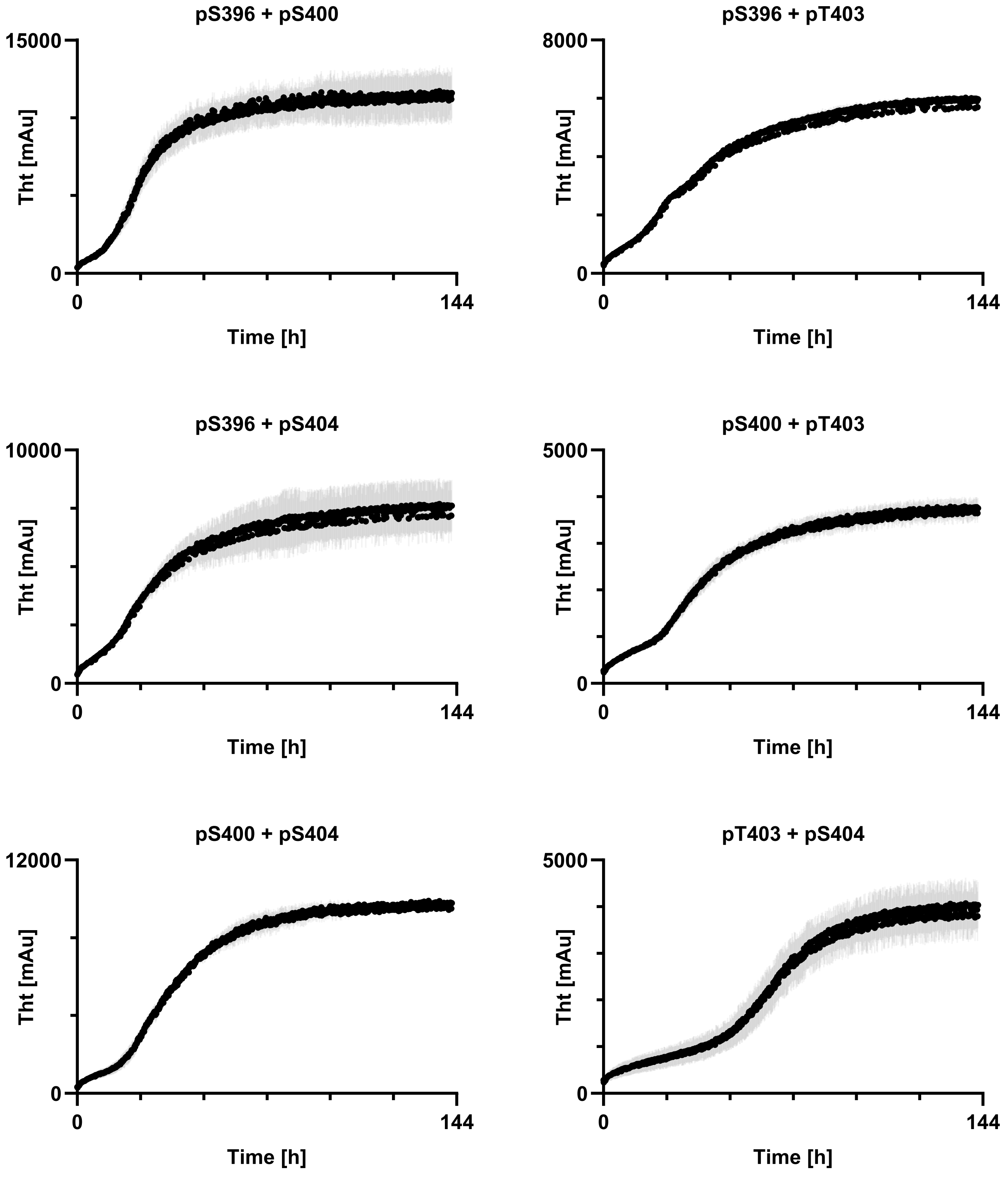

pS396 + pS400
15000
0
Tht [mAu]
0
144
Time [h]
pS396 + pT403
8000
0
Tht [mAu]
0
144
Time [h]
pS396 + pS404
10000
0
Tht [mAu]
0
144
Time [h]
pS400 + pT403
5000
0
Tht [mAu]
0
144
Time [h]
pS400 + pS404
12000
0
Tht [mAu]
0
144
Time [h]
pT403 + pS404
5000
0
Tht [mAu]
0
144
Time [h]

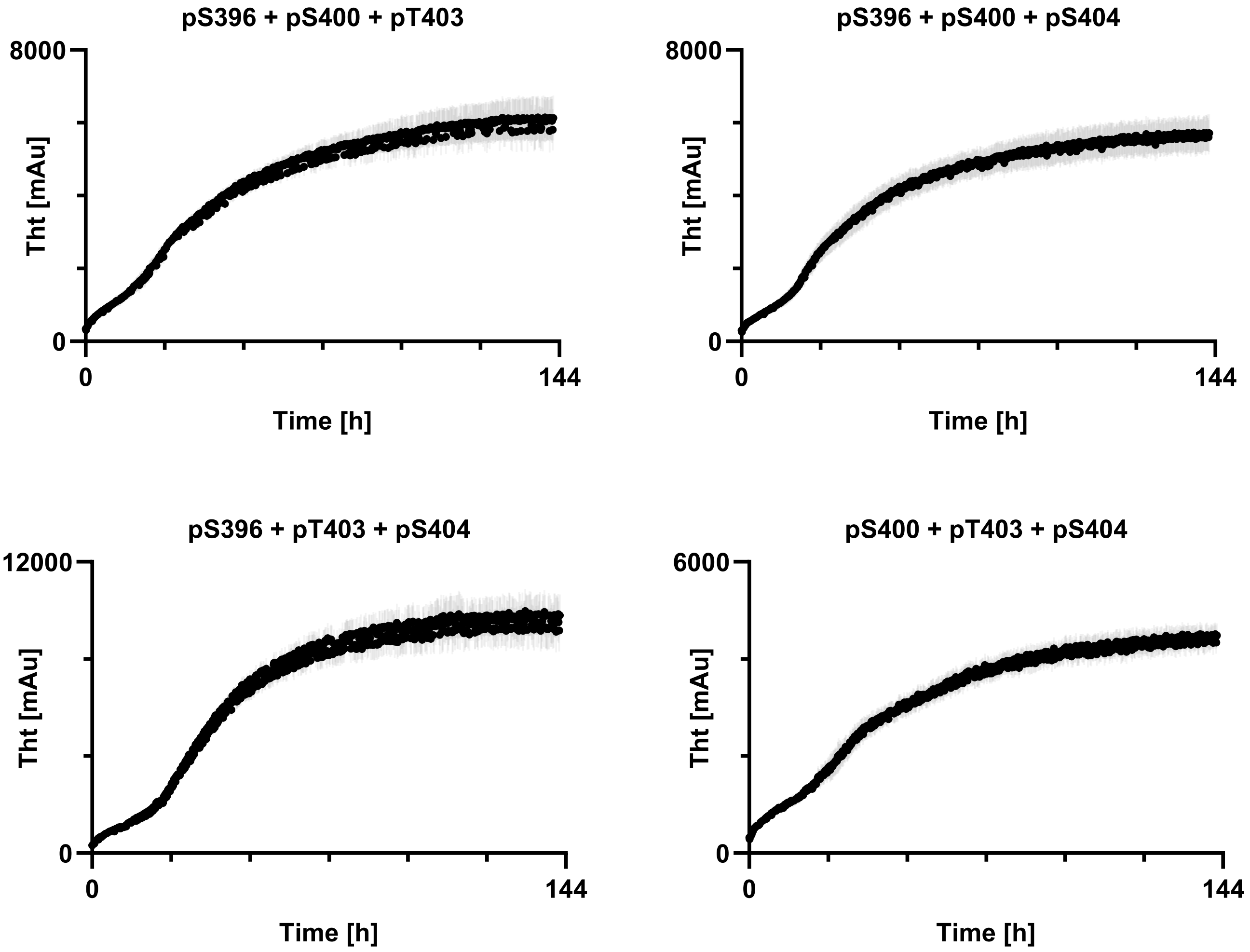

Graphical representation of $t_{1/2}$ values.

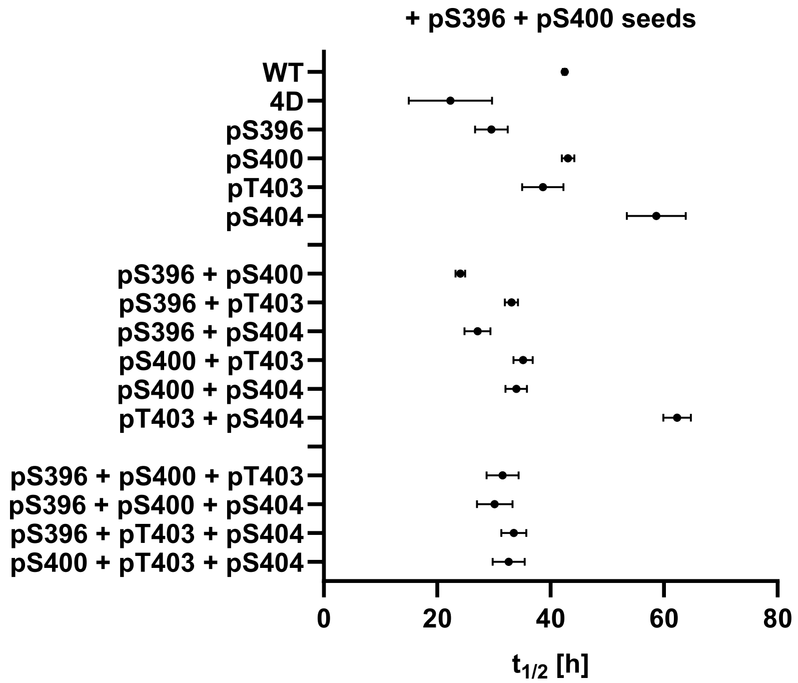

**Table of $t_{1/2}$ values and standard deviation after seeding with Tau(297-407) pS396 + pS400 seeds.**

| tau proteoform | 1 [h] | 2 [h] | 3 [h] | average [h] | STDEV [h] |
|---|---|---|---|---|---|
| WT | 42.1 | 42.7 | 42.6 | 42.5 | 0.3 |
| 4D | 19.2 | 17.1 | 30.7 | 22.3 | 7.3 |
| pS396 | 28.0 | 27.8 | 32.9 | 29.6 | 2.9 |
| pS400 | 42.2 | 42.7 | 44.3 | 43.1 | 1.1 |
| pT403 | 42.2 | 34.9 | 38.8 | 38.6 | 3.7 |
| pS404 | 53.0 | 63.2 | 59.7 | 58.6 | 5.2 |
| pS396 + pS400 | 23.2 | 24.1 | 24.9 | 24.1 | 0.9 |
| pS396 + pT403 | 32.0 | 33.0 | 34.3 | 33.1 | 1.2 |
| pS396 + pS404 | 24.7 | 27.4 | 29.2 | 27.1 | 2.3 |
| pS400 + pT403 | 34.1 | 34.2 | 37.1 | 35.1 | 1.7 |
| pS400 + pS404 | 31.9 | 34.2 | 35.7 | 33.9 | 1.9 |
| pT403 + pS404 | 62.4 | 59.8 | 64.7 | 62.3 | 2.5 |
| pS396 + pS400 + pT403 | 28.3 | 33.5 | 32.8 | 31.5 | 2.8 |
| pS396 + pS400 + pS404 | 26.8 | 30.6 | 33.0 | 30.1 | 3.1 |
| pS396 + pT403 + pS404 | 31.2 | 35.6 | 33.7 | 33.5 | 2.2 |
| pS400 + pT403 + pS404 | 31.4 | 30.6 | 35.8 | 32.6 | 2.8 |

**ThT traces after seeding with Tau(297-407) WT seeds.**

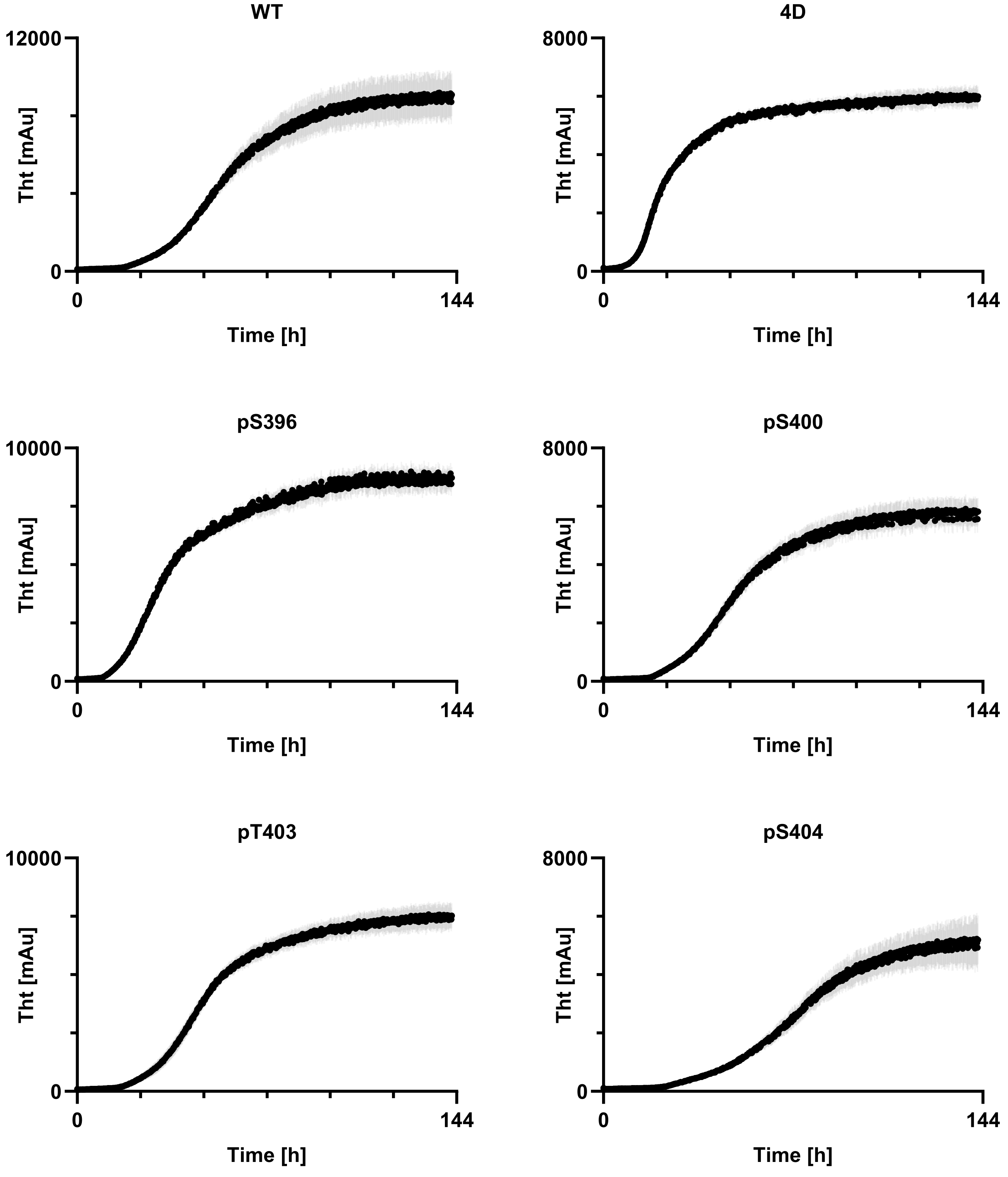

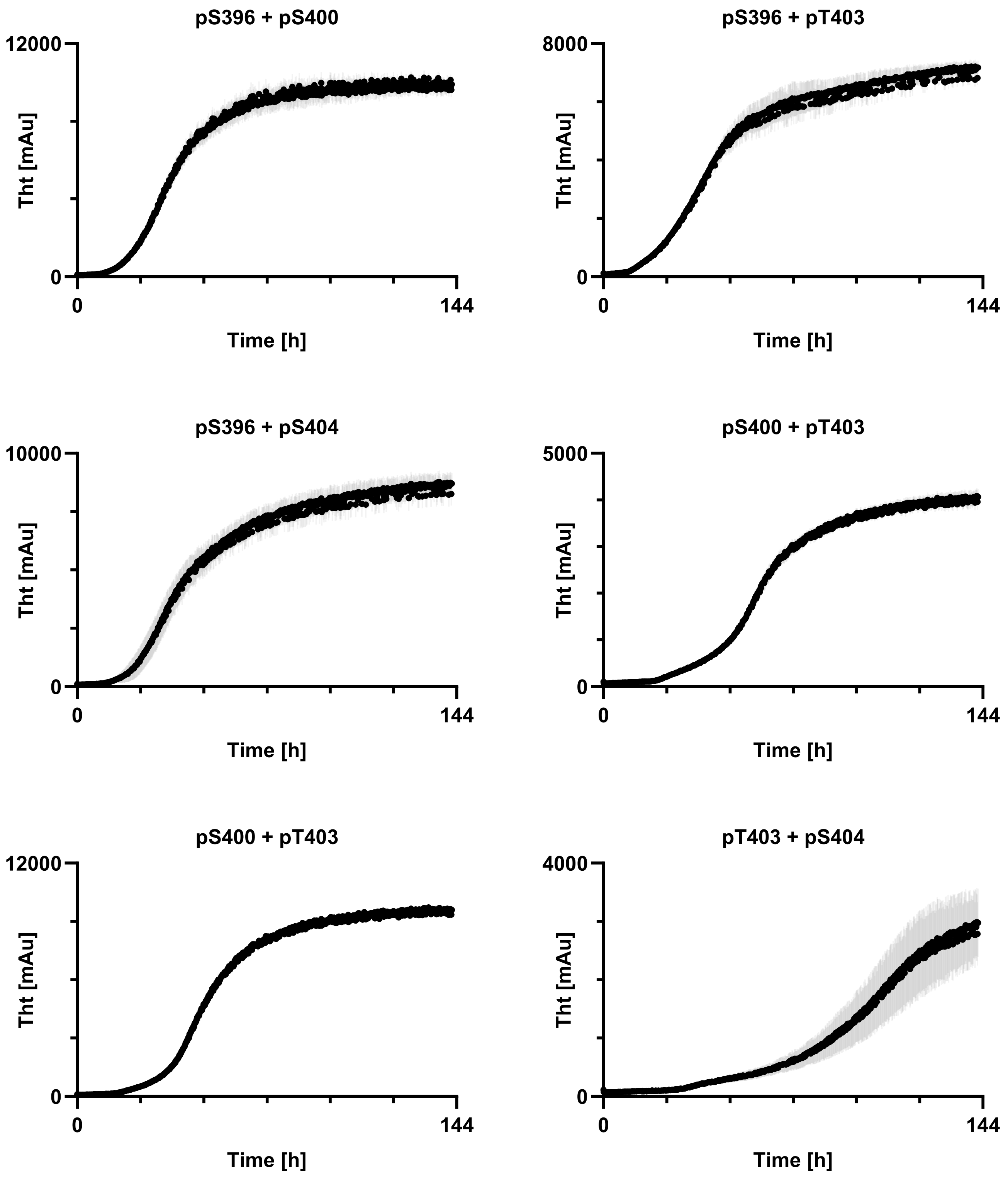
pS396 + pS400
12000
0
Tht [mAu]
0
144
Time [h]
pS396 + pT403
8000
0
Tht [mAu]
0
144
Time [h]
pS396 + pS404
10000
0
Tht [mAu]
0
144
Time [h]
pS400 + pT403
5000
0
Tht [mAu]
0
144
Time [h]
pS400 + pT403
12000
0
Tht [mAu]
0
144
Time [h]
pT403 + pS404
4000
0
Tht [mAu]
0
144
Time [h]

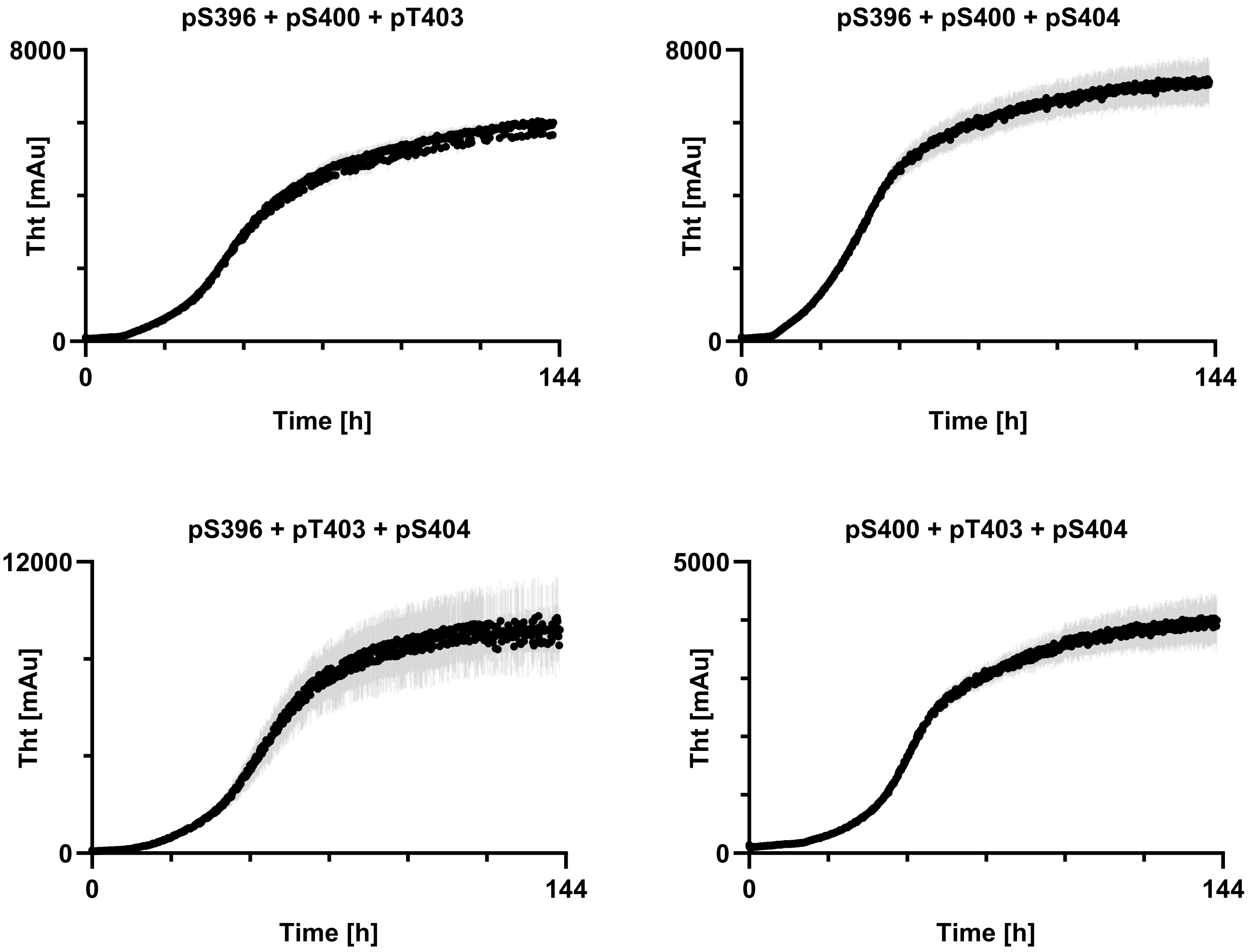

Graphical representation of $t_{1/2}$ values.

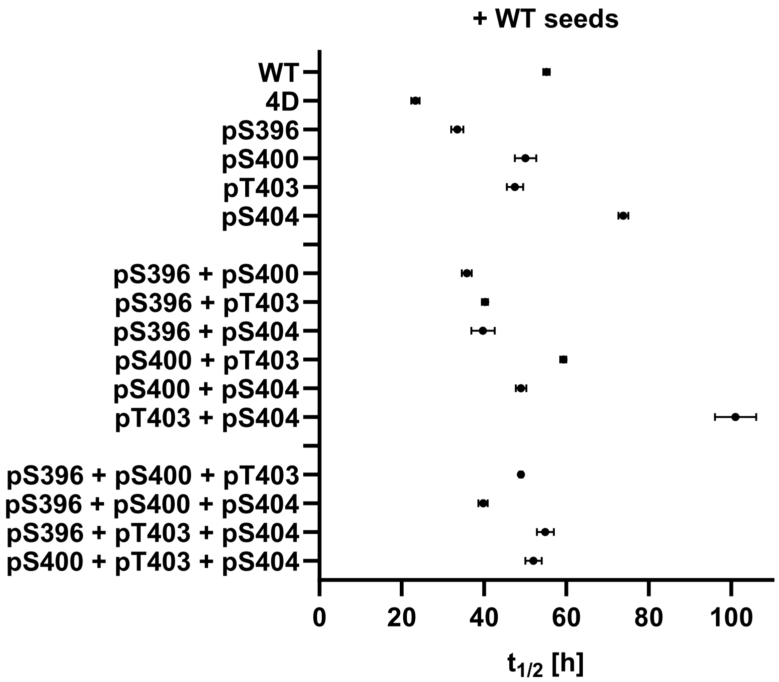

**Table of $t_{1/2}$ values and standard deviation after seeding with Tau(297-407) WT seeds.**

| tau proteoform | 1 [h] | 2 [h] | 3 [h] | average [h] | STDEV [h] |
|---|---|---|---|---|---|
| WT | 55.1 | 54.4 | 56.0 | 55.2 | 0.8 |
| 4D | 22.2 | 23.8 | 24.1 | 23.4 | 1.0 |
| pS396 | 33.9 | 34.7 | 31.9 | 33.5 | 1.4 |
| pS400 | 47.9 | 49.4 | 52.9 | 50.1 | 2.6 |
| pT403 | 46.5 | 46.2 | 49.8 | 47.5 | 2.0 |
| pS404 | 74.5 | 72.4 | 74.5 | 73.8 | 1.2 |
| pS396 + pS400 | 35.6 | 37.1 | 34.7 | 35.8 | 1.2 |
| pS396 + pT403 | 40.6 | 39.4 | 40.7 | 40.2 | 0.7 |
| pS396 + pS404 | 41.0 | 41.7 | 36.5 | 39.7 | 2.8 |
| pS400 + pT403 | 59.7 | 58.4 | 59.6 | 59.2 | 0.7 |
| pS400 + pS404 | 47.9 | 50.4 | 48.6 | 49.0 | 1.3 |
| pT403 + pS404 | 97.1 | 99.4 | 106.7 | 101.1 | 5.0 |
| pS396 + pS400 + pT403 | 48.8 | 49.4 | 48.7 | 49.0 | 0.4 |
| pS396 + pS400 + pS404 | 40.1 | 38.5 | 40.7 | 39.8 | 1.1 |
| pS396 + pT403 + pS404 | 53.0 | 54.5 | 57.1 | 54.9 | 2.1 |
| pS400 + pT403 + pS404 | 53.7 | 49.8 | 52.5 | 52.0 | 2.0 |

### S6. SOLUBILITY EQUILIBRIUM DETERMINATION.

**Calibration curve and HPLC method.** The calibration curves were generated using known concentrations of Tau(297-407)-4D, determined by A280. Each data point on the calibration curve is the average of three 10 µL injections, at the listed tau concentrations. The absorbance signal was generated by integrating the HPLC chromatogram at 214 nm.

**Analytical HPLC.** Measured on an Agilent Eclipse Plus C18 column (4.6 x 100 mm; 5 µm; 100 Å; 55 °C; 1 mL/min) with a gradient of 5-55% MeCN/$H_2O$/0.1% TFA over 10 minutes.

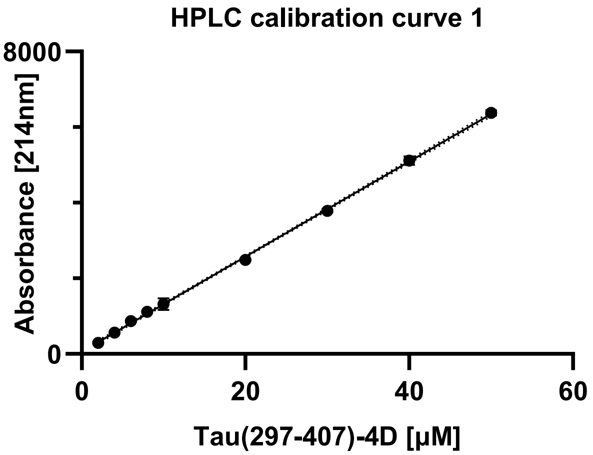

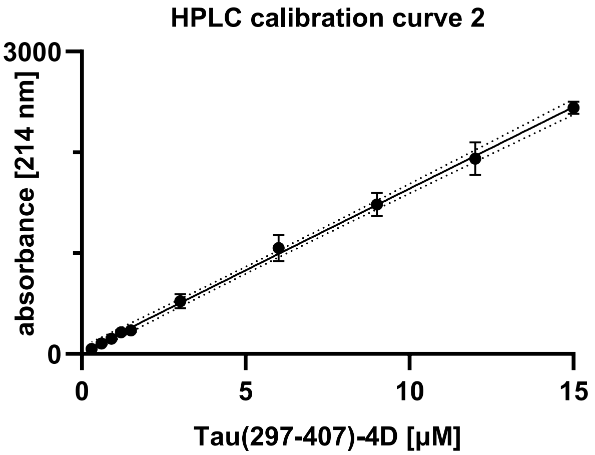

**Sedimentation assay.** Upon completion of the incubation, the assembly reactions (22 µL) were transferred to 1.6 mL Eppendorf tubes (Beckmann Coulter, Ref 357488) and centrifuged at 100 k x g for 60 min at 4 ºC. The supernatant (20 µL) was collected and transferred to a 0.3 mL HPLC vial. The concentration of tau remaining in the supernatant was determined by HPLC using a single 10 µL injection per sample. Each data point is the average of 3-6 10 µL injections. The monomer concentrations are reported as the average, and the error bars represent the standard deviation.

**Graphical representation of $C_{sat}$ values after primary nucleation.**

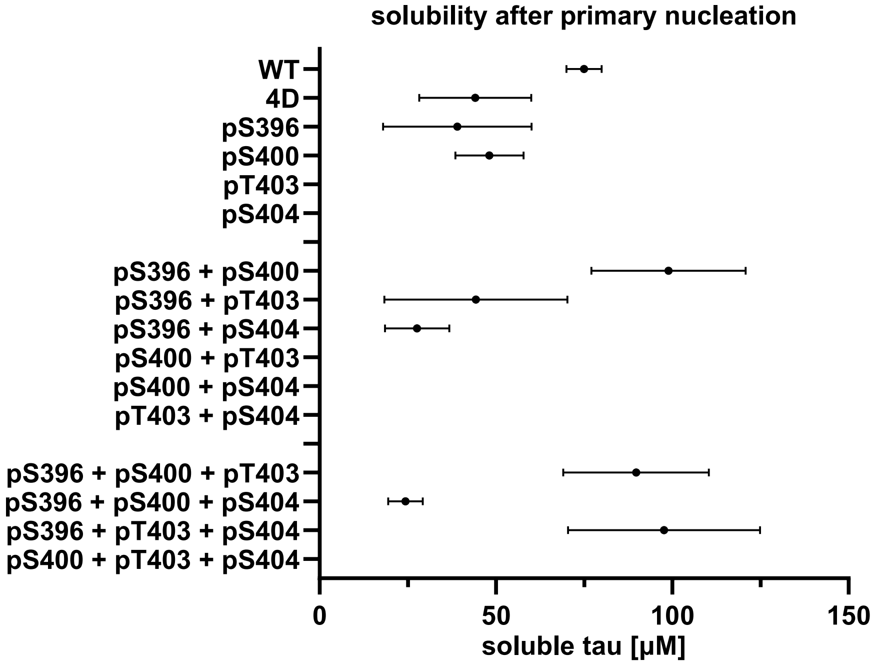

**Table of $C_{sat}$ values after primary nucleation.**

| tau proteoform | 1 [µM] | 2 [µM] | 3 [µM] | 4 [µM] | 5 [µM] | 6 [µM] | average [µM] | STDEV [µM] |
|---|---|---|---|---|---|---|---|---|
| WT | 79.7 | 75.5 | 69.8 | | | | 75.0 | 5.0 |
| 4D | 23.8 | 45.4 | 38.2 | 66.4 | 33.0 | 58.1 | 44.2 | 15.9 |
| pS396 | 63.8 | 25.5 | 24.7 | 27.8 | 24.2 | 68.5 | 39.1 | 21.1 |
| pS400 | 65.1 | 48.3 | 45.0 | 51.0 | 43.2 | 36.5 | 48.2 | 9.7 |
| pS396 + pS400 | 79.4 | 116.0 | 119.8 | 80.7 | | | 99.0 | 21.9 |
| pS396 + pT403 | 31.2 | 83.1 | 34.0 | 28.9 | | | 44.3 | 26.0 |
| pS396 + pS404 | 18.4 | 36.7 | 27.8 | | | | 27.7 | 9.1 |
| pS396 + pS400 + pT403 | 65.4 | 101.8 | 111.2 | 80.7 | | | 89.8 | 20.7 |
| pS396 + pS400 + pS404 | 28.9 | 28.2 | 19.5 | 20.8 | | | 24.4 | 4.9 |
| pS396 + pT403 + pS404 | 101.7 | 110.5 | 58.4 | 120.0 | | | 97.7 | 27.2 |

- The solubility equilibrium data indicate that there is no clear trend in solubility between the constructs.

**Graphical representation of $C_{sat}$ values after seeding with Tau(297-407)-4D seeds.**

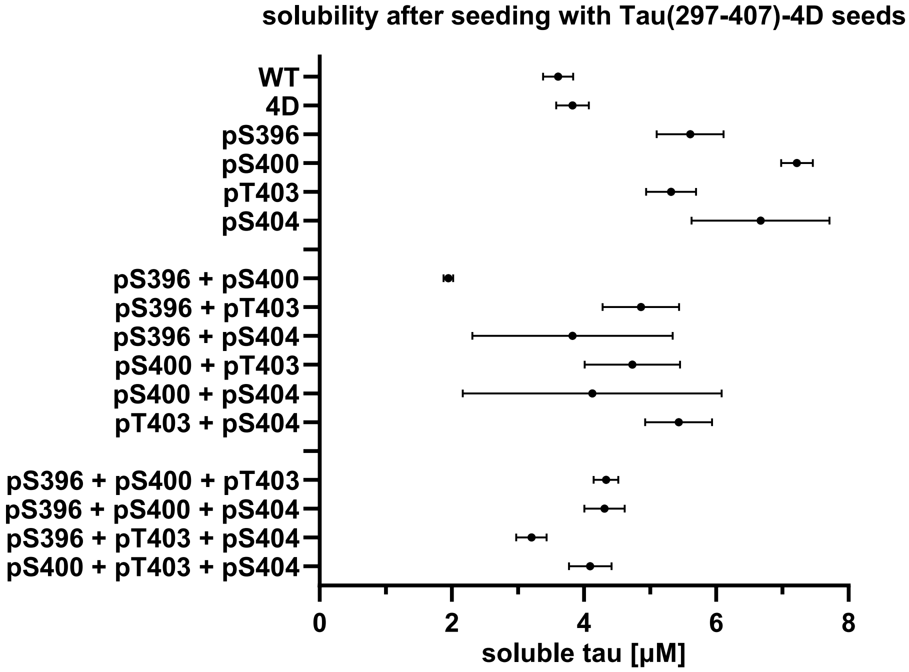

**Table of $C_{sat}$ values after seeding with Tau(297-407)-4D seeds.**

| tau proteoform | 1 [μM] | 2 [μM] | 3 [μM] | average [μM] | STDEV [μM] |
|---|---|---|---|---|---|
| WT | 3.54 | 3.42 | 3.86 | 3.61 | 0.23 |
| 4D | 4.08 | 3.80 | 3.59 | 3.83 | 0.25 |
| pS396 | 5.87 | 5.92 | 5.02 | 5.60 | 0.51 |
| pS400 | 7.47 | 7.19 | 7.00 | 7.22 | 0.24 |
| pT403 | 5.31 | 4.94 | 5.70 | 5.32 | 0.38 |
| pS404 | 7.36 | 5.47 | 7.18 | 6.67 | 1.04 |
| pS396 + pS400 | 2.02 | 1.95 | 1.87 | 1.95 | 0.07 |
| pS396 + pT403 | 4.56 | 5.53 | 4.49 | 4.86 | 0.58 |
| pS396 + pS404 | 3.14 | 5.56 | 2.77 | 3.83 | 1.51 |
| pS400 + pT403 | 5.51 | 4.59 | 4.09 | 4.73 | 0.72 |
| pS400 + pS404 | 6.38 | 3.09 | 2.90 | 4.12 | 1.96 |
| pT403 + pS404 | 5.00 | 5.99 | 5.31 | 5.43 | 0.51 |
| pS396 + pS400 + pT403 | 4.13 | 4.36 | 4.50 | 4.33 | 0.19 |
| pS396 + pS400 + pS404 | 4.00 | 4.61 | 4.32 | 4.31 | 0.30 |
| pS396 + pT403 + pS404 | 3.00 | 3.45 | 3.16 | 3.20 | 0.23 |
| pS400 + pT403 + pS404 | 4.05 | 4.44 | 3.80 | 4.09 | 0.32 |

**Graphical representation of $C_{sat}$ values after seeding with Tau(297-407) pS396 pS400 seeds.**

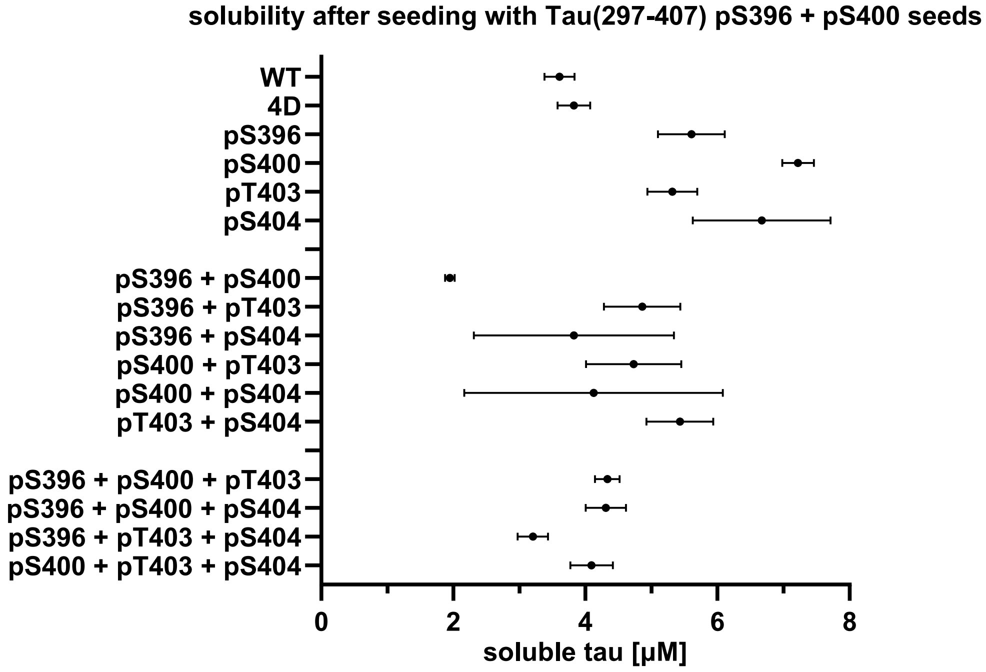

**Table of $C_{sat}$ values after seeding with Tau(297-407) pS396 pS400 seeds.**

| tau proteoform | 1 [µM] | 2 [µM] | 3 [µM] | average [µM] | STDEV [µM] |
|---|---|---|---|---|---|
| WT | 1.85 | 1.67 | 1.99 | 1.83 | 0.16 |
| 4D | 8.89 | 6.57 | 6.13 | 7.20 | 1.48 |
| pS396 | 3.88 | 2.89 | 3.90 | 3.56 | 0.58 |
| pS400 | 6.19 | 5.81 | 6.59 | 6.20 | 0.39 |
| pT403 | 3.45 | 4.18 | 3.65 | 3.76 | 0.38 |
| pS404 | 5.64 | 5.49 | 5.52 | 5.55 | 0.08 |
| pS396 + pS400 | 0.57 | 0.44 | 0.95 | 0.65 | 0.26 |
| pS396 + pT403 | 3.70 | 3.87 | 3.35 | 3.64 | 0.26 |
| pS396 + pS404 | 2.35 | 2.60 | 2.52 | 2.49 | 0.12 |
| pS400 + pT403 | 4.41 | 3.80 | 4.02 | 4.08 | 0.31 |
| pS400 + pS404 | 1.49 | 1.23 | 1.58 | 1.44 | 0.18 |
| pT403 + pS404 | 5.32 | 4.57 | 5.61 | 5.17 | 0.54 |
| pS396 + pS400 + pT403 | 3.03 | 3.11 | 3.39 | 3.18 | 0.19 |
| pS396 + pS400 + pS404 | 4.70 | 3.59 | 3.49 | 3.93 | 0.67 |
| pS396 + pT403 + pS404 | 1.73 | 2.02 | 1.79 | 1.84 | 0.15 |
| pS400 + pT403 + pS404 | 3.06 | 3.45 | 3.34 | 3.28 | 0.20 |

**Graphical representation of $C_{sat}$ values after seeding with Tau(297-407) WT seeds.**

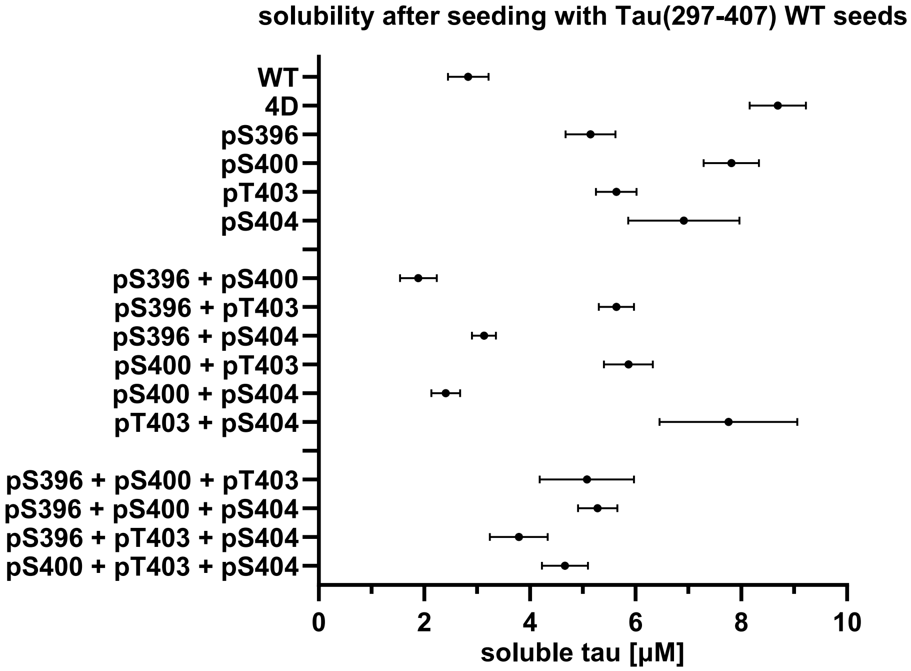

**Table of $C_{sat}$ values after seeding with Tau(297-407) WT seeds.**

| tau proteoform | 1 [µM] | 2 [µM] | 3 [µM] | average [µM] | STDEV [µM] |
|---|---|---|---|---|---|
| WT | 2.45 | 2.82 | 3.22 | 2.83 | 0.38 |
| 4D | 9.11 | 8.09 | 8.87 | 8.69 | 0.53 |
| pS396 | 4.74 | 5.66 | 5.03 | 5.14 | 0.47 |
| pS400 | 7.26 | 7.86 | 8.31 | 7.81 | 0.52 |
| pT403 | 5.19 | 5.81 | 5.90 | 5.63 | 0.38 |
| pS404 | 7.47 | 5.70 | 7.57 | 6.91 | 1.05 |
| pS396 + pS400 | 2.06 | 1.49 | 2.11 | 1.89 | 0.35 |
| pS396 + pT403 | 5.26 | 5.89 | 5.76 | 5.64 | 0.33 |
| pS396 + pS404 | 2.91 | 3.37 | 3.11 | 3.13 | 0.23 |
| pS400 + pT403 | 6.09 | 5.33 | 6.17 | 5.86 | 0.46 |
| pS400 + pS404 | 2.10 | 2.62 | 2.49 | 2.40 | 0.27 |
| pT403 + pS404 | 6.93 | 7.08 | 9.26 | 7.76 | 1.31 |
| pS396 + pS400 + pT403 | 4.26 | 6.03 | 4.94 | 5.08 | 0.89 |
| pS396 + pS400 + pS404 | 5.36 | 4.87 | 5.60 | 5.28 | 0.37 |
| pS396 + pT403 + pS404 | 3.43 | 3.52 | 4.42 | 3.79 | 0.55 |
| pS400 + pT403 + pS404 | 4.35 | 4.47 | 5.16 | 4.66 | 0.44 |

### S7. CHEMICAL DENATURATION.

**Denaturation conditions.** Denaturant gradient, 2 μM Tau(297-407) seeds, 10 mM KPB-KOH pH 7.4, 10 mM DTT, 10 μM ThT, 23 ºC, 16 h.

**Stock solutions.**

- buffer: 10 mM KPB-KOH, 10 mM DTT, 10 μM ThT pH 7.4.
- 2x fibril: 4 μM Tau(297-407) fibrils diluted in buffer.
- 2x Gnd·HCl: 5 M Gnd·HCl in buffer pH 7.4.
- 2x urea: 7 M urea in buffer pH 7.4.

**Chemical denaturation.** The denaturation was performed in black, non-binding, low-volume 384-well microplates (Greiner 784900), sealed with adhesive microplate film (VWR 7659). A 1.6 mL Eppendorf tube was filled with 2x fibril master mix, and three wells were filled with 5 μL of 2x fibril and 2x denaturant, for each denaturant concentration. The plate was incubated at 23 °C for 16 h, then analyzed for ThT fluorescence using a microplate reader. The reactions were performed in quadruplicate. The denaturation curve ThT values were averaged, and the error bars represent the standard error of the mean. The $IC_{50}$ values were obtained by fitting the curves to sigmoidal functions, and the midpoints are reported with their standard errors.

**Denaturation curves after primary nucleation.**

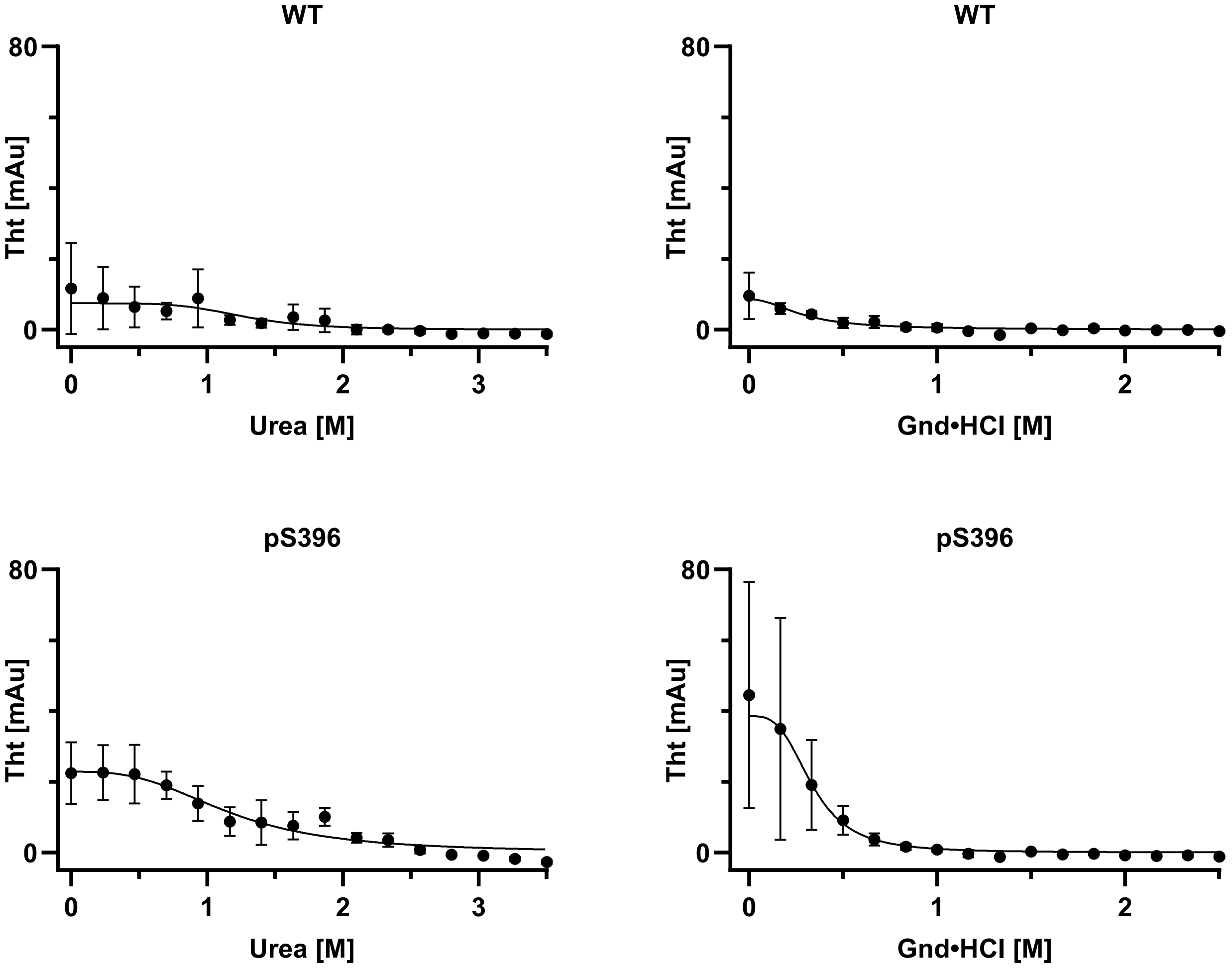

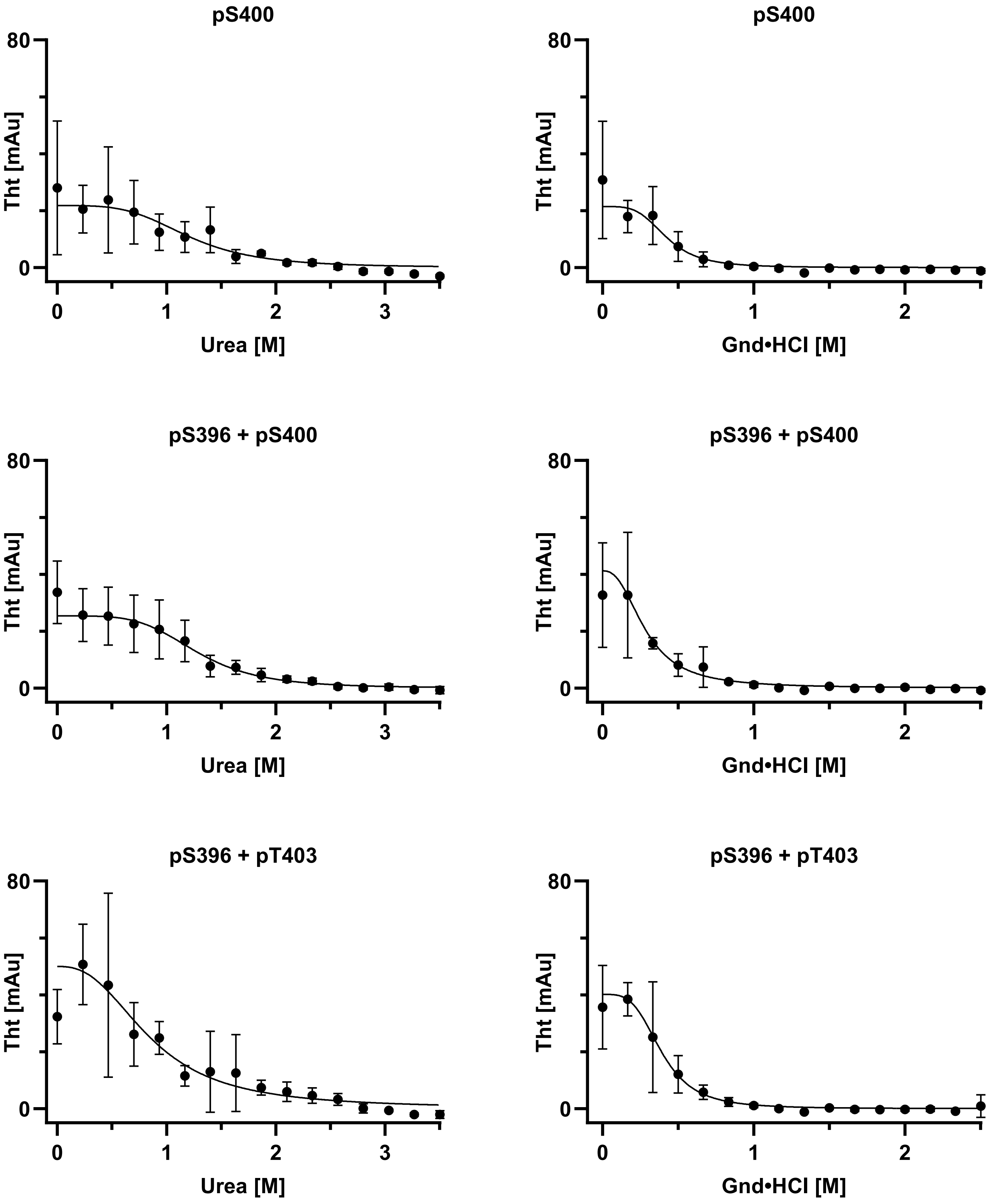
pS400
Tht [mAu]
Urea [M]
pS400
Tht [mAu]
Gnd•HCl [M]
pS396 + pS400
Tht [mAu]
Urea [M]
pS396 + pS400
Tht [mAu]
Gnd•HCl [M]
pS396 + pT403
Tht [mAu]
Urea [M]
pS396 + pT403
Tht [mAu]
Gnd•HCl [M]

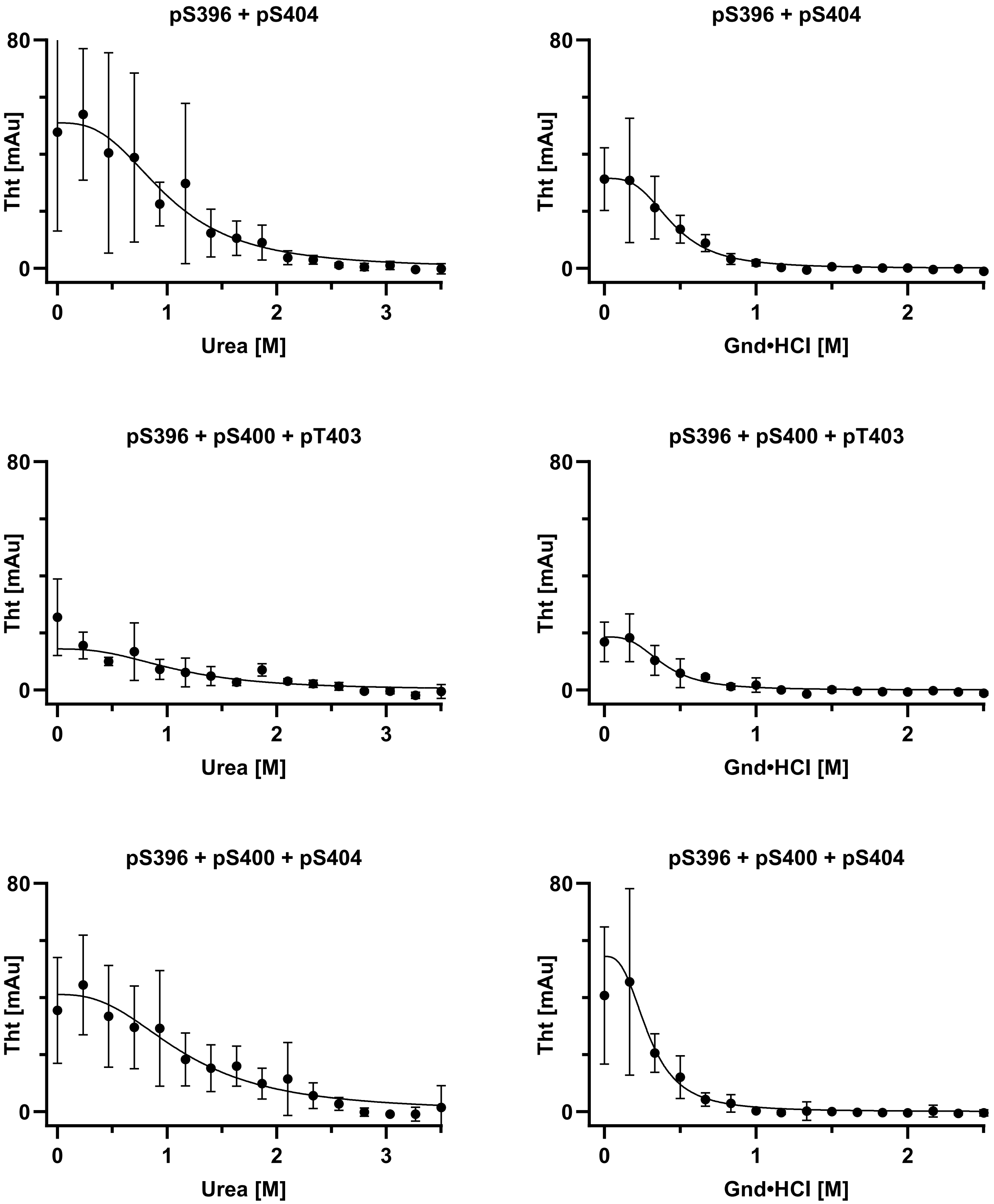

pS396 + pS404
pS396 + pS400 + pT403
pS396 + pS400 + pS404
Tht [mAu]
80
0
Urea [M]
Gnd•HCl [M]
0
1
2
3

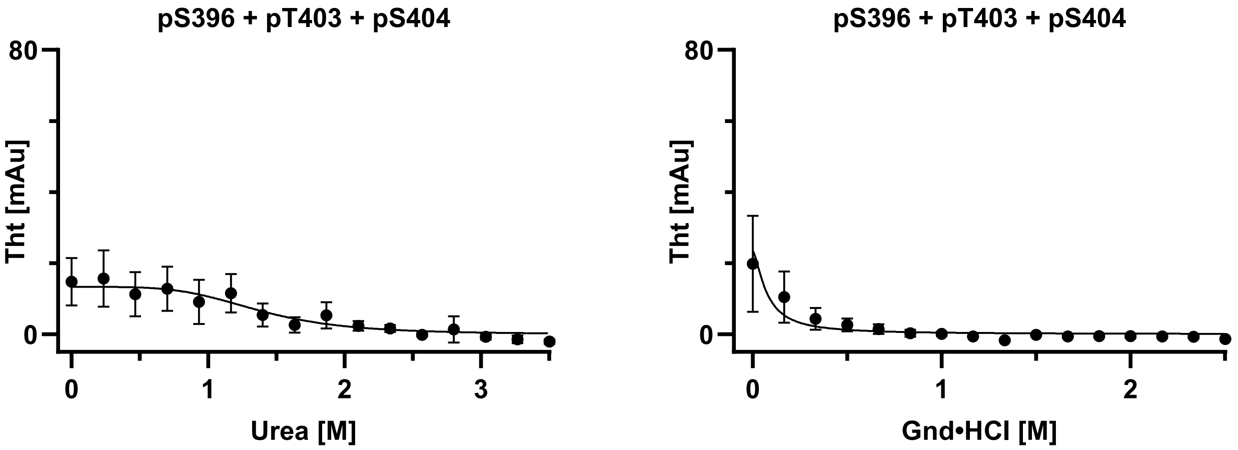

Graphical $IC_{50}$ values for chemical denaturation after primary nucleation.

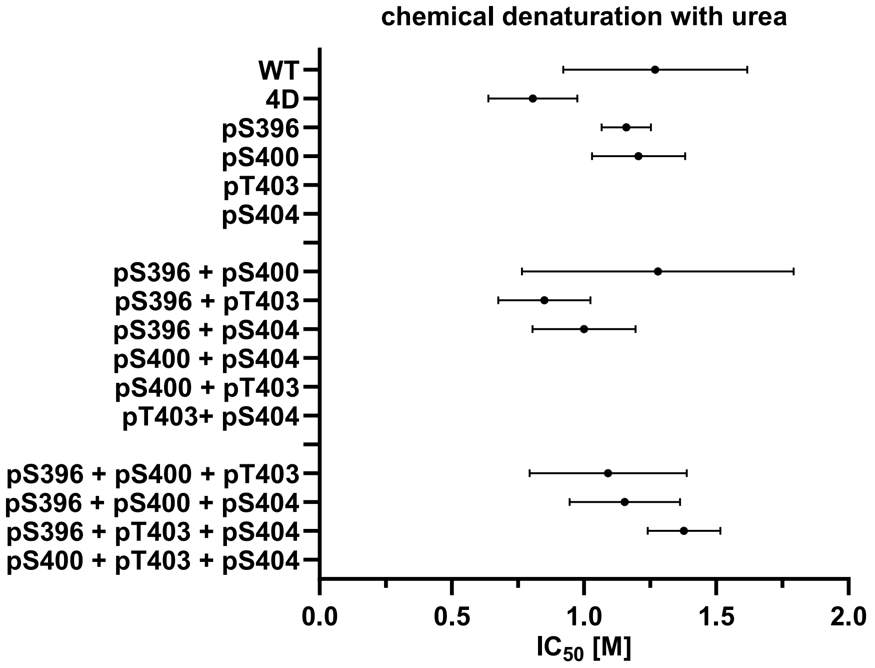

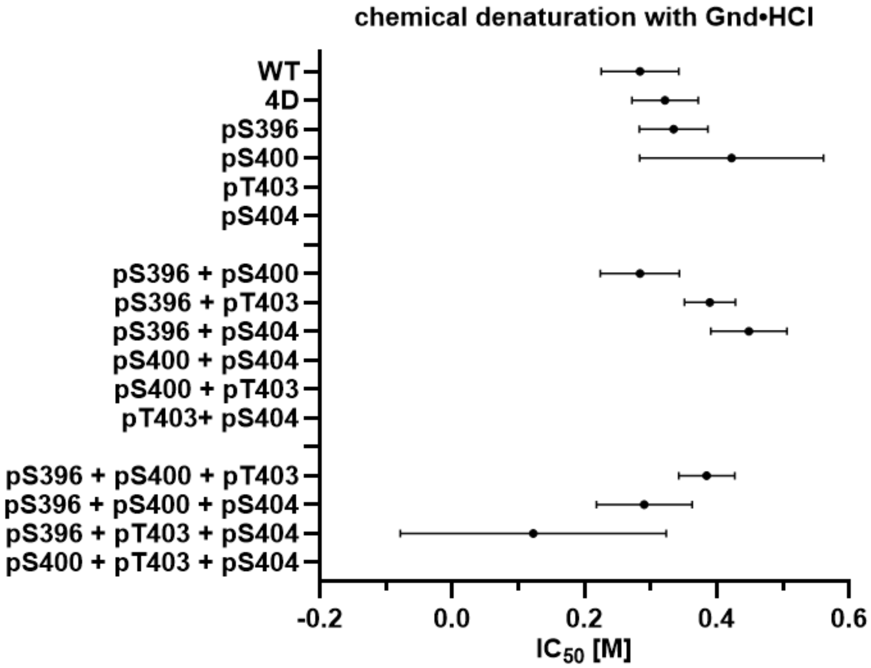

**Graphical $IC_{50}$ values for chemical denaturation after primary nucleation.**

| tau proteoform | urea $IC_{50}$ [M] | urea error [M] | Gnd·HCl $IC_{50}$ [M] | Gnd·HCl error [M] |
|---|---|---|---|---|
| WT | 1.27 | 0.35 | 0.28 | 0.06 |
| 4D | 0.81 | 0.17 | 0.32 | 0.05 |
| pS396 | 1.16 | 0.09 | 0.33 | 0.05 |
| pS400 | 1.21 | 0.18 | 0.42 | 0.14 |
| pS396 + pS400 | 1.28 | 0.51 | 0.28 | 0.06 |
| pS396 + pT403 | 0.85 | 0.17 | 0.39 | 0.04 |
| pS396 + pS404 | 1.00 | 0.20 | 0.45 | 0.06 |
| pS396 + pS400 + pT403 | 1.09 | 0.30 | 0.39 | 0.04 |
| pS396 + pS400 + pS404 | 1.15 | 0.21 | 0.29 | 0.07 |
| pS396 + pT403 + pS404 | 1.39 | 0.14 | 0.12 | 0.20 |

### S8. DETAILS OF THE CRYO-EM PROCESSING.

**Cryo-electron microscopy sample preparation (cryo-EM).** Various dilutions (i.e. 1:5, 1:10, 1:20) of fibrillized phosphorylated Tau(297-407) in fibrillization buffer (10 mM KPB-KOH pH 7.5, 200 mM potassium citrate tribasic, 10 mM DTT, 3 μM ThT) were prepared after 1 week of quiescent incubation at 37 °C and followed by an additional 4 weeks of quiescent incubation at 23 °C. At the time points, 3 μL of diluted sample was applied to the front side of holey carbon grids (Quantifoil R1.2/1.3 on a gold 300 mesh support) that were glow-discharged for 120 s. Then, 1.5 μL of fibrillization buffer was applied to the back side. The grids were back-blotted (i.e., the blotting paper contacted the back side only) for 8 s at 4 °C and 100% humidity using a Leica GP2 (Leica Microsystems, Waltham, MA), followed by plunge freezing in liquid ethane.

All grids were screened on a Glacios (Thermo Fisher Scientific, Waltham, MA) operated at 200 kV. Two constructs (pS396+pT403; pS400) resulted in sufficient quantities of twisted filaments with well-defined crossovers suitable for structural determination, while all other samples gave non-twisting ribbons or filaments with irregular widths and crossovers. For the pS396 + pT403 sample, a total of 10,331 movies were collected at a nominal magnification of 165,000× (physical pixel size: 0.728 Å/pixel) on a Titan Krios G4 (Thermo Fisher Scientific, Waltham, MA) operated at 300 kV and equipped with a Falcon 4i direct electron detector and Selectris X energy filter (Thermo Fisher Scientific, Waltham, MA) set to a slit width of 20 eV. A defocus range of −0.8 to −1 μm was used with a total exposure time of 4.5-5 s fractionated into 90 TIFF frames. The total dose for each movie was 45 electrons/Å$^2$. Movies were motion corrected using MotionCor2 [1] in Scipion.[2] Motion-corrected and dose-weighted micrographs were manually curated in Scipion to remove micrographs lacking filaments, with too high filament density, at low resolution, or with significant ice contamination, resulting in 832 remaining micrographs. For the pS400 sample, a total of 9,823 movies were collected using the same settings as the pS396+pT403 sample. After curation, 1,805 micrographs were used for further processing.

**Cryo-EM data processing.** All image processing was done in RELION 5 and 5.1. [3-6] Dose-weighted summed micrographs were imported into RELION 5. The contrast transfer function was estimated using CTFFIND-4.1.[7] For the pS396+pT403 sample, filaments were picked manually, and segments were extracted from the CTF-corrected micrographs with a box size of 900 pixels downscaled to 300 pixels, resulting in 122,440 segments. Reference-free 2D classification was used to remove contaminants and segments contributing to straight filaments, resulting in 26,038 remaining segments. From these 2D classes, we generated a 3D ab initio model using RELION's relion_helix_inimodel2d feature. The segments were then re-extracted with a box size of 288 pixels without downscaling. One round of 3D classification with image alignment was performed on the segments using the ab initio model, 3 classes (i.e. k=3), a regularization parameter (T) of 15, and allowing helical rise and twist parameters to vary, starting from an initial rise of 4.78 Å and twist of -1.3°. After 35 iterations, the highest-resolution class (12,980 segments) was subjected to one round of 3D auto-refinement imposing $C2_1$ symmetry with the helical parameters and using Blush regularization with the AmyBlush (amy-v1.0) network in RELION 5.1. The refined map underwent standard mask creation and post-processing, to give the final pS396+pT403 map with a global resolution of 3.6 Å, a helical rise of 2.39 Å, and a twist of 179.47°.

For the pS400 sample, filaments were picked manually and segments were extracted with a box size of 900 pixels downscaled to 300 pixels, resulting in 372,025 segments. Multiple rounds of reference-free 2D classification were then performed to identify 2 different major polymorphs: polymorph 1 (122,580 segments, 33% of the data) and polymorph 2 (103,301 segments, 28% of the data). From the highest resolution 2D classes, we generated separate 3D ab initio models as with the pS396+pT403 sample and re-extracted segments with a box size of 288 pixels.

Segments in polymorph 1 underwent two rounds of 3D classification. The first round used the ab initio model as a reference, 25 iterations, k=5, T=10, initial twist -1.2°, initial rise 4.75 Å, allowing helical parameters to vary. The best class (38,134 segments) was used for a second round of 3D classification (40 iterations, k=3, T=20, Blush regularization with the AmyBlush network, initial twist -1.4°, initial rise 4.76 Å, allowing helical parameters to vary). The best class (15,578 segments) underwent 3D auto-refinement using Blush regularization (AmyBlush) then standard mask creation and post-processing, to give the final pS400-1 map with a global resolution of 3.7 Å, a helical rise of 4.76 Å, and a twist of -1.40°.

Segments in polymorph 2 underwent three rounds of 3D classification. The first round used the ab initio model as a reference, 38 iterations, k=5, T=10, initial twist -1.1°, initial rise 4.75 Å, allowing helical parameters to vary. The best class (52,936 segments) was used for a second round of 3D classification (25 iterations, k=4, T=10, initial twist -1.0°, initial rise 4.78 Å, fixed helical parameters). The best class from the second round (32,780 segments) was used for a third round of 3D classification (40 iterations, k=4, T=10, Blush regularization with AmyBlush network, initial twist -1.0°, initial rise 4.78 Å, allowing helical parameters to vary). The best class from the third round (24,468 segments) underwent 3D auto-refinement using Blush regularization (AmyBlush) then standard mask creation and post-processing, to give the final pS400-2 map with a global resolution of 3.6 Å, a helical rise of 4.77 Å, and a twist of -1.36°.

**Model building and analysis.** Initial model building was performed by fitting existing in vitro tau filament models into the refined maps (pS396+pT403: PDB 7QJW, pS400-1: PDB 8Q9F, pS400-2: 8QJJ). Subsequent model fitting and refinement was performed using ISOLDE.[8] For pS400-2, ModelAngelo was used to determine the identity of the 19-residue peptide flanking the main protofilament. [9] After fitting into the density, each protofilament dimer was translated to give a stack of three rungs (6 strands total per model). After Ramachandran parameters, rotamers, and clashes were satisfied in the middle rung, the outer two rungs were deleted, and the middle rung was re-translated into a new eight-rung stack (16 strands per model). Afterwards, the ISOLDE command "isolde write phenixRsrInput #<model> <map resolution> #<map>" was used to export the model and generate a rigid-body refinement settings file for real-space refinement in PHENIX,[10] giving the final refined models. (Note that the validation of 3-rung models often gave spurious map-model FSC curves that started below FSC 0.5). Data were deposited into the Protein Data Bank (PDB) under accession codes 13GX (pS396+pT403), 13GY (pS400-1), 13GZ (pS400-2) and into the Electron Microscopy Data Bank (EMDB) under accession codes EMD-77062 (pS396+pT403), EMD-77063 (pS400-1), and EMD-77064 (pS400-2).

Amyloid packing difference (APD) scores were calculated and associated figures were generated as previously described. [11] Other structural figures were made using UCSF ChimeraX.[12]

**S8a.** Cryo-EM micrographs.

**Cryo-EM micrographs of pS396 + pS400 + pT403 (1 week).**

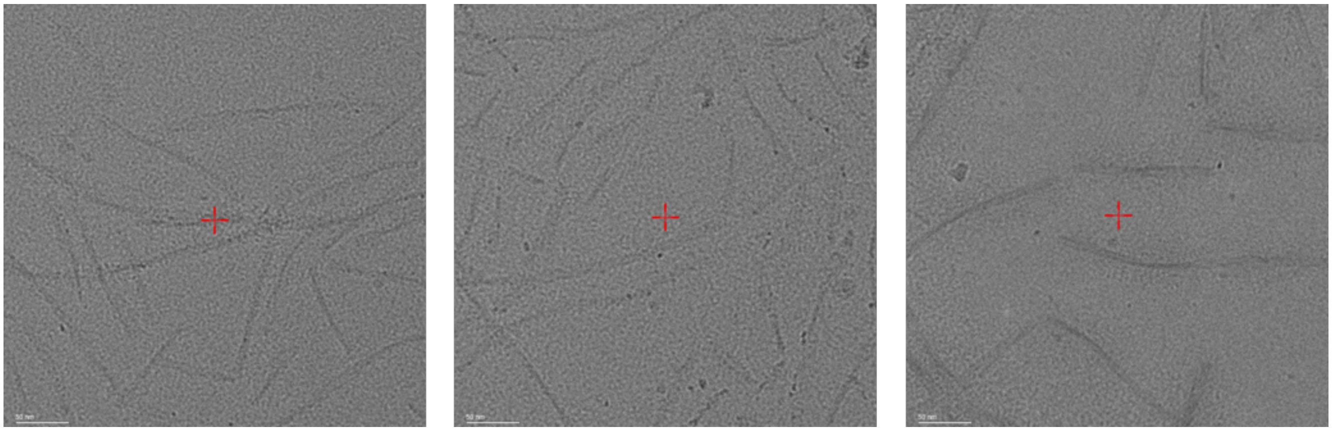

**Cryo-EM micrographs of pS396 + pS404 (1 week).**

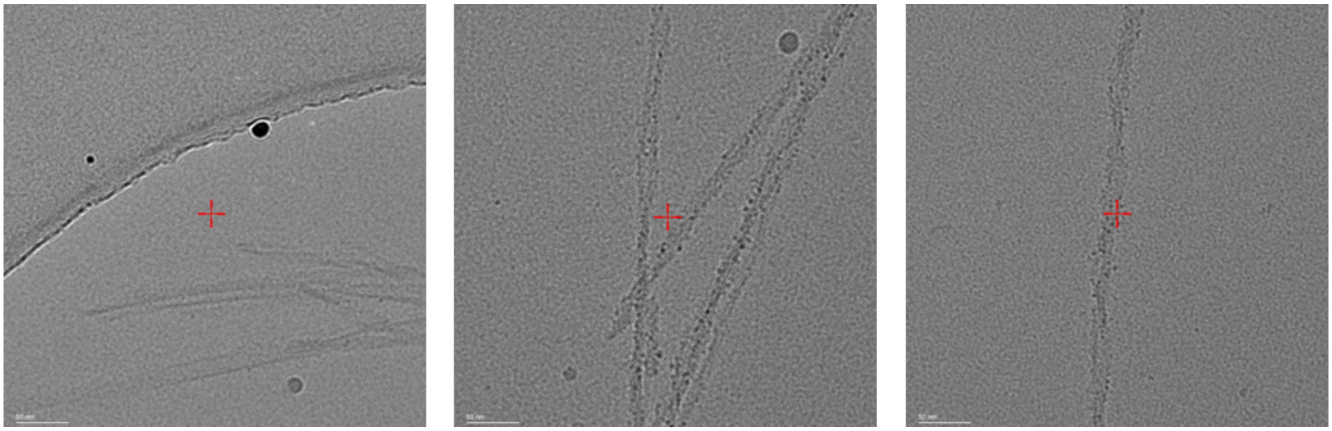

**Cryo-EM micrographs of pS396 + pT403 (1 weeks).**

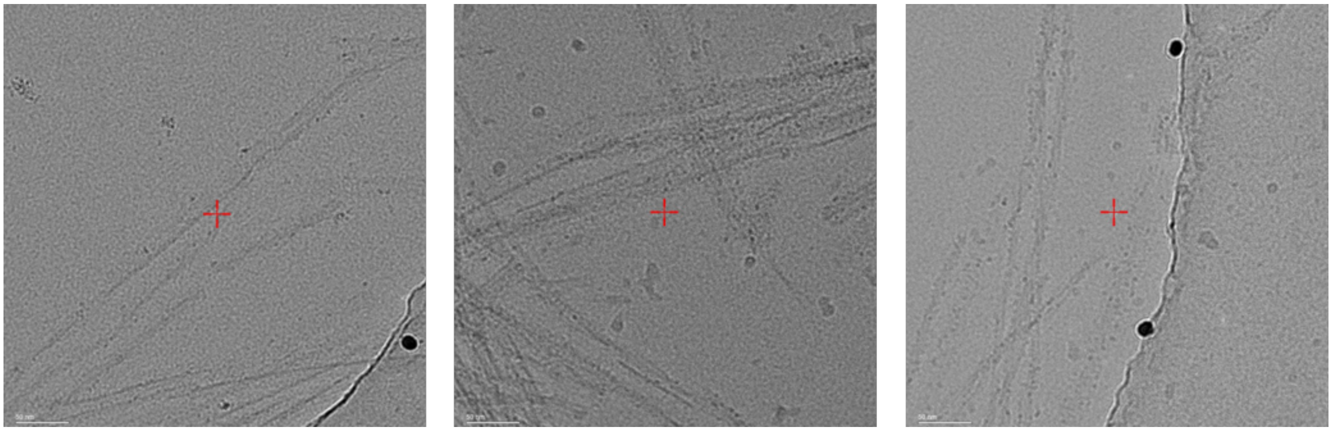

**Cryo-EM micrographs of pS400 (1 weeks).**

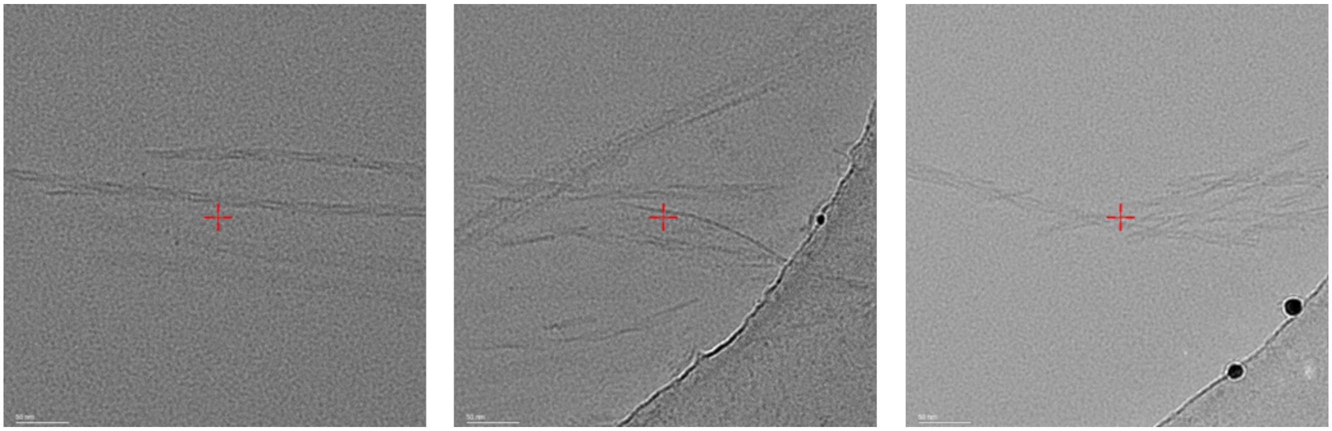

**Cryo-EM micrographs of pS396 + pS404 (5 weeks).**

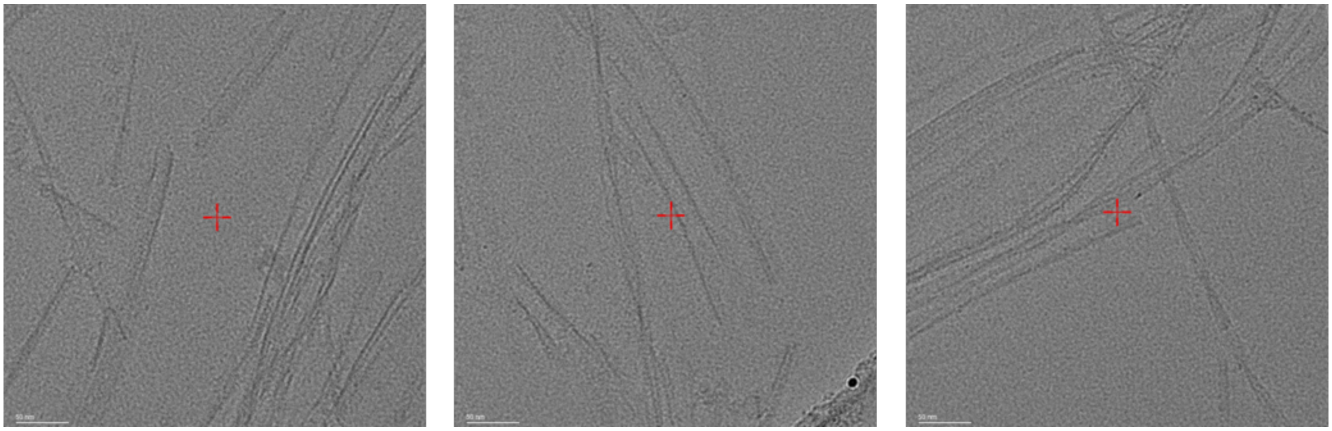

**Cryo-EM micrographs of pS396 + pT403 (5 weeks).**

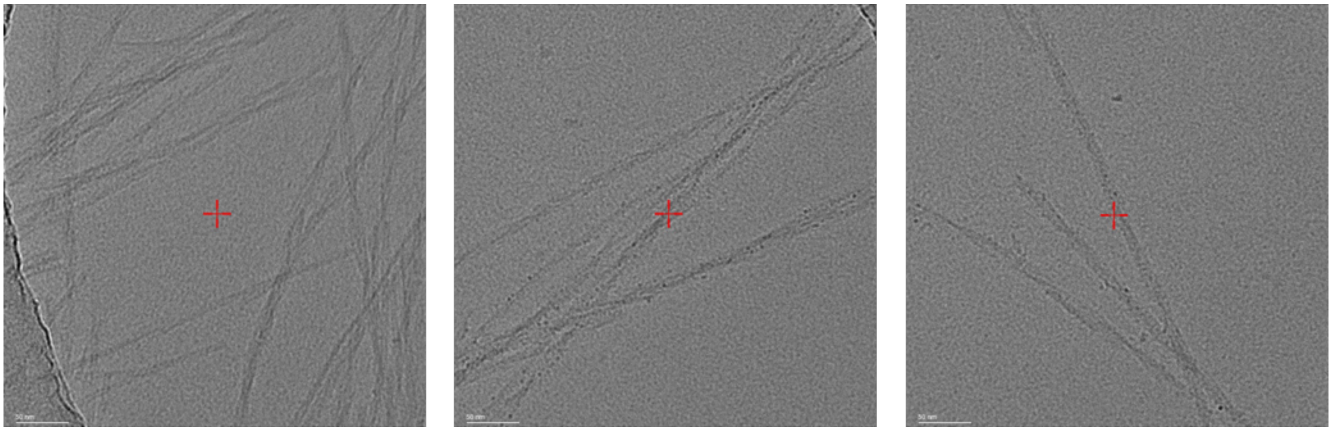

**Cryo-EM micrographs of pS400 (5 weeks).**

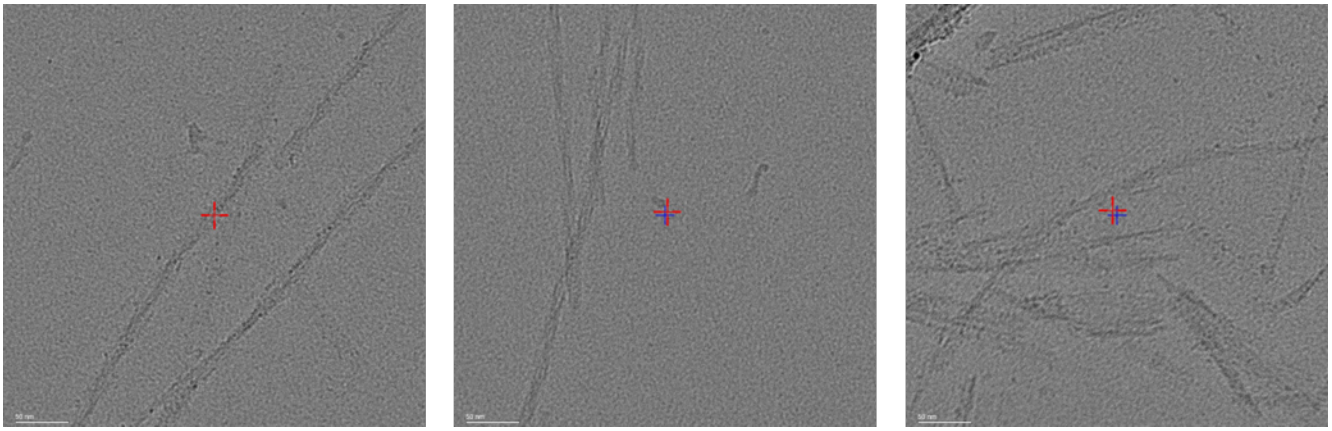

**S8b.** Cryo-EM data collection and refinement statistics.

| | **pS396+pT403** | **pS400-1** | **pS400-2** |
|---|---|---|---|
| **PDB accession code** | **13GX** | **13GY** | **13GZ** |
| **EMDB accession code** | EMD-77062 | EMD-77063 | EMD-77064 |
| **Data collection** | | | |
| Microscope and camera | Krios G4, Falcon 4i | | |
| Nominal magnification | 165,000x | | |
| Voltage (kV) | 300 | | |
| Data acquisition software | EPU | | |
| Exposure navigation | Image shift | | |
| Electron exposure ($e^{-}/Å^2$) | 45 | | |
| Defocus range (μm) | -0.8 to -1.0 | | |
| Pixel size (Å) | 0.728 | | |
| **Reconstruction** | | | |
| Box size (pixels) | 288 | 288 | 288 |
| Interbox distance (Å) | 14.25 | 14.25 | 14.25 |
| Initial segments extracted (no.) | 26,038 | 122,580 | 103,301 |
| Final segments (no.) | 12,980 | 15,578 | 24,468 |
| Helical rise (Å) | 2.39 | 4.76 | 4.77 |
| Helical twist (°) | 179.47 | -1.40 | -1.36 |
| Map resolution (Å) | 3.6 | 3.7 | 3.6 |
| FSC threshold | 0.143 | 0.143 | 0.143 |
| Map sharpening B factor ($Å^2$) | -91.65 | -97.47 | -86.58 |
| **Modeling** | | | |
| Model resolution (Å) | 3.9 | 3.8 | 3.8 |
| **Model composition** | | | |
| Non-hydrogen atoms | 9,184 | 8,352 | 5,728 |
| Protein residues | 1,200 | 1,104 | 752 |
| **B factors ($Å^2$)** | | | |
| Protein | 65.9 | 54.6 | 69.9 |
| **R.M.S. deviations** | | | |
| Bond lengths (Å) | 0.009 | 0.011 | 0.005 |
| Bond angles (°) | 1.128 | 1.317 | 1.170 |
| MolProbity score | 1.54 | 1.67 | 1.73 |
| Clash score | 4.96 | 6.11 | 4.86 |
| Rotamer outliers (%) | 0 | 0 | 0 |
| **Ramachandran plot** | | | |
| Favored (%) | 95.89 | 95.15 | 92.22 |
| Allowed (%) | 4.11 | 4.85 | 7.78 |
| Outliers (%) | 0 | 0 | 0 |

**S8c.** 2D class averages.

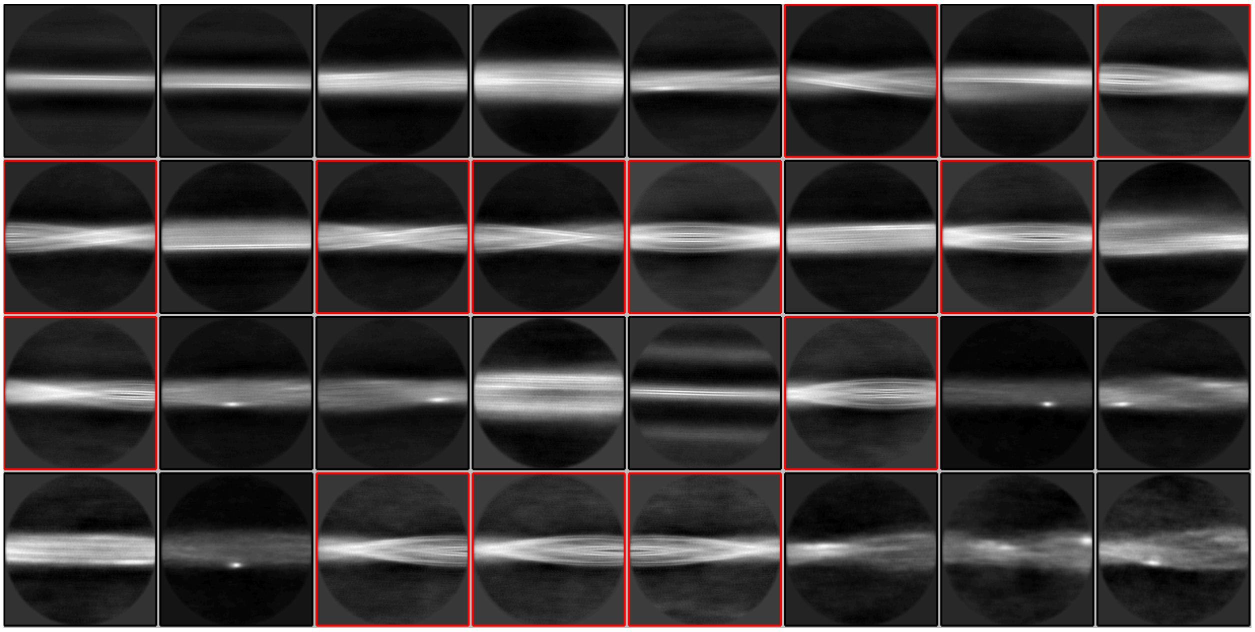

**Supplemental Figure S8c-1.** Initial reference-free 2D class averages of pS396+pT403 dataset (box size 900px binned to 300px). Highlighted classes correspond to twisted filaments used for further processing.

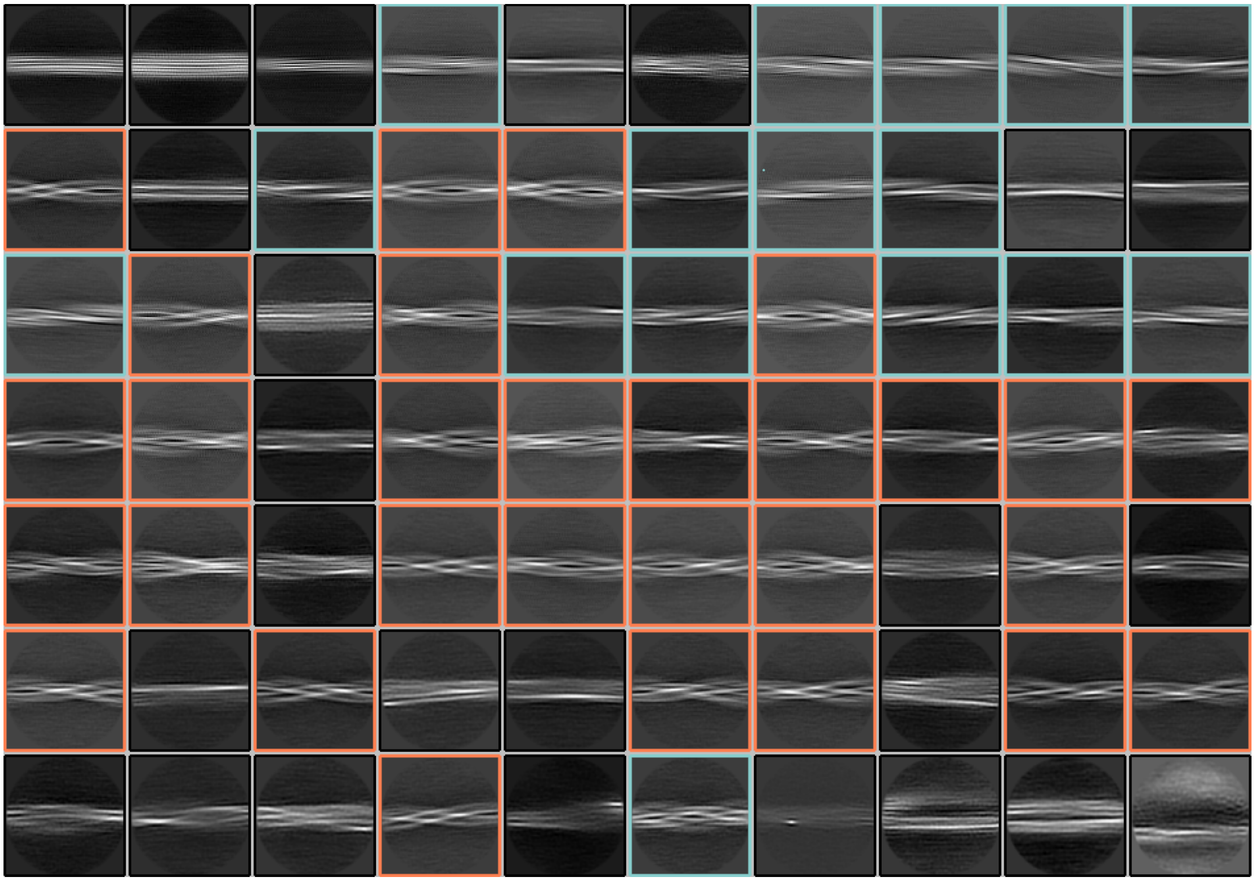

**Supplemental Figure S8c-2.** Initial reference-free 2D class averages of pS400 dataset (box size 900px binned to 300px). Highlighted classes correspond to resolvable polymorphs: orange (polymorph **1**) or light blue (polymorph **2**). The averages were derived from segments extracted from non-CTF-corrected micrographs, which allows easier identification of polymorphs despite the lower resolution.

**S8d.** Model building.

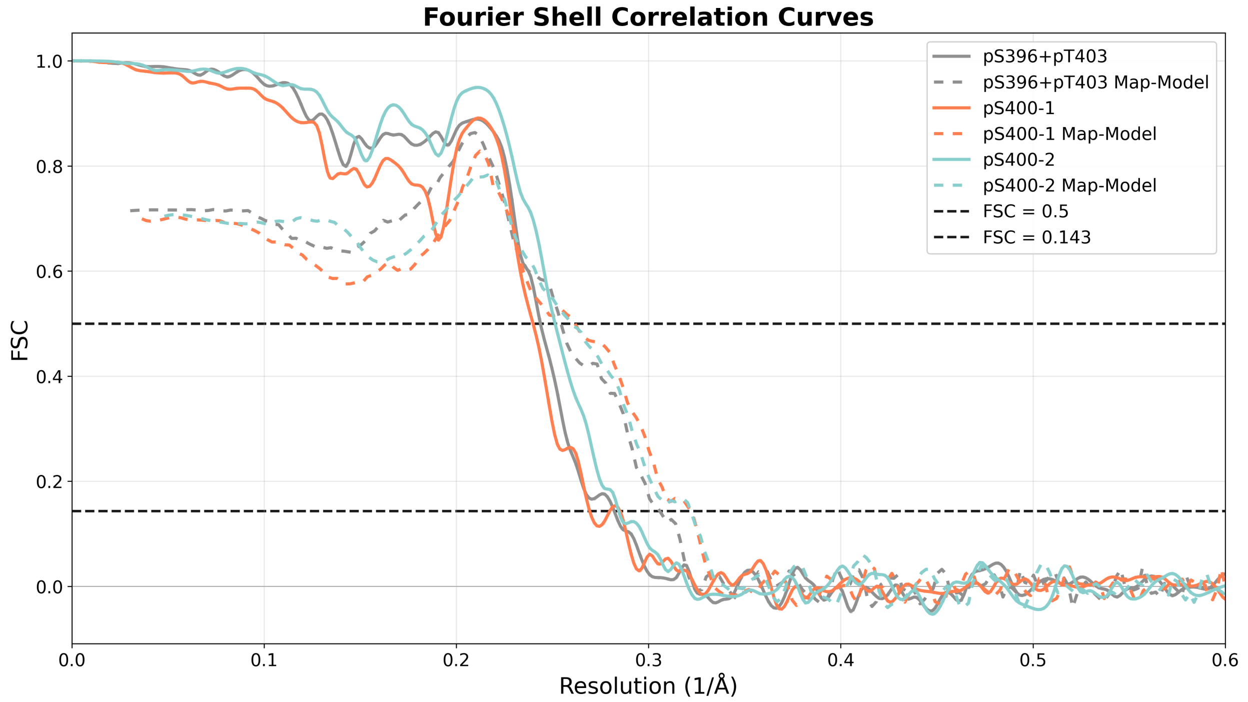

**Supplemental Figure S8d-1.** Fourier Shell Correlation (FSC) curves of post-processed, masked cryo-EM maps (comparing independently refined half-maps) and map-to-model FSC curves.

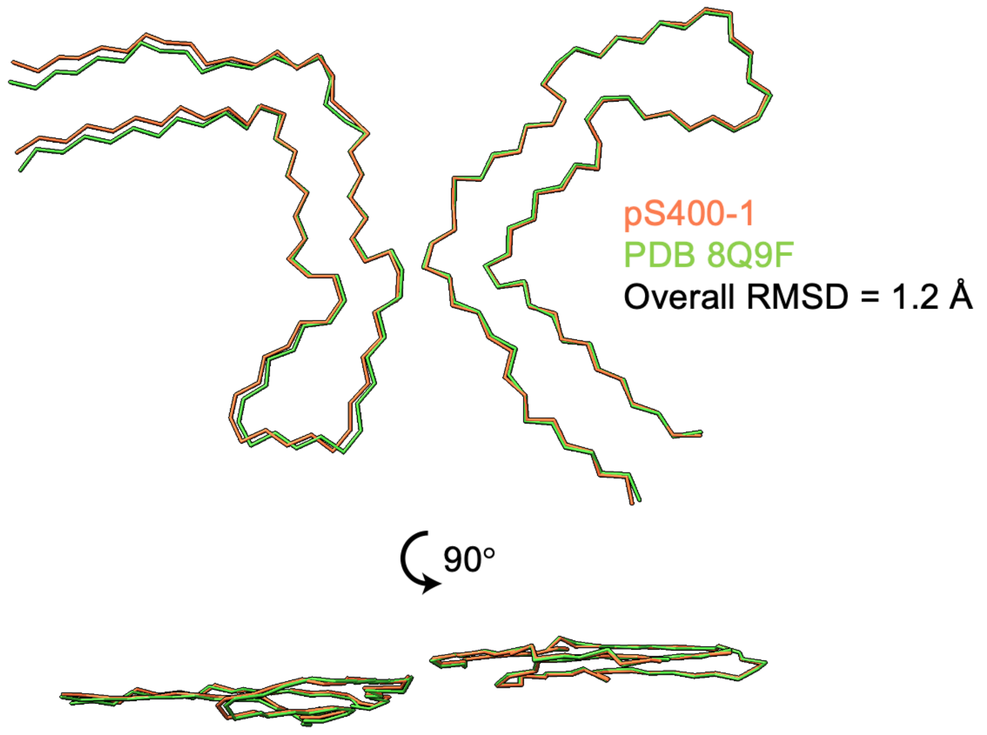

**Supplemental Figure S8d-2.** Comparison between pS400-1 (orange) and CTE LIA-3 (PDB: 8Q9F; green). Both adopt very similar protofilament folds and interfaces. The overall RMSD of pS400-1 to 8Q9F is 1.2 Å.

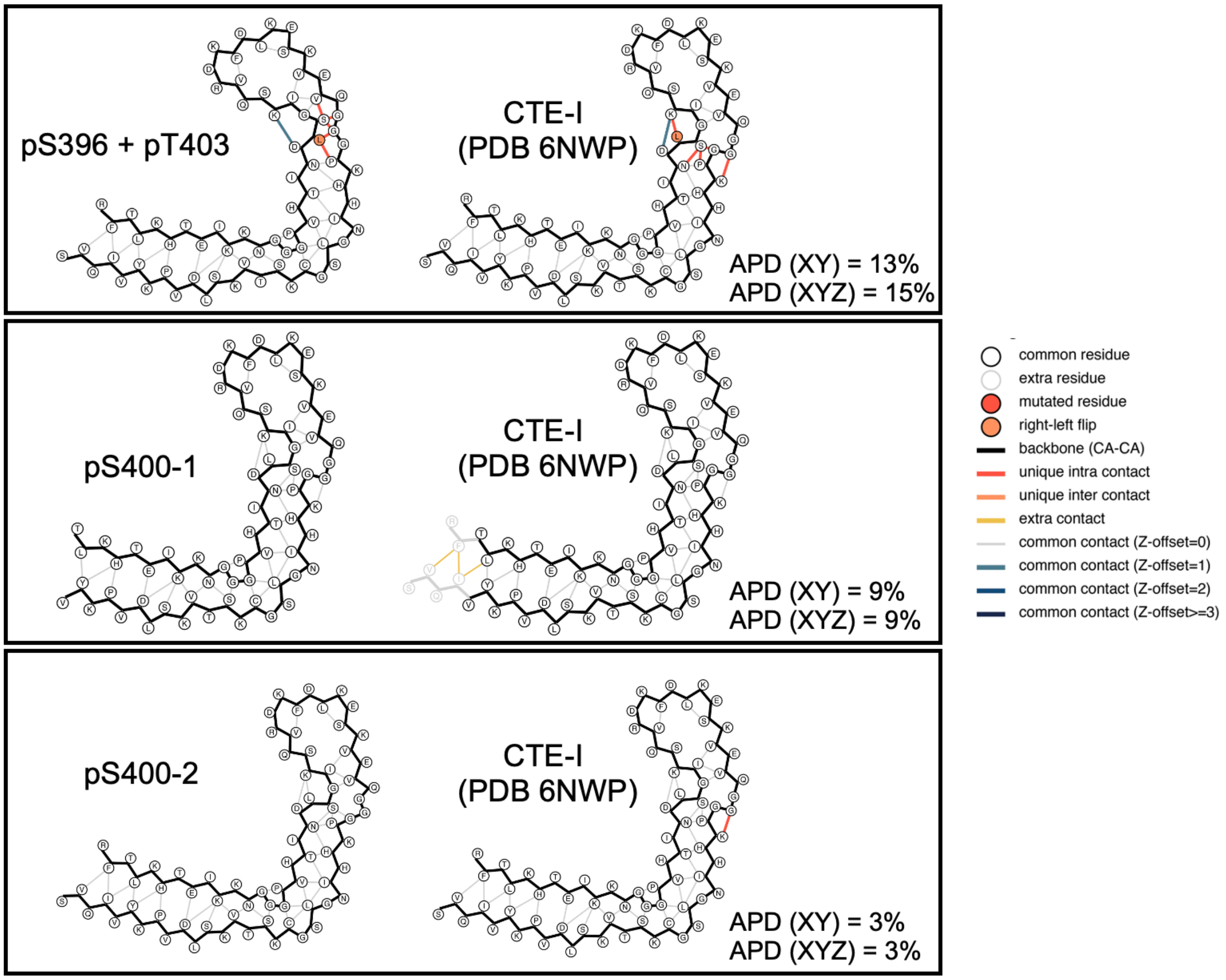

**Supplemental Figure S8d-3.** Amyloid packing differences (APD) between phosphorylated tau filaments (main protofilaments) and CTE protofilaments (PDB 6NWP).

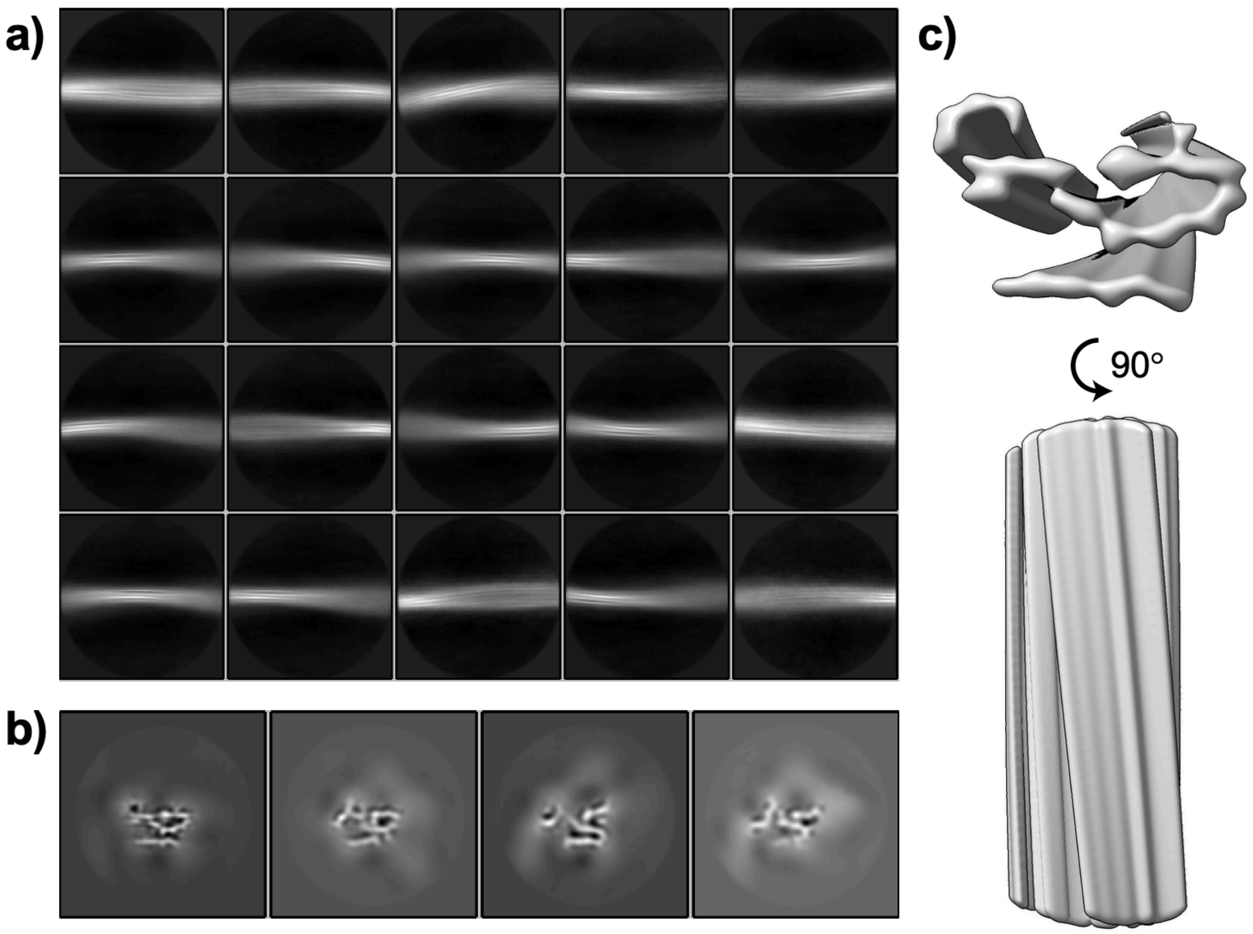

**Supplemental Figure S8d-4.** Attempted cryo-EM structural determination of tau(297-407) pS396+pS400+pT403 filaments. a) Reference-free 2D class averages (900px box binned to 300px). b) 3D classification with AmyBlush c) The highest-resolution 3D class obtained.

### References.
